# Proteomic Profiling of Human Omental and Subcutaneous Adipose Tissue in Individuals with a Broad Range of BMI

**DOI:** 10.64898/2026.01.14.699533

**Authors:** Alex Zelter, Yue Winnie Wen, Michael Riffle, Lindsay C. Czuba, Aprajita S. Yadav, Jerry Zhu, Jessica M Snyder, Aaron Maurais, Jeffrey LaFrance, Saurabh Khandelwal, Judy Y. Chen, Estell Williams, Zoe Parr, Daniel Kim, Katya B. Rubinow, Michael J MacCoss, Nina Isoherranen

## Abstract

Obesity is a major public health challenge affecting an ever-increasing proportion of the global population. It is associated with numerous comorbidities. Progressive expansion and remodeling of adipose tissue may lead to depot specific changes in adipose tissue biology and energy partitioning. Such changes likely precede the development of obesity-related complications. To facilitate a deeper understanding of adipose tissue biology, a comprehensive and quantitative proteomic dataset at the peptide and protein level is presented. Data-independent acquisition LC-MS/MS data were acquired from matched subcutaneous and omental adipose tissues from metabolically healthy individuals with no comorbidities and covering a wide range of body mass indexes. Adipose tissue samples were collected during elective surgeries and immediately processed for histology or frozen until proteomic analysis. Internal and external quality control systems ensured high quality data. All data presented are available via ProteomeXchange. This dataset will allow new insights into biological changes that evolve with increasing adiposity captured before the onset of comorbidities. Matched sampling across fat depots provides an opportunity to uncover depot-specific physiological signatures.

## Background and Summary

Obesity is a significant public health problem with increasing prevalence. It is estimated that 42% of the US adult population has obesity and 10% has severe (class III) obesity.^1^ Obesity is characterized by adipose tissue expansion, increased adipogenesis, and increase in adipocyte size due to accumulation of fat in the cells.^2^ Progressive obesity is associated with metabolic impairment and development of comorbidities such as diabetes, cardiometabolic syndrome and metabolic dysfunction-associated fatty liver disease (MAFLD) that result in morbidity.^2,3^ The progression of metabolic impairment associated with obesity varies considerably between individuals. The metabolic derangements in adipose tissue that precede the development of obesity-related comorbidities are not well defined.

Adipose tissue is a key endocrine organ that serves important functions of energy regulation and thermogenesis. Adipose tissue is, however, not a single homogenous organ but a heterogeneous assembly of different depots that play different roles in endocrine regulation.^4^ In the context of obesity, the subcutaneous and visceral adipose tissue depots are of particular interest. Progressive obesity results in differential expansion of subcutaneous and visceral adipose tissue with substantial interindividual differences in the adipose tissue depots. Metabolic impairment that occurs with obesity is suggested to be associated with increased visceral adipose tissue while subcutaneous adipose tissue may be metabolically protective in the setting of obesity.^4^ Adipose tissue depot-specific data are lacking in human obesity, and the protein signatures of adipose tissue expansion have not been defined in clinical samples. Analysis of adipose tissue from individuals with wide range of adiposity may lend insight into potentially protective and maladaptive pathways in the subcutaneous and visceral depots.

Several studies have characterized adipose tissue proteomes in humans to determine how the adipose tissue proteome may differ between tissue depots or between individuals with and without obesity or with and without type 2 diabetes.^5–8^ However, most prior studies focused on obese patients only^5–7^ or specifically on individuals with type 2 diabetes.^8^ None of the prior studies included men and women across a broad range of adiposity and simultaneously analyzed adipose tissue from multiple depots. In most studies, comorbidities such as type 2 diabetes were not excluded, and hence the proteomic differences observed in these datasets may reflect changes driven by other comorbidities rather than obesity alone.

The differences in the proteomes of adipocytes isolated from omental visceral and subcutaneous tissue from women with obesity who underwent bariatric surgery have been previously reported.^5,6^ Both studies identified enriched protein networks within the two depots. These two studies implicate depot difference in adipose tissue in women with obesity, but they do not show how comorbidities present in most of the participants impact the adipose tissue proteomes. This is important as presence of diabetes or exercise training has been shown to impact the adipose tissue proteome.^7,8^ Proteomic, data dependent acquisition (DDA) based analysis of mouse abdominal visceral and subcutaneous adipose tissues has shown profound changes in the adipose tissue proteome with diet-induced obesity compared with chow-fed mice.^9^ In mice, intermittent fasting was also found to have depot specific effects in adipose tissue proteome when evaluated by DDA and followed by targeted quantification of beta-adrenergic receptors.^10^ Whether these findings in mice translate to humans is not known. The current study provides a data independent acquisition (DIA) based quantitative proteomic analysis of adipose tissue from subcutaneous and visceral depots in men and women along a continuum of adiposity but in the absence of obesity-related comorbidities.

In the current work, we performed proteomic analysis of paired subcutaneous and omental adipose tissue samples from male and female participants with a broad range of adiposity but without significant obesity related comorbidities such as diabetes, kidney or liver disease, or cardiovascular disease. Body mass index (BMI) was used as a surrogate of adiposity. The data were collected using data independent analysis (DIA)^11–13^ which enables untargeted and quantitative assessment of the adipose tissue proteome. DIA enables the quantitation of extracted precursor > product ion chromatograms (XICs) for all detected peptides. These XICs facilitate the quantitative analysis of peptides detected without a list of pre-specified protein or peptide targets. In the current study, we performed DIA analysis on all samples, allowing the detection and differential analysis of all peptides in the sampled m/z range (400-1000 m/z).

These systematically collected mass spectrometry data provide a powerful resource for investigating the biology of human obesity. Critically, the paired nature of the dataset enables direct comparisons between adipose tissue depots, offering insights into depot-specific biology across the spectrum of human obesity.

## Methods

### Ethics Oversight, Study Participants and Human Tissue Samples

Men and women ages 18-65 years old who were planning elective abdominal surgical procedures with access to visceral fat and liver were recruited through University of Washington Medical Center surgery clinics. The study was conducted in accordance with the Declaration of Helsinki principles and approved by the UW Institutional Review Board (ID: CR00006134, study ID: STUDY00005135). All participants signed written informed consent and provided HIPAA authorization prior to any study procedures. Enrollment inclusion criteria included a body-mass index (BMI) of ≤27.5 kg/m^2^ or ≥30 kg/m^2^. Some participants undergoing metabolic surgery were required to adhere to a very low-calorie diet over 3-6 weeks prior to surgery and lost weight before surgery. These participants are indicated in the associated metadata. For any participant with an underlying health condition that could increase risk of liver disease, laboratory data were required within 6 months of pre-screening. For participants with a BMI <30kg/m^2^ and without risk factors for underlying liver disease, laboratory tests were not required unless requested by the surgical team.

Exclusion criteria included a history of endocrine/secondary causes of obesity, diabetes, current use (within 3 months) of any weight loss, lipid- or glucose-lowering medication, major systemic illness or infection, significant heart, kidney or liver disease, exposure to any form of cancer chemotherapy (intravenous or oral) or radiation therapy (within 2 months), daily vitamin A supplementation ≥10,000 international units, use of anti-platelet or anti-coagulation therapy, malabsorptive gastroenterological disease, anemia, or bleeding disorders. Alcohol use >140 gram for men or >70 gram for women per week was also exclusionary. For participants with a BMI ≥30 kg/m^2^, the non-alcoholic fatty liver disease fibrosis score (NAFLD-FS) was used to estimate risk of the presence of significant fibrosis due to metabolic dysfunction-associated steatohepatitis, and a score >0.675 was exclusionary.^14^

Demographic information for the 31 participants, including age, sex and body mass index (BMI), is listed in Table 1. Race and ethnicity were collected from electronic medical records. Other clinical characteristics are summarized in Table 2.

**Table 1:** Demographic information for study participants and surgical procedure characteristics.

|  |  |
| --- | --- |
| Number of participants | n=31 (20 female, 11 male) |
|  | **Mean ±SD** |
| Age (yr) | 42.8 ± 10.2 |
| BMI (kg/m<sup>2</sup>) | 38.2 ± 9.7 |
| Self-reported Race/Ethnicity | **n (%)** |
| White; Not Hispanic or Latino | 19 (61%) |
| White; Hispanic or Latino | 6 (19%) |
| Black or African American; Not Hispanic or Latino | 5 (16%) |
| Asian; Unknown | 1 (3%) |
| Surgical procedure | **n (%)** |
| Metabolic surgery | 20 (65%) |
| Cholecystectomy | 4 (13%) |
| Hernia repair | 3 (10%) |
| Inguinal hernia repair | 3 (10%) |
| Umbilical hernia repair | 1 (3%) |

**Table 2.** Clinical characteristics for study participants.

|  | **Median (IQR: 25%; 75%)** | **n** |
| --- | --- | --- |
| **Serum Albumin (g/dL)** | 4.0 (4.0; 4.0) | 27 |
| **Serum AST (U/L)** | 18.0 (15.0; 23.0) | 27 |
| **Serum ALT (U/L)** | 21.0 (15.0; 26.0) | 27 |
| **Estimated glomerular filtration rate (eGFR; mL/min/1.73 m<sup>2</sup>)** | 113.6 (100.0; 117.6) | 31 |
| **Hematocrit (%)** | 43.0 (40.0; 45.0) | 29 |
| **Platelets (per μL)** | 271 (219; 307) | 29 |
| **Glucose (ELISA) (mM)** | 4.7 (4.4; 5.0) | 31 |
| **Insulin (mU/L)** | 10.7 (6.5; 19.2) | 31 |
| **Homeostatic Model Assessment of Insulin Resistance (HOMA-IR)** | 2.0 (1.4; 4.1) | 31 |
| **Non-Alcoholic Fatty Liver Disease Fibrosis Score (NAFLD-FS)** | -1.76 (-2.24; -1.26) | 26 |

Two adverse events were reported during the study that were possibly or likely associated with the study. One participant experienced a hematoma at the surgical incision site where the subcutaneous fat biopsy was obtained but completely resolved within 1-week post-surgery. One participant had a superficial wound dehiscence at the surgical incision that was unlikely to be study related but was treated by the surgical team and subsequently resolved.

### Sample Acquisition and Post-excision Adipose Tissue Processing

All participants consumed no food for at least 8 hours prior to transfer to the operating room. Some participants were instructed by the surgical team to consume 8 oz of apple juice ≥2 hours prior to hospital arrival (≥3 hours prior to operating room transfer). Subcutaneous (n=31) and visceral (from omentum) (n=28) adipose tissue biopsies were collected during the elective surgery. For three participants with a BMI <30 kg/m^2^, visceral adipose tissue could not be collected as the indicated surgical procedure did not provide access to the omental depot for biopsy. Biopsies of omental and subcutaneous adipose tissues were performed by surgeons during the elective surgical procedures, with tissue yields of 2-23 grams from each depot. Upon acquisition, tissue was immediately rinsed with ice-cold phosphate-buffered saline. From biopsies from each depot, aliquots of adipose tissue were flash-frozen on dry ice and stored at -80°C for whole-tissue analysis or fixed in formalin for histological analyses. A flowchart of the adipose tissue processing from the biopsy samples is shown in Fig.1. Rinsing and freezing or fixing of the adipose tissue samples by research staff was completed within 5 minutes of surgical excision from the patient.

**Figure 1.**
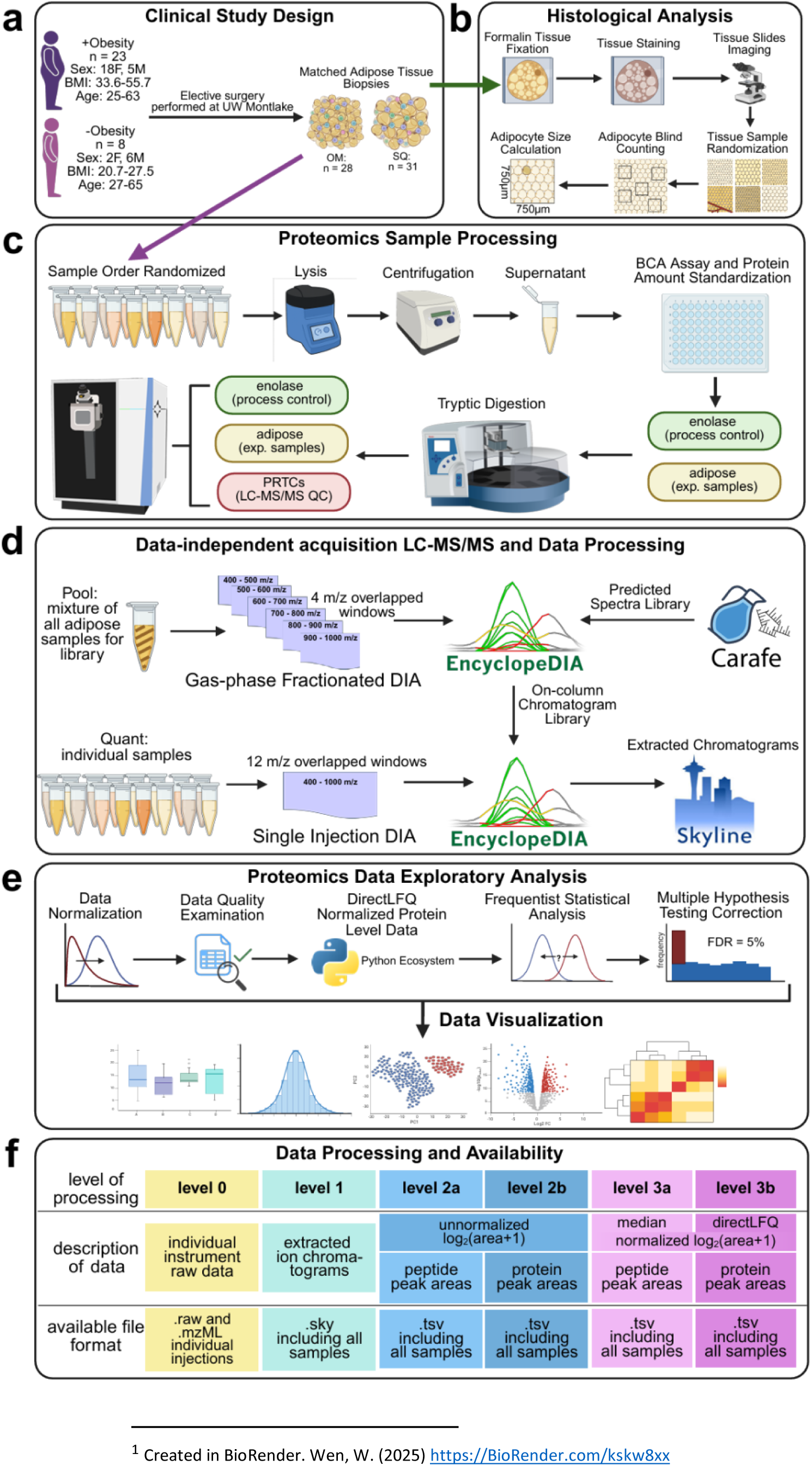
Clinical study design, experimental scheme, data acquisition and processing workflow. (a) Matched omental (OM) and subcutaneous (SQ) adipose tissues were collected during elective surgery from participants with and without obesity. (b) Histological analysis was performed to estimate the adipocyte size in each adipose tissue depot. (c) Samples were randomized and analyzed in a single batch of 58. Yeast enolase was added as a process control to each adipose protein lysate before digestion. After digestion, Pierce PRTC synthetic peptide mixture was added as a system suitability control. (d) An on-column chromatogram library was built from six gas-phase fractionated LC-MS/MS injections using narrow-window DIA (4 m/z overlapping windows) of a pooled mixture of all individual samples. Single injections of each individual sample were acquired for quantitative analysis using wide-window (12 m/z overlapping windows) DIA. Peptide detection and scoring was done using EncyclopeDIA and extracted ion chromatograms produced and integrated with Skyline. (e) Peptide and protein peak areas were normalized using directLFQ prior to data analysis and visualization using python. (f) The proteomics data is publicly available on the Panorama web server at 5 different levels of processing^1^.

### Slide Preparation and Adipocyte Analysis

Formalin-fixed samples were embedded in paraffin blocks, sectioned onto slides, and routinely hematoxylin and eosin (HE) stained by the Pathology Research Services Laboratory in the University of Washington Department of Pathology. Anti-CD45 immunohistochemistry was performed at the University of Washington Histology and Imaging Core using mouse anti-human CD45 antibody, clone PD7/26+2B11 (Abcam, Waltham, MA, #ab781) at 1:100 using the Bond automated immunostainer and the Bond Polymer Refine (DAB) Detection Kit (Leica Microsystems; Deer Park, IL). HE and anti-CD45 stained slides were then scanned in bright-field with a 20× objective using a Nanozoomer Digital Pathology slide scanner (Hamamatsu; Bridgewater, NJ) and the scanned anti-CD45 stained slides were used for adipocyte analysis. Representative HE images are shown for each group and depot (Fig. 2).

**Figure 2.**
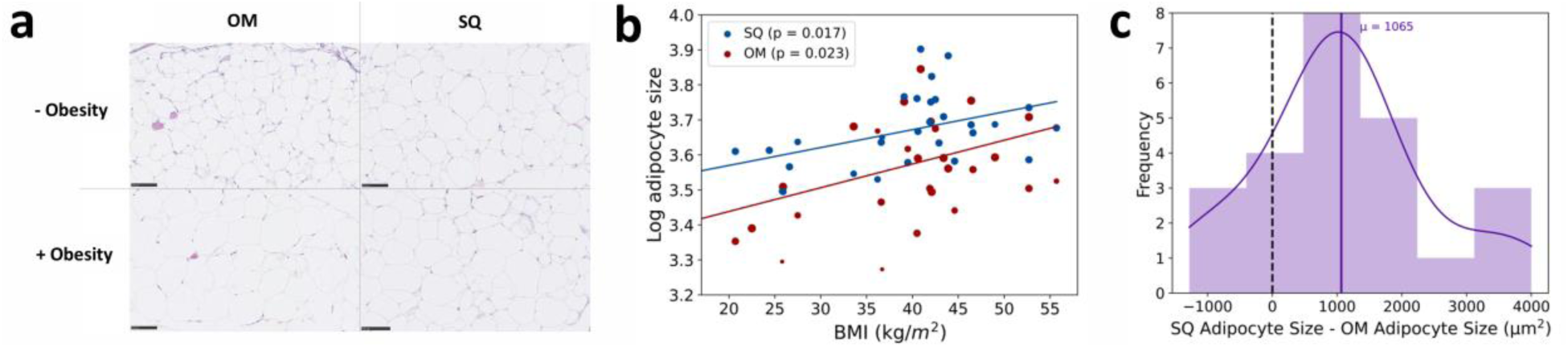
Adipose tissue histology suggests that omental (OM) and subcutaneous (SQ) adipocyte size correlates with BMI and that SQ adipocytes are bigger in size than OM. (a) Representative histology images of adipocytes. (b) Weighted linear regression with log adipocyte size plotted against BMI (SQ = 31 and OM = 28). The data weight/dot size was calculated based on the number of squares available for adipocyte counting out of all the squares drawn. (c) Histogram for the size difference between SQ and OM adipocytes with the kernel density estimation. On average, the SQ adipocytes are 1,065 µm^2^ greater than the OM adipocytes (Wilcoxon signed-rank test: *p-value* < 0.05). The estimated mean adipocyte size in each individual sample is included in the associated metadata for the adipose tissue biopsies analyzed.

All the slides across study participants (n = 31) and adipose tissue types (omental = 28; subcutaneous = 31) were analyzed in a blinded manner where the analyzer (JZ) was blinded from the identity of the slide, the adipose tissue depot and participant BMI. Up to five squares (750 µm x 750 µm) were randomly drawn for each slide imaged for every study participant and each depot. The number of adipocytes were counted for each square. Adipocytes within the squared regions were counted if at least half of the adipocyte was within the area of the square. The average number of adipocytes per square in each study participant and adipose tissue depot was calculated. The arithmetic mean number of adipocytes per square was calculated for all the squares analyzed in each omental and subcutaneous sample. A second investigator (AY) reviewed all squares and counting. The size of adipocytes was calculated by finding the area for the square drawn (0.5625 mm^2^) and then dividing the area by the number of adipocytes counted. The arithmetic mean of adipocyte size from all the squares counted per imaged slide was reported, which assumes that there is no space between each cell.

### Sample Preparation

Frozen adipose tissue samples were weighed, and approximately 100 mg of frozen tissue was placed in a 1.5 mL Eppendorf Safe-Lock tube (cat no. 022363212) on ice. A 3:1 ratio of chilled lysis buffer to tissue weight was added to each tube. The lysis buffer was 100 mM tris pH 8.5 containing 1x EDTA-free HALT protease inhibitor cocktail (sold as 100X, Invitrogen cat no. 78425). Approximately 10 beads (Stainless Steel Beads 0.9 – 2.0 mm blend, Next Advance, Inc. cat no. SSB14B) were added to each Eppendorf tube and the tissues were lysed at max power in a Next Advance Bullet Blender Tissue Homogenizer in a 4°C cold room for 4 cycles of 5 minutes lysing and 1 minute rest between each cycle. The resulting lysate, complete with beads, was centrifuged at 9,000 g at 4°C for 30 minutes in a benchtop centrifuge. After centrifugation, samples consisted of 3 layers: the pellet, a middle liquid layer and a lipid layer on top. From the middle layer, 150 µL of lysate was moved to a fresh 1.5 mL Eppendorf LoBind tube (cat no. 022431081). This solution was brought up to 2.5% sodium dodecyl sulfate (SDS) by addition of 21.3 µL of 20% SDS solution (Invitrogen cat no. AM9820). A BCA assay was performed on all samples using Invitrogen’s Pierce BCA Protein Assay Kit according to the manufacturer’s instructions (cat no. 23227).

### Protein Digestion

Samples were prepared for mass spectrometry analysis using protein aggregation capture (PAC).^15,16^ For each sample a variable volume of diluent (100 mM tris pH 8.5, 2.5% SDS) and adipose lysate corresponding to 35 µg total protein was added to a 96 well, deep well-plate (Thermo Fisher Scientific catalogue no. 95040450). Diluent was added first, followed by adipose lysate. The final volume per well was 120 µL. In three cases, lysate concentration was insufficient for addition of 35 µg. In these cases, the maximum lysate volume of 120 µL was added corresponding to the following protein amounts: 51-3505-SQ-2-obese (34 µg); 51-3502-SQ-2-obese (27 µg) and 51-3504-SQ-4-lean (11 µg). Data from 51-3504-SQ-4-lean was included in all analyses except Fig. 7c (the hierarchical clustering panel) where it was removed for visual clarity.

To each well of the 96-well plate, 15 µL of enolase (Sigma, cat no. E6126) was added as a process control,^17^ along with a tris(2-carboxyethyl)phosphine (TCEP) solution (Invitrogen, Bond-Breaker TCEP Solution, Neutral pH, cat no. 77720) in 100 mM Tris buffer at pH 8.5. This brought the final concentrations to 560 ng of enolase and 10 mM TCEP per well. A lid was placed on the 96-well plate (Corning, cat no. 3099) and samples were mixed in the plate by shaking at 1,200 rpm for 30 seconds using an Eppendorf ThermoMixer followed by reduction for 1 hour at 37°C in the ThermoMixer with shaking at 650 rpm. To each well, 15 µL of iodoacetamide (Sigma, cat no. I1149) was added to bring the final concentration to 10 mM. Alkylation was performed at room temperature in the dark for 30 minutes. To each well, 12.5 µL of MagResyn Hydroxyl particles (ReSyn Biosciences, cat no. MR-HYX002) were added. Protein was aggregated to the beads by addition of 290 µL of 100% ethanol followed by 547.5 µL MTBE (methyl-tert-butylether, Sigma, cat no. 34875) bringing the final volume per well to 1 mL. The beads were cycled through 5 KingFisher deep well wash plates containing 1 mL of wash solution per well (3 plates containing 95% acetonitrile followed by 2 plates containing 70% ethanol) with each wash step taking 2.5 minutes in a KingFisher Apex (Thermofisher Scientific). Protein-containing beads were released into 150 µL 50 mM tris, pH 8.5 containing 1.75 µg trypsin (Pierce, cat no. PI90058) per well for a 1:20 trypsin to substrate ratio. Digestion was performed at 47°C for 1 hour in the KingFisher. After digestion, 120 µL per sample was added to a 1.5 mL LoBind Eppendorf tube containing 7 µL 10% trifluoroacetic acid (TFA). A further 12 µL of 500 fmol/µL Pierce Peptide Retention Time Calibration Mixture (PRTC) (Thermo Scientific, cat no. PI-88321) dissolved in 0.1% TFA was added as a process control.^17^

From each adipose digest, 15 µL of peptides + PRTCs were pooled into a single Eppendorf LoBind tube for generation of a chromatogram library^4^ as described below.

All peptide samples were centrifuged in a benchtop centrifuge at 4°C at max speed for 10 minutes and supernatants were transferred to autosampler vials and stored at -80°C until analysis.

### Mass Spectrometry

Prepared samples were analyzed by data independent acquisition-mass spectrometry (DIA-MS). Each sample contained 2 µg total adipose protein plus 32 ng yeast enolase and 450 fmol of heavy labelled Pierce PRTC Mixture. Enolase and PRTCs were added to all samples as process controls.^17^ Using a Thermo Vanquish Neo UHPLC System, 9 µL of sample was analyzed using “trap and elute” with a Thermo Scientific PepMap Neo Trap Cartridge (cat no. 174500) and a Bruker PepSep C18 15 cm x 150 µm, 1.9 µm column (cat no. 1893471). Separate System Suitability injections were performed by injecting 6 µL of a mix containing 300 fmol PRTCs and 1,200 fmol BSA in 0.1% TFA.^17^ The output of the HPLC column was connected to a 5 cm x 20 µm ID Sharp Singularity Fossil Ion Tech tapered tip mounted in a custom constructed microspray source heated to 45°C. Peptides were eluted from the column at 0.8 µL/min using the following acetonitrile gradient: (1) 0-0.7 mins; 3-5% B; flow 1.3 µL/min; (2) 0.7-1 mins; 5-5.5% B; flow 1.3 µL/min; (3) 1-57 mins; 5.5-40% B; flow 0.8 µL/min; (4) 57-57.5 mins; 40-55% B; flow 1.3 µL/min; (5) 57.5-58 mins; 55-99% B; flow 1.3 µL/min; (6) 58-60 mins; 99% B; flow 1.3 µL/min. Buffer A was 0.1% formic acid in water. Buffer B was 0.1% formic acid, 80% acetonitrile and 20% water.

Mass spectrometry data were collected using a Thermo Fisher Scientific Orbitrap Exploris 480. DIA data was collected on each individual sample using a method with 12 m/z staggered precursor isolation windows.^18,19^

Data on individual samples were collected using the following parameters. The Orbitrap resolving power for MS1 and MS/MS were 30,000 at *m/z* 200. Each cycle consisted of 1 MS scan followed by 50 MS/MS scans. MS scan range was *m/z* 395–1,005. Automatic gain control was set to standard for MS scans and to a normalized value of 1000% for MS/MS scans. MS/MS spectra were acquired using overlapping isolation windows of 12 *m/z* with 50 MS/MS scans per cycle and a normalized collision energy of 27 assuming charge state 3. Cycle 1 spanned 406.4347 *m/z* through 994.7021 *m/z* in 12 *m/z* isolation windows. Cycle 2 spanned 400.4319 *m/z* through 1000.7048 *m/z* in 12 *m/z* windows. All spectra were collected in centroid mode.

A chromatogram library was created to improve the analysis of the 12 m/z DIA data collected on the individual samples. To create this library, gas phase fractionation on a pooled adipose tissue digest was performed.^20,21^ The chromatogram library was collected by performing six LC-MS/MS runs each covering a subset of the total m/z range measured by the individual sample analyses. Each of the six LC-MS/MS runs were performed using scan ranges of 395-505, 495-605, 595-705, 695-805, 795-905 and 895-1005 *m/z*. These data were acquired using 4 *m/z* staggered windows. Other settings, particularly the HPLC separation conditions, remained the same as the analyses performed on the individual samples using 12 m/z precursor isolation windows.

### Data Processing and Normalization

Acquired spectra were demultiplexed^18,19^ and converted to mzML format using ProteoWizard’s msConvert^22^ version 3.0.24172 using the following arguments: --mzML --zlib --ignoreUnknownInstrumentError --filter “peakPicking true 1-” --64 --filter “demultiplex optimization=overlap_only” –simAsSpectra. An instrument-specific, optimized, spectral library in dlib format was produced using a FASTA file containing the UniProt reviewed human proteome (UP000005640) plus common contaminants (https://www.thegpm.org/crap/) and yeast enolase (P00924) using DIA-NN^23^ version 1.8.1 and Carafe version 0.0.1 as previously described.^24^ This step was performed using an automated Nextflow^25^ workflow, nf-carafe-ai-ms version 85ee6c5, available at: https://nf-carafe-ai-ms.readthedocs.io/en/latest/. A second Nextflow workflow, nf-skyline-dia-ms version eaf4febe75, (available at: https://nf-teirex-dia.readthedocs.io/en/latest/) was used to generate an on-column chromatogram library using the narrow-window DIA data followed by quantification of the wide-window data using EncyclopeDIA version 2.12.30 as previously described.^20^ The final set of transitions were quantified using Skyline.^26,27^

Skyline documents were made from the data corresponding to the complete sample set. Protein grouping was performed by Skyline (as described here: https://skyline.ms/wiki/home/software/Skyline/page.view?name=Skyline%20Protein%20Association%2022.1) using logic based on the IDPicker algorithm whereby proteins that match the same set of peptides are merged into indistinguishable protein groups.^28,29^ Two Skyline documents were made from the same complete dataset. These documents differed only by different treatment of peptides shared between multiple proteins or protein groups: For the first Skyline document, peptides mapping to more than one protein were kept (documents tagged “keep_dups”). For the second document peptides mapping to more than one protein were removed (tagged “throw_dups”). The figures and analysis presented in this publication were done using the keep_dups Skyline file. The throw_dups file is shared as a convenience for cases where proteins of interest have peptides common to more than one protein but is not otherwise referenced in the current work. Both Skyline files are available on Panorama Public^30^ as described in the Data Records section, below.

Peptide and protein level normalization was performed on the two Skyline documents corresponding to the complete sample set using a third Nextflow workflow (available at: https://github.com/uw-maccosslab/nf-dia-batch-correction). This normalization was designed to compensate for unavoidable systematic biases introduced during sample processing and data collection. The workflow applied Median Deviation (MD) normalization at the precursor (peptide) level^31^ and directLFQ^32^ normalization at the protein level. Normalization was performed across the entire sample set and both unnormalized and normalized precursor and protein values were output as tsv files, which were used in downstream analyses. Unnormalized and normalized files are available for both Skyline documents (Fig. 1f, level 2a through 3b data) and are provided as log_2_(area+1), where area is the sum of the total area of all MS/MS transitions for a given peptide (peptide level) or all measured peptides in a given protein (protein level).

### Statistical Analysis

Exploratory statistical analysis and visualization were conducted in Python (v.3.11.9). Numeric and tabular data transformations were performed through NumPy and Pandas modules.^33,34^ Statistical analyses like weighted linear regression, principal component analysis (PCA), Wilcoxon signed-rank test, hierarchical clustering analysis and Spearman’s rank correlations were conducted with SciPy, Scikit-Learn, and Seaborn packages.^35–37^ Data visualization was completed using Matplotlib and Seaborn packages.^37,38^

## Data Records

All data presented in the current work are available at several levels of data processing, from raw data as recorded by the mass spectrometer (level 0) through to normalized peptide and protein level quantities suitable for downstream analysis (levels 3a and 3b, respectively). These levels are highlighted in Fig. 1f and the complete dataset can be accessed via Panorama Public^30^ here: https://panoramaweb.org/human-adipose.url and was assigned the ProteomeXchange^39^ ID: PXD067514 (https://doi.org/10.6069/sq0n-1x32).

**Level 0** represents raw data as recorded by the mass spectrometer, deposited in two formats: Format 1: raw format as output directly mass spectrometer; Format 2: mzML format which is an open, XML-based format for mass spectrometer output files and was additionally demultiplexed during conversion from the original raw files using (Proteowizard version 3.0.24172). **Level 1** represents zipped Skyline documents made from data corresponding to the complete sample set. Two skyline documents were deposited and differed only by use of different protein parsimony options as described in the previous section. **Level 2** represents unnormalized peptide and protein level data and **Level 3** represents normalized peptide and protein level data.

Skyline documents and raw data tracking both system and sample quality are provided in the System Suitability and Process Controls sections of the Panorama Public site. The raw files associated with the process controls are the same as those for the sample data because the process controls were spiked into the samples themselves. The System Suitability raw files are separate as those represent samples specifically designed to test only the performance of the LC-MS/MS system and as such are run separately.

## Technical Validation

### Experimental Design and Internal Controls

The adipose tissue samples analyzed for this dataset were collected by the investigative team during elective surgeries. All samples were flash frozen immediately after collection except for those designated for histological analyses, which were immediately fixed in formalin. This ensured high quality tissue collection and carefully controlled processing. The clinical characteristics of all tissue donors were well defined.

All study samples were randomized prior to proteomic analyses. This randomized order, listed in Supplementary Table S1, was used throughout the sample preparation, digestion and mass spectrometry data acquisition steps.

High quality of the complete dataset was ensured through a system of process controls and system suitability standards, which were added at different stages of sample preparation as previously described (Fig. 1c).^17^ Briefly, all samples included internal process controls consisting of Pierce Retention Time Calibration peptide mixture (PRTC). These controls allow measurement of retention time and signal intensity stability across all chromatographic runs independent of protein digestion efficiency. In addition, whole yeast enolase protein was added to all samples prior to digestion to capture variation in sample workup, digestion, chromatography and MS data acquisition. To this end, five enolase peptides were monitored across all runs. The CV% for these peptide peak areas ranged from 12.3% to 30.4% (Fig. 3a). The CV% of the 15 PRTC peptide peak areas was similar and ranged from 8.4% to 33.2% (Fig. 3a) indicating that sample workup and digestion did not add significantly to run-to-run variation. Sample-independent system suitability injections were also interspersed between the sample injections using a mixture of a tryptic digest of BSA plus PRTCs. These injections were performed every 6 to 18 samples. The CV% of the 15 PRTC peptide areas in these sample-independent injections were similar to the same peptides measured within the samples and ranged from 11.3% to 27.4% (Fig. 3b). These sample-independent controls allow tracking of instrument performance longitudinally over time regardless of sample type.

**Figure 3.**
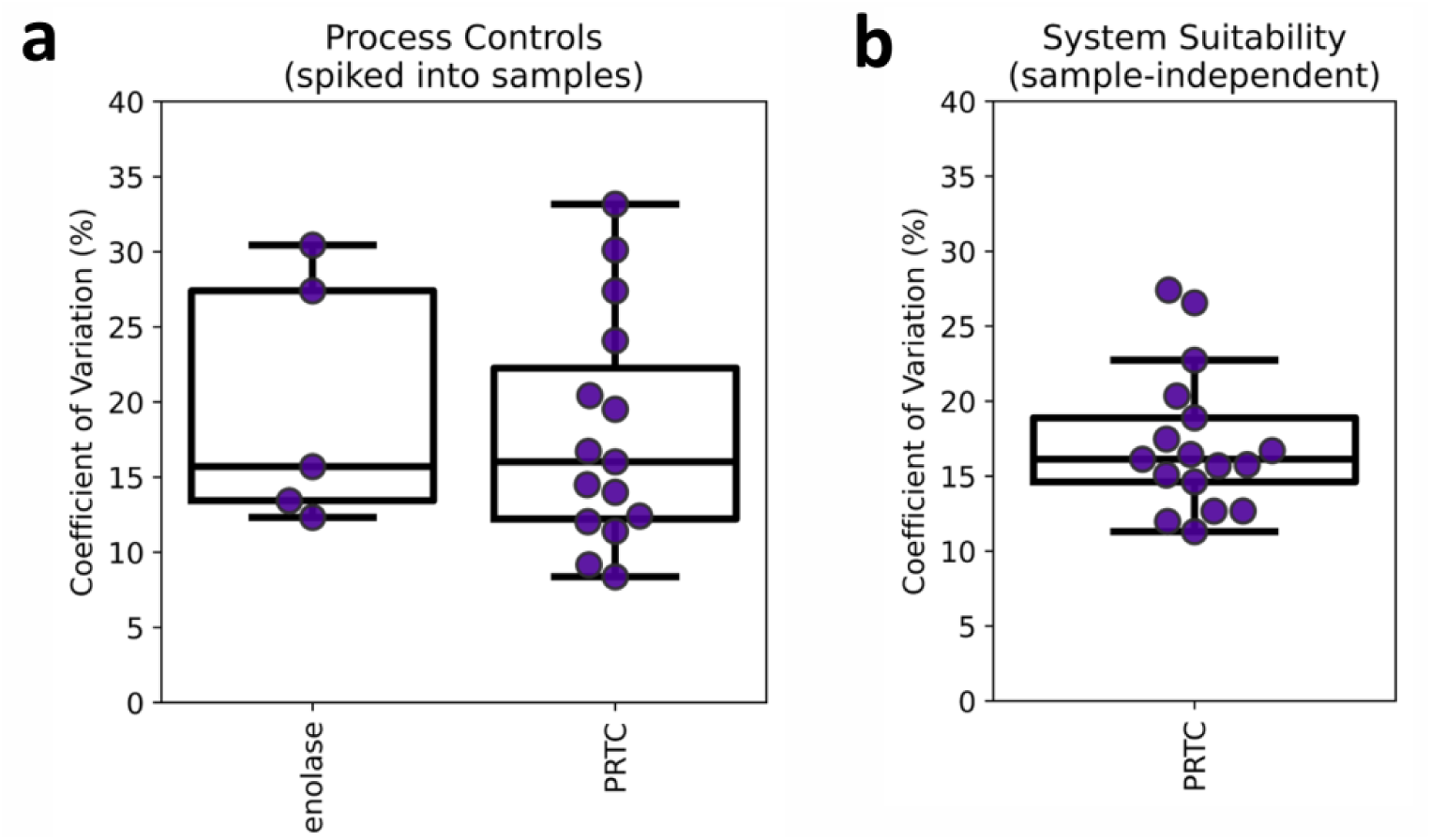
Overview of peptide variance in process control (yeast enolase), internal LC-MS/MS quality control (PRTC), and external system suitability control (PRTC). (a) Box and Whisker plots of coefficient of variation of enolase (added as whole protein) and PRTC peptides spiked into all adipose samples. Each point represents an individual peptide. (b) Coefficient of variation of PRTC peptides measured in sample-independent injections interspersed between the sample injections.

To minimize technical variation and the introduction of confounding factors when comparing different tissue groups, per-sample adipose tissue protein amount was measured and normalized during sample preparation. All samples were prepared, digested and run in the same batch. Despite these precautions, some level of residual technical noise is expected. Computational Median Deviation (MD) normalization^31^ was performed on the peptide total area fragment peak areas to remove this noise. The distribution of peptide peak areas in each sample is shown in Fig. 4 before and after MD normalization. For protein level data, directLFQ normalization was used, which is a ratio-based approach previously shown to have excellent normalization properties when applied to DIA data.^32^

**Figure 4.**
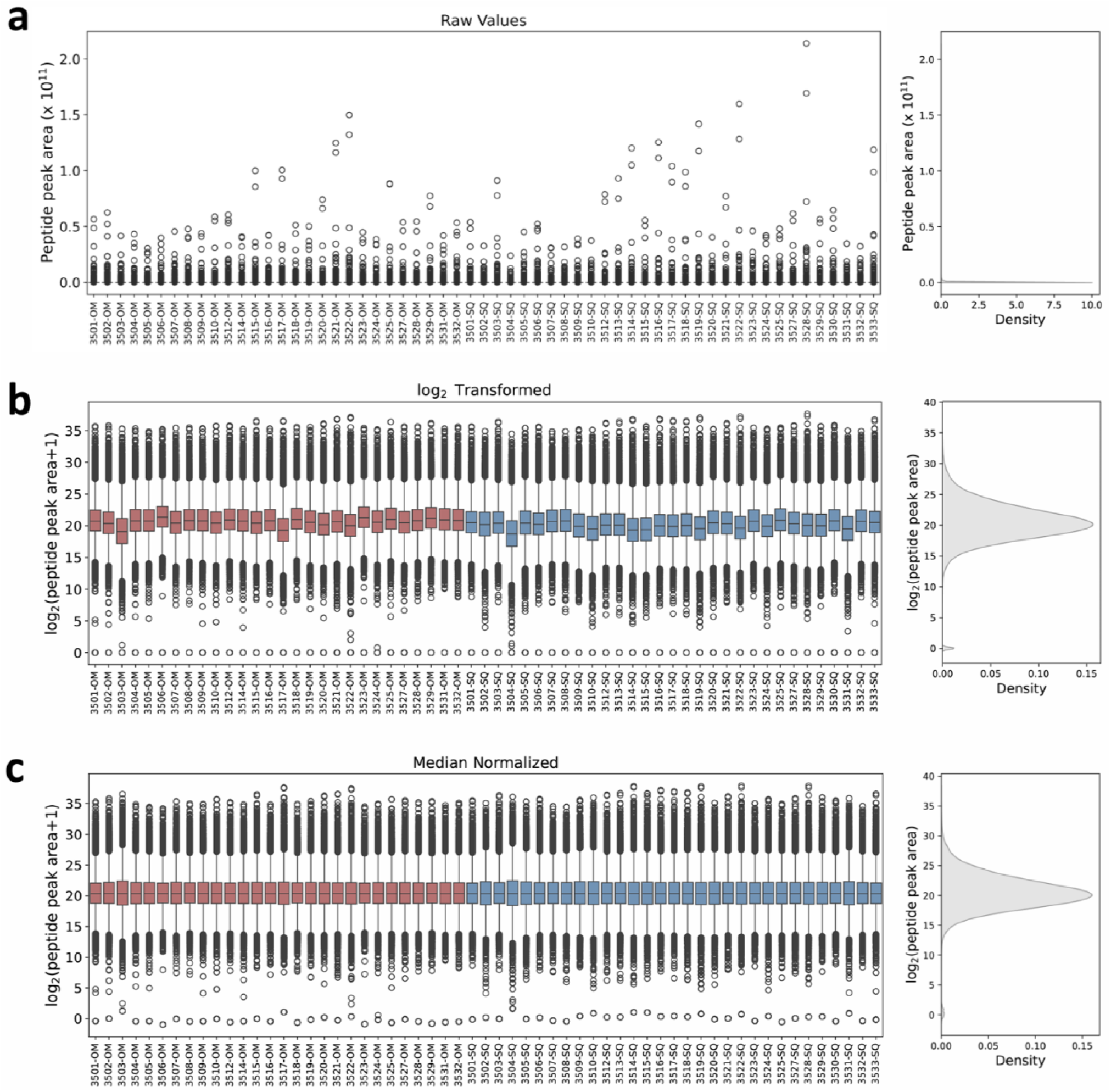
Peptides detected in individual samples. Box and Whisker plots (left) for each individual sample and kernel density plots (right) show the distribution of (a) raw peptide peak area, (b) log_2_ transformed peptide peak area, (c) median normalized log_2_ transformed peptide peak area. Sample groups are highlighted with red (omental, OM) and blue (subcutaneous, SQ) in the log_2_ transformation and median normalization plots. Sample presentation order was organized for clarity and does not reflect sample run order, which was randomized. Figure is generated with level 2 and level 3a data.

### Peptide and Protein Detections in Adipose Tissue

Using a 1% false discovery rate (FDR) cut-off, 47,965 unique tryptic peptides were detected in the entire experiment corresponding to 4,438 proteins. The number of tryptic peptides in individual omental adipose tissue ranged from 21,786 to 39,807 and in subcutaneous tissue from 20,741 to 36,336 (Fig. 5). When the dataset was split to the adipose tissue from donors with and without obesity 21,271-36,188 tryptic peptides were detected in the individual samples from individuals without obesity and 20,741-39,807 peptides were detected in the samples from donors with obesity. The detected peptides in the adipose tissue map to 2,249-4,051 proteins in the individual samples (Fig. 5).

**Figure 5.**
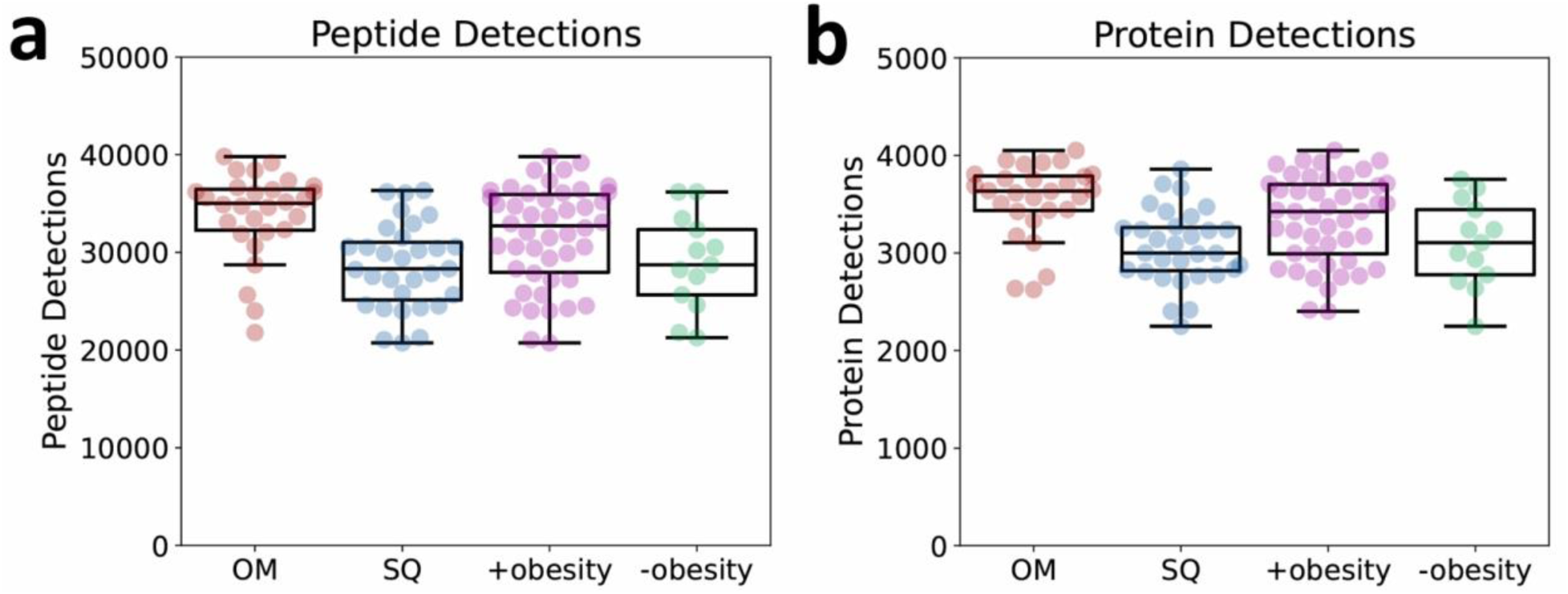
Box and Whisker plots of peptide and protein detections in different sample groups. (a) The number of peptides identified by EncyclopeDIA in each LC-MS/MS injection. (b) The number of proteins identified by EncyclopeDIA identified per LC-MS/MS injection. Samples are grouped by type: omental (OM), subcutaneous (SQ), with obesity or without obesity. Each sample is a member of two groups: either OM or SQ and either with obesity or without obesity. Data is presented at 1% FDR. All values are shown in Supplementary Table S2.

### Differences Between Subcutaneous and Omental Adipose Tissue

The overall proteomes of subcutaneous and omental adipose tissue had considerable overlap. This overlap was reflected in the principal component analysis of the omental and subcutaneous adipose tissue samples (Fig. 6a). Overall, 4,799 proteins were detected in the omental and subcutaneous adipose tissue. The protein abundance covered a wide dynamic range in both depots (Fig. 6b and c). Examples of adipose tissue marker proteins detected in both adipose tissue depots and in all donors include: FABP4, VINC, PLIN1, ADIPO, and LEP.^40–44^

**Figure 6.**
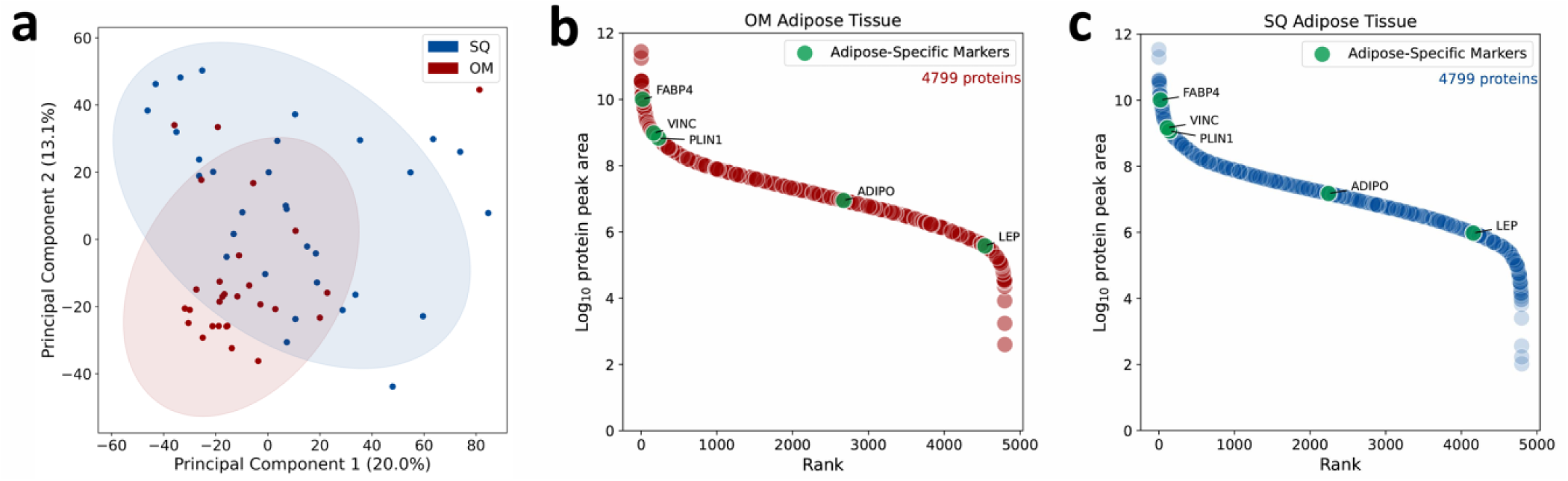
Overview of protein level adipose proteomics data. (a) Principal component analysis (PCA) on the entire dataset. The analysis was performed on directLFQ normalized protein peak areas without feature selection (level 3b data). The shaded area represents 95% confidence ellipses for omental (OM) and subcutaneous (SQ) adipose tissues. (b) Dynamic range plot of ranked, directLFQ normalized protein peak area in OM adipose tissue (level 3b data). (c) Dynamic range plot of ranked, directLFQ normalized protein peak area in SQ adipose tissue (level 3b data). Adipose tissue-specific proteins are labeled in both OM and SQ adipose tissues.

Despite a large overlap, many proteins were enriched in either subcutaneous or omental adipose tissue depots (Fig. 7). Overall, 200 proteins were enriched over 2-fold in the omental adipose tissue, and an additional 599 proteins were significantly enriched in omental adipose tissue but by less than 2-fold. In subcutaneous adipose tissue 93 proteins were enriched by over 2-fold and an additional 768 proteins were significantly enriched but by less than 2-fold when compared with omental adipose tissue. Hierarchical clustering of the differentially expressed proteins showed distinct omental and subcutaneous tissue clusters (Fig. 7).

**Figure 7.**
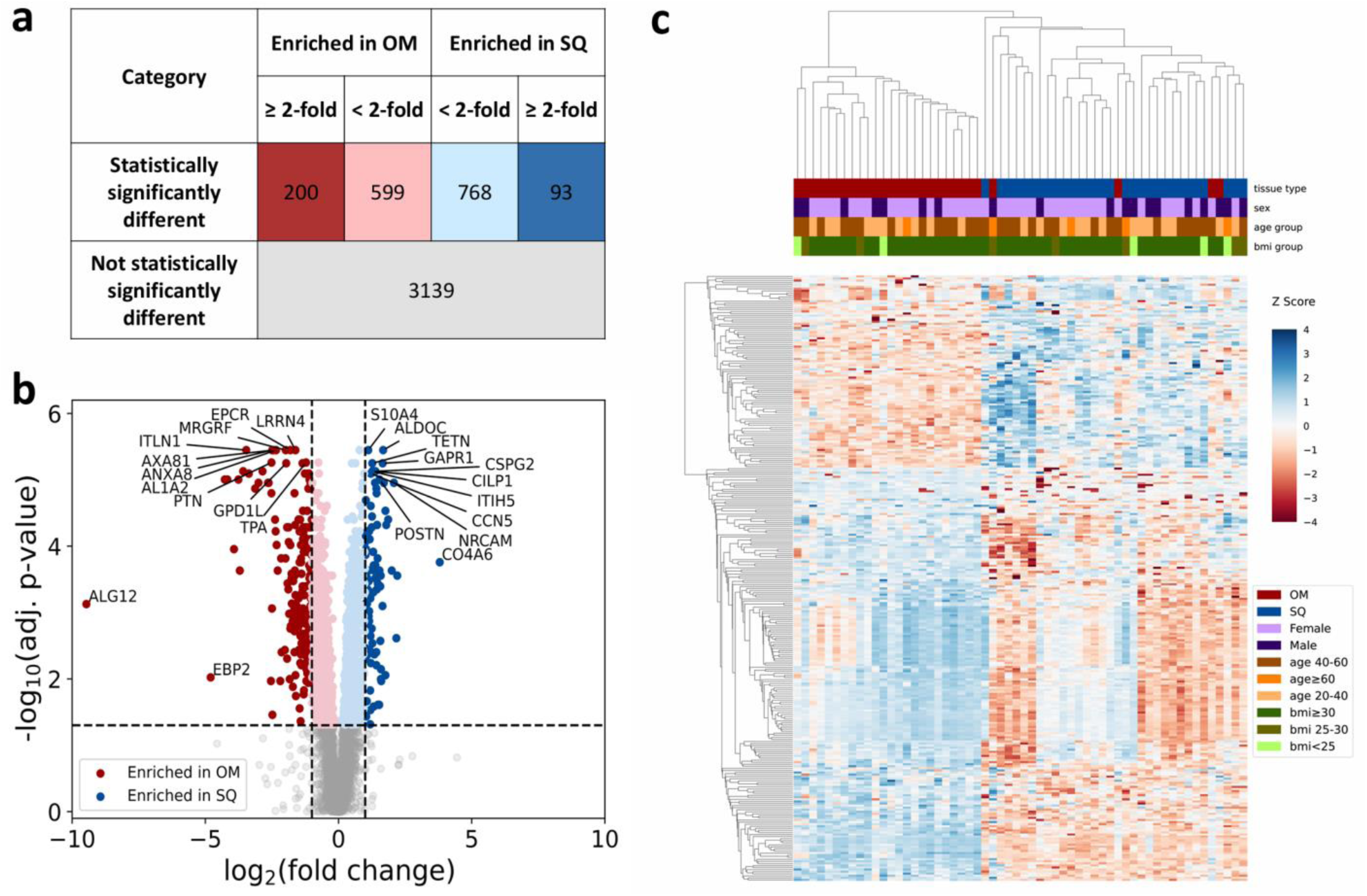
Summary data of proteomic differences observed between adipose tissue depots. (a) Table of numbers of proteins that are significally enriched (adjusted *p-value* < 0.05 and fold change > 2) in each depot or show no statistically meaningful difference in normalized protein peak area at 5% FDR. (b) Volcano plot of paired log2 fold changes in protein abundance between omental (OM) and subcutaneous (SQ) adipose tissues for the 28 study participants that have matched adipose tissue collected. A Wilcoxon signed-rank test was used to generate p-values. Statistically significantly different proteins are shown using a cut-off of log_2_ fold-change > 1 or < -1 and Benjamini-Hochberg adjusted *p-value* < 0.05. (c) Hierarchical clustering of the differentially expressed proteins in OM and SQ adipose tissue that were statistically significantly different as highlighted in b. A single sample was removed from (c) due to low protein abundance (see Methods). For visualization, Z scores were capped at -4 and +4 to limit the influence of extreme outliers on the color scale. Clustering analysis was performed on the uncapped values. Complete data is shown in Supplementary Table S3.

### Reproducibility and Verification

Using targeted proteomics quantification, we previously described depot-specific expression of retinoic acid synthesizing ALDH1A2, ALDH1A1 and aldehyde oxidase (AOX) in a limited number of human adipose tissue samples.^45^ The proteomics quantification was coupled with a measurement of the *all-trans-*retinoic acid synthesis activity associated with ALDH1A2, ALDH1A1 and AOX. In our previous study ADLH1A2, ALDH1A1, and AOX were found to have higher expression in omental than in subcutaneous adipose tissue. In the current dataset, untargeted DIA analysis of the depot-specific proteomes identified ALDH1A2 protein as significantly enriched in omental adipose tissue (Fig. 8), reproducing our previous findings. Similarly, ALDH1A1 and AOX expression was higher in omental than in subcutaneous adipose tissue in the current analysis.

**Figure 8.**
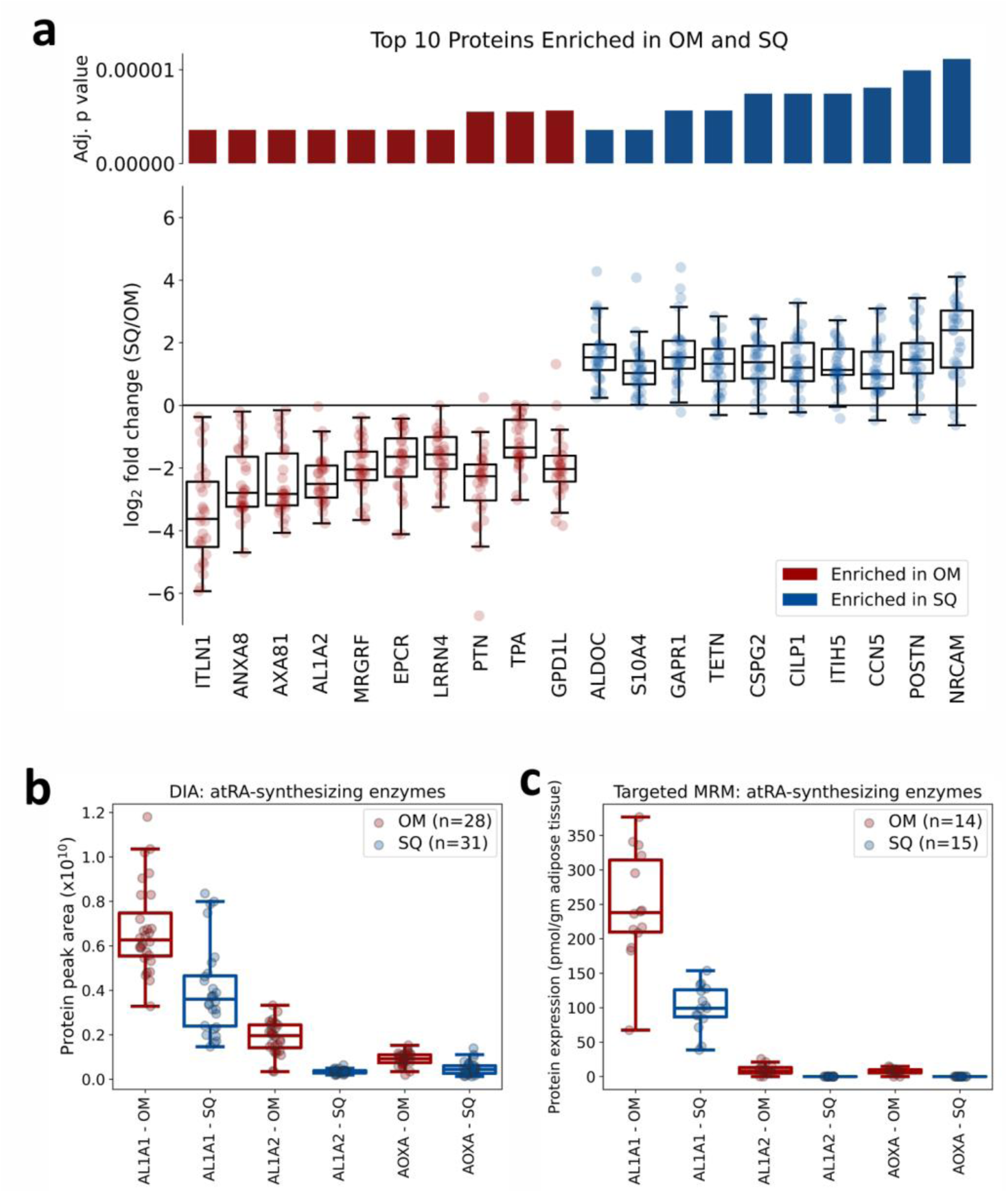
Differential expression of proteins in omental (OM) and subcutaneous (SQ) adipose tissues. (a) Box and Whisker plot for the log_2_ fold-change of the top 10 proteins that are enriched in omental (OM) adipose tissue and in subcutaneous (SQ) adipose tissue between 28 study participants. The proteins were filtered based on the adjusted *p-value* (lowest 10) and log_2_ fold-change (< -1 or >1) for significantly different proteins in each depot. Adjusted p values are shown in the bar chart above each protein shown. (b) Box and Whisker plot of *all-trans* retinoic acid-(*at*RA-) synthesizing protein peak area in all adipose tissue samples using data independent acquisition-mass spectrometry collected in the current study. (c) Previously published *at*RA-synthesizing enzymes’ expression in a subset of adipose tissue samples using targeted MRM method.^45^ The subset of samples analyzed in the previous study were from the same clinical study as the current work. The original observations from the 2022 study showing enrichment of the three *at*RA-synthesizing enzymes in omental adipose tissue were reproduced in the current study.

## Usage Notes

We provide peptide and protein level data of human subcutaneous and omental adipose tissue in individuals with a wide range of BMI. This dataset allows cross-evaluation of proteomic changes detected in adipose tissue in mouse models of obesity with human adipose tissue. The availability of data from individuals with a wide range of BMI allows exploratory analysis of potential proteomic changes that occur in each adipose tissue depot with increasing BMI.

### Use Case 1. Identification of Depot Differences Between Omental and Subcutaneous Adipose Tissue

The dataset presented here is from paired samples where each omental and subcutaneous sample is matched and from a single individual. The dataset can be used to mine depot differences in specific biochemical pathways that are associated with increasing adiposity in the absence of major comorbidities or organ impairment. An example of the differential protein expression detected in these data in subcutaneous versus omental adipose tissue is depicted in Figure 8. We show the top ten proteins significantly enriched in subcutaneous or omental adipose tissue. The associated metadata includes the adipocyte size measurements and key clinical characteristics in each participant expanding the opportunities to explore proteomic changes in human adipose tissue in depot specific manner.

### Use Case 2. Differential Protein and Peptide Abundance with Increasing BMI

Prior studies in mice have shown differences in adipose tissue proteome in lean and obese mice^9^ and with intermittent fasting.^10^ The current dataset can be used to assess whether similar proteome level differences are found in association with BMI in humans. A total of 19 proteins were found to significantly correlate with BMI (11 negative and 8 positive correlation) in omental adipose tissue (Figure 9). In subcutaneous adipose tissue 103 proteins correlated significantly with BMI (48 negative and 55 positive) (Figure 9). We show the four proteins (ITSN1, AGFG1, LDHD, and TXTP) that significantly correlated with BMI in both omental and subcutaneous adipose tissue in Figure 9. In our dataset, all individuals with BMI ≥30 had obesity; therefore BMI was used as a surrogate of adiposity. BMI has limitations as a surrogate marker for adiposity as it does not account for differences in muscle, bone and fat mass as contributors to height to weight ratio. Nevertheless, it provides a satisfactory measure of the adiposity in our study population with confirmed obesity in individuals with BMI ≥30 kg/m^2^ and confirmed lack of obesity in participants with BMI ≤27.5 kg/m^2^. The overall dataset expands the knowledge of human adipose tissue proteome as it includes tissue samples from both men and women spanning a wide range of BMIs. The metadata included with the proteomic dataset includes the key clinical features in the tissue donors that will aid in analysis of the proteomic changes along a continuum of increasing obesity.

**Figure 9.**
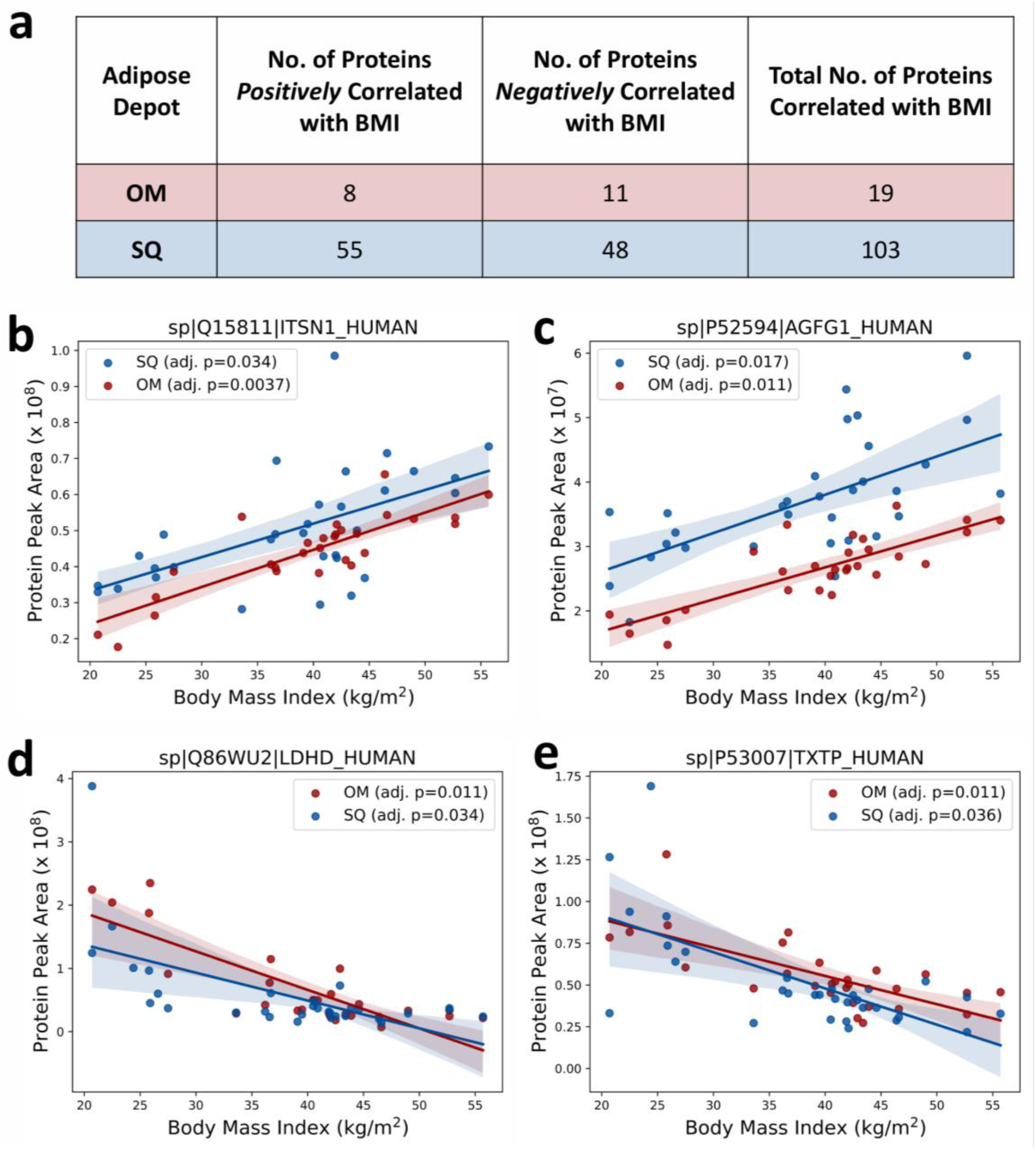
Summary data of proteomic signatures correlated with the level of adiposity observed in omental (OM) and subcutaneous (SQ) adipose tissues. (a) Table of numbers of proteins that statistically correlated with BMI (adjusted *p-value* < 0.05) in each depot. (b-e) Scatter plots of protein peak area against BMI for ITSN1, AGFG1, LDHD, and TXTP that showed correlations with BMI in both OM and SQ adipose tissues. The line and the shaded region represent fitted linear regression and 95% confidence intervals, respectively, for each panel.

### Use Case 3. Reuse and Reanalysis of the Mass Spectrometry Data

All the presented mass spectrometry data at the peptide and protein level have been collated for reuse, and all extracted ion chromatograms and peptide and protein identifications are available through the Panorama server for reanalysis. The DIA LC-MS/MS analysis allows the raw data to be easily reanalyzed, and the data can be used to explore any protein of interest in adipose tissue depots. This may be important for example for exploration on whether a specific potential new drug target is expressed in subcutaneous or omental adipose tissue or both. Our dataset also allows testing whether expression of a specific protein is associated with increasing adiposity or obesity using BMI as a surrogate marker of adiposity or by use of other metadata included. The dataset can also be further analyzed for presence of potential post translational modifications or different proteoforms in the adipose tissue depots.

## Code availability

MSConvert is available from https://proteowizard.sourceforge.io/. DIA-NN is available from https://github.com/vdemichev/DiaNN. Carafe is available from: https://github.com/Noble-Lab/Carafe. EncyclopeDIA is available from https://bitbucket.org/searleb/encyclopedia/. Skyline is available from https://skyline.ms/skyline.url.

All Nextflow workflows used in the current work to generate spectral libraries, search, quantify and normalize the MS data presented here are publicly available at: https://nf-carafe-ai-ms.readthedocs.io/en/latest/, for generation of spectral libraries using Carafe and DIA-NN; https://nf-teirex-dia.readthedocs.io/en/latest/, for generation of on-column chromatogram libraries followed by quantification using EncyclopeDIA and Skyline; https://github.com/uw-maccosslab/nf-dia-batch-correction, for peptide and protein level normalization.

All in house code written for the analysis of the data presented in this work is available online at https://github.com/Isoherranen-Lab/Human-Adipose-Depot-DIA.

## Funding resources

This work was supported by grants from the National Institutes of Health: R01GM111772, R01DK143511, T32GM007750, T32DK007247, TL1TR002318 R24GM141156, and U01DK137097. This work is supported in part by the University of Washington’s Proteomics Resource (UWPR95794).

## Abbreviations

OM: omental
SQ: subcutaneous
FDR: false discovery rate

## References

1. Stierman, B. et al. National Health and Nutrition Examination Survey 2017–March 2020 Prepandemic Data Files -- Development of Files and Prevalence Estimates for Selected Health Outcomes. Natl Health Stat Report 2021, (2021).

2. Busebee, B., Ghusn, W., Cifuentes, L. & Acosta, A. Obesity: A Review of Pathophysiology and Classification. Mayo Clin Proc 98, 1842–1857 (2023).

3. Heymsfield, S. B. & Wadden, T. A. Mechanisms, Pathophysiology, and Management of Obesity. New England Journal of Medicine 376, 254–266 (2017).

4. Ibrahim, M. M. Subcutaneous and visceral adipose tissue: Structural and functional differences. Obesity Reviews 11, 11–18 (2010).

5. Hruska, P. et al. Proteomic Signatures of Human Visceral and Subcutaneous Adipocytes. Journal of Clinical Endocrinology and Metabolism 107, 755–775 (2022).

6. Raajendiran, A. et al. Proteome analysis of human adipocytes identifies depot-specific heterogeneity at metabolic control points. Am J Physiol Endocrinol Metab 320, E1068–E1084 (2021).

7. Gómez-Serrano, M. et al. Proteome-wide alterations on adipose tissue from obese patients as age-, diabetes- and gender-specific hallmarks. Scientific Reports 2016 6:1 6, 1–15 (2016).

8. Larsen, J. K. et al. High-throughput proteomics uncovers exercise training and type 2 diabetes-induced changes in human white adipose tissue. Sci Adv 9, eadi7548 (2023).

9. Madsen, S. et al. Deep Proteome Profiling of White Adipose Tissue Reveals Marked Conservation and Distinct Features Between Different Anatomical Depots. Molecular and Cellular Proteomics 22, (2023).

10. Harney, D. J. et al. Proteomics analysis of adipose depots after intermittent fasting reveals visceral fat preservation mechanisms. Cell Rep 34, 108804 (2021).

11. Venable, J. D., Dong, M.-Q., Wohlschlegel, J., Dillin, A. & Yates, J. R. Automated approach for quantitative analysis of complex peptide mixtures from tandem mass spectra. Nat Methods 1, 39–45 (2004).

12. Egertson, J. D., MacLean, B., Johnson, R., Xuan, Y. & MacCoss, M. J. Multiplexed peptide analysis using data-independent acquisition and Skyline. Nat Protoc 10, 887–903 (2015).

13. Gillet, L. C. et al. Targeted Data Extraction of the MS/MS Spectra Generated by Data-independent Acquisition: A New Concept for Consistent and Accurate Proteome Analysis. Molecular & Cellular Proteomics 11, O111.016717 (2012).

14. Angulo, P. et al. The NAFLD fibrosis score: A noninvasive system that identifies liver fibrosis in patients with NAFLD. Hepatology 45, 846–854 (2007).

15. Batth, T. S. et al. Protein Aggregation Capture on Microparticles Enables Multipurpose Proteomics Sample Preparation. Mol Cell Proteomics 18, 1027–1035 (2019).

16. Hughes, C. S. et al. Single-pot, solid-phase-enhanced sample preparation for proteomics experiments. Nat Protoc 14, 68–85 (2019).

17. Tsantilas, K. A. et al. A Framework for Quality Control in Quantitative Proteomics. J Proteome Res (2024) doi:10.1021/acs.jproteome.4c00363.

18. Egertson, J. D. et al. Multiplexed MS/MS for improved data-independent acquisition. Nature Methods 2013 10:8 10, 744–746 (2013).

19. Amodei, D. et al. Improving Precursor Selectivity in Data-Independent Acquisition Using Overlapping Windows. J Am Soc Mass Spectrom 30, 669–684 (2019).

20. Searle, B. C. et al. Chromatogram libraries improve peptide detection and quantification by data independent acquisition mass spectrometry. Nat Commun 9, 5128 (2018).

21. Pino, L. K., Just, S. C., MacCoss, M. J. & Searle, B. C. Acquiring and Analyzing Data Independent Acquisition Proteomics Experiments without Spectrum Libraries. Molecular and Cellular Proteomics 19, 1088–1103 (2020).

22. Chambers, M. C. et al. A cross-platform toolkit for mass spectrometry and proteomics. Nat Biotechnol 30, 918–920 (2012).

23. Demichev, V., Messner, C. B., Vernardis, S. I., Lilley, K. S. & Ralser, M. DIA-NN: neural networks and interference correction enable deep proteome coverage in high throughput. Nat Methods 17, 41–44 (2020).

24. Wen, B. et al. Carafe Enables High Quality in Silico Spectral Library Generation for Data-Independent Acquisition Proteomics.

25. DI Tommaso, P., et al. Nextflow enables reproducible computational workflows. Nat Biotechnol 35, 316–319 (2017).

26. MacLean, B., et al. Skyline: An open source document editor for creating and analyzing targeted proteomics experiments. Bioinformatics 26, 966–968 (2010).

27. Pino, L. K. et al. The Skyline ecosystem: Informatics for quantitative mass spectrometry proteomics. Mass Spectrom Rev (2017) doi:10.1002/mas.21540.

28. Sharma, V., Eng, J. K., Maccoss, M. J. & Riffle, M. A mass spectrometry proteomics data management platform. Mol Cell Proteomics 11, 824–831 (2012).

29. Webb-Robertson, B. J. M., Matzke, M. M., Jacobs, J. M., Pounds, J. G. & Waters, K. M. A statistical selection strategy for normalization procedures in LC-MS proteomics experiments through dataset-dependent ranking of normalization scaling factors. Proteomics 11, 4736–4741 (2011).

30. Sharma, V. et al. Panorama Public: A Public Repository for Quantitative Data Sets Processed in Skyline. Mol Cell Proteomics 17, 1239–1244 (2018).

31. Merrihew, G. E. et al. A peptide-centric quantitative proteomics dataset for the phenotypic assessment of Alzheimer’s disease. Sci Data 10, (2023).

32. Ammar, C., Schessner, J. P., Willems, S., Michaelis, A. C. & Mann, M. Accurate Label-Free Quantification by directLFQ to Compare Unlimited Numbers of Proteomes. Molecular and Cellular Proteomics 22, (2023).

33. Harris, C. R. et al. Array programming with NumPy. Nature 2020 585:7825 585, 357–362 (2020).

34. McKinney, W. Data Structures for Statistical Computing in Python. scipy 56–61 (2010) doi:10.25080/MAJORA-92BF1922-00A.

35. Virtanen, P. et al. SciPy 1.0: fundamental algorithms for scientific computing in Python. Nat Methods 17, 261–272 (2020).

36. Pedregosa FABIANPEDREGOSA, F. et al. Scikit-learn: Machine Learning in Python. Journal of Machine Learning Research 12, 2825–2830 (2011).

37. Waskom, M. L. seaborn: statistical data visualization. J Open Source Softw 6, 3021 (2021).

38. Hunter, J. D. Matplotlib: A 2D graphics environment. Comput Sci Eng 9, 90–95 (2007).

39. Vizcaíno, J. A. et al. ProteomeXchange provides globally coordinated proteomics data submission and dissemination. Nature Biotechnology 2014 32:3 32, 223–226 (2014).

40. Furuhashi, M., Saitoh, S., Shimamoto, K. & Miura, T. Fatty Acid-Binding Protein 4 (FABP4): Pathophysiological Insights and Potent Clinical Biomarker of Metabolic and Cardiovascular Diseases. Clin Med Insights Cardiol 8, 23 (2015).

41. Li, J. et al. Vinculin: A new target for the diagnosis and treatment of disease. Prog Biophys Mol Biol 195, 157–166 (2025).

42. Sohn, J. H. et al. Perilipin 1 (Plin1) deficiency promotes inflammatory responses in lean adipose tissue through lipid dysregulation. Journal of Biological Chemistry 293, 13974–13988 (2018).

43. Straub, L. G. & Scherer, P. E. Metabolic Messengers: Adiponectin. Nat Metab 1, 334 (2019).

44. Bäckdahl, J. et al. Spatial mapping reveals human adipocyte subpopulations with distinct sensitivities to insulin. Cell Metab 33, 1869–1882.e6 (2021).

45. Rubinow, K. B. et al. Evidence of depot-specific regulation of all-trans-retinoic acid biosynthesis in human adipose tissue. Clin Transl Sci 15, 1460–1471 (2022).

