## Supplemental table 3 for "Proteomic Profiling of Human Omental and Subcutaneous Adipose Tissue in Individuals with a Broad Range of BMI"

| Protein | Mean OM Prot | Mean SC Prot | log2(Fold Cha | P Value | Adjusted P Value |
|---|---|---|---|---|---|
| sp\|P13928\|A | 225655632 | 34253294.4 | -2.48864 | 7.45E-09 | 3.58E-06 |
| sp\|Q96AM1\|I | 9785571.31 | 2869711.68 | -1.9800854 | 7.45E-09 | 3.58E-06 |
| sp\|P09110\|T | 40000507.4 | 70564662.6 | 0.78782531 | 7.45E-09 | 3.58E-06 |
| sp\|P09972\|A | 399992958 | 1432554000 | 1.67510951 | 7.45E-09 | 3.58E-06 |
| sp\|Q5VT79\|A | 230245784 | 35262195.6 | -2.444365 | 7.45E-09 | 3.58E-06 |
| sp\|Q8WWA0 | 128637845 | 11864321.3 | -3.4626645 | 7.45E-09 | 3.58E-06 |
| sp\|Q8WUT4\| | 29244081.3 | 9781089.92 | -1.6166913 | 7.45E-09 | 3.58E-06 |
| sp\|O94788\|A | 1876587699 | 345944607 | -2.3447325 | 7.45E-09 | 3.58E-06 |
| sp\|P26447\|S | 199130829 | 479179025 | 1.11718759 | 7.45E-09 | 3.58E-06 |
| sp\|Q9UNN8\| | 50729821.5 | 16035310.6 | -1.8105663 | 7.45E-09 | 3.58E-06 |
| sp\|P21246\|P | 6624284.83 | 1094611.32 | -2.5157111 | 1.49E-08 | 5.50E-06 |
| sp\|P50453\|S | 175828510 | 101569303 | -0.7531661 | 1.49E-08 | 5.50E-06 |
| sp\|P00750\|T | 2143704.55 | 1031509.28 | -1.2186597 | 1.49E-08 | 5.50E-06 |
| sp\|P05452\|T | 151269873 | 382267856 | 1.26695523 | 2.24E-08 | 5.65E-06 |
| sp\|Q03135\|C | 145363274 | 229913377 | 0.63499877 | 2.24E-08 | 5.65E-06 |
| sp\|Q9H4G4\| | 18810029.1 | 66360580.8 | 1.65716001 | 2.24E-08 | 5.65E-06 |
| sp\|Q8N335\|C | 228086956 | 64031038 | -1.9634084 | 2.24E-08 | 5.65E-06 |
| sp\|P52594\|A | 26608677.5 | 37895679.2 | 0.50229129 | 2.24E-08 | 5.65E-06 |
| sp\|O00592\|P | 19952444.8 | 7551190.14 | -1.3526283 | 2.24E-08 | 5.65E-06 |
| sp\|P05783\|K | 1155311270 | 100320320 | -3.5785475 | 3.73E-08 | 7.45E-06 |
| sp\|Q13421\|M | 21076694.9 | 2998484.88 | -2.8518372 | 3.73E-08 | 7.45E-06 |
| sp\|O75339\|C | 8330950.65 | 21798314.3 | 1.29445471 | 3.73E-08 | 7.45E-06 |
| sp\|P13611\|C | 54331585 | 158971739 | 1.38642746 | 3.73E-08 | 7.45E-06 |
| sp\|Q86UX2\|I | 21511738.9 | 55321289.2 | 1.27116097 | 3.73E-08 | 7.45E-06 |
| sp\|Q6UVK1\|C | 45987757.1 | 91201391.3 | 0.96343805 | 5.22E-08 | 8.07E-06 |
| sp\|P05023\|A | 71134191.4 | 31608011.8 | -1.1018877 | 5.22E-08 | 8.07E-06 |
| sp\|Q9NZR1\|T | 34749798.6 | 14854935.5 | -1.2637786 | 5.22E-08 | 8.07E-06 |
| sp\|Q8IWA5\|C | 19021853.6 | 11357666.9 | -0.735352 | 5.22E-08 | 8.07E-06 |
| sp\|O76076\|C | 2368612.02 | 6246244.84 | 1.21351538 | 5.22E-08 | 8.07E-06 |
| sp\|P19012\|K | 433266992 | 52612469.8 | -3.3661902 | 5.22E-08 | 8.07E-06 |
| sp\|Q9NZA1\|C | 75752251 | 34330612 | -1.1426431 | 5.22E-08 | 8.07E-06 |
| sp\|Q9BVJ7\|D | 8365463.6 | 3660379.29 | -1.0891871 | 7.45E-08 | 9.93E-06 |
| sp\|P05787\|K | 6662121549 | 520763822 | -3.7561099 | 7.45E-08 | 9.93E-06 |
| sp\|Q15063\|P | 118163612 | 357756812 | 1.53828857 | 7.45E-08 | 9.93E-06 |
| sp\|P01303\|N | 920336.242 | 79533.7077 | -4.2688158 | 7.45E-08 | 9.93E-06 |
| sp\|P08727\|K | 1203244116 | 75104010.6 | -4.1715613 | 7.45E-08 | 9.93E-06 |
| sp\|Q9HCU0\| | 70948685.8 | 286732958 | 1.70016559 | 1.04E-07 | 1.11E-05 |
| sp\|Q92823\|N | 7868912.73 | 43289543.4 | 2.07848511 | 1.04E-07 | 1.11E-05 |

| | | | | | |
|---|---|---|---|---|---|
| sp\|Q9UJ83\|H | 5934331.12 | 16006139.4 | 1.36978501 | 1.04E-07 | 1.11E-05 |
| sp\|Q13219\|F | 13425531.3 | 6200620.53 | -1.0527487 | 1.04E-07 | 1.11E-05 |
| sp\|P13646\|K | 319340631 | 46881451.5 | -3.0030335 | 1.04E-07 | 1.11E-05 |
| sp\|Q9BY76\|A | 1541163.19 | 2960248.87 | 0.90128449 | 1.04E-07 | 1.11E-05 |
| sp\|Q9NRN5\| | 48374000.7 | 24335796.2 | -0.9770139 | 1.04E-07 | 1.11E-05 |
| sp\|Q04695\|K | 337178600 | 61526199.7 | -2.6205235 | 1.04E-07 | 1.11E-05 |
| sp\|Q8IVN8\|S | 1292070.74 | 176482.417 | -2.6460327 | 1.04E-07 | 1.11E-05 |
| sp\|Q5TZA2\|C | 16528595.2 | 2173285.31 | -3.1287221 | 1.42E-07 | 1.36E-05 |
| sp\|P15311\|E | 1357181850 | 643683176 | -1.0580068 | 1.42E-07 | 1.36E-05 |
| sp\|P01033\|T | 238163175 | 111821918 | -1.131836 | 1.42E-07 | 1.36E-05 |
| sp\|P14384\|C | 20329491.7 | 53522609.3 | 1.42192373 | 1.42E-07 | 1.36E-05 |
| sp\|Q5JPB2\|Z | 33781636 | 62885312.1 | 0.89271154 | 1.42E-07 | 1.36E-05 |
| sp\|Q9C0B1\|I | 22557115.7 | 12942608.2 | -0.8009288 | 1.86E-07 | 1.60E-05 |
| sp\|Q14CN4\| | 109188756 | 20404122.1 | -2.518389 | 1.86E-07 | 1.60E-05 |
| sp\|O94875\|S | 99658609.1 | 57967751.5 | -0.7712713 | 1.86E-07 | 1.60E-05 |
| sp\|P41159\|L | 417932.283 | 1259926.64 | 1.42929626 | 1.86E-07 | 1.60E-05 |
| sp\|Q13308\|F | 18368264.3 | 5454897.76 | -1.6518939 | 1.86E-07 | 1.60E-05 |
| sp\|P83916\|C | 36646282.4 | 21799571.2 | -0.7490479 | 1.86E-07 | 1.60E-05 |
| sp\|O95297\|N | 772386.43 | 456855.357 | -0.8533697 | 2.46E-07 | 2.03E-05 |
| sp\|P55268\|L | 429571880 | 862643902 | 1.02843018 | 2.46E-07 | 2.03E-05 |
| sp\|O95861\|E | 99692808.8 | 63308362.6 | -0.6688755 | 3.20E-07 | 2.48E-05 |
| sp\|P09486\|S | 90864620.1 | 216573978 | 1.21007443 | 3.20E-07 | 2.48E-05 |
| sp\|P23193\|T | 18057469.1 | 13348207.5 | -0.4398488 | 3.20E-07 | 2.48E-05 |
| sp\|O60240\|F | 704080236 | 1256757176 | 0.80886481 | 3.20E-07 | 2.48E-05 |
| sp\|P16870\|C | 1407815.64 | 576360.727 | -1.3357115 | 4.10E-07 | 2.94E-05 |
| sp\|Q6UB28\|I | 709841.881 | 2413410.95 | 1.76126424 | 4.10E-07 | 2.94E-05 |
| sp\|O95848\|N | 12302150.5 | 26111977 | 0.8968913 | 4.10E-07 | 2.94E-05 |
| sp\|O43175\|S | 111799225 | 53117967.7 | -1.1763161 | 4.10E-07 | 2.94E-05 |
| sp\|O00423\|E | 92388899.8 | 64761335.7 | -0.5148202 | 4.10E-07 | 2.94E-05 |
| sp\|O60664\|F | 69650720.1 | 139007175 | 0.86582939 | 5.22E-07 | 3.58E-05 |
| sp\|A2A2Z9\|A | 3084362.9 | 7960327.56 | 1.25816091 | 5.22E-07 | 3.58E-05 |
| sp\|P55011\|S | 8771024.14 | 12603548 | 0.52562392 | 5.22E-07 | 3.58E-05 |
| sp\|Q07666\|K | 31356776 | 20215788.6 | -0.6335967 | 6.56E-07 | 3.98E-05 |
| sp\|Q16647\|F | 22705391.1 | 7384595.8 | -1.5028169 | 6.56E-07 | 3.98E-05 |
| sp\|P00352\|A | 6718894506 | 3946298199 | -0.8751378 | 6.56E-07 | 3.98E-05 |
| sp\|Q6UWP8\| | 144601.954 | 742178.743 | 1.863291 | 6.56E-07 | 3.98E-05 |
| sp\|Q969G5\|C | 459990453 | 689725141 | 0.57611332 | 6.56E-07 | 3.98E-05 |
| sp\|P20648\|A | 7709470.99 | 2975826.92 | -1.4402085 | 6.56E-07 | 3.98E-05 |
| sp\|Q9Y5K6\|C | 71970740.3 | 91867262.6 | 0.33977052 | 6.56E-07 | 3.98E-05 |

| | | | | | |
|---|---|---|---|---|---|
| sp\|Q9NXD2\| | 6826030.64 | 2442952.52 | -2.381243 | 6.56E-07 | 3.98E-05 |
| sp\|Q9H6N6\| | 9891802.53 | 5361219.36 | -1.0091646 | 6.56E-07 | 3.98E-05 |
| sp\|Q9HBR0\|! | 864179.038 | 3511783.63 | 1.79223096 | 8.20E-07 | 4.80E-05 |
| sp\|Q99685\|N | 64489077.4 | 179512445 | 1.44826525 | 8.20E-07 | 4.80E-05 |
| sp\|P62861\|R | 3133531.79 | 1138368.48 | -1.6275118 | 8.20E-07 | 4.80E-05 |
| sp\|Q9Y3Y2\|C | 7737539.62 | 4500371.82 | -0.8206602 | 1.02E-06 | 5.21E-05 |
| sp\|P08729\|K | 4779321957 | 1013930885 | -1.8745112 | 1.02E-06 | 5.21E-05 |
| sp\|P55899\|F | 19175311.3 | 12831335.1 | -0.6564414 | 1.02E-06 | 5.21E-05 |
| sp\|Q8NFW8\| | 11910998.8 | 5104158 | -1.2268302 | 1.02E-06 | 5.21E-05 |
| sp\|Q9H2U1\| | 22778089.5 | 12184194.8 | -1.0300989 | 1.02E-06 | 5.21E-05 |
| sp\|Q8TD08\|N | 17216772.4 | 8990241.72 | -1.1282172 | 1.02E-06 | 5.21E-05 |
| sp\|Q9H330\|T | 3126883.82 | 1531887.78 | -1.3194417 | 1.02E-06 | 5.21E-05 |
| sp\|P98194\|A | 1335493.45 | 399244.412 | -1.9303614 | 1.02E-06 | 5.21E-05 |
| sp\|O96019\|A | 25224637.1 | 15609116.3 | -0.7117756 | 1.02E-06 | 5.21E-05 |
| sp\|P22413\|E | 4150080.66 | 9492385.21 | 1.2474031 | 1.02E-06 | 5.21E-05 |
| sp\|O75643\|U | 10205118.7 | 6175746.08 | -0.6861611 | 1.02E-06 | 5.21E-05 |
| sp\|P39656\|C | 71668760.2 | 35628438.1 | -1.0918498 | 1.02E-06 | 5.21E-05 |
| sp\|P78559\|M | 17412740.5 | 31812059.9 | 0.79204039 | 1.26E-06 | 5.98E-05 |
| sp\|Q8IYS5\|O | 213851.413 | 548713.325 | 1.08438738 | 1.26E-06 | 5.98E-05 |
| sp\|P51531\|S | 5273160.1 | 2037551.93 | -1.465157 | 1.26E-06 | 5.98E-05 |
| sp\|Q9UKX2\|N | 52056998.9 | 11947929.5 | -2.3593906 | 1.26E-06 | 5.98E-05 |
| sp\|Q5M775\|( | 4387299 | 6880416.14 | 0.63540042 | 1.26E-06 | 5.98E-05 |
| sp\|P00167\|C | 14777352.2 | 30862031.4 | 1.14861987 | 1.26E-06 | 5.98E-05 |
| sp\|P21980\|T | 67516805.4 | 28044110.4 | -1.3116773 | 1.26E-06 | 5.98E-05 |
| sp\|Q9Y6R7\|F | 10187318.9 | 16371168.4 | 0.67268767 | 1.54E-06 | 7.05E-05 |
| sp\|P27487\|D | 97736417.4 | 41843263.1 | -1.3012217 | 1.54E-06 | 7.05E-05 |
| sp\|Q07820\|N | 14386406.8 | 9074112.68 | -0.6954734 | 1.54E-06 | 7.05E-05 |
| sp\|Q96CG8\|( | 20920679.5 | 46605393.5 | 1.02327764 | 1.54E-06 | 7.05E-05 |
| sp\|P00387\|N | 69083839.9 | 115396975 | 0.69814249 | 1.88E-06 | 7.87E-05 |
| sp\|Q9BQI9\|N | 23533582.5 | 12448043.2 | -0.9080178 | 1.88E-06 | 7.87E-05 |
| sp\|O14745\|N | 157318549 | 68938694 | -1.1764638 | 1.88E-06 | 7.87E-05 |
| sp\|Q96F85\|C | 77600660.9 | 38291129 | -1.0585201 | 1.88E-06 | 7.87E-05 |
| sp\|O94911\|A | 15945632 | 10314925.5 | -0.6647968 | 1.88E-06 | 7.87E-05 |
| sp\|O75891\|A | 1020420938 | 449124662 | -1.3349385 | 1.88E-06 | 7.87E-05 |
| sp\|Q8N5R6\|( | 13097034.6 | 18648397.3 | 0.48934815 | 1.88E-06 | 7.87E-05 |
| sp\|Q9BX68\|H | 86842556.5 | 140152745 | 0.62614948 | 1.88E-06 | 7.87E-05 |
| sp\|P12111\|C | 476348619 | 1129226069 | 1.17090646 | 1.88E-06 | 7.87E-05 |
| sp\|P54289\|C | 46899332.1 | 72366095.4 | 0.60446553 | 1.88E-06 | 7.87E-05 |
| sp\|Q6UWP2\| | 33539039.7 | 65788906.8 | 0.82783354 | 2.29E-06 | 8.71E-05 |

| | | | | | |
|---|---|---|---|---|---|
| sp\|P08237\|P | 184200370 | 104921768 | -0.8928371 | 2.29E-06 | 8.71E-05 |
| sp\|Q9Y2S2\|( | 51390144.5 | 33895221.8 | -0.6328527 | 2.29E-06 | 8.71E-05 |
| sp\|P43121\|M | 364621747 | 729481222 | 1.003312 | 2.29E-06 | 8.71E-05 |
| sp\|Q08211\|D | 173055520 | 92221271.9 | -0.9239712 | 2.29E-06 | 8.71E-05 |
| sp\|P18031\|P | 15398050.6 | 4965263.87 | -1.868916 | 2.29E-06 | 8.71E-05 |
| sp\|Q52LC2\|\ | 1633388.48 | 789075.875 | -1.0439076 | 2.29E-06 | 8.71E-05 |
| sp\|Q06828\|F | 25965558 | 44638301 | 0.74451262 | 2.29E-06 | 8.71E-05 |
| sp\|Q14315\|F | 701694343 | 345345659 | -0.9826772 | 2.29E-06 | 8.71E-05 |
| sp\|Q96IJ6\|GI | 35074673.8 | 24860674.6 | -0.5039814 | 2.29E-06 | 8.71E-05 |
| sp\|P08294\|S | 158849023 | 263558759 | 0.71196051 | 2.29E-06 | 8.71E-05 |
| sp\|P62277\|R | 128398197 | 35454281.5 | -2.32273 | 2.76E-06 | 9.61E-05 |
| sp\|P08138\|T | 962932.624 | 333285.035 | -1.8060332 | 2.76E-06 | 9.61E-05 |
| sp\|P53384\|N | 5565447.53 | 3860710.85 | -0.5610528 | 2.76E-06 | 9.61E-05 |
| sp\|P07101\|T | 2591428.4 | 992847.091 | -1.2667193 | 2.76E-06 | 9.61E-05 |
| sp\|P46952\|3 | 40626468.3 | 25539597.2 | -0.7267477 | 2.76E-06 | 9.61E-05 |
| sp\|Q7Z4H3\|I | 11088332.3 | 7381270.15 | -0.608992 | 2.76E-06 | 9.61E-05 |
| sp\|Q8IUX7\|A | 30943803.2 | 16605632.8 | -0.8012674 | 2.76E-06 | 9.61E-05 |
| sp\|O43491\|E | 353097992 | 249172436 | -0.5032888 | 2.76E-06 | 9.61E-05 |
| sp\|P30039\|P | 21499835.5 | 33480202.2 | 0.62029585 | 2.76E-06 | 9.61E-05 |
| sp\|Q9UEY8\|/ | 112171422 | 61096219.1 | -0.9098126 | 2.76E-06 | 9.61E-05 |
| sp\|P28827\|P | 22082014.8 | 34796709.8 | 0.59325427 | 2.76E-06 | 9.61E-05 |
| sp\|Q53GG5\| | 32068504.7 | 13329821.1 | -1.4132609 | 2.76E-06 | 9.61E-05 |
| sp\|Q15124\|F | 307646253 | 153226690 | -1.0325109 | 3.33E-06 | 0.00011099 |
| sp\|Q14116\|I | 109225573 | 50278914.9 | -1.1873212 | 3.33E-06 | 0.00011099 |
| sp\|P17302\|C | 232032.325 | 66589.8356 | -3.9231658 | 3.33E-06 | 0.00011099 |
| sp\|P51151\|R | 12517631.2 | 20591190.3 | 0.64946191 | 3.33E-06 | 0.00011099 |
| sp\|P61019\|R | 103297565 | 153805126 | 0.54434847 | 3.33E-06 | 0.00011099 |
| sp\|Q9UI17\|M | 19271970.3 | 42129970.6 | 1.03267127 | 3.33E-06 | 0.00011099 |
| sp\|P10643\|C | 266180383 | 139595835 | -1.1915513 | 3.99E-06 | 0.00012207 |
| sp\|Q5MNZ9\| | 1370487.16 | 2479003.17 | 0.97584958 | 3.99E-06 | 0.00012207 |
| sp\|Q02790\|F | 73440325.1 | 49616194.8 | -0.6114334 | 3.99E-06 | 0.00012207 |
| sp\|P04040\|C | 5801423001 | 9666183463 | 0.69555446 | 3.99E-06 | 0.00012207 |
| sp\|P07942\|L | 577976280 | 307053033 | -1.0090921 | 3.99E-06 | 0.00012207 |
| sp\|Q9P2B2\|F | 20586467.4 | 9205954.57 | -1.1169453 | 3.99E-06 | 0.00012207 |
| sp\|Q03252\|L | 276532936 | 165421002 | -0.7349059 | 3.99E-06 | 0.00012207 |
| sp\|Q9H6B4\|( | 2129495.83 | 5791087.62 | 1.30242746 | 3.99E-06 | 0.00012207 |
| sp\|Q99523\|S | 8465805.3 | 12376362.5 | 0.63564049 | 3.99E-06 | 0.00012207 |
| sp\|Q9H5N1\| | 69354776.4 | 91457285.1 | 0.41185466 | 3.99E-06 | 0.00012207 |
| sp\|Q13547\|H | 10415350.1 | 5947345.64 | -0.8350025 | 3.99E-06 | 0.00012207 |

| | | | | | |
|---|---|---|---|---|---|
| sp\|Q9NXG2\|' | 5374942.65 | 2499019.66 | -1.2346141 | 3.99E-06 | 0.00012207 |
| sp\|Q71UM5\| | 24396451.5 | 9895018.76 | -1.3072521 | 3.99E-06 | 0.00012207 |
| sp\|Q9UI09\|N | 7099465.14 | 3913389.72 | -1.0061121 | 4.77E-06 | 0.00013541 |
| sp\|Q7Z3D6\|( | 59206815.6 | 35035297.9 | -0.7273359 | 4.77E-06 | 0.00013541 |
| sp\|P39059\|C | 62687085.8 | 128583531 | 1.01834334 | 4.77E-06 | 0.00013541 |
| sp\|P43490\|N | 44357784.8 | 82850333.8 | 0.85351328 | 4.77E-06 | 0.00013541 |
| sp\|P98160\|P | 382082871 | 581180778 | 0.56528448 | 4.77E-06 | 0.00013541 |
| sp\|P00488\|F | 505034700 | 239162375 | -1.0089062 | 4.77E-06 | 0.00013541 |
| sp\|P16050\|L | 39981859.5 | 23414699.4 | -0.8210708 | 4.77E-06 | 0.00013541 |
| sp\|O43556\|S | 4485348.4 | 8369776.41 | 0.84705014 | 4.77E-06 | 0.00013541 |
| sp\|Q96S97\|N | 22303736.7 | 15894792.4 | -0.5405073 | 4.77E-06 | 0.00013541 |
| sp\|Q04726\|T | 7777080.81 | 5506090.51 | -0.528597 | 4.77E-06 | 0.00013541 |
| sp\|P30041\|P | 2393806159 | 1195797523 | -0.9639013 | 4.77E-06 | 0.00013541 |
| sp\|O15254\|A | 19319955.5 | 29715066.9 | 0.58096542 | 4.77E-06 | 0.00013541 |
| sp\|A6NDG6\| | 62782594.9 | 90765482.5 | 0.52256697 | 5.68E-06 | 0.00015307 |
| sp\|Q13231\|( | 5109557.89 | 21427175.6 | 1.46815861 | 5.68E-06 | 0.00015307 |
| sp\|Q96S19\|N | 19786500.3 | 11451999.1 | -0.874341 | 5.68E-06 | 0.00015307 |
| sp\|P14174\|N | 836466293 | 570474229 | -0.6194694 | 5.68E-06 | 0.00015307 |
| sp\|P55263\|A | 132482010 | 232820416 | 0.70528011 | 5.68E-06 | 0.00015307 |
| sp\|O95613\|F | 7010758.5 | 1943439.83 | -2.1564923 | 5.68E-06 | 0.00015307 |
| sp\|O60279\|S | 8162443.57 | 2910955.97 | -1.4071536 | 5.68E-06 | 0.00015307 |
| sp\|Q13451\|F | 692145.188 | 1391619.62 | 0.81077374 | 5.68E-06 | 0.00015307 |
| sp\|P62280\|R | 116875615 | 34771356.6 | -1.9902717 | 5.68E-06 | 0.00015307 |
| sp\|Q8WXE0\| | 1750615.68 | 3812559.25 | 0.9230018 | 6.74E-06 | 0.00017472 |
| sp\|Q00839\|H | 175682083 | 85530284.2 | -0.9974572 | 6.74E-06 | 0.00017472 |
| sp\|Q9NXC2\|' | 755298.191 | 458824.273 | -0.8130989 | 6.74E-06 | 0.00017472 |
| sp\|Q14031\|( | 198609.403 | 1096073.39 | 3.80018606 | 6.74E-06 | 0.00017472 |
| sp\|P62263\|R | 74864578.7 | 29740806.2 | -1.4043879 | 6.74E-06 | 0.00017472 |
| sp\|P61247\|R | 249937046 | 100429730 | -1.4623286 | 6.74E-06 | 0.00017472 |
| sp\|O43493\|T | 11717445 | 20560343.5 | 0.74397027 | 6.74E-06 | 0.00017472 |
| sp\|Q07065\|( | 31883431.7 | 18392188.5 | -0.7822971 | 7.96E-06 | 0.00019501 |
| sp\|P31949\|S | 191315925 | 303155424 | 0.64141768 | 7.96E-06 | 0.00019501 |
| sp\|P17661\|D | 5869035292 | 2254352231 | -1.1040868 | 7.96E-06 | 0.00019501 |
| sp\|Q96A49\|S | 1575648.6 | 2914909.84 | 0.73748176 | 7.96E-06 | 0.00019501 |
| sp\|P98179\|R | 48088874 | 123191339 | 1.31855102 | 7.96E-06 | 0.00019501 |
| sp\|O76070\|S | 405510679 | 1088118376 | 1.38602346 | 7.96E-06 | 0.00019501 |
| sp\|Q92522\|H | 38594310.7 | 17988805.4 | -1.0850154 | 7.96E-06 | 0.00019501 |
| sp\|O15400\|S | 99469516.8 | 130914756 | 0.39395194 | 7.96E-06 | 0.00019501 |
| sp\|Q9NPY3\|( | 28159207.6 | 38796654.6 | 0.45174338 | 7.96E-06 | 0.00019501 |

| | | | | | |
|---|---|---|---|---|---|
| sp\|Q2KHR3\|( | 6440627.17 | 13321142.7 | 0.91494281 | 7.96E-06 | 0.00019501 |
| sp\|Q8WWY3\| | 3055935.78 | 1428242.75 | -1.086624 | 7.96E-06 | 0.00019501 |
| sp\|P54802\|A | 16670556.1 | 30262895.5 | 0.75768223 | 9.40E-06 | 0.00021781 |
| sp\|P12955\|P | 90995947.3 | 150590648 | 0.59552898 | 9.40E-06 | 0.00021781 |
| sp\|Q5JS37\|N | 11375724.9 | 21226896.9 | 0.82879276 | 9.40E-06 | 0.00021781 |
| sp\|O00764\|F | 421239115 | 789181871 | 0.70014344 | 9.40E-06 | 0.00021781 |
| sp\|Q13822\|E | 6756839.55 | 11379041.9 | 0.70187994 | 9.40E-06 | 0.00021781 |
| sp\|Q08174\|F | 17352706.3 | 24625287.9 | 0.53311658 | 9.40E-06 | 0.00021781 |
| sp\|Q92841\|D | 221057273 | 116912764 | -0.8355455 | 9.40E-06 | 0.00021781 |
| sp\|P49189\|A | 1598856625 | 2848845780 | 0.66105615 | 9.40E-06 | 0.00021781 |
| sp\|A4D0S4\|L | 16622474.8 | 32742970.7 | 0.99482252 | 9.40E-06 | 0.00021781 |
| sp\|Q99759\|N | 13617587.4 | 7217343.5 | -1.0595949 | 9.40E-06 | 0.00021781 |
| sp\|P49407\|A | 46829685.4 | 29442327.3 | -0.6626293 | 9.40E-06 | 0.00021781 |
| sp\|Q96J01\|T | 577015.852 | 274588.419 | -2.2830133 | 1.10E-05 | 0.00023359 |
| sp\|Q15067\|A | 52116634.6 | 104176457 | 1.13336001 | 1.10E-05 | 0.00023359 |
| sp\|Q00577\|F | 55771756.3 | 36142118.2 | -0.651384 | 1.10E-05 | 0.00023359 |
| sp\|P61978\|H | 294391966 | 184595917 | -0.6833418 | 1.10E-05 | 0.00023359 |
| sp\|P02452\|C | 114573594 | 689234044 | 1.99801344 | 1.10E-05 | 0.00023359 |
| sp\|P43351\|R | 5388057.17 | 14038617.3 | 1.34719601 | 1.10E-05 | 0.00023359 |
| sp\|P02461\|C | 52100575 | 170780349 | 1.53912362 | 1.10E-05 | 0.00023359 |
| sp\|Q8N684\|( | 4658995.13 | 3103453.18 | -0.6070535 | 1.10E-05 | 0.00023359 |
| sp\|P07203\|G | 608015668 | 1074166286 | 0.81275517 | 1.10E-05 | 0.00023359 |
| sp\|Q13561\|D | 167442602 | 225496723 | 0.42991419 | 1.10E-05 | 0.00023359 |
| sp\|P36969\|G | 63341458 | 109361670 | 0.78790055 | 1.10E-05 | 0.00023359 |
| sp\|P33151\|C | 17117081.7 | 28885405.5 | 0.78244704 | 1.10E-05 | 0.00023359 |
| sp\|P0DMV8\|I | 3863806486 | 2501981507 | -0.5854261 | 1.10E-05 | 0.00023359 |
| sp\|Q9H0W5\| | 158856.02 | 42289.6797 | -3.7055064 | 1.10E-05 | 0.00023359 |
| sp\|Q13554\|K | 26669597.2 | 15466664.5 | -0.7935801 | 1.10E-05 | 0.00023359 |
| sp\|Q6GTX8\|L | 5048389.89 | 8831309.97 | 0.76509709 | 1.10E-05 | 0.00023359 |
| sp\|P56181\|N | 2197781.39 | 3480305.54 | 0.80393909 | 1.10E-05 | 0.00023359 |
| sp\|Q3KQU3\| | 27262046.7 | 37694858.6 | 0.49326962 | 1.10E-05 | 0.00023359 |
| sp\|P12532\|K | 66173219.7 | 15998412.4 | -1.6383027 | 1.10E-05 | 0.00023359 |
| sp\|P62987\|R | 568843554 | 771417545 | 0.40122343 | 1.10E-05 | 0.00023359 |
| sp\|O75347\|T | 210034589 | 283749404 | 0.47896037 | 1.29E-05 | 0.0002611 |
| sp\|Q96AP7\|E | 12979489.1 | 20071041.2 | 0.62691954 | 1.29E-05 | 0.0002611 |
| sp\|P34931\|H | 2619314604 | 1778324830 | -0.5287973 | 1.29E-05 | 0.0002611 |
| sp\|P20700\|L | 229438402 | 139565058 | -0.7282612 | 1.29E-05 | 0.0002611 |
| sp\|Q6UXG3\|( | 3299400.08 | 8425115.49 | 1.46814476 | 1.29E-05 | 0.0002611 |
| sp\|Q8TD47\|F | 82711475.2 | 29616621.3 | -1.765124 | 1.29E-05 | 0.0002611 |

| | | | | | |
|---|---|---|---|---|---|
| sp\|O00233\|F | 96216239.4 | 145968357 | 0.62007465 | 1.29E-05 | 0.0002611 |
| sp\|Q9Y2T7\|Y | 25217886.3 | 40459035.8 | 0.59894009 | 1.29E-05 | 0.0002611 |
| sp\|Q8WUD1\| | 84008896.8 | 118822915 | 0.48583523 | 1.29E-05 | 0.0002611 |
| sp\|P62081\|R | 90966717.1 | 37401314 | -1.3895979 | 1.29E-05 | 0.0002611 |
| sp\|Q12906\|I | 86016470.1 | 52230694.2 | -0.6786664 | 1.29E-05 | 0.0002611 |
| sp\|Q9NRA2\|S | 447183.778 | 1399494.73 | 1.02690521 | 1.51E-05 | 0.00027837 |
| sp\|P37108\|S | 6555433.79 | 3132443.99 | -1.0767189 | 1.51E-05 | 0.00027837 |
| sp\|Q92747\|A | 72463347.2 | 54870466.8 | -0.4226517 | 1.51E-05 | 0.00027837 |
| sp\|Q9BRF8\|C | 59292948.8 | 102206438 | 0.76363503 | 1.51E-05 | 0.00027837 |
| sp\|Q13332\|F | 37054469.2 | 29190262.4 | -0.3418535 | 1.51E-05 | 0.00027837 |
| sp\|P51610\|H | 28485409.1 | 24293973.1 | -0.2350946 | 1.51E-05 | 0.00027837 |
| sp\|Q96Q06\|I | 1620530855 | 2971011457 | 0.9187423 | 1.51E-05 | 0.00027837 |
| sp\|Q9BPW8\| | 44474407.2 | 25592090.3 | -0.7626312 | 1.51E-05 | 0.00027837 |
| sp\|P62854\|R | 80972480.2 | 27796099.4 | -1.6433899 | 1.51E-05 | 0.00027837 |
| sp\|P27361\|M | 170920205 | 98636973 | -0.876755 | 1.51E-05 | 0.00027837 |
| sp\|Q2MKA7\| | 1050522.17 | 9719690.58 | 2.2059896 | 1.51E-05 | 0.00027837 |
| sp\|O43143\|D | 61106128.3 | 40697749.4 | -0.5692304 | 1.51E-05 | 0.00027837 |
| sp\|Q1KMD3\| | 85946495.3 | 50744946.4 | -0.7599596 | 1.51E-05 | 0.00027837 |
| sp\|Q12905\|I | 45291921.8 | 26042768.1 | -0.7790083 | 1.51E-05 | 0.00027837 |
| sp\|O15231\|Z | 54740063 | 35248503.7 | -0.6612868 | 1.51E-05 | 0.00027837 |
| sp\|O00468\|A | 8046596.31 | 3627608.38 | -1.0593241 | 1.51E-05 | 0.00027837 |
| sp\|Q07507\|D | 43985406.6 | 126572313 | 1.58631076 | 1.51E-05 | 0.00027837 |
| sp\|Q9BXF6\|F | 12549775.3 | 9633642.37 | -0.3986841 | 1.51E-05 | 0.00027837 |
| sp\|P17858\|P | 265417859 | 165414196 | -0.8200504 | 1.51E-05 | 0.00027837 |
| sp\|Q5SSJ5\|H | 21729479.2 | 12200071.1 | -0.8429991 | 1.51E-05 | 0.00027837 |
| sp\|P13716\|H | 1008051261 | 1603086392 | 0.73984992 | 1.51E-05 | 0.00027837 |
| sp\|P15104\|G | 97020150.9 | 185429431 | 0.73302796 | 1.51E-05 | 0.00027837 |
| sp\|Q52LJ0\|F | 9418841.15 | 4714256.29 | -1.0371232 | 1.51E-05 | 0.00027837 |
| sp\|Q15942\|Z | 337945495 | 531459947 | 0.62377422 | 1.77E-05 | 0.00030469 |
| sp\|Q9BWJ5\|S | 4386303.7 | 2657683.17 | -0.8127846 | 1.77E-05 | 0.00030469 |
| sp\|Q8WUF8\| | 3613324.44 | 2377927.26 | -0.6119916 | 1.77E-05 | 0.00030469 |
| sp\|O43390\|H | 278551442 | 145455339 | -0.9594117 | 1.77E-05 | 0.00030469 |
| sp\|Q6UWY5\| | 11942294.5 | 5661688.45 | -0.9403904 | 1.77E-05 | 0.00030469 |
| sp\|P15586\|G | 122194847 | 227277074 | 0.79992384 | 1.77E-05 | 0.00030469 |
| sp\|P15880\|R | 129716799 | 46260199.9 | -1.7663462 | 1.77E-05 | 0.00030469 |
| sp\|Q6QNY1\| | 8968256.38 | 12684968.8 | 0.4530598 | 1.77E-05 | 0.00030469 |
| sp\|Q4KMQ2\| | 14947880.6 | 10662247.3 | -0.4908838 | 1.77E-05 | 0.00030469 |
| sp\|P52272\|H | 143372297 | 79152597.9 | -0.9273144 | 1.77E-05 | 0.00030469 |
| sp\|Q16143\|S | 63948077.4 | 131058729 | 1.0798274 | 1.77E-05 | 0.00030469 |

| | | | | | |
|---|---|---|---|---|---|
| sp\|Q13510\|A | 138646685 | 283021611 | 1.0055984 | 1.77E-05 | 0.00030469 |
| sp\|Q9BVC6\|T | 22558808.1 | 12166039.3 | -1.0925056 | 1.77E-05 | 0.00030469 |
| sp\|P06396\|G | 1475461275 | 2099969580 | 0.4963981 | 1.77E-05 | 0.00030469 |
| sp\|Q16610\|E | 60210476.1 | 146278425 | 1.3827161 | 1.77E-05 | 0.00030469 |
| sp\|P55769\|N | 12315695 | 6197169.04 | -0.9202556 | 1.77E-05 | 0.00030469 |
| sp\|Q96NY7\|C | 33879556.8 | 62069003.7 | 0.82435726 | 1.77E-05 | 0.00030469 |
| sp\|Q86XE5\|H | 2737628.65 | 4844418.22 | 0.78706075 | 2.05E-05 | 0.00033608 |
| sp\|P09455\|R | 12660899.5 | 6505678.98 | -0.9024966 | 2.05E-05 | 0.00033608 |
| sp\|Q9Y2D5\|A | 156388124 | 283958238 | 0.87325 | 2.05E-05 | 0.00033608 |
| sp\|Q9HB19\|I | 233891.065 | 563045.584 | 1.13833163 | 2.05E-05 | 0.00033608 |
| sp\|P10809\|C | 699158655 | 504944295 | -0.4756497 | 2.05E-05 | 0.00033608 |
| sp\|P43251\|B | 46752702.4 | 61344741.2 | 0.4472446 | 2.05E-05 | 0.00033608 |
| sp\|P31947\|1 | 1022292382 | 1374745013 | 0.38180478 | 2.05E-05 | 0.00033608 |
| sp\|P62241\|R | 156856961 | 55977797.6 | -1.6963896 | 2.05E-05 | 0.00033608 |
| sp\|Q8N1G4\| | 46780109.5 | 28535193.3 | -0.738566 | 2.05E-05 | 0.00033608 |
| sp\|Q9Y2T3\|G | 3445906.72 | 8778762.43 | 1.1147198 | 2.05E-05 | 0.00033608 |
| sp\|P11216\|P | 949311430 | 1596486292 | 0.68321574 | 2.05E-05 | 0.00033608 |
| sp\|Q14194\|D | 374386765 | 262252530 | -0.5785173 | 2.05E-05 | 0.00033608 |
| sp\|Q6UXB8\|I | 16774397.8 | 38903894.7 | 1.04839478 | 2.05E-05 | 0.00033608 |
| sp\|Q6UY14\|A | 6245904.54 | 10611868.8 | 0.70710711 | 2.05E-05 | 0.00033608 |
| sp\|P03950\|A | 31307956.8 | 19828892.9 | -0.7925841 | 2.05E-05 | 0.00033608 |
| sp\|P62701\|R | 193786683 | 67018378.2 | -1.9141897 | 2.38E-05 | 0.00036498 |
| sp\|P49903\|S | 29620346.2 | 21649768 | -0.468787 | 2.38E-05 | 0.00036498 |
| sp\|Q08170\|S | 29911730.3 | 18285639.8 | -0.6989953 | 2.38E-05 | 0.00036498 |
| sp\|Q8IXQ3\|C | 7814075.63 | 16966716.1 | 1.04409148 | 2.38E-05 | 0.00036498 |
| sp\|Q9BV79\|M | 6546946.96 | 8630698.54 | 0.41136572 | 2.38E-05 | 0.00036498 |
| sp\|Q86SQ4\|A | 492698.205 | 284898.811 | -0.8661668 | 2.38E-05 | 0.00036498 |
| sp\|Q8NHP8\| | 48473707.7 | 105498157 | 0.97620322 | 2.38E-05 | 0.00036498 |
| sp\|Q9NRV9\|I | 93235473.1 | 145163907 | 0.6773371 | 2.38E-05 | 0.00036498 |
| sp\|P20810\|IC | 338070945 | 476769698 | 0.56116657 | 2.38E-05 | 0.00036498 |
| sp\|Q92769\|H | 26769270.1 | 16217108.6 | -0.7408542 | 2.38E-05 | 0.00036498 |
| sp\|P56199\|IT | 51540213.5 | 81513054.8 | 0.61240135 | 2.38E-05 | 0.00036498 |
| sp\|THIO_HUN | 373624558 | 539526463 | 0.5092395 | 2.38E-05 | 0.00036498 |
| sp\|Q9NRR5\|U | 34043476.4 | 47719794.5 | 0.48040561 | 2.38E-05 | 0.00036498 |
| sp\|Q06278\|A | 906149638 | 485857155 | -1.0597705 | 2.38E-05 | 0.00036498 |
| sp\|Q9HCN4\| | 730650.416 | 1431059.44 | 0.79882764 | 2.38E-05 | 0.00036498 |
| sp\|Q15334\|L | 69602992.6 | 47638455.8 | -0.5583754 | 2.38E-05 | 0.00036498 |
| sp\|Q13867\|B | 26552196 | 41592196.9 | 0.5888831 | 2.38E-05 | 0.00036498 |
| sp\|P12109\|C | 96461878 | 178036343 | 0.7790829 | 2.38E-05 | 0.00036498 |

| | | | | | |
|---|---|---|---|---|---|
| sp\|Q8N392\|[illegible] | 4825825.24 | 2807165.03 | -0.7162421 | 2.38E-05 | 0.00036498 |
| sp\|P25940\|C | 1275985.22 | 3342571.55 | 1.47192208 | 2.38E-05 | 0.00036498 |
| sp\|Q53EL6\|F | 2609432.79 | 1669317.4 | -0.7259047 | 2.75E-05 | 0.00040424 |
| sp\|Q9NP81\|[illegible] | 8889431.28 | 5594638.28 | -0.6918083 | 2.75E-05 | 0.00040424 |
| sp\|Q14117\|[illegible] | 118757397 | 77245111.3 | -0.6210866 | 2.75E-05 | 0.00040424 |
| sp\|P47989\|X | 69931136.7 | 40674534.7 | -0.9842315 | 2.75E-05 | 0.00040424 |
| sp\|Q9BY43\|[illegible] | 86796877.7 | 130416904 | 0.66441598 | 2.75E-05 | 0.00040424 |
| sp\|Q9NS86\|[illegible] | 24273551.2 | 32464364 | 0.4008469 | 2.75E-05 | 0.00040424 |
| sp\|Q6P1J9\|C | 153388.719 | 62483.1397 | -1.3511479 | 2.75E-05 | 0.00040424 |
| sp\|Q13557\|K | 49657074.8 | 28892172.8 | -0.8031172 | 2.75E-05 | 0.00040424 |
| sp\|Q8TD55\|F | 36434902.4 | 50470642.6 | 0.41758916 | 2.75E-05 | 0.00040424 |
| sp\|Q06481\|A | 12077176.3 | 19200111.8 | 0.82329728 | 2.75E-05 | 0.00040424 |
| sp\|Q96AJ9\|V | 144212317 | 222573278 | 1.56943681 | 2.75E-05 | 0.00040424 |
| sp\|P62753\|R | 88550312.7 | 38391271.1 | -1.3382996 | 2.75E-05 | 0.00040424 |
| sp\|Q16629\|S | 25418186.1 | 10957146 | -1.2329226 | 2.75E-05 | 0.00040424 |
| sp\|Q9BRT3\|N | 8319797.53 | 10692438.3 | 0.36933195 | 2.75E-05 | 0.00040424 |
| sp\|P62851\|R | 138814011 | 62490737.5 | -1.3407284 | 3.18E-05 | 0.00044752 |
| sp\|Q2M2I8\|A | 2533762.62 | 3796420.59 | 0.6171896 | 3.18E-05 | 0.00044752 |
| sp\|Q8WVV9\| | 29541999.3 | 16875161.7 | -0.9061825 | 3.18E-05 | 0.00044752 |
| sp\|P09382\|L | 2505819826 | 3461462170 | 0.49575829 | 3.18E-05 | 0.00044752 |
| sp\|Q3SY69\|A | 213479452 | 120447561 | -0.8796652 | 3.18E-05 | 0.00044752 |
| sp\|Q8TAV4\|S | 12712426.7 | 20333626 | 0.73015901 | 3.18E-05 | 0.00044752 |
| sp\|O76031\|[illegible] | 2084533.71 | 5843667.09 | 1.44032589 | 3.18E-05 | 0.00044752 |
| sp\|P17066\|H | 1927256228 | 1446976306 | -0.3575626 | 3.18E-05 | 0.00044752 |
| sp\|P06737\|P | 1924411971 | 3128368339 | 0.60787438 | 3.18E-05 | 0.00044752 |
| sp\|Q08AF3\|S | 609305.204 | 395725.86 | -0.6644152 | 3.18E-05 | 0.00044752 |
| sp\|Q7Z7H3\|[illegible] | 8742089.84 | 14617139 | 0.81940421 | 3.18E-05 | 0.00044752 |
| sp\|P62266\|R | 36798076.2 | 12719851.7 | -1.8834855 | 3.18E-05 | 0.00044752 |
| sp\|Q8N1W1\| | 2906012.75 | 5906982.99 | 1.08909394 | 3.18E-05 | 0.00044752 |
| sp\|Q9NQ39\| | 26893387.8 | 14389709.2 | -0.9538745 | 3.18E-05 | 0.00044752 |
| sp\|Q7Z417\|N | 3930661.41 | 6695871.8 | 0.73328391 | 3.66E-05 | 0.00049664 |
| sp\|Q53H82\|[illegible] | 22620128.6 | 32214795.4 | 0.48826832 | 3.66E-05 | 0.00049664 |
| sp\|P08913\|A | 334280.952 | 803113.032 | 1.29198658 | 3.66E-05 | 0.00049664 |
| sp\|Q12841\|F | 15785909.3 | 30026750.2 | 0.908265 | 3.66E-05 | 0.00049664 |
| sp\|P42330\|A | 1250049308 | 3304296878 | 1.30742463 | 3.66E-05 | 0.00049664 |
| sp\|P05090\|A | 193830713 | 337138718 | 0.85625619 | 3.66E-05 | 0.00049664 |
| sp\|P61026\|R | 124573421 | 154702754 | 0.31188605 | 3.66E-05 | 0.00049664 |
| sp\|O60763\|U | 16140366.6 | 22140993.6 | 0.44576079 | 3.66E-05 | 0.00049664 |
| sp\|P16671\|C | 682488895 | 1415558895 | 1.08347107 | 3.66E-05 | 0.00049664 |

| | | | | | |
|---|---|---|---|---|---|
| sp\|P08708\|R | 46590143.1 | 20549007.4 | -1.3663655 | 3.66E-05 | 0.00049664 |
| sp\|Q16620\|N | 274090.571 | 509187.616 | 0.99319396 | 3.66E-05 | 0.00049664 |
| sp\|P84103\|S | 93466262.2 | 45275616.7 | -0.9999019 | 3.66E-05 | 0.00049664 |
| sp\|P61587\|R | 740934.856 | 1335532.01 | 0.81635237 | 3.66E-05 | 0.00049664 |
| sp\|Q9UNL2\|S | 784808.086 | 407170.957 | -1.6172679 | 4.21E-05 | 0.00054767 |
| sp\|Q6NZI2\|C | 2232328115 | 3640512953 | 0.67911972 | 4.21E-05 | 0.00054767 |
| sp\|P28074\|P | 192016627 | 312986676 | 0.65750117 | 4.21E-05 | 0.00054767 |
| sp\|P62857\|R | 76322990.5 | 40077401.9 | -1.0525544 | 4.21E-05 | 0.00054767 |
| sp\|Q9Y6T7\|D | 1939121.08 | 4011140.99 | 0.35486855 | 4.21E-05 | 0.00054767 |
| sp\|P22307\|S | 99340424.3 | 137522291 | 0.44380234 | 4.21E-05 | 0.00054767 |
| sp\|Q5IJ48\|CI | 1712256.57 | 645689.589 | -1.3466118 | 4.21E-05 | 0.00054767 |
| sp\|P34059\|G | 8676135.32 | 18596715.6 | 0.94979983 | 4.21E-05 | 0.00054767 |
| sp\|Q13595\|T | 10321024.8 | 6660602.08 | -0.6329568 | 4.21E-05 | 0.00054767 |
| sp\|Q96DB5\|I | 22923875 | 34177736.6 | 0.56442452 | 4.21E-05 | 0.00054767 |
| sp\|P28072\|P | 183941255 | 284907188 | 0.59802307 | 4.21E-05 | 0.00054767 |
| sp\|P22570\|A | 43598681 | 27113347.7 | -0.6172164 | 4.21E-05 | 0.00054767 |
| sp\|P12429\|A | 302375362 | 134628598 | -1.2863885 | 4.21E-05 | 0.00054767 |
| sp\|Q08623\|H | 7724740.98 | 5533496.1 | -0.5038323 | 4.21E-05 | 0.00054767 |
| sp\|P47813\|IF | 16553885.3 | 10585641.4 | -0.6978731 | 4.21E-05 | 0.00054767 |
| sp\|P20930\|F | 6664905.73 | 28771829.1 | 1.19856503 | 4.83E-05 | 0.00061001 |
| sp\|P50851\|L | 261390.806 | 476068.06 | 0.72459612 | 4.83E-05 | 0.00061001 |
| sp\|P62244\|R | 130568282 | 54191637.9 | -1.4929972 | 4.83E-05 | 0.00061001 |
| sp\|Q9UK59\|D | 2552008.92 | 1681072.84 | -0.6332795 | 4.83E-05 | 0.00061001 |
| sp\|Q9H299\|S | 489775955 | 884504021 | 0.69547502 | 4.83E-05 | 0.00061001 |
| sp\|P31942\|H | 80943447 | 45452003.7 | -0.8312039 | 4.83E-05 | 0.00061001 |
| sp\|Q9Y3L5\|R | 1427883.9 | 1887623.09 | 0.40483854 | 4.83E-05 | 0.00061001 |
| sp\|P62829\|R | 58583402.3 | 26471586.6 | -1.044998 | 4.83E-05 | 0.00061001 |
| sp\|Q96B97\|S | 51239087.9 | 73514046 | 0.51232719 | 4.83E-05 | 0.00061001 |
| sp\|P13804\|E | 335970947 | 254405847 | -0.471464 | 4.83E-05 | 0.00061001 |
| sp\|P14866\|H | 189080287 | 98726613.6 | -0.9955296 | 4.83E-05 | 0.00061001 |
| sp\|Q9NP90\|I | 14527923 | 21426151.5 | 0.43740425 | 5.53E-05 | 0.00067184 |
| sp\|P56556\|N | 4223460.37 | 2763578.42 | -0.6472844 | 5.53E-05 | 0.00067184 |
| sp\|P00746\|C | 78864807.6 | 128430681 | 0.61081239 | 5.53E-05 | 0.00067184 |
| sp\|O15355\|P | 14242460 | 7923565.03 | -0.8510213 | 5.53E-05 | 0.00067184 |
| sp\|P62834\|R | 127492803 | 192105659 | 0.57429833 | 5.53E-05 | 0.00067184 |
| sp\|O95639\|C | 3093308.8 | 4232381.1 | 0.43454377 | 5.53E-05 | 0.00067184 |
| sp\|P35568\|IF | 2125350.29 | 2972393.31 | 0.45338771 | 5.53E-05 | 0.00067184 |
| sp\|Q14847\|L | 486546423 | 791035864 | 0.66218246 | 5.53E-05 | 0.00067184 |
| sp\|P22676\|C | 380137779 | 582808721 | 0.58888765 | 5.53E-05 | 0.00067184 |

| | | | | | |
|---|---|---|---|---|---|
| sp\|Q96FJ2\|D | 39567242.2 | 30413285.9 | -0.3801037 | 5.53E-05 | 0.00067184 |
| sp\|Q9Y4G6\|T | 110345725 | 141816360 | 0.32877686 | 5.53E-05 | 0.00067184 |
| sp\|P23468\|P | 11418714.3 | 7071733.78 | -0.682821 | 5.53E-05 | 0.00067184 |
| sp\|Q15181\|I | 252992616 | 400763661 | 0.63140553 | 5.53E-05 | 0.00067184 |
| sp\|Q8N5K1\|( | 987324.18 | 717127.515 | -0.7514312 | 5.53E-05 | 0.00067184 |
| sp\|O00142\|K | 35327293.1 | 49196222.9 | 0.48596568 | 5.53E-05 | 0.00067184 |
| sp\|O43920\|N | 3925263.04 | 2438721.48 | -0.653992 | 6.32E-05 | 0.00073594 |
| sp\|Q27J81\|IN | 9500482.09 | 13286195.7 | 0.46062665 | 6.32E-05 | 0.00073594 |
| sp\|Q5VUM1\| | 629128.265 | 1290779.82 | 1.07204499 | 6.32E-05 | 0.00073594 |
| sp\|Q9UJJ9\|G | 564446.073 | 806237.023 | 0.55308142 | 6.32E-05 | 0.00073594 |
| sp\|Q86Y82\|S | 38686078.3 | 52348567 | 0.41661808 | 6.32E-05 | 0.00073594 |
| sp\|Q9UGI8\|T | 125614478 | 83159882.3 | -0.5519055 | 6.32E-05 | 0.00073594 |
| sp\|O00192\|A | 2782083.52 | 5902059.04 | 1.0105019 | 6.32E-05 | 0.00073594 |
| sp\|P61224\|R | 133918703 | 200331531 | 0.55288348 | 6.32E-05 | 0.00073594 |
| sp\|Q9Y2V2\|( | 86100240.8 | 63868053.1 | -0.4287217 | 6.32E-05 | 0.00073594 |
| sp\|O60784\|T | 18945035.6 | 26468034.1 | 0.45069038 | 6.32E-05 | 0.00073594 |
| sp\|Q14195\|D | 1092978994 | 755494974 | -0.6256769 | 6.32E-05 | 0.00073594 |
| sp\|P48163\|M | 469647641 | 868361385 | 0.6771515 | 6.32E-05 | 0.00073594 |
| sp\|P46781\|R | 226819004 | 87676984.7 | -1.755065 | 6.32E-05 | 0.00073594 |
| sp\|Q96IU4\|A | 225383355 | 146632298 | -0.7580181 | 6.32E-05 | 0.00073594 |
| sp\|P31323\|K | 223721498 | 124950335 | -0.9622618 | 6.32E-05 | 0.00073594 |
| sp\|Q9UNZ5\|I | 4580128.84 | 1546825.29 | -1.608329 | 6.32E-05 | 0.00073594 |
| sp\|P04179\|S | 792449331 | 1543801561 | 0.78472206 | 6.32E-05 | 0.00073594 |
| sp\|Q9BV10\|A | 117199.897 | 23590.8775 | -9.4651537 | 6.42E-05 | 0.00074654 |
| sp\|P62070\|R | 107303741 | 173358145 | 0.62639476 | 7.20E-05 | 0.00079851 |
| sp\|P04843\|R | 64686476.8 | 34881843.7 | -1.053114 | 7.20E-05 | 0.00079851 |
| sp\|Q13907\|I | 5125415.84 | 9933858.9 | 0.89343996 | 7.20E-05 | 0.00079851 |
| sp\|O95980\|F | 606897.443 | 331263.495 | -0.9316814 | 7.20E-05 | 0.00079851 |
| sp\|Q16853\|A | 2814688963 | 4474815195 | 0.7019802 | 7.20E-05 | 0.00079851 |
| sp\|O75368\|S | 160797480 | 235599374 | 0.45782337 | 7.20E-05 | 0.00079851 |
| sp\|P62847\|R | 48734514 | 18621620.5 | -1.5645881 | 7.20E-05 | 0.00079851 |
| sp\|Q9UPZ6\|T | 18058439.6 | 13153974.3 | -0.4892693 | 7.20E-05 | 0.00079851 |
| sp\|P11279\|L | 78850212.1 | 121336653 | 0.53939897 | 7.20E-05 | 0.00079851 |
| sp\|P08473\|N | 1317052.02 | 3598907.76 | 1.6519039 | 7.20E-05 | 0.00079851 |
| sp\|O94905\|E | 28687490.4 | 17862737.7 | -0.7732797 | 7.20E-05 | 0.00079851 |
| sp\|P83731\|R | 36426257.9 | 18215874.5 | -0.9851142 | 7.20E-05 | 0.00079851 |
| sp\|P09525\|A | 63664700.7 | 41444765.7 | -0.6348233 | 7.20E-05 | 0.00079851 |
| sp\|A6NIH7\|U | 11175134.4 | 7568251.65 | -0.5812198 | 7.20E-05 | 0.00079851 |
| sp\|Q09666\|A | 4666761332 | 7075937550 | 0.57245384 | 7.20E-05 | 0.00079851 |

| | | | | | |
|---|---|---|---|---|---|
| sp\|Q7Z4H8\|F | 85252240.2 | 51100775 | -0.8710335 | 7.20E-05 | 0.00079851 |
| sp\|Q13522\|F | 14106073.7 | 21492463.9 | 0.6149214 | 7.20E-05 | 0.00079851 |
| sp\|P30101\|P | 908726723 | 588421028 | -0.6941652 | 7.20E-05 | 0.00079851 |
| sp\|P16144\|IT | 9898542.93 | 6606868.22 | -0.64158 | 7.20E-05 | 0.00079851 |
| sp\|P15531\|N | 1030871323 | 1442746557 | 0.44727839 | 7.20E-05 | 0.00079851 |
| sp\|Q9NZB2\|F | 58140042.8 | 41621342.2 | -0.4756122 | 8.20E-05 | 0.00087079 |
| sp\|Q15046\|S | 192198730 | 137834052 | -0.4776 | 8.20E-05 | 0.00087079 |
| sp\|P27695\|A | 52543265.6 | 28881108.4 | -1.0523701 | 8.20E-05 | 0.00087079 |
| sp\|Q9BS40\|L | 41666223.2 | 21389059.1 | -1.0100161 | 8.20E-05 | 0.00087079 |
| sp\|Q6ZMP0\|T | 236220.63 | 386023.297 | -2.4977554 | 8.20E-05 | 0.00087079 |
| sp\|P27635\|R | 57170269.3 | 18819812.4 | -1.6087923 | 8.20E-05 | 0.00087079 |
| sp\|Q92597\|N | 33895662.5 | 20070056.9 | -0.7176497 | 8.20E-05 | 0.00087079 |
| sp\|P62273\|R | 46355040 | 20185659.5 | -1.6617699 | 8.20E-05 | 0.00087079 |
| sp\|P12235\|A | 83449154 | 52260456.9 | -0.6592582 | 8.20E-05 | 0.00087079 |
| sp\|O76021\|F | 4543236.92 | 2540439.84 | -0.7841221 | 8.20E-05 | 0.00087079 |
| sp\|P07741\|A | 103059367 | 60969507.7 | -0.7859642 | 8.20E-05 | 0.00087079 |
| sp\|Q99436\|F | 58649238.6 | 86098439 | 0.51122531 | 8.20E-05 | 0.00087079 |
| sp\|Q8IXS6\|P | 47055234.7 | 81077634.7 | 0.77187329 | 8.20E-05 | 0.00087079 |
| sp\|P00846\|A | 764329.671 | 423022.659 | -0.9760741 | 8.20E-05 | 0.00087079 |
| sp\|Q9NX24\|I | 874330.788 | 348709.009 | -1.3344393 | 8.20E-05 | 0.00087079 |
| sp\|Q8TEP8\|C | 13917861 | 19612922.5 | 0.56562078 | 8.20E-05 | 0.00087079 |
| sp\|A1L4H1\|S | 5523069.54 | 12171123.1 | 0.78813105 | 8.20E-05 | 0.00087079 |
| sp\|P49458\|S | 38622626.3 | 25366683.3 | -0.6400489 | 8.20E-05 | 0.00087079 |
| sp\|Q15848\|A | 10120940.6 | 17965600.3 | 0.75969034 | 8.20E-05 | 0.00087079 |
| sp\|Q9UEU0\|V | 8370747.01 | 12195259.4 | 0.50193462 | 9.32E-05 | 0.00095766 |
| sp\|Q08397\|L | 747326.143 | 1735078.68 | 1.15191846 | 9.32E-05 | 0.00095766 |
| sp\|Q9NQ66\| | 1862275.16 | 1337314.72 | -0.4770106 | 9.32E-05 | 0.00095766 |
| sp\|P39019\|R | 187758116 | 99175466.4 | -1.0391465 | 9.32E-05 | 0.00095766 |
| sp\|P0DMM9\| | 102581706 | 60154515.8 | -0.7797963 | 9.32E-05 | 0.00095766 |
| sp\|P26639\|S | 61397391.7 | 39560463.1 | -0.6289852 | 9.32E-05 | 0.00095766 |
| sp\|Q9GZT8\|N | 74512338 | 99290412.4 | 0.39405116 | 9.32E-05 | 0.00095766 |
| sp\|Q13449\|L | 5089814.47 | 2492869.4 | -1.1152735 | 9.32E-05 | 0.00095766 |
| sp\|P11532\|D | 9438674.85 | 12960674.8 | 0.45903018 | 9.32E-05 | 0.00095766 |
| sp\|P45880\|V | 52639756.3 | 34814769.6 | -0.6131243 | 9.32E-05 | 0.00095766 |
| sp\|P46783\|R | 66155713.9 | 31154516.6 | -1.1647593 | 9.32E-05 | 0.00095766 |
| sp\|P61088\|U | 29851704.7 | 43940146.4 | 0.4955482 | 9.32E-05 | 0.00095766 |
| sp\|P25398\|R | 108234602 | 58349768.9 | -0.9257132 | 9.32E-05 | 0.00095766 |
| sp\|P23246\|S | 148490718 | 96015312.3 | -0.6352493 | 9.32E-05 | 0.00095766 |
| sp\|Q969Z3\|N | 29121860.8 | 21893093.5 | -0.4226856 | 9.32E-05 | 0.00095766 |

| | | | | | |
|---|---|---|---|---|---|
| sp\|B7ZW38\|I | 116122909 | 43224755.6 | -1.4486163 | 0.0001057 | 0.0010459 |
| sp\|Q92611\|E | 2993562.95 | 1826906.11 | -0.9423316 | 0.0001057 | 0.0010459 |
| sp\|Q9P2M7\|( | 614150.763 | 439736.855 | -1.1103635 | 0.0001057 | 0.0010459 |
| sp\|Q14244\|M | 3267797.35 | 5180690.59 | 0.71729121 | 0.0001057 | 0.0010459 |
| sp\|Q9UHB6\| | 60014317.2 | 89128870.5 | 0.49934772 | 0.0001057 | 0.0010459 |
| sp\|P05141\|A | 132595536 | 88617246.8 | -0.5845496 | 0.0001057 | 0.0010459 |
| sp\|Q8IVN3\|M | 4090147.17 | 1993953.8 | -0.9324871 | 0.0001057 | 0.0010459 |
| sp\|Q9Y3I0\|R | 85163322.8 | 57742917.8 | -0.5600839 | 0.0001057 | 0.0010459 |
| sp\|Q969Q0\|I | 12061506.2 | 3630693.42 | -1.8535869 | 0.0001057 | 0.0010459 |
| sp\|Q96CT7\|( | 10355048.4 | 16704135.2 | 0.65480513 | 0.0001057 | 0.0010459 |
| sp\|P49590\|S | 50220400.2 | 24925109.1 | -1.170314 | 0.0001057 | 0.0010459 |
| sp\|P61106\|R | 105701831 | 137848962 | 0.37845319 | 0.0001057 | 0.0010459 |
| sp\|P83881\|R | 10373715.7 | 3905019.25 | -1.4516282 | 0.0001057 | 0.0010459 |
| sp\|O43865\|S | 91436523.7 | 50872326.9 | -0.8579398 | 0.0001057 | 0.0010459 |
| sp\|P16284\|P | 49739078.7 | 68868297.7 | 0.44572195 | 0.0001057 | 0.0010459 |
| sp\|P54725\|R | 98346602.8 | 130356599 | 0.44147527 | 0.0001057 | 0.0010459 |
| sp\|Q9NP74\|I | 81392452.1 | 139715616 | 0.75733245 | 0.0001057 | 0.0010459 |
| sp\|O75113\|M | 2375615.12 | 3782129.95 | 0.67463836 | 0.0001057 | 0.0010459 |
| sp\|O00217\|M | 6661942.33 | 4391054.03 | -0.5600818 | 0.0001197 | 0.00113752 |
| sp\|P28482\|M | 124045582 | 90672442.4 | -0.5354999 | 0.0001197 | 0.00113752 |
| sp\|P13010\|X | 128488082 | 69891439.9 | -0.8783426 | 0.0001197 | 0.00113752 |
| sp\|O94885\|S | 3065085.11 | 4431892.42 | 0.46990588 | 0.0001197 | 0.00113752 |
| sp\|Q7Z7G0\|T | 20221074.3 | 50256232.9 | 0.94382209 | 0.0001197 | 0.00113752 |
| sp\|Q15185\|T | 76023400.6 | 46615287.1 | -0.7733648 | 0.0001197 | 0.00113752 |
| sp\|P22090\|R | 44074234.5 | 17066300.9 | -1.6235047 | 0.0001197 | 0.00113752 |
| sp\|Q14CZ8\|H | 28315184.7 | 46848118.6 | 0.73280979 | 0.0001197 | 0.00113752 |
| sp\|P62249\|R | 134208909 | 55879805.5 | -1.5503469 | 0.0001197 | 0.00113752 |
| sp\|Q8NBJ7\|S | 52556115.6 | 70051788.6 | 0.36978702 | 0.0001197 | 0.00113752 |
| sp\|Q13740\|( | 11817992.8 | 20130845.2 | 0.62741372 | 0.0001197 | 0.00113752 |
| sp\|Q8IWS0\|F | 754797.151 | 2234772.61 | 1.13511806 | 0.0001197 | 0.00113752 |
| sp\|P78324\|S | 11726233.2 | 17530982.2 | 0.44777163 | 0.0001197 | 0.00113752 |
| sp\|P61769\|B | 81078352.3 | 119827132 | 0.5180984 | 0.0001197 | 0.00113752 |
| sp\|P29590\|P | 33222524.4 | 41781789.6 | 0.30759144 | 0.0001197 | 0.00113752 |
| sp\|P15144\|A | 43141441.4 | 60392060.4 | 0.45601328 | 0.0001197 | 0.00113752 |
| sp\|Q9P266\|J | 5302119.53 | 7596867.76 | 0.449028 | 0.0001197 | 0.00113752 |
| sp\|O00244\|A | 24005934.3 | 40480304.6 | 0.68548285 | 0.0001197 | 0.00113752 |
| sp\|P00441\|S | 287649437 | 409331588 | 0.51849087 | 0.0001197 | 0.00113752 |
| sp\|Q8NG11\|T | 355146.591 | 238932.132 | -0.6012271 | 0.0001197 | 0.00113752 |
| sp\|Q07157\|Z | 505678971 | 433671355 | -0.2191039 | 0.00013533 | 0.00123936 |

| | | | | | |
|---|---|---|---|---|---|
| sp\|Q9Y2X3\|N | 16820116.7 | 11261540.4 | -0.6657661 | 0.00013533 | 0.00123936 |
| sp\|P16070\|C | 91641125 | 157634166 | 0.65710928 | 0.00013533 | 0.00123936 |
| sp\|P30533\|A | 37082920.2 | 53529682.7 | 0.4737219 | 0.00013533 | 0.00123936 |
| sp\|Q15276\|F | 49583483.2 | 61439347.7 | 0.30089646 | 0.00013533 | 0.00123936 |
| sp\|P51687\|S | 62979632.9 | 113543911 | 0.57608423 | 0.00013533 | 0.00123936 |
| sp\|Q8N142\|I | 11434613.1 | 5323315.06 | -1.2571102 | 0.00013533 | 0.00123936 |
| sp\|P11277\|S | 151151387 | 357998001 | 0.88855753 | 0.00013533 | 0.00123936 |
| sp\|P67870\|C | 30578257.6 | 21687308.9 | -0.5434287 | 0.00013533 | 0.00123936 |
| sp\|P02656\|A | 33621983.1 | 62085429.1 | 0.64381854 | 0.00013533 | 0.00123936 |
| sp\|P02649\|A | 47294343.9 | 110909626 | 1.07863891 | 0.00013533 | 0.00123936 |
| sp\|Q9Y580\|F | 21042731.8 | 35780165.8 | 0.75199854 | 0.00013533 | 0.00123936 |
| sp\|P48449\|L | 15936154.9 | 23771931.6 | 0.48980714 | 0.00013533 | 0.00123936 |
| sp\|P49247\|R | 208165865 | 350361623 | 0.79921121 | 0.00013533 | 0.00123936 |
| sp\|P62820\|R | 181248136 | 233842195 | 0.36309003 | 0.00013533 | 0.00123936 |
| sp\|Q15075\|E | 305798634 | 444231842 | 0.50194154 | 0.00013533 | 0.00123936 |
| sp\|P23396\|R | 596050181 | 265792152 | -1.4374218 | 0.00013533 | 0.00123936 |
| sp\|P49792\|R | 27662336.6 | 20123011.9 | -0.4961115 | 0.00013533 | 0.00123936 |
| sp\|P54727\|R | 319457202 | 481804043 | 0.56843683 | 0.00013533 | 0.00123936 |
| sp\|Q96GA7\|S | 12661339.2 | 27460836.5 | 0.96236552 | 0.00015274 | 0.0013474 |
| sp\|P26599\|P | 160896593 | 104311353 | -0.6588896 | 0.00015274 | 0.0013474 |
| sp\|P15289\|A | 31706479 | 54886223.9 | 0.78019996 | 0.00015274 | 0.0013474 |
| sp\|Q9BQB6\|V | 2005655.03 | 1089980.63 | -1.288435 | 0.00015274 | 0.0013474 |
| sp\|Q13151\|F | 70046082.6 | 37781615.6 | -0.8942431 | 0.00015274 | 0.0013474 |
| sp\|P07910\|H | 300270467 | 114294356 | -1.3924665 | 0.00015274 | 0.0013474 |
| sp\|P51659\|D | 104958384 | 146295278 | 0.41327081 | 0.00015274 | 0.0013474 |
| sp\|P62269\|R | 256342187 | 112593139 | -1.5390509 | 0.00015274 | 0.0013474 |
| sp\|Q9UM47\| | 3226119.48 | 5334714.86 | 0.7339011 | 0.00015274 | 0.0013474 |
| sp\|Q9BXK5\|E | 861841.719 | 1557301.17 | 0.77146646 | 0.00015274 | 0.0013474 |
| sp\|P39687\|A | 73443283.2 | 39118837.5 | -0.9754919 | 0.00015274 | 0.0013474 |
| sp\|Q9NP72\|I | 7339220.31 | 11258342.1 | 0.55755813 | 0.00015274 | 0.0013474 |
| sp\|P55083\|M | 66625677.2 | 126622046 | 0.79309262 | 0.00015274 | 0.0013474 |
| sp\|P55036\|P | 74039309.1 | 90836354 | 0.28553678 | 0.00015274 | 0.0013474 |
| sp\|Q04828\|A | 1486562779 | 3552109908 | 1.15218345 | 0.00015274 | 0.0013474 |
| sp\|P46782\|R | 158553001 | 64449179.2 | -1.7215554 | 0.00015274 | 0.0013474 |
| sp\|Q9NWU5\| | 2693532.96 | 3735724.41 | 0.48023423 | 0.00015274 | 0.0013474 |
| sp\|Q96C01\|F | 5209405.82 | 7328131.96 | 0.45851405 | 0.00015274 | 0.0013474 |
| sp\|Q9UKM9\| | 62869726.9 | 35949255.7 | -0.7679937 | 0.00015274 | 0.0013474 |
| sp\|P04080\|C | 323503508 | 490693147 | 0.52769785 | 0.00015274 | 0.0013474 |
| sp\|P09012\|S | 30917915.1 | 23812282 | -0.4082239 | 0.00017213 | 0.00147247 |

| | | | | | |
|---|---|---|---|---|---|
| sp\|Q969E4\|T | 29179427.1 | 15092804.1 | -1.0926389 | 0.00017213 | 0.00147247 |
| sp\|P11234\|R | 18849599.7 | 25481230.2 | 0.43273768 | 0.00017213 | 0.00147247 |
| sp\|P08572\|C | 38141332.3 | 69341763.3 | 0.89831165 | 0.00017213 | 0.00147247 |
| sp\|Q9Y547\|I | 1427702.4 | 2565355.72 | 0.73419632 | 0.00017213 | 0.00147247 |
| sp\|P02747\|C | 33752741.9 | 16434739.4 | -0.9458079 | 0.00017213 | 0.00147247 |
| sp\|P02655\|A | 8882834.09 | 19865865.3 | 0.75180547 | 0.00017213 | 0.00147247 |
| sp\|Q9Y285\|S | 7224832.87 | 4006975.83 | -0.7871703 | 0.00017213 | 0.00147247 |
| sp\|P30049\|A | 108542656 | 166105882 | 0.6532695 | 0.00017213 | 0.00147247 |
| sp\|Q15599\|N | 42007213.9 | 56131948.6 | 0.44368723 | 0.00017213 | 0.00147247 |
| sp\|P59768\|G | 10874278.2 | 19931333.2 | 0.90576778 | 0.00017213 | 0.00147247 |
| sp\|Q9UMX0\| | 53092401 | 74286502.3 | 0.48581014 | 0.00017213 | 0.00147247 |
| sp\|P61927\|R | 1694599.81 | 982241.808 | -0.852331 | 0.00017213 | 0.00147247 |
| sp\|P07358\|C | 121752371 | 64092855.9 | -1.243442 | 0.00017213 | 0.00147247 |
| sp\|Q14011\|C | 36881520 | 70733964.7 | 0.74048157 | 0.00017213 | 0.00147247 |
| sp\|O95628\|C | 3127499.4 | 4176881.97 | 0.40152477 | 0.00017213 | 0.00147247 |
| sp\|O95817\|E | 100103619 | 151471377 | 0.58706093 | 0.00017213 | 0.00147247 |
| sp\|P67812\|S | 1510573.12 | 749683.136 | -1.4955925 | 0.00019369 | 0.00160535 |
| sp\|P63279\|U | 27603353.8 | 17098705.9 | -0.7422665 | 0.00019369 | 0.00160535 |
| sp\|Q96QK1\|V | 97248404.6 | 54805308 | -0.8566941 | 0.00019369 | 0.00160535 |
| sp\|P50440\|G | 40999430.2 | 28807740.2 | -0.5484565 | 0.00019369 | 0.00160535 |
| sp\|P08574\|C | 21178147.2 | 13044090.2 | -0.6704725 | 0.00019369 | 0.00160535 |
| sp\|Q14247\|S | 246955508 | 369161162 | 0.53900398 | 0.00019369 | 0.00160535 |
| sp\|Q8N668\|C | 4810346.69 | 6325198.24 | 0.37918486 | 0.00019369 | 0.00160535 |
| sp\|P06733\|E | 6109799186 | 9374886684 | 0.47051539 | 0.00019369 | 0.00160535 |
| sp\|Q9BQI4\|C | 522371.474 | 261764.277 | -1.0459443 | 0.00019369 | 0.00160535 |
| sp\|Q07955\|S | 81477435.2 | 46015648.2 | -0.7875237 | 0.00019369 | 0.00160535 |
| sp\|Q9P2R7\|S | 251166042 | 161294755 | -0.7450488 | 0.00019369 | 0.00160535 |
| sp\|P19838\|N | 27483609.6 | 19489401.2 | -0.4551141 | 0.00019369 | 0.00160535 |
| sp\|Q9Y3F4\|S | 65201309.4 | 86455093.6 | 0.34223926 | 0.00019369 | 0.00160535 |
| sp\|Q15287\|F | 11989085.7 | 6975363.7 | -0.8705885 | 0.00019369 | 0.00160535 |
| sp\|Q01105\|S | 22720091.2 | 11404617.5 | -1.1329139 | 0.00019369 | 0.00160535 |
| sp\|P55957\|B | 769604.4 | 1883997.61 | 1.03838932 | 0.00019369 | 0.00160535 |
| sp\|P05230\|F | 2739865.11 | 902070.629 | -1.7939011 | 0.00019369 | 0.00160535 |
| sp\|Q96SB3\|N | 8541462.75 | 11348510.8 | 0.409194 | 0.00019369 | 0.00160535 |
| sp\|P07237\|P | 425937964 | 577041233 | 0.39302442 | 0.00021761 | 0.00174633 |
| sp\|O00625\|F | 49658104.3 | 68182105.3 | 0.48057502 | 0.00021761 | 0.00174633 |
| sp\|Q8NBK3\|S | 6681824.96 | 10103619.5 | 0.47443024 | 0.00021761 | 0.00174633 |
| sp\|P46776\|R | 19114213.1 | 6111625.4 | -1.8077603 | 0.00021761 | 0.00174633 |
| sp\|P30876\|R | 8943891.41 | 6160942.92 | -0.5112814 | 0.00021761 | 0.00174633 |

| | | | | | |
|---|---|---|---|---|---|
| sp\|P25789\|P | 342843182 | 450308730 | 0.338615 | 0.00021761 | 0.00174633 |
| sp\|O00186\|S | 715309.079 | 410870.547 | -1.646855 | 0.00021761 | 0.00174633 |
| sp\|O95433\|A | 1539331.13 | 885825.595 | -0.9034225 | 0.00021761 | 0.00174633 |
| sp\|O75094\|S | 5573091.66 | 3587607.8 | -0.5887129 | 0.00021761 | 0.00174633 |
| sp\|O14879\|I | 3224609.36 | 1881180.25 | -0.760338 | 0.00021761 | 0.00174633 |
| sp\|P19338\|N | 287408039 | 172305164 | -0.8206798 | 0.00021761 | 0.00174633 |
| sp\|P38117\|E | 617595539 | 484823533 | -0.3768403 | 0.00021761 | 0.00174633 |
| sp\|Q14108\|S | 71901776.7 | 98699289 | 0.44122099 | 0.00021761 | 0.00174633 |
| sp\|Q13790\|A | 373032.321 | 809657.894 | 1.24723121 | 0.00021761 | 0.00174633 |
| sp\|Q15020\|S | 906847.317 | 502922.042 | -0.8211728 | 0.00021761 | 0.00174633 |
| sp\|P62906\|R | 54327725.8 | 19602729.8 | -1.643176 | 0.00021761 | 0.00174633 |
| sp\|P26640\|S | 82857305.7 | 52284318.5 | -0.629098 | 0.00021761 | 0.00174633 |
| sp\|Q03518\|T | 29932561.3 | 47745220.7 | 0.52358228 | 0.00021761 | 0.00174633 |
| sp\|P25787\|P | 303707750 | 422054765 | 0.40294731 | 0.00021761 | 0.00174633 |
| sp\|Q96QR8\|I | 11397147.3 | 8541656.53 | -0.41888 | 0.00024414 | 0.00191131 |
| sp\|P12236\|A | 116291585 | 76626917.3 | -0.5931461 | 0.00024414 | 0.00191131 |
| sp\|P43243\|M | 30670833.2 | 17323404.6 | -0.7667184 | 0.00024414 | 0.00191131 |
| sp\|Q16643\|D | 41913413.9 | 58261341.1 | 0.43224531 | 0.00024414 | 0.00191131 |
| sp\|P19623\|S | 44039079 | 29840505.6 | -0.7164837 | 0.00024414 | 0.00191131 |
| sp\|Q9NUQ9\| | 2670900.29 | 3615509.3 | 0.43474816 | 0.00024414 | 0.00191131 |
| sp\|Q9BVM4\| | 87888606.2 | 113185698 | 0.36194607 | 0.00024414 | 0.00191131 |
| sp\|Q9NUJ1\|A | 13729527.3 | 8218008.17 | -0.7046914 | 0.00024414 | 0.00191131 |
| sp\|Q96A99\|F | 594723.337 | 981720.349 | 0.70388239 | 0.00024414 | 0.00191131 |
| sp\|P36222\|C | 2875419.65 | 7293859.47 | 0.98670484 | 0.00024414 | 0.00191131 |
| sp\|Q9BZK7\|T | 8025019.1 | 6293054.91 | -0.3742049 | 0.00024414 | 0.00191131 |
| sp\|Q16762\|T | 165886969 | 88189671.6 | -1.1281071 | 0.00024414 | 0.00191131 |
| sp\|P40429\|R | 65787678.9 | 23385805.1 | -1.650704 | 0.00024414 | 0.00191131 |
| sp\|Q9UKF6\|C | 1373294.27 | 3903602.15 | 1.0315157 | 0.00024414 | 0.00191131 |
| sp\|Q99447\|F | 144430044 | 113833194 | -0.4077899 | 0.00024414 | 0.00191131 |
| sp\|P49411\|E | 143690016 | 91867153.2 | -0.7576528 | 0.00027351 | 0.00208677 |
| sp\|P35268\|R | 47407927.2 | 29932288.7 | -0.722129 | 0.00027351 | 0.00208677 |
| sp\|P08621\|R | 81752685.4 | 53960098.8 | -0.5769348 | 0.00027351 | 0.00208677 |
| sp\|P08579\|R | 8358829.06 | 5842876.09 | -0.6949368 | 0.00027351 | 0.00208677 |
| sp\|Q969R8\|I | 1074313.15 | 657295.832 | -0.7297041 | 0.00027351 | 0.00208677 |
| sp\|Q14118\|D | 17424343.8 | 28280901.5 | 0.71283171 | 0.00027351 | 0.00208677 |
| sp\|Q9UMX5\| | 24905515.5 | 36466415.7 | 0.50847712 | 0.00027351 | 0.00208677 |
| sp\|P60866\|R | 21870390.7 | 9946204.38 | -1.2866018 | 0.00027351 | 0.00208677 |
| sp\|P60900\|P | 431312312 | 592597436 | 0.38539209 | 0.00027351 | 0.00208677 |
| sp\|P62995\|T | 48183029.1 | 32077648.7 | -0.6038545 | 0.00027351 | 0.00208677 |

| | | | | | |
|---|---|---|---|---|---|
| sp\|P15088\|C | 175141687 | 96010103.1 | -0.9470071 | 0.00027351 | 0.00208677 |
| sp\|Q86SE5\|F | 17489921.1 | 7100670.5 | -1.3743639 | 0.00027351 | 0.00208677 |
| sp\|P29401\|T | 4994502328 | 8373343279 | 0.64646483 | 0.00027351 | 0.00208677 |
| sp\|Q8TC44\|F | 6527319.46 | 10239328.1 | 0.53402856 | 0.00027351 | 0.00208677 |
| sp\|P62917\|R | 91150105.1 | 38097522.8 | -1.2096958 | 0.00027351 | 0.00208677 |
| sp\|P25786\|P | 826575253 | 1160200045 | 0.4116574 | 0.00027351 | 0.00208677 |
| sp\|P00492\|H | 105617292 | 170466987 | 0.65347603 | 0.00030598 | 0.00225561 |
| sp\|P12956\|X | 85129292.2 | 51100301.8 | -0.739409 | 0.00030598 | 0.00225561 |
| sp\|Q96CM8\| | 59410560.1 | 39115890.8 | -0.5917771 | 0.00030598 | 0.00225561 |
| sp\|O14818\|F | 277227726 | 376104136 | 0.36220952 | 0.00030598 | 0.00225561 |
| sp\|P61513\|R | 12178948.3 | 5040054.14 | -1.1840176 | 0.00030598 | 0.00225561 |
| sp\|Q6WCQ1\| | 14309478.3 | 18221839.1 | 0.33547082 | 0.00030598 | 0.00225561 |
| sp\|Q8TE76\|M | 2455939.64 | 4227732.45 | 0.61338949 | 0.00030598 | 0.00225561 |
| sp\|P25788\|P | 345327052 | 469907412 | 0.37851133 | 0.00030598 | 0.00225561 |
| sp\|A4FU69\|E | 933565.082 | 2541958.69 | 1.20393269 | 0.00030598 | 0.00225561 |
| sp\|Q9HCE6\|. | 527109.512 | 918635.326 | 0.72175967 | 0.00030598 | 0.00225561 |
| sp\|P46778\|R | 44787983.8 | 15270437.5 | -1.4310121 | 0.00030598 | 0.00225561 |
| sp\|P12821\|A | 7600828.33 | 10404083.3 | 0.44410486 | 0.00030598 | 0.00225561 |
| sp\|P62841\|R | 54830625.8 | 26252292.5 | -1.4137496 | 0.00030598 | 0.00225561 |
| sp\|Q9H3M7\| | 355788.897 | 148371.338 | -1.2019855 | 0.00030598 | 0.00225561 |
| sp\|Q9BRP8\|F | 47667086.3 | 71884605 | 0.49936119 | 0.00030598 | 0.00225561 |
| sp\|Q13492\|F | 10380396.6 | 12723661.4 | 0.29411589 | 0.00030598 | 0.00225561 |
| sp\|O00483\|N | 14048834.6 | 9077609.52 | -0.6670902 | 0.00030598 | 0.00225561 |
| sp\|P61129\|Z | 2455939.64 | 4227732.45 | 0.61338949 | 0.00030598 | 0.00225561 |
| sp\|Q9NZN4\|I | 867416385 | 1341361727 | 0.51067942 | 0.00030598 | 0.00225561 |
| sp\|P02533\|K | 337925323 | 129121740 | -1.293685 | 0.00030598 | 0.00225561 |
| sp\|P10636\|T | 57628744 | 91048665.4 | 0.71306605 | 0.00030598 | 0.00225561 |
| sp\|P0DP24\|C | 1312246467 | 1953874567 | 0.49736388 | 0.00030598 | 0.00225561 |
| sp\|P11117\|P | 30029718.8 | 47970562.6 | 0.60771357 | 0.00034185 | 0.00243763 |
| sp\|P15529\|M | 4823198.34 | 7522952.55 | 0.59446304 | 0.00034185 | 0.00243763 |
| sp\|Q9NSD9\|: | 420763178 | 256854867 | -0.6548156 | 0.00034185 | 0.00243763 |
| sp\|O60488\|A | 4406913.86 | 5597495.76 | 0.28238544 | 0.00034185 | 0.00243763 |
| sp\|P13667\|P | 224996515 | 128876395 | -0.8914838 | 0.00034185 | 0.00243763 |
| sp\|Q96RS6\|N | 8976258.31 | 6784455.84 | -0.4829405 | 0.00034185 | 0.00243763 |
| sp\|P27816\|M | 1043531715 | 1727865346 | 0.63350123 | 0.00034185 | 0.00243763 |
| sp\|Q9H0X4\|I | 4469412.14 | 6144817.81 | 0.51436728 | 0.00034185 | 0.00243763 |
| sp\|P31150\|G | 286191463 | 158567596 | -0.9343497 | 0.00034185 | 0.00243763 |
| sp\|P07814\|S | 117129436 | 78864576.8 | -0.6045732 | 0.00034185 | 0.00243763 |
| sp\|Q8N163\|( | 36685681.8 | 26012767 | -0.4675506 | 0.00034185 | 0.00243763 |

| | | | | | |
|---|---|---|---|---|---|
| sp\|P12081\|H | 46370493.1 | 25781686 | -0.9134944 | 0.00034185 | 0.00243763 |
| sp\|P51532\|S | 2276972.45 | 1087482.78 | -1.1063921 | 0.00034185 | 0.00243763 |
| sp\|P12107\|C | 792061.214 | 2948331.64 | 2.17157861 | 0.00034185 | 0.00243763 |
| sp\|P32969\|R | 80218032.3 | 36797757 | -1.0610135 | 0.00034185 | 0.00243763 |
| sp\|P48059\|L | 164251968 | 217680475 | 0.3703611 | 0.00034185 | 0.00243763 |
| sp\|P30622\|C | 219195868 | 306193469 | 0.45732055 | 0.00034185 | 0.00243763 |
| sp\|Q8IXL7\|M | 27323625.5 | 48111386.2 | 0.65629038 | 0.00034185 | 0.00243763 |
| sp\|Q15428\|S | 9273445.46 | 7352102.33 | -0.3714538 | 0.00034185 | 0.00243763 |
| sp\|O43353\|F | 206724.797 | 310231.339 | 0.63325916 | 0.00034185 | 0.00243763 |
| sp\|P63104\|1 | 1542095769 | 1885415428 | 0.27651039 | 0.00034185 | 0.00243763 |
| sp\|Q5VTU8\|A | 30096369.8 | 42884701.9 | 0.58178477 | 0.00034185 | 0.00243763 |
| sp\|O94973\|A | 44820259.6 | 32132795.8 | -0.4439515 | 0.0003814 | 0.00267987 |
| sp\|Q13526\|F | 32790627.2 | 41414224.6 | 0.32929011 | 0.0003814 | 0.00267987 |
| sp\|Q5T5P2\|S | 18431229.7 | 24687941.4 | 0.41427905 | 0.0003814 | 0.00267987 |
| sp\|Q9H0N5\| | 11741388 | 15829822.4 | 0.51451954 | 0.0003814 | 0.00267987 |
| sp\|P02549\|S | 106043421 | 282340325 | 0.88251142 | 0.0003814 | 0.00267987 |
| sp\|P08123\|C | 113926447 | 379150051 | 1.5706041 | 0.0003814 | 0.00267987 |
| sp\|P39023\|R | 241942891 | 100888421 | -1.2893915 | 0.0003814 | 0.00267987 |
| sp\|Q9UHX1\| | 41727999.1 | 30258790.4 | -0.4449837 | 0.0003814 | 0.00267987 |
| sp\|Q02978\|N | 15279226.4 | 10779716.3 | -0.4909104 | 0.0003814 | 0.00267987 |
| sp\|Q9UJY1\|H | 12933440.3 | 18218548.3 | 0.52402292 | 0.0003814 | 0.00267987 |
| sp\|Q9P0P0\|F | 6064818.23 | 10222435.8 | 0.64788522 | 0.00042497 | 0.00290929 |
| sp\|Q9Y3U8\|F | 15841369.8 | 6019704.4 | -1.3908333 | 0.00042497 | 0.00290929 |
| sp\|NQO2_HU | 155873238 | 279052536 | 0.59051524 | 0.00042497 | 0.00290929 |
| sp\|P63010\|A | 53815163.7 | 37555515 | -0.4904337 | 0.00042497 | 0.00290929 |
| sp\|Q12882\|D | 158688200 | 102401868 | -0.7240235 | 0.00042497 | 0.00290929 |
| sp\|P49327\|F | 2045336047 | 1278527446 | -1.2002272 | 0.00042497 | 0.00290929 |
| sp\|Q6ZVM7\|T | 13718129.3 | 18390074.3 | 0.4275043 | 0.00042497 | 0.00290929 |
| sp\|Q8TCT8\|S | 1297509.4 | 1737159.32 | 0.41098279 | 0.00042497 | 0.00290929 |
| sp\|Q15233\|N | 81424374 | 54585269.8 | -0.5457849 | 0.00042497 | 0.00290929 |
| sp\|O95757\|H | 51308286.1 | 35467496.1 | -0.6431415 | 0.00042497 | 0.00290929 |
| sp\|P46777\|R | 88519491.2 | 52442470.6 | -0.8393289 | 0.00042497 | 0.00290929 |
| sp\|Q9Y5J9\|TI | 2399623.12 | 3220019.72 | 0.53250905 | 0.00042497 | 0.00290929 |
| sp\|P48047\|A | 73294582 | 56291417.4 | -0.3722113 | 0.00042497 | 0.00290929 |
| sp\|Q4ZHG4\| | 1002618.28 | 4537847.58 | 1.27141296 | 0.00042497 | 0.00290929 |
| sp\|Q9UL42\|F | 33240346.4 | 18363484.5 | -1.1033521 | 0.00042497 | 0.00290929 |
| sp\|Q92734\|T | 38021327.1 | 47249311.4 | 0.29803546 | 0.00042497 | 0.00290929 |
| sp\|Q9NRG1\| | 1595209.26 | 3160600.4 | 0.89066527 | 0.00042497 | 0.00290929 |
| sp\|Q16795\|N | 9899198.71 | 6824019.51 | -0.5039869 | 0.00042497 | 0.00290929 |

| | | | | | |
|---|---|---|---|---|---|
| sp\|Q96DC8\|[illegible] | 149023422 | 101623176 | -0.6696065 | 0.0004729 | 0.00317407 |
| sp\|Q9NVD7\| | 6983073.52 | 9036058.88 | 0.34026163 | 0.0004729 | 0.00317407 |
| sp\|P42704\|L | 5463320.89 | 3938058.68 | -0.4942358 | 0.0004729 | 0.00317407 |
| sp\|Q9Y2Z0\|S | 19107583.4 | 25317972.2 | 0.40040111 | 0.0004729 | 0.00317407 |
| sp\|Q6P2Q9\|[illegible] | 56442288.5 | 41061170.4 | -0.4255092 | 0.0004729 | 0.00317407 |
| sp\|Q8N4T0\|[illegible] | 12340361.7 | 5710890.96 | -1.2855007 | 0.0004729 | 0.00317407 |
| sp\|Q9H4A4\|[illegible] | 844491206 | 571988814 | -0.6417771 | 0.0004729 | 0.00317407 |
| sp\|Q9Y275\|T | 15211144.6 | 20655571.2 | 0.42490157 | 0.0004729 | 0.00317407 |
| sp\|P36578\|R | 113111272 | 49884008.4 | -1.3049915 | 0.0004729 | 0.00317407 |
| sp\|P01034\|C | 53413795.9 | 74733634.8 | 0.44792773 | 0.0004729 | 0.00317407 |
| sp\|P62424\|R | 79440806.9 | 36359110 | -1.2435798 | 0.0004729 | 0.00317407 |
| sp\|P22392\|N | 1218295531 | 1551817970 | 0.32638994 | 0.0004729 | 0.00317407 |
| sp\|O75954\|T | 1064959.17 | 648374.667 | -0.6590551 | 0.0004729 | 0.00317407 |
| sp\|P05388\|R | 37041477.1 | 20233140.4 | -0.9268406 | 0.0004729 | 0.00317407 |
| sp\|Q63HR2\|[illegible] | 22891148.2 | 17481635.9 | -0.4074193 | 0.00052558 | 0.00342697 |
| sp\|O95299\|N | 25827238.8 | 19101532 | -0.4531887 | 0.00052558 | 0.00342697 |
| sp\|Q15717\|E | 120768916 | 89130406.5 | -0.4119331 | 0.00052558 | 0.00342697 |
| sp\|O95716\|F | 68036021.9 | 83709663.3 | 0.29885291 | 0.00052558 | 0.00342697 |
| sp\|P08670\|V | 2.6281E+10 | 3.4455E+10 | 0.35531053 | 0.00052558 | 0.00342697 |
| sp\|P00966\|A | 541317761 | 852993803 | 0.61446266 | 0.00052558 | 0.00342697 |
| sp\|P0CW20\| | 1941794.36 | 1421884.48 | -0.5815631 | 0.00052558 | 0.00342697 |
| sp\|Q9BSA9\|T | 267263.241 | 528998.54 | 1.00771201 | 0.00052558 | 0.00342697 |
| sp\|Q9BTV4\|T | 27201015.3 | 16142342.1 | -0.9341743 | 0.00052558 | 0.00342697 |
| sp\|P61313\|R | 57636750.4 | 30038000.9 | -1.0721451 | 0.00052558 | 0.00342697 |
| sp\|P55884\|E | 45411525 | 30369475.1 | -0.6008627 | 0.00052558 | 0.00342697 |
| sp\|P23141\|E | 3252055052 | 5498909409 | 0.77643612 | 0.00052558 | 0.00342697 |
| sp\|P06576\|A | 2260261502 | 3062709225 | 0.5003684 | 0.00052558 | 0.00342697 |
| sp\|O14737\|F | 72256463 | 92015328.9 | 0.38961438 | 0.00052558 | 0.00342697 |
| sp\|Q96C11\|F | 5054586.38 | 7218343.49 | 0.5073611 | 0.00052558 | 0.00342697 |
| sp\|Q9BXN1\|[illegible] | 221338136 | 123527498 | -1.1934049 | 0.00052558 | 0.00342697 |
| sp\|P16930\|F | 1437771857 | 2552851994 | 0.68409911 | 0.00052558 | 0.00342697 |
| sp\|Q53GQ0\| | 16746974.1 | 12136155 | -0.4842694 | 0.00052558 | 0.00342697 |
| sp\|Q92629\|S | 2172894.21 | 3486349.61 | 0.58939516 | 0.00052558 | 0.00342697 |
| sp\|P08047\|S | 76032.967 | 133457.627 | 0.89496685 | 0.00052558 | 0.00342697 |
| sp\|Q8WVE0\| | 916628.605 | 619747.088 | -0.4855191 | 0.00052558 | 0.00342697 |
| sp\|P49593\|P | 22218886.1 | 15111265.2 | -0.6273648 | 0.00058339 | 0.00367412 |
| sp\|P06703\|S | 515705869 | 701916035 | 0.41480276 | 0.00058339 | 0.00367412 |
| sp\|P07602\|S | 35885277.5 | 57605960.7 | 0.60830551 | 0.00058339 | 0.00367412 |
| sp\|P39060\|C | 69336682.8 | 91322367.5 | 0.41404489 | 0.00058339 | 0.00367412 |

| | | | | | |
|---|---|---|---|---|---|
| sp\|Q8IVF2\|A | 78729803.1 | 126973071 | 0.57986867 | 0.00058339 | 0.00367412 |
| sp\|Q9Y4P1\|A | 1653101.06 | 889015.625 | -0.9049432 | 0.00058339 | 0.00367412 |
| sp\|O95429\|B | 435975.741 | 1206812.69 | 1.1707911 | 0.00058339 | 0.00367412 |
| sp\|O95319\|C | 45265305 | 34441542.8 | -0.4045351 | 0.00058339 | 0.00367412 |
| sp\|Q99623\|P | 135683213 | 91900603.3 | -0.5493262 | 0.00058339 | 0.00367412 |
| sp\|Q9BQ67\|( | 182256.428 | 102435.285 | -2.0131573 | 0.00058339 | 0.00367412 |
| sp\|Q9Y2J2\|E | 127660635 | 103317075 | -0.3201209 | 0.00058339 | 0.00367412 |
| sp\|P23284\|P | 126154053 | 90476164.8 | -0.495939 | 0.00058339 | 0.00367412 |
| sp\|O75381\|F | 1008259.74 | 1937842.91 | 0.91938881 | 0.00058339 | 0.00367412 |
| sp\|P13674\|P | 24790611.6 | 19203825.5 | -0.4130494 | 0.00058339 | 0.00367412 |
| sp\|P06865\|H | 42681827.5 | 62831954.9 | 0.45401646 | 0.00058339 | 0.00367412 |
| sp\|Q92688\|A | 70636400.1 | 32132543.2 | -1.3077159 | 0.00058339 | 0.00367412 |
| sp\|P05556\|IT | 233328421 | 312921531 | 0.42471744 | 0.00058339 | 0.00367412 |
| sp\|P40855\|P | 11302702.2 | 15356097.2 | 0.46269195 | 0.00058339 | 0.00367412 |
| sp\|P24539\|A | 44911383.8 | 32508667.4 | -0.4917538 | 0.00058339 | 0.00367412 |
| sp\|P18621\|R | 48457108.2 | 20099306.7 | -1.3667222 | 0.00058339 | 0.00367412 |
| sp\|P18077\|R | 47084411.5 | 19489715.4 | -1.485151 | 0.00058339 | 0.00367412 |
| sp\|Q02241\|K | 4408924.48 | 8316291.2 | 0.60830342 | 0.00058339 | 0.00367412 |
| sp\|P27105\|S | 91688462.5 | 140451717 | 0.59902831 | 0.00058339 | 0.00367412 |
| sp\|Q13361\|M | 103792803 | 187662300 | 1.01902437 | 0.00058339 | 0.00367412 |
| sp\|P61254\|R | 57855337.5 | 24349076.8 | -1.3849926 | 0.00058339 | 0.00367412 |
| sp\|P55285\|C | 3251074.17 | 4663035.84 | 0.49046502 | 0.00058339 | 0.00367412 |
| sp\|Q6UXH9\| | 3405386.55 | 2207467.97 | -0.6155033 | 0.00064679 | 0.00396414 |
| sp\|P42574\|C | 15800783.6 | 24022453.5 | 0.64401723 | 0.00064679 | 0.00396414 |
| sp\|O60762\|D | 658115.569 | 305874.067 | -2.1363613 | 0.00064679 | 0.00396414 |
| sp\|P84095\|R | 52135740.6 | 71242578.8 | 0.41496252 | 0.00064679 | 0.00396414 |
| sp\|Q7Z5L9\|I2 | 7133913.91 | 5341279.76 | -0.4643386 | 0.00064679 | 0.00396414 |
| sp\|P51857\|A | 75355175.5 | 202370305 | 1.42043113 | 0.00064679 | 0.00396414 |
| sp\|Q99439\|C | 31984626.3 | 43838032 | 0.40051447 | 0.00064679 | 0.00396414 |
| sp\|P02746\|C | 21084347.8 | 12568971.9 | -0.7722159 | 0.00064679 | 0.00396414 |
| sp\|P0DJI8\|SA | 4353832.07 | 9243678.47 | 1.02307915 | 0.00064679 | 0.00396414 |
| sp\|Q14157\|U | 24056903.6 | 33365422.2 | 0.46580276 | 0.00064679 | 0.00396414 |
| sp\|P17844\|D | 169131381 | 99212705.1 | -0.8307415 | 0.00064679 | 0.00396414 |
| sp\|P49207\|R | 33063312 | 13782344.5 | -1.3211973 | 0.00064679 | 0.00396414 |
| sp\|Q15393\|S | 400525556 | 273943725 | -0.5728463 | 0.00064679 | 0.00396414 |
| sp\|Q9H7C9\|. | 59676455.8 | 114925893 | 0.74813921 | 0.00064679 | 0.00396414 |
| sp\|Q5ZPR3\|C | 2928457.7 | 4579520.5 | 0.58839834 | 0.00064679 | 0.00396414 |
| sp\|Q8N1G2\| | 369469.457 | 158436.084 | -1.2181778 | 0.00064679 | 0.00396414 |
| sp\|P68402\|P | 116490045 | 85613701.3 | -0.5082696 | 0.00064679 | 0.00396414 |

| | | | | | |
|---|---|---|---|---|---|
| sp\|P50135\|H | 196011705 | 126849542 | -0.6506639 | 0.00064679 | 0.00396414 |
| sp\|P50914\|R | 133931001 | 47495834 | -1.654658 | 0.00064679 | 0.00396414 |
| sp\|Q9Y624\|J | 46217739.6 | 65446756.4 | 0.45659118 | 0.00064679 | 0.00396414 |
| sp\|O00267\|S | 3766259.85 | 2642697.72 | -0.5352072 | 0.00064679 | 0.00396414 |
| sp\|Q9HCE1\| | 1312680.59 | 900358.296 | -0.8369638 | 0.00071621 | 0.00426968 |
| sp\|Q6EMK4\|' | 11237650.2 | 15246006.9 | 0.43539456 | 0.00071621 | 0.00426968 |
| sp\|P52895\|A | 1287831336 | 2843217739 | 1.09128801 | 0.00071621 | 0.00426968 |
| sp\|P02753\|R | 396377447 | 593092085 | 0.65113142 | 0.00071621 | 0.00426968 |
| sp\|Q07020\|F | 50012362.7 | 20922630.8 | -1.4133072 | 0.00071621 | 0.00426968 |
| sp\|Q92879\|C | 13453279.4 | 9446211.08 | -0.5280136 | 0.00071621 | 0.00426968 |
| sp\|Q92890\|U | 25216026.3 | 32363239.6 | 0.30807113 | 0.00071621 | 0.00426968 |
| sp\|P42765\|T | 818606276 | 1069069359 | 0.39003367 | 0.00071621 | 0.00426968 |
| sp\|Q14980\|N | 92267526.9 | 72213799.2 | -0.3306303 | 0.00071621 | 0.00426968 |
| sp\|Q9H4X1\|I | 1259056.15 | 1849362.26 | 0.59197777 | 0.00071621 | 0.00426968 |
| sp\|Q6UWE0\| | 1398853.58 | 2073357.2 | 0.45266074 | 0.00071621 | 0.00426968 |
| sp\|P34896\|G | 63694694 | 96990303.2 | 0.53492395 | 0.00071621 | 0.00426968 |
| sp\|Q15293\|F | 50048641.6 | 30150576.1 | -0.8026509 | 0.00071621 | 0.00426968 |
| sp\|Q8IXQ4\|G | 207823.471 | 108928.008 | -1.315017 | 0.00071621 | 0.00426968 |
| sp\|Q96I59\|S' | 4981497.25 | 2976177.32 | -0.8108927 | 0.00071621 | 0.00426968 |
| sp\|P47929\|L | 6025284.08 | 41599947.5 | 1.34479842 | 0.00071621 | 0.00426968 |
| sp\|P84085\|A | 46230032.1 | 57950389.5 | 0.31434345 | 0.00071621 | 0.00426968 |
| sp\|Q9UM22\| | 8447105.87 | 14395532.8 | 0.93997671 | 0.00071621 | 0.00426968 |
| sp\|P68036\|U | 19827487.1 | 28390206.9 | 0.51296643 | 0.00071621 | 0.00426968 |
| sp\|P18124\|R | 72740669.1 | 27765609.7 | -1.3381279 | 0.00071621 | 0.00426968 |
| sp\|Q9Y600\|C | 86272971.3 | 132809177 | 0.50866506 | 0.00071621 | 0.00426968 |
| sp\|P41227\|N | 1570698.46 | 947319.261 | -0.8351616 | 0.00071621 | 0.00426968 |
| sp\|Q9Y5P6\|C | 45742014.2 | 36084204.7 | -0.337851 | 0.00079215 | 0.00463034 |
| sp\|Q99470\|S | 4167113.19 | 6528974.75 | 0.50272336 | 0.00079215 | 0.00463034 |
| sp\|P10619\|P | 20119194.7 | 27301998.1 | 0.38041936 | 0.00079215 | 0.00463034 |
| sp\|P22105\|T | 232912160 | 171605185 | -0.460982 | 0.00079215 | 0.00463034 |
| sp\|P28288\|A | 434123.459 | 688562.812 | 0.60492453 | 0.00079215 | 0.00463034 |
| sp\|Q13247\|S | 36025747.8 | 22568581.8 | -0.6011061 | 0.00079215 | 0.00463034 |
| sp\|Q7Z6M1\|I | 15453904.1 | 23138792 | 0.48211968 | 0.00079215 | 0.00463034 |
| sp\|P09001\|R | 5989827.12 | 9153809.82 | 0.47732131 | 0.00079215 | 0.00463034 |
| sp\|Q9NP77\|S | 2596370.24 | 1678382.59 | -0.7777669 | 0.00079215 | 0.00463034 |
| sp\|Q13405\|F | 3830833.43 | 2864300.61 | -0.682053 | 0.00079215 | 0.00463034 |
| sp\|P28066\|P | 182837414 | 259684422 | 0.38626509 | 0.00079215 | 0.00463034 |
| sp\|A5YM72\|C | 1262266.23 | 783667.997 | -0.977379 | 0.00079215 | 0.00463034 |
| sp\|P02745\|C | 14213785 | 8918487.64 | -0.5920488 | 0.00079215 | 0.00463034 |

| | | | | | |
|---|---|---|---|---|---|
| sp\|O75131\|( | 37386721 | 27405212.2 | -0.4689137 | 0.00079215 | 0.00463034 |
| sp\|P05181\|C | 563952.817 | 909190.051 | 0.58593252 | 0.00079215 | 0.00463034 |
| sp\|Q7L3T8\|S | 624086.326 | 472596.51 | -0.4560761 | 0.00079215 | 0.00463034 |
| sp\|Q8IXH6\|T | 493809.002 | 201769.727 | -1.9290493 | 0.00084923 | 0.00495797 |
| sp\|Q9UKU7\|/ | 35790636.8 | 50573389.8 | 0.50260776 | 0.00087515 | 0.004982 |
| sp\|P42766\|R | 9228642.77 | 4951961.2 | -0.8524301 | 0.00087515 | 0.004982 |
| sp\|P26006\|I | 6913881.37 | 11051898.7 | 0.46122093 | 0.00087515 | 0.004982 |
| sp\|P78356\|P | 22526517.4 | 13773898.2 | -0.8407232 | 0.00087515 | 0.004982 |
| sp\|P26373\|R | 78916741.6 | 36901512.3 | -1.0805862 | 0.00087515 | 0.004982 |
| sp\|P63167\|D | 31700693.3 | 26488274.2 | -0.2288025 | 0.00087515 | 0.004982 |
| sp\|P50148\|G | 10241003.8 | 8325530.58 | -0.3301957 | 0.00087515 | 0.004982 |
| sp\|P49721\|P | 193031148 | 266420789 | 0.38210649 | 0.00087515 | 0.004982 |
| sp\|P08631\|H | 66753346 | 53235684.4 | -0.3553317 | 0.00087515 | 0.004982 |
| sp\|O75970\|N | 4231203.99 | 3000372.7 | -0.5214254 | 0.00087515 | 0.004982 |
| sp\|P35443\|T | 6597318.45 | 9718014.3 | 0.50758092 | 0.00087515 | 0.004982 |
| sp\|P48304\|R | 154178.472 | 694085.446 | 1.01939008 | 0.00087515 | 0.004982 |
| sp\|Q8WTS6\|S | 20128792.5 | 12491147.1 | -0.8889452 | 0.00087515 | 0.004982 |
| sp\|Q15029\|U | 110800105 | 85690496.7 | -0.3680534 | 0.00087515 | 0.004982 |
| sp\|Q8WTS1\|/ | 12876775 | 20718537.5 | 0.4884993 | 0.00087515 | 0.004982 |
| sp\|P20618\|P | 122515965 | 177726964 | 0.43916005 | 0.00087515 | 0.004982 |
| sp\|P54920\|S | 40004877.6 | 48628084.5 | 0.30054939 | 0.00087515 | 0.004982 |
| sp\|O14964\|H | 16319870.3 | 19555304.7 | 0.26956003 | 0.00087515 | 0.004982 |
| sp\|Q6IAA8\|L | 12777823.5 | 18248815.6 | 0.48819987 | 0.00087515 | 0.004982 |
| sp\|Q15056\|I | 49060643.1 | 72739155.4 | 0.48358924 | 0.00087515 | 0.004982 |
| sp\|P62888\|R | 89534780.2 | 43262557.1 | -1.1665727 | 0.00087515 | 0.004982 |
| sp\|P61204\|A | 104131661 | 128650872 | 0.27791183 | 0.00096574 | 0.00538904 |
| sp\|P47224\|N | 9053558.68 | 13462898.1 | 0.50160193 | 0.00096574 | 0.00538904 |
| sp\|Q9H0E2\|T | 27646362.8 | 36937243.7 | 0.34331781 | 0.00096574 | 0.00538904 |
| sp\|P53007\|T | 55867004.2 | 44551827.5 | -0.3383314 | 0.00096574 | 0.00538904 |
| sp\|P54136\|S | 54391927.2 | 37268921.3 | -0.5737218 | 0.00096574 | 0.00538904 |
| sp\|P29992\|G | 12991604.2 | 10456432.7 | -0.3296317 | 0.00096574 | 0.00538904 |
| sp\|Q4U2R6\|I | 877358.676 | 599570.843 | -0.9648611 | 0.00096574 | 0.00538904 |
| sp\|P07333\|C | 5279246.9 | 8869645.37 | 0.52256968 | 0.00096574 | 0.00538904 |
| sp\|P62487\|R | 8210943.02 | 6576001.5 | -0.3439411 | 0.00096574 | 0.00538904 |
| sp\|Q9NVG8\| | 7042760.76 | 5035173.5 | -0.5525091 | 0.00096574 | 0.00538904 |
| sp\|P60484\|P | 2932154.44 | 1799191.64 | -0.7692847 | 0.00096574 | 0.00538904 |
| sp\|Q9UNK0\|S | 30763731 | 43433861.9 | 0.47609074 | 0.00096574 | 0.00538904 |
| sp\|Q9UDW1\| | 1476943.19 | 850997.893 | -0.7900049 | 0.00096574 | 0.00538904 |
| sp\|P10586\|P | 52114541.8 | 41897687.2 | -0.3427823 | 0.00096574 | 0.00538904 |

| | | | | | |
|---|---|---|---|---|---|
| sp\|Q9BX97\|F | 1254264.75 | 1768529.65 | 0.54381117 | 0.00096574 | 0.00538904 |
| sp\|O00170\|A | 10109484.2 | 6467572.46 | -0.6358711 | 0.00096574 | 0.00538904 |
| sp\|Q9UJY5\|G | 239783.272 | 440491.814 | 0.86891832 | 0.00096574 | 0.00538904 |
| sp\|Q02878\|F | 119145406 | 53569672.2 | -1.192879 | 0.00106452 | 0.00577898 |
| sp\|Q8N8S7\|E | 31591202.8 | 40723283.4 | 0.37055499 | 0.00106452 | 0.00577898 |
| sp\|Q13148\|T | 17473158.6 | 10358093 | -0.8823253 | 0.00106452 | 0.00577898 |
| sp\|Q9BR76\|C | 352453796 | 261525556 | -0.549033 | 0.00106452 | 0.00577898 |
| sp\|Q00325\|N | 309053970 | 223027641 | -0.483414 | 0.00106452 | 0.00577898 |
| sp\|Q01433\|A | 190672028 | 152036519 | -0.3381499 | 0.00106452 | 0.00577898 |
| sp\|P06702\|S | 56703455.8 | 249000800 | 1.21276169 | 0.00106452 | 0.00577898 |
| sp\|Q13555\|K | 36152662.6 | 25146940.7 | -0.5554376 | 0.00106452 | 0.00577898 |
| sp\|Q6IB77\|G | 22922787.5 | 34142049.2 | 0.61148001 | 0.00106452 | 0.00577898 |
| sp\|Q15154\|F | 1785131.05 | 2703275.76 | 0.62606717 | 0.00106452 | 0.00577898 |
| sp\|P62910\|R | 10336972.5 | 4125258.49 | -1.2162375 | 0.00106452 | 0.00577898 |
| sp\|Q8TBE0\|E | 448850.34 | 792592.086 | 0.67196563 | 0.00106452 | 0.00577898 |
| sp\|P61353\|R | 80180330.3 | 36302359.9 | -1.4365453 | 0.00106452 | 0.00577898 |
| sp\|O60506\|H | 242290735 | 169859614 | -0.5478799 | 0.00106452 | 0.00577898 |
| sp\|Q16798\|N | 58231019.7 | 81101565.5 | 0.38772297 | 0.00106452 | 0.00577898 |
| sp\|Q9NSE4\|S | 174251043 | 130673574 | -0.4859643 | 0.00106452 | 0.00577898 |
| sp\|P38919\|IF | 39816433.7 | 24623618 | -0.7490064 | 0.00106452 | 0.00577898 |
| sp\|Q04206\|T | 25828550.5 | 20752544.5 | -0.3221683 | 0.00106452 | 0.00577898 |
| sp\|O75964\|A | 14877504.7 | 9596994.14 | -0.6245798 | 0.00106452 | 0.00577898 |
| sp\|P60468\|S | 863888.439 | 423656.786 | -0.962204 | 0.00106452 | 0.00577898 |
| sp\|P18206\|V | 1038932099 | 1513234558 | 0.52531771 | 0.00106452 | 0.00577898 |
| sp\|Q9NPG3\| | 1390742.04 | 3260569.76 | 0.79984848 | 0.00106452 | 0.00577898 |
| sp\|Q9H8H3\| | 39609222.2 | 26940534.7 | -0.7101588 | 0.00106452 | 0.00577898 |
| sp\|Q13444\|A | 1276919.98 | 1999005.39 | 0.63905165 | 0.00106452 | 0.00577898 |
| sp\|Q6YP21\|K | 96379143.8 | 121662006 | 0.38985332 | 0.00117213 | 0.00622242 |
| sp\|Q7L1Q6\|E | 4012373.44 | 8098518.95 | 1.42222048 | 0.00117213 | 0.00622242 |
| sp\|Q04727\|T | 1895688.06 | 1431463.49 | -0.462668 | 0.00117213 | 0.00622242 |
| sp\|P84098\|R | 19124692.3 | 11509905.5 | -0.7191526 | 0.00117213 | 0.00622242 |
| sp\|Q13263\|T | 33809634.5 | 25560462.4 | -0.4142546 | 0.00117213 | 0.00622242 |
| sp\|Q01813\|F | 92909749.8 | 66547017.8 | -0.5333575 | 0.00117213 | 0.00622242 |
| sp\|P61006\|R | 115493814 | 143689442 | 0.2896596 | 0.00117213 | 0.00622242 |
| sp\|Q9H7E2\|T | 34843082.7 | 62756583.1 | 0.65584618 | 0.00117213 | 0.00622242 |
| sp\|P51808\|D | 15589247.1 | 21408978.7 | 0.4547986 | 0.00117213 | 0.00622242 |
| sp\|P49760\|C | 1910438.86 | 976085.571 | -1.1554246 | 0.00117213 | 0.00622242 |
| sp\|P11182\|C | 24864856.7 | 20296235.6 | -0.3212351 | 0.00117213 | 0.00622242 |
| sp\|O15389\|S | 2651835.95 | 3505212.68 | 0.4303866 | 0.00117213 | 0.00622242 |

| | | | | | |
|---|---|---|---|---|---|
| sp\|P49821\|N | 140575128 | 185478603 | 0.30663701 | 0.00117213 | 0.00622242 |
| sp\|Q9UPU7\|T | 7593819.69 | 10873015.2 | 0.44118166 | 0.00117213 | 0.00622242 |
| sp\|Q96C86\|I | 3667866.34 | 2151497.75 | -0.8156854 | 0.00117213 | 0.00622242 |
| sp\|Q02543\|F | 51757451.1 | 21190492.9 | -1.395107 | 0.00117213 | 0.00622242 |
| sp\|O60925\|F | 42053945.2 | 54780234.8 | 0.39717486 | 0.00117213 | 0.00622242 |
| sp\|Q8TD30\|A | 58356017.9 | 46107861.3 | -0.3605201 | 0.00117213 | 0.00622242 |
| sp\|O00203\|A | 3879057.05 | 2876167.67 | -0.4993256 | 0.00117213 | 0.00622242 |
| sp\|Q9NVJ2\|A | 9863721 | 13241388.2 | 0.38161457 | 0.00117213 | 0.00622242 |
| sp\|Q9H0U4\| | 170427583 | 217409962 | 0.34035893 | 0.00128923 | 0.00669592 |
| sp\|P22897\|M | 110471867 | 79501099.2 | -0.4886874 | 0.00128923 | 0.00669592 |
| sp\|Q9UKY3\|C | 480619514 | 809592133 | 0.7537289 | 0.00128923 | 0.00669592 |
| sp\|P45984\|M | 10417489.2 | 8099098.2 | -0.3868359 | 0.00128923 | 0.00669592 |
| sp\|P07686\|H | 102937227 | 160736742 | 0.50899237 | 0.00128923 | 0.00669592 |
| sp\|P04275\|V | 481182563 | 599446806 | 0.33177181 | 0.00128923 | 0.00669592 |
| sp\|P62913\|R | 49733996.6 | 24884022.3 | -0.9983138 | 0.00128923 | 0.00669592 |
| sp\|P61225\|R | 3286453.14 | 4392423.16 | 0.36167455 | 0.00128923 | 0.00669592 |
| sp\|P18085\|A | 50976482.4 | 63015418.5 | 0.29590121 | 0.00128923 | 0.00669592 |
| sp\|P26368\|U | 23431706 | 17036518.6 | -0.4925973 | 0.00128923 | 0.00669592 |
| sp\|Q9HBL0\|T | 345886976 | 439759965 | 0.33172474 | 0.00128923 | 0.00669592 |
| sp\|O95782\|A | 47860403.8 | 37513754.6 | -0.3499505 | 0.00128923 | 0.00669592 |
| sp\|Q8NBL1\|F | 993149.542 | 674038.252 | -0.6428075 | 0.00128923 | 0.00669592 |
| sp\|Q8NFP9\|N | 6371300.37 | 11901864.5 | 0.8976435 | 0.00128923 | 0.00669592 |
| sp\|P47756\|C | 237275392 | 188268111 | -0.4341326 | 0.00128923 | 0.00669592 |
| sp\|Q16595\|F | 53061533.1 | 68863626.3 | 0.30759487 | 0.00128923 | 0.00669592 |
| sp\|P08238\|H | 263582804 | 167614026 | -0.7547576 | 0.00128923 | 0.00669592 |
| sp\|P56134\|A | 29493519.3 | 23171128.8 | -0.376161 | 0.00128923 | 0.00669592 |
| sp\|P17568\|N | 1436964.13 | 1017428.03 | -0.4733721 | 0.00128923 | 0.00669592 |
| sp\|P34949\|M | 65479644.5 | 50477964.1 | -0.4352482 | 0.00128923 | 0.00669592 |
| sp\|P19388\|R | 1801947.66 | 1146940.7 | -0.7705523 | 0.00141653 | 0.0071557 |
| sp\|Q15847\|A | 89639622 | 117144265 | 0.49954946 | 0.00141653 | 0.0071557 |
| sp\|Q9P0J1\|P | 14266339 | 11058441.9 | -0.4068868 | 0.00141653 | 0.0071557 |
| sp\|P63244\|R | 306343573 | 220689061 | -0.575454 | 0.00141653 | 0.0071557 |
| sp\|P55084\|E | 266831505 | 185919203 | -0.6219469 | 0.00141653 | 0.0071557 |
| sp\|P23946\|C | 89202138.1 | 49287714.1 | -0.8943681 | 0.00141653 | 0.0071557 |
| sp\|Q7Z434\|N | 1466607.06 | 2220943.42 | 0.62268949 | 0.00141653 | 0.0071557 |
| sp\|Q9UBG0\| | 42195517.3 | 32984713.7 | -0.4285301 | 0.00141653 | 0.0071557 |
| sp\|Q9UNZ2\|I | 135905114 | 171211681 | 0.31419407 | 0.00141653 | 0.0071557 |
| sp\|Q9Y4H2\|I | 1674079.42 | 2451281.78 | 0.62754352 | 0.00141653 | 0.0071557 |
| sp\|Q8NEV1\|C | 14705831.9 | 11103720.1 | -0.4401456 | 0.00141653 | 0.0071557 |

| | | | | | |
|---|---|---|---|---|---|
| sp\|P52888\|T | 34817085.3 | 43397506.3 | 0.28917123 | 0.00141653 | 0.0071557 |
| sp\|Q96CW1\| | 29876099 | 21046254.2 | -0.5289114 | 0.00141653 | 0.0071557 |
| sp\|Q53QV2\|I | 1888728.06 | 945358.243 | -1.206064 | 0.00141653 | 0.0071557 |
| sp\|O14561\|A | 212644.249 | 384067.879 | 1.59457023 | 0.00141653 | 0.0071557 |
| sp\|Q13427\|F | 2443810.69 | 1567689.64 | -0.7107056 | 0.00141653 | 0.0071557 |
| sp\|Q8TDQ0\|I | 1468328.51 | 2365626.46 | 0.57112318 | 0.00141653 | 0.0071557 |
| sp\|P40939\|E | 316487325 | 217902359 | -0.6265647 | 0.00141653 | 0.0071557 |
| sp\|P06727\|A | 790750058 | 1362054020 | 0.85964079 | 0.00141653 | 0.0071557 |
| sp\|Q9UMS4\| | 12924048.8 | 9746057.74 | -0.4210262 | 0.00141653 | 0.0071557 |
| sp\|P63151\|2 | 21903009.5 | 17084640.2 | -0.3528436 | 0.00141653 | 0.0071557 |
| sp\|P19367\|H | 173642118 | 210439708 | 0.31220043 | 0.00141653 | 0.0071557 |
| sp\|P50402\|E | 7206779.06 | 8964282.51 | 0.32876129 | 0.00141653 | 0.0071557 |
| sp\|P49720\|P | 185300863 | 259439344 | 0.39764402 | 0.00141653 | 0.0071557 |
| sp\|Q7Z5L7\|P | 8160318.02 | 4780285.09 | -0.8513282 | 0.00141653 | 0.0071557 |
| sp\|O75844\|F | 107492.001 | 173785.847 | 0.66532183 | 0.00141653 | 0.0071557 |
| sp\|Q15691\|M | 12845661.4 | 17223959 | 0.37192571 | 0.00155479 | 0.00770013 |
| sp\|Q6P587\|F | 71104662.3 | 103096901 | 0.45795091 | 0.00155479 | 0.00770013 |
| sp\|Q13185\|C | 25330906.7 | 19623110.2 | -0.3866021 | 0.00155479 | 0.00770013 |
| sp\|P04075\|A | 1639716690 | 2261757911 | 0.43312509 | 0.00155479 | 0.00770013 |
| sp\|Q9UHD8\| | 149373667 | 107052765 | -0.4161155 | 0.00155479 | 0.00770013 |
| sp\|Q9H3G5\| | 24779432.3 | 18260549.6 | -0.4399546 | 0.00155479 | 0.00770013 |
| sp\|Q8WUM4 | 10099596.6 | 13129063.6 | 0.34801688 | 0.00155479 | 0.00770013 |
| sp\|Q9BQI0\|A | 6164183.46 | 8151788.89 | 0.44086946 | 0.00155479 | 0.00770013 |
| sp\|P28330\|A | 84693222.2 | 97915030.7 | 0.47359993 | 0.00155479 | 0.00770013 |
| sp\|Q92835\|S | 1281801.98 | 980307.734 | -0.5496983 | 0.00155479 | 0.00770013 |
| sp\|P12110\|C | 55237596.3 | 99044988 | 0.72138976 | 0.00155479 | 0.00770013 |
| sp\|P17900\|S | 29986590.5 | 44580197.4 | 0.57795138 | 0.00155479 | 0.00770013 |
| sp\|Q14151\|S | 4724298.55 | 3570188.82 | -0.3832734 | 0.00155479 | 0.00770013 |
| sp\|Q9H477\|I | 15143112.4 | 22232517.4 | 0.49601572 | 0.00155479 | 0.00770013 |
| sp\|Q9UQ03\| | 3449611.11 | 5013906.57 | 0.41866379 | 0.00155479 | 0.00770013 |
| sp\|P31943\|H | 208610112 | 145445779 | -0.6096053 | 0.00155479 | 0.00770013 |
| sp\|P09619\|P | 20671831.1 | 15691636 | -0.417621 | 0.00155479 | 0.00770013 |
| sp\|Q99614\|T | 9948550.85 | 12178167.8 | 0.28760428 | 0.00155479 | 0.00770013 |
| sp\|P16278\|B | 62057841.9 | 103815830 | 0.53311431 | 0.00155479 | 0.00770013 |
| sp\|Q96E17\|F | 66483669.9 | 78700823 | 0.2409299 | 0.00170479 | 0.00825559 |
| sp\|P63096\|G | 113872126 | 89886540.7 | -0.3734855 | 0.00170479 | 0.00825559 |
| sp\|P33908\|M | 4615127.94 | 5852930.84 | 0.30871353 | 0.00170479 | 0.00825559 |
| sp\|P10155\|R | 145282352 | 113089536 | -0.4373482 | 0.00170479 | 0.00825559 |
| sp\|P81605\|D | 978797.927 | 2649432.1 | 0.9683162 | 0.00170479 | 0.00825559 |

| | | | | | |
|---|---|---|---|---|---|
| sp\|P62318\|S | 73897993.1 | 52265318.7 | -0.5109893 | 0.00170479 | 0.00825559 |
| sp\|P48637\|G | 296494375 | 386592471 | 0.33983369 | 0.00170479 | 0.00825559 |
| sp\|Q9NUM4\| | 3651784.51 | 4622598 | 0.33387368 | 0.00170479 | 0.00825559 |
| sp\|Q6L8Q7\|F | 18144102.6 | 11688039.1 | -0.7184206 | 0.00170479 | 0.00825559 |
| sp\|Q7Z4G4\|T | 698615.979 | 1486451.6 | 0.92876936 | 0.00170479 | 0.00825559 |
| sp\|Q9Y2G1\|I | 4908721.73 | 6445423.66 | 0.39019 | 0.00170479 | 0.00825559 |
| sp\|P07900\|H | 323023975 | 199488719 | -0.808877 | 0.00170479 | 0.00825559 |
| sp\|Q9HC38\| | 256191608 | 191787297 | -0.6290573 | 0.00170479 | 0.00825559 |
| sp\|Q99424\|A | 23765388.6 | 31782514.2 | 0.37520158 | 0.00170479 | 0.00825559 |
| sp\|Q49A26\|C | 2303984.15 | 1611234.75 | -0.5343582 | 0.00170479 | 0.00825559 |
| sp\|Q13485\|S | 5962135 | 4273184.69 | -1.1728347 | 0.00170479 | 0.00825559 |
| sp\|P24592\|IE | 7602305.68 | 12175904.9 | 0.56380268 | 0.00170479 | 0.00825559 |
| sp\|Q02252\|M | 520685731 | 332554029 | -0.9330307 | 0.00170479 | 0.00825559 |
| sp\|Q9BT73\|F | 15702854.6 | 19598111.9 | 0.3172471 | 0.00170479 | 0.00825559 |
| sp\|Q9BYT8\|N | 15766179.6 | 26943411.2 | 0.61586947 | 0.00170479 | 0.00825559 |
| sp\|Q13326\|S | 17979901 | 28639604.1 | 0.69402174 | 0.00170479 | 0.00825559 |
| sp\|O60610\|D | 1183529.6 | 2948121.07 | 0.5326975 | 0.00170479 | 0.00825559 |
| sp\|Q8ND24\| | 2128508.45 | 4256629.02 | 0.8573408 | 0.00186737 | 0.0088117 |
| sp\|Q15493\|F | 36496071.8 | 60441265.6 | 0.71148643 | 0.00186737 | 0.0088117 |
| sp\|Q6DKI1\|R | 6425449.22 | 2411884 | -1.6153784 | 0.00186737 | 0.0088117 |
| sp\|Q8NDI1\|E | 2257466.74 | 3209789.18 | 0.45439535 | 0.00186737 | 0.0088117 |
| sp\|Q9BQ69\|I | 7886392.27 | 4504760.2 | -1.0428562 | 0.00186737 | 0.0088117 |
| sp\|Q9P2Y5\|U | 2087930.36 | 2859230.42 | 0.46284962 | 0.00186737 | 0.0088117 |
| sp\|Q00796\|D | 73366238.1 | 62588778.5 | -0.2529021 | 0.00186737 | 0.0088117 |
| sp\|P46939\|U | 160709845 | 186397674 | 0.22047572 | 0.00186737 | 0.0088117 |
| sp\|P49591\|S | 44067323.5 | 34324577.4 | -0.401945 | 0.00186737 | 0.0088117 |
| sp\|Q9BWH2\| | 1409806.94 | 2063492.96 | 0.53836636 | 0.00186737 | 0.0088117 |
| sp\|Q9NRX5\|S | 668555.537 | 1053136.35 | 0.51847549 | 0.00186737 | 0.0088117 |
| sp\|P41271\|N | 1088203.79 | 8846395.99 | 1.7610478 | 0.00186737 | 0.0088117 |
| sp\|Q96CD2\| | 4858027.54 | 6098139.61 | 0.32254904 | 0.00186737 | 0.0088117 |
| sp\|P02652\|A | 167115561 | 270094782 | 0.5146351 | 0.00186737 | 0.0088117 |
| sp\|Q92945\|F | 134773533 | 103977804 | -0.4074081 | 0.00186737 | 0.0088117 |
| sp\|Q7L266\|A | 6357346.95 | 10222165.4 | 0.56199919 | 0.00186737 | 0.0088117 |
| sp\|O14949\|C | 3265366.36 | 2350567.98 | -0.4934393 | 0.00186737 | 0.0088117 |
| sp\|P13693\|T | 45114439 | 54758302.3 | 0.27937452 | 0.00186737 | 0.0088117 |
| sp\|P51571\|S | 3200766.91 | 2001895.5 | -0.6111521 | 0.00186737 | 0.0088117 |
| sp\|P11215\|IT | 14154557.2 | 22249170.1 | 0.5812874 | 0.00186737 | 0.0088117 |
| sp\|O95169\|N | 1771539.68 | 1248756.79 | -0.8100354 | 0.00186737 | 0.0088117 |
| sp\|O15371\|E | 71186886 | 49692950.3 | -0.6128828 | 0.00186737 | 0.0088117 |

| | | | | | |
|---|---|---|---|---|---|
| sp\|Q9HBW9\| | 2505982.04 | 3646500.45 | 0.6033251 | 0.00186737 | 0.0088117 |
| sp\|P62072\|T | 4723821.52 | 6042973.43 | 0.44478965 | 0.00186737 | 0.0088117 |
| sp\|P51991\|R | 101212033 | 71090081.9 | -0.5235202 | 0.00186737 | 0.0088117 |
| sp\|P51888\|P | 1299926561 | 991387894 | -0.5047722 | 0.00186737 | 0.0088117 |
| sp\|O43399\|T | 86846770.3 | 116581545 | 0.4352406 | 0.00204344 | 0.0094932 |
| sp\|Q12765\|S | 52988837.3 | 39914990.5 | -0.4932237 | 0.00204344 | 0.0094932 |
| sp\|Q99848\|E | 356542.059 | 715572.36 | -4.798917 | 0.00204344 | 0.0094932 |
| sp\|P62807\|H | 436368837 | 251664805 | -1.0154789 | 0.00204344 | 0.0094932 |
| sp\|Q9P0J7\|K | 1115836.97 | 1645003 | 0.51827752 | 0.00204344 | 0.0094932 |
| sp\|P10620\|M | 58116353.6 | 34911313 | -1.0645323 | 0.00204344 | 0.0094932 |
| sp\|P33778\|H | 417529561 | 235923084 | -0.9882043 | 0.00204344 | 0.0094932 |
| sp\|A0FGR8\|E | 31792529.8 | 23634312.4 | -0.4344632 | 0.00204344 | 0.0094932 |
| sp\|Q9P270\|S | 7925814.38 | 10651480.6 | 0.38458439 | 0.00204344 | 0.0094932 |
| sp\|P01040\|C | 6545415.82 | 11914216.1 | 0.71888957 | 0.00204344 | 0.0094932 |
| sp\|Q92558\|V | 88995139.4 | 176957330 | 0.75083978 | 0.00204344 | 0.0094932 |
| sp\|Q9H7C4\|: | 2072314.55 | 3194534.72 | 0.7340124 | 0.00204344 | 0.0094932 |
| sp\|P02647\|A | 6267958859 | 1.1202E+10 | 0.81217848 | 0.00204344 | 0.0094932 |
| sp\|O60547\|( | 46079164.7 | 61017359.4 | 0.34368871 | 0.00204344 | 0.0094932 |
| sp\|P02462\|C | 29588687.1 | 45764838.2 | 0.64800059 | 0.00204344 | 0.0094932 |
| sp\|A5D8V7\|( | 579408.45 | 1442447.13 | 0.41908765 | 0.00204344 | 0.0094932 |
| sp\|P23229\|IT | 10982133.6 | 15240475 | 0.43646022 | 0.00223392 | 0.01010424 |
| sp\|Q9UMS0\| | 10507012.6 | 14379324.4 | 0.488201 | 0.00223392 | 0.01010424 |
| sp\|Q13277\|S | 7871180.21 | 11387174.5 | 0.57910411 | 0.00223392 | 0.01010424 |
| sp\|Q12959\|D | 21082466.5 | 17200478.5 | -0.3101108 | 0.00223392 | 0.01010424 |
| sp\|Q8TCS8\|F | 11582355.3 | 8720561.11 | -0.497925 | 0.00223392 | 0.01010424 |
| sp\|Q8TD19\|N | 28167362.9 | 21473530.2 | -0.4436778 | 0.00223392 | 0.01010424 |
| sp\|Q9NX55\|I | 6261696.43 | 7932464.34 | 0.35799029 | 0.00223392 | 0.01010424 |
| sp\|Q96QU6\| | 1397206.14 | 1219268.76 | 0.69757983 | 0.00223392 | 0.01010424 |
| sp\|O60568\|F | 19373291 | 13770993.2 | -0.5021519 | 0.00223392 | 0.01010424 |
| sp\|P22314\|U | 635430086 | 423872196 | -0.7978848 | 0.00223392 | 0.01010424 |
| sp\|P43686\|P | 44238160.5 | 66407004.4 | 0.49949986 | 0.00223392 | 0.01010424 |
| sp\|Q32P44\|E | 30240904.6 | 24038352.1 | -0.3806879 | 0.00223392 | 0.01010424 |
| sp\|Q92576\|F | 1813770.76 | 5455225.83 | 1.58857053 | 0.00223392 | 0.01010424 |
| sp\|Q96PK6\|F | 7055126.47 | 4343994.34 | -0.7019777 | 0.00223392 | 0.01010424 |
| sp\|Q9UBI6\|G | 19923013.7 | 14700049 | -0.5435067 | 0.00223392 | 0.01010424 |
| sp\|Q86V88\|N | 3350153.19 | 2461539.92 | -0.5860142 | 0.00223392 | 0.01010424 |
| sp\|O00193\|S | 3033067.9 | 1830644.84 | -0.643801 | 0.00223392 | 0.01010424 |
| sp\|Q99766\|A | 1069743.9 | 1748474.42 | 0.66119312 | 0.00223392 | 0.01010424 |
| sp\|Q96HN2\|: | 67361292 | 43472610.4 | -0.5924201 | 0.00223392 | 0.01010424 |

| | | | | | |
|---|---|---|---|---|---|
| sp\|P50281\|M | 2306433.59 | 3564936.02 | 0.5305516 | 0.00223392 | 0.01010424 |
| sp\|Q8WWB7 | 2950287.79 | 3904936.71 | 0.34058378 | 0.00223392 | 0.01010424 |
| sp\|A0A087W | 3487377.06 | 6077345.25 | 0.57892406 | 0.00223392 | 0.01010424 |
| sp\|Q9Y6K9\|N | 18647260.2 | 22530364.2 | 0.28259348 | 0.00223392 | 0.01010424 |
| sp\|P21333\|F | 1797152520 | 1119678454 | -0.6657132 | 0.00223392 | 0.01010424 |
| sp\|P49773\|H | 122802169 | 149321891 | 0.25063282 | 0.00223392 | 0.01010424 |
| sp\|Q9BWT6\| | 746298.864 | 362931.653 | -1.831754 | 0.00223392 | 0.01010424 |
| sp\|Q9GZP4\|I | 62352036 | 82434541.7 | 0.39657618 | 0.00223392 | 0.01010424 |
| sp\|P55060\|X | 165354654 | 90091649.3 | -1.0537235 | 0.00223392 | 0.01010424 |
| sp\|Q14728\|N | 796282.966 | 412537.156 | -2.1845943 | 0.00243979 | 0.01077144 |
| sp\|P31040\|S | 218858205 | 293344617 | 0.37473957 | 0.00243979 | 0.01077144 |
| sp\|Q9BUT1\|E | 126697030 | 85793629.9 | -0.6607946 | 0.00243979 | 0.01077144 |
| sp\|Q6GMV3\| | 206856463 | 280093747 | 0.5053979 | 0.00243979 | 0.01077144 |
| sp\|Q8TCC3\|I | 4451646.13 | 8295124.24 | 1.60449765 | 0.00243979 | 0.01077144 |
| sp\|O60551\|N | 1281467963 | 1632968261 | 0.31505688 | 0.00243979 | 0.01077144 |
| sp\|Q06418\|T | 1678994.57 | 2484298.75 | 0.60037976 | 0.00243979 | 0.01077144 |
| sp\|Q05707\|C | 187850568 | 117744712 | -0.7576266 | 0.00243979 | 0.01077144 |
| sp\|Q8WUW1 | 905624.125 | 1335709.63 | 0.46906597 | 0.00243979 | 0.01077144 |
| sp\|P35237\|S | 466415208 | 366252569 | -0.4411916 | 0.00243979 | 0.01077144 |
| sp\|P49588\|S | 125941565 | 91546225.9 | -0.574608 | 0.00243979 | 0.01077144 |
| sp\|Q8N2H3\| | 23158161.1 | 34523782.6 | 0.51385054 | 0.00243979 | 0.01077144 |
| sp\|P00403\|C | 25609122.6 | 17184777.8 | -0.5989939 | 0.00243979 | 0.01077144 |
| sp\|O95149\|S | 647764.564 | 410386.458 | -1.2080933 | 0.00243979 | 0.01077144 |
| sp\|Q9NQP4\| | 12222861.2 | 16210378.1 | 0.38907381 | 0.00243979 | 0.01077144 |
| sp\|Q9NQR4\| | 272584426 | 397805726 | 0.35988012 | 0.00243979 | 0.01077144 |
| sp\|P05067\|A | 547139.36 | 376018.05 | -0.6557376 | 0.00243979 | 0.01077144 |
| sp\|O14735\|C | 139388.636 | 53227.1899 | -2.5358279 | 0.00243979 | 0.01077144 |
| sp\|O43657\|T | 462391.581 | 715594.171 | 0.53718237 | 0.00243979 | 0.01077144 |
| sp\|P05107\|IT | 20475138.3 | 33229484.8 | 0.52250932 | 0.00243979 | 0.01077144 |
| sp\|O14772\|F | 7203429.42 | 5712327.24 | -0.411856 | 0.00243979 | 0.01077144 |
| sp\|P07195\|L | 8041285658 | 1.0319E+10 | 0.362386 | 0.00243979 | 0.01077144 |
| sp\|Q8TDI0\|C | 6680928.29 | 9245715.93 | 0.50790413 | 0.00243979 | 0.01077144 |
| sp\|Q96RP9\|E | 20762978.4 | 15758336.8 | -0.4565098 | 0.00243979 | 0.01077144 |
| sp\|P18859\|A | 24835540.6 | 35348628.9 | 0.73365585 | 0.00243979 | 0.01077144 |
| sp\|Q16836\|H | 743493437 | 582305305 | -0.4016709 | 0.00243979 | 0.01077144 |
| sp\|P33241\|L | 19381830.9 | 40455812 | 0.77529574 | 0.00266212 | 0.01143735 |
| sp\|Q9UDT6\|C | 3849413.02 | 3056482.15 | -0.4019972 | 0.00266212 | 0.01143735 |
| sp\|Q5M9N0\| | 1574622.24 | 2193784.08 | 0.53377677 | 0.00266212 | 0.01143735 |
| sp\|O95470\|S | 6057409.11 | 8053149.64 | 0.34709601 | 0.00266212 | 0.01143735 |

| | | | | | |
|---|---|---|---|---|---|
| sp\|P50991\|T | 42734678.4 | 32668081.8 | -0.4688528 | 0.00266212 | 0.01143735 |
| sp\|P32942\|I( | 19678334 | 10509450.3 | -0.8101647 | 0.00266212 | 0.01143735 |
| sp\|P00742\|F | 36704224.5 | 56362575.1 | 0.56888763 | 0.00266212 | 0.01143735 |
| sp\|O75348\|\ | 2623595.03 | 4249620.32 | 0.62762773 | 0.00266212 | 0.01143735 |
| sp\|P46934\|N | 944459.737 | 625989.832 | -0.6269386 | 0.00266212 | 0.01143735 |
| sp\|O43464\|H | 37158778.2 | 29187380.9 | -0.3471281 | 0.00266212 | 0.01143735 |
| sp\|P04114\|A | 61697662.7 | 111057847 | 0.57692726 | 0.00266212 | 0.01143735 |
| sp\|Q9P2D8\|U | 2714439.27 | 4454617.26 | 0.60869097 | 0.00266212 | 0.01143735 |
| sp\|Q14344\|( | 78219881.5 | 63539152.2 | -0.347215 | 0.00266212 | 0.01143735 |
| sp\|P09496\|C | 95028742.6 | 125938046 | 0.35496714 | 0.00266212 | 0.01143735 |
| sp\|Q8WUH6\| | 3003067.78 | 4181918.91 | 0.41713912 | 0.00266212 | 0.01143735 |
| sp\|P51814\|Z | 11075643 | 15638298 | 0.4722182 | 0.00266212 | 0.01143735 |
| sp\|O15212\|F | 17176317.8 | 21978037.1 | 0.41027098 | 0.00266212 | 0.01143735 |
| sp\|Q12849\|( | 1676078.44 | 1077024.63 | -0.7912922 | 0.00266212 | 0.01143735 |
| sp\|Q9Y608\|L | 4780536.03 | 7201456.07 | 0.62010475 | 0.00266212 | 0.01143735 |
| sp\|P63272\|S | 459057.21 | 237605.422 | -1.0165932 | 0.00266212 | 0.01143735 |
| sp\|P13639\|E | 797925483 | 561958521 | -0.5702093 | 0.00266212 | 0.01143735 |
| sp\|Q13243\|S | 19837850.2 | 10447454 | -1.1365918 | 0.00266212 | 0.01143735 |
| sp\|P04844\|R | 6482377.57 | 4242939.2 | -0.7083033 | 0.00266212 | 0.01143735 |
| sp\|P05165\|P | 154682296 | 110302861 | -0.5896583 | 0.00266212 | 0.01143735 |
| sp\|P06756\|I1 | 44598640.2 | 60255621.6 | 0.37272717 | 0.00266212 | 0.01143735 |
| sp\|Q8TCZ2\|( | 631969.893 | 969943.929 | 0.86189486 | 0.00266212 | 0.01143735 |
| sp\|P07339\|C | 1022441512 | 1430795701 | 0.39255665 | 0.00266212 | 0.01143735 |
| sp\|P40925\|M | 3329234655 | 4936400999 | 0.43741204 | 0.00266212 | 0.01143735 |
| sp\|Q13501\|S | 6325015.41 | 8739037.4 | 0.37249332 | 0.00266212 | 0.01143735 |
| sp\|O00754\|N | 43527367.4 | 67356282.1 | 0.53249974 | 0.00266212 | 0.01143735 |
| sp\|P31946\|1 | 1250064209 | 1459012960 | 0.20037772 | 0.00290199 | 0.01230268 |
| sp\|Q14956\|( | 36738530.2 | 57540082 | 0.51565603 | 0.00290199 | 0.01230268 |
| sp\|P15260\|IN | 3952373.03 | 6246757.75 | 0.55212687 | 0.00290199 | 0.01230268 |
| sp\|P63313\|T | 58853597.2 | 105902277 | 0.60573612 | 0.00290199 | 0.01230268 |
| sp\|P60903\|S | 71968137.8 | 90295438.8 | 0.2791259 | 0.00290199 | 0.01230268 |
| sp\|Q9BV57\|N | 8304106.28 | 13957084.6 | 0.54477769 | 0.00290199 | 0.01230268 |
| sp\|Q9H488\|( | 57567607.2 | 73163280.8 | 0.34012696 | 0.00290199 | 0.01230268 |
| sp\|P16219\|A | 462664082 | 702950789 | 0.31581176 | 0.00290199 | 0.01230268 |
| sp\|P41219\|P | 4064579649 | 4950464444 | 0.26006678 | 0.00290199 | 0.01230268 |
| sp\|Q8IWE2\|N | 278031775 | 350468598 | 0.30064825 | 0.00290199 | 0.01230268 |
| sp\|P51572\|B | 51473616.1 | 39730907.9 | -0.522104 | 0.00290199 | 0.01230268 |
| sp\|P52597\|H | 124720514 | 91522036.8 | -0.4870651 | 0.00290199 | 0.01230268 |
| sp\|Q4VC31\|I | 7854562.28 | 10892442.9 | 0.5149196 | 0.00290199 | 0.01230268 |

| | | | | | |
|---|---|---|---|---|---|
| sp\|Q9UJN7\|Z | 2095742.57 | 2904560.7 | 0.46758774 | 0.00290199 | 0.01230268 |
| sp\|P06730\|IF | 33966031.6 | 25502925 | -0.4660225 | 0.00290199 | 0.01230268 |
| sp\|Q9H1P3\|( | 1793762.18 | 2437259.51 | 0.46120065 | 0.00316053 | 0.01316613 |
| sp\|P46821\|M | 253779145 | 330815772 | 0.3158113 | 0.00316053 | 0.01316613 |
| sp\|O75781\|F | 146954037 | 183566858 | 0.28535521 | 0.00316053 | 0.01316613 |
| sp\|P41218\|M | 9932898.71 | 21607822 | 0.59344365 | 0.00316053 | 0.01316613 |
| sp\|P25098\|A | 449552.618 | 711095.172 | 0.59956939 | 0.00316053 | 0.01316613 |
| sp\|Q9NYF8\|E | 29661495.1 | 20976354.8 | -0.6603503 | 0.00316053 | 0.01316613 |
| sp\|P78536\|A | 1838449.6 | 2803013.39 | 0.49471251 | 0.00316053 | 0.01316613 |
| sp\|O43719\|H | 2240283.31 | 1428211.1 | -0.5769987 | 0.00316053 | 0.01316613 |
| sp\|Q9H2D6\|T | 37861796.3 | 33001568.3 | -0.2129397 | 0.00316053 | 0.01316613 |
| sp\|Q9Y3B8\|( | 47599885.5 | 56128639.3 | 0.21440695 | 0.00316053 | 0.01316613 |
| sp\|P19404\|N | 67185594.9 | 101711216 | 0.40835205 | 0.00316053 | 0.01316613 |
| sp\|Q8N987\|I | 1430646.93 | 897747.316 | -0.8250232 | 0.00316053 | 0.01316613 |
| sp\|Q8NDW4\| | 1824270.67 | 1004758.68 | -1.7130387 | 0.00316053 | 0.01316613 |
| sp\|A0A0J9YV | 479284570 | 626476095 | 0.40616085 | 0.00316053 | 0.01316613 |
| sp\|Q86WU2\| | 67603111.4 | 43563314.9 | -0.3812457 | 0.00316053 | 0.01316613 |
| sp\|P60896\|S | 165311.472 | 242568.498 | 0.69095899 | 0.00316053 | 0.01316613 |
| sp\|P12694\|C | 40725664.6 | 29877175.1 | -0.4501313 | 0.00316053 | 0.01316613 |
| sp\|Q01955\|( | 4481432.45 | 8145629.33 | 0.94181807 | 0.00316053 | 0.01316613 |
| sp\|P02679\|F | 2591637335 | 1886466626 | -0.6974436 | 0.00316053 | 0.01316613 |
| sp\|P62899\|R | 42251660.7 | 25844568.3 | -0.6843357 | 0.00316053 | 0.01316613 |
| sp\|O15118\|N | 809244.557 | 1109159.77 | 0.44339961 | 0.00343899 | 0.01406968 |
| sp\|P62750\|R | 39879335.2 | 27607700.3 | -0.5405927 | 0.00343899 | 0.01406968 |
| sp\|Q8N0X4\|( | 8386102.5 | 6322120.2 | -0.3146233 | 0.00343899 | 0.01406968 |
| sp\|P17655\|C | 79483744.2 | 57900182.6 | -0.4614871 | 0.00343899 | 0.01406968 |
| sp\|P16401\|H | 80568197.3 | 135638164 | 0.61330953 | 0.00343899 | 0.01406968 |
| sp\|Q8IY95\|TI | 1733027.11 | 2487103.89 | 0.44818755 | 0.00343899 | 0.01406968 |
| sp\|Q9H7D7\|' | 1201190.75 | 1584108.42 | 0.36992178 | 0.00343899 | 0.01406968 |
| sp\|Q9NZL4\|H | 2325133.62 | 1664243.4 | -0.5370659 | 0.00343899 | 0.01406968 |
| sp\|Q5JTH9\|R | 2441567.84 | 3244673.34 | 0.39927229 | 0.00343899 | 0.01406968 |
| sp\|Q8NEZ2\|V | 838066.215 | 1105404.91 | 0.33555754 | 0.00343899 | 0.01406968 |
| sp\|Q92598\|H | 128348807 | 102047679 | -0.3584616 | 0.00343899 | 0.01406968 |
| sp\|Q9UMS6\| | 55314850 | 81660082.6 | 0.47293767 | 0.00343899 | 0.01406968 |
| sp\|Q16585\|S | 1189659.92 | 1776062.42 | 0.5651733 | 0.00343899 | 0.01406968 |
| sp\|P61619\|S | 2193047.39 | 1403906.34 | -0.6543456 | 0.00343899 | 0.01406968 |
| sp\|Q9Y3C4\|T | 988475.416 | 657728.869 | -0.6724412 | 0.00343899 | 0.01406968 |
| sp\|Q9NSK7\|( | 1998770.92 | 2950204.15 | 0.52957015 | 0.00343899 | 0.01406968 |
| sp\|Q9NP61\|/ | 4722401.6 | 6233656.95 | 0.3870801 | 0.00343899 | 0.01406968 |

| | | | | | |
|---|---|---|---|---|---|
| sp\|O43809\|( | 27648982.9 | 22519051.4 | -0.3162203 | 0.00343899 | 0.01406968 |
| sp\|P16109\|L | 10673674.8 | 14924336.2 | 0.50107991 | 0.00343899 | 0.01406968 |
| sp\|Q6P2I3\|F/ | 94609951.5 | 134063922 | 0.33819456 | 0.00343899 | 0.01406968 |
| sp\|Q9Y6M9\|I | 10583063 | 7052086.29 | -0.5755153 | 0.00343899 | 0.01406968 |
| sp\|Q9P0V9\|S | 88589589.7 | 67941729.5 | -0.3686191 | 0.00373862 | 0.01493891 |
| sp\|P10114\|R | 928452.661 | 1187984.18 | 0.39733683 | 0.00373862 | 0.01493891 |
| sp\|Q6P1N9\|T | 55620232.9 | 77316019.7 | 0.3442021 | 0.00373862 | 0.01493891 |
| sp\|P10109\|A | 23937869.8 | 34428285.4 | 0.4073438 | 0.00373862 | 0.01493891 |
| sp\|P02675\|F | 2178592415 | 1596228652 | -0.6939556 | 0.00373862 | 0.01493891 |
| sp\|Q7Z4I7\|LI | 102806432 | 128584929 | 0.29522292 | 0.00373862 | 0.01493891 |
| sp\|Q71UI9\|H | 161811334 | 94798874.9 | -0.7145544 | 0.00373862 | 0.01493891 |
| sp\|Q6UX04\|( | 2019725.87 | 1474378.82 | -0.5480061 | 0.00373862 | 0.01493891 |
| sp\|P50748\|K | 2668385.92 | 2128955.97 | -0.3488279 | 0.00373862 | 0.01493891 |
| sp\|Q15291\|F | 4536251.39 | 6456349.71 | 0.45302218 | 0.00373862 | 0.01493891 |
| sp\|Q9H4F8\|S | 6129118.8 | 9588149.11 | 0.47546516 | 0.00373862 | 0.01493891 |
| sp\|P25705\|A | 1106349993 | 1387067880 | 0.37342948 | 0.00373862 | 0.01493891 |
| sp\|Q9C0C2\|T | 305367482 | 380584933 | 0.27611267 | 0.00373862 | 0.01493891 |
| sp\|Q07866\|K | 1642746.29 | 1305362.67 | -0.5061453 | 0.00373862 | 0.01493891 |
| sp\|Q8N5G2\| | 519092.624 | 778289.392 | 0.47555603 | 0.00373862 | 0.01493891 |
| sp\|Q8N3V7\|S | 10033918.1 | 13478268.5 | 0.403716 | 0.00373862 | 0.01493891 |
| sp\|Q96N66\|I | 161098.425 | 105109.21 | -1.2976111 | 0.00373862 | 0.01493891 |
| sp\|P61158\|A | 169124405 | 119962225 | -0.6016083 | 0.00373862 | 0.01493891 |
| sp\|Q96AG4\|I | 2751688.47 | 1830368.31 | -0.6905791 | 0.00373862 | 0.01493891 |
| sp\|A0A0B4J2 | 325633644 | 369218057 | 0.23239643 | 0.00373862 | 0.01493891 |
| sp\|O60216\|F | 7161029.72 | 10554859.2 | 0.5399978 | 0.00373862 | 0.01493891 |
| sp\|O60271\|J | 120024021 | 141672413 | 0.23076067 | 0.00373862 | 0.01493891 |
| sp\|Q9Y2T2\|A | 20758120.1 | 28574500.5 | 0.40701195 | 0.00373862 | 0.01493891 |
| sp\|Q96S99\|F | 6324554.32 | 4268747.4 | -0.6604412 | 0.00373862 | 0.01493891 |
| sp\|O75191\|X | 21027180.4 | 29283395.5 | 0.3602536 | 0.00373862 | 0.01493891 |
| sp\|P20853\|C | 2067950 | 1528590.7 | -0.488747 | 0.00373862 | 0.01493891 |
| sp\|O75369\|F | 1215407516 | 919032581 | -0.4653113 | 0.00373862 | 0.01493891 |
| sp\|Q8NFV4\|/ | 196878.916 | 421598.388 | 1.18876657 | 0.00373862 | 0.01493891 |
| sp\|Q13642\|F | 3633610831 | 4695543439 | 0.3100507 | 0.00406074 | 0.01603908 |
| sp\|Q9BUJ2\|H | 223961814 | 166545198 | -0.4946515 | 0.00406074 | 0.01603908 |
| sp\|P32456\|G | 1630676.53 | 2064210.58 | 0.34383151 | 0.00406074 | 0.01603908 |
| sp\|P11233\|R | 21239065.9 | 26786528.3 | 0.33263051 | 0.00406074 | 0.01603908 |
| sp\|Q6PI48\|S | 12870467.9 | 10139779.8 | -0.3958047 | 0.00406074 | 0.01603908 |
| sp\|P05997\|C | 1154829.1 | 1751792.47 | 0.45055927 | 0.00406074 | 0.01603908 |
| sp\|P46779\|R | 20801333.2 | 10462908.7 | -0.9112782 | 0.00406074 | 0.01603908 |

| | | | | | |
|---|---|---|---|---|---|
| sp\|P04066\|F | 35563476.2 | 27649181.4 | -0.4598031 | 0.00406074 | 0.01603908 |
| sp\|P19174\|P | 4496480.26 | 3769934.93 | -0.2378502 | 0.00406074 | 0.01603908 |
| sp\|Q12792\|T | 39446366.2 | 30559801.1 | -0.4553239 | 0.00406074 | 0.01603908 |
| sp\|P00338\|L | 7377152331 | 9336381270 | 0.2887762 | 0.00406074 | 0.01603908 |
| sp\|O75533\|S | 50322787.8 | 40983696.9 | -0.2717537 | 0.00406074 | 0.01603908 |
| sp\|O75618\|D | 2716129.54 | 4592607.78 | 0.84537574 | 0.00406074 | 0.01603908 |
| sp\|O43639\|N | 10366665.6 | 8095054.34 | -0.4143014 | 0.00406074 | 0.01603908 |
| sp\|P52758\|R | 92542186.4 | 106670954 | 0.28326611 | 0.00440677 | 0.01708247 |
| sp\|Q8WUA2\| | 2973539.95 | 2329313.29 | -0.4878991 | 0.00440677 | 0.01708247 |
| sp\|O43684\|E | 30518977.3 | 23202946.4 | -0.3094598 | 0.00440677 | 0.01708247 |
| sp\|P53597\|S | 187997321 | 140335992 | -0.5020531 | 0.00440677 | 0.01708247 |
| sp\|P19634\|S | 1085668.38 | 1735824.29 | 0.65474606 | 0.00440677 | 0.01708247 |
| sp\|Q8N5M9\| | 98442.2651 | 60062.3955 | -1.3066531 | 0.00440677 | 0.01708247 |
| sp\|Q99784\|N | 1084805.95 | 1350062.21 | 0.3034597 | 0.00440677 | 0.01708247 |
| sp\|Q9NS15\|L | 2663227.93 | 3901535.75 | 0.46148928 | 0.00440677 | 0.01708247 |
| sp\|P19525\|E | 525926.665 | 715293.515 | 0.45584697 | 0.00440677 | 0.01708247 |
| sp\|P04792\|H | 2880497206 | 2311430863 | -0.2932173 | 0.00440677 | 0.01708247 |
| sp\|Q86V48\|L | 3896796.15 | 5213591.95 | 0.40619676 | 0.00440677 | 0.01708247 |
| sp\|Q9NZ01\|T | 14575340.5 | 11088191.7 | -0.5564231 | 0.00440677 | 0.01708247 |
| sp\|P99999\|C | 186943404 | 254199556 | 0.4351596 | 0.00440677 | 0.01708247 |
| sp\|P04156\|P | 1292613.5 | 1715809.45 | 0.36063155 | 0.00440677 | 0.01708247 |
| sp\|Q9NUQ3\| | 171664.102 | 300923.721 | 0.67990483 | 0.00440677 | 0.01708247 |
| sp\|P41252\|S | 25041867.5 | 19526753.5 | -0.3636173 | 0.00440677 | 0.01708247 |
| sp\|P35270\|S | 20422380.4 | 30927973.3 | 0.35809616 | 0.00440677 | 0.01708247 |
| sp\|P35232\|P | 31023603.4 | 24506909.3 | -0.3219443 | 0.00440677 | 0.01708247 |
| sp\|P11171\|E | 78051782.1 | 151085380 | 0.71785785 | 0.00440677 | 0.01708247 |
| sp\|Q9UBV2\|S | 3744463.98 | 5044350.73 | 0.44552542 | 0.00440677 | 0.01708247 |
| sp\|P17213\|B | 1668329.44 | 4444687.33 | 0.75719264 | 0.00440677 | 0.01708247 |
| sp\|Q6P1L8\|F | 5440014.94 | 3373559.61 | -0.7843791 | 0.00440677 | 0.01708247 |
| sp\|Q93077\|H | 250351492 | 143838959 | -0.824939 | 0.00440677 | 0.01708247 |
| sp\|O00151\|P | 132277912 | 183771378 | 0.37092126 | 0.00477815 | 0.01811245 |
| sp\|Q8WWX9 | 11470945.3 | 15761779 | 0.32482913 | 0.00477815 | 0.01811245 |
| sp\|Q92665\|F | 2332633.01 | 1643009.99 | -0.5498979 | 0.00477815 | 0.01811245 |
| sp\|Q96I99\|S | 264843551 | 191826486 | -0.5349214 | 0.00477815 | 0.01811245 |
| sp\|Q8WZA0\| | 19365991.1 | 24981726.7 | 0.35847347 | 0.00477815 | 0.01811245 |
| sp\|Q8TBF8\|F | 24212508.5 | 36331419 | 0.4685741 | 0.00477815 | 0.01811245 |
| sp\|P98172\|E | 4821362.24 | 5896322.63 | 0.2987954 | 0.00477815 | 0.01811245 |
| sp\|Q86Y46\|K | 1100661031 | 633184919 | -0.7880698 | 0.00477815 | 0.01811245 |
| sp\|P02743\|S | 299601136 | 208708079 | -0.6325582 | 0.00477815 | 0.01811245 |

| | | | | | |
|---|---|---|---|---|---|
| sp\|O95445\|A | 33148722.8 | 51147381.5 | 0.430301 | 0.00477815 | 0.01811245 |
| sp\|Q92974\|A | 9683491.16 | 13544133.9 | 0.47335055 | 0.00477815 | 0.01811245 |
| sp\|Q7L0Y3\|T | 15914774.7 | 12871106.9 | -0.3652188 | 0.00477815 | 0.01811245 |
| sp\|O95678\|K | 1492697373 | 697829600 | -0.9721945 | 0.00477815 | 0.01811245 |
| sp\|P47895\|A | 1125408713 | 817432136 | -0.5404204 | 0.00477815 | 0.01811245 |
| sp\|Q10567\|A | 101253598 | 83481736.9 | -0.2462553 | 0.00477815 | 0.01811245 |
| sp\|Q9UHV9\| | 41355657.3 | 52687397 | 0.32166259 | 0.00477815 | 0.01811245 |
| sp\|Q13496\|M | 2823163.71 | 2319722.14 | -0.3088588 | 0.00477815 | 0.01811245 |
| sp\|Q9P016\|T | 7026779.35 | 4993578.05 | -0.5273817 | 0.00477815 | 0.01811245 |
| sp\|P35080\|P | 48881598.3 | 37797676.1 | -0.4141219 | 0.00477815 | 0.01811245 |
| sp\|P30304\|M | 4124052.69 | 5022041.56 | 0.29990212 | 0.00477815 | 0.01811245 |
| sp\|Q9BZ23\|F | 4122753.93 | 3380238.99 | -0.3418313 | 0.00477815 | 0.01811245 |
| sp\|Q9BXJ8\|T | 1031562.38 | 612462.831 | -1.6002063 | 0.00477815 | 0.01811245 |
| sp\|Q9NX58\|L | 748892.594 | 590425.496 | -0.6251941 | 0.00477815 | 0.01811245 |
| sp\|Q15746\|M | 92785165.3 | 63633873.9 | -0.4224702 | 0.00477815 | 0.01811245 |
| sp\|Q9HCX4\|T | 314635.721 | 423378.73 | 0.55164259 | 0.00477815 | 0.01811245 |
| sp\|O75175\|C | 6853533.79 | 9955300.54 | 0.42891409 | 0.00477815 | 0.01811245 |
| sp\|Q9NXA8\|S | 9770142.25 | 7482525.81 | -0.4305587 | 0.00477815 | 0.01811245 |
| sp\|Q8N474\|S | 13518605.1 | 9549532.46 | -0.5621952 | 0.00477815 | 0.01811245 |
| sp\|Q9BYC9\|I | 5579342.42 | 8557868.68 | 0.5263639 | 0.0051764 | 0.01930191 |
| sp\|Q4J6C6\|P | 16829049.8 | 14565397.2 | -0.2454246 | 0.0051764 | 0.01930191 |
| sp\|P12277\|K | 563044340 | 392293960 | -0.7315919 | 0.0051764 | 0.01930191 |
| sp\|P19387\|R | 988914.587 | 893320.893 | -0.5139139 | 0.0051764 | 0.01930191 |
| sp\|Q08495\|D | 59716364.6 | 97298519.6 | 0.69663861 | 0.0051764 | 0.01930191 |
| sp\|O95819\|M | 1726404.85 | 2588068.48 | 0.7151492 | 0.0051764 | 0.01930191 |
| sp\|Q9NRY5\|I | 5047512.7 | 6480064.59 | 0.38260823 | 0.0051764 | 0.01930191 |
| sp\|P02786\|T | 33591028.3 | 43645587.5 | 0.34930126 | 0.0051764 | 0.01930191 |
| sp\|Q9H8L6\|M | 18103167.2 | 22140803 | 0.33180387 | 0.0051764 | 0.01930191 |
| sp\|Q5NDL2\|I | 6590269.6 | 9326300.98 | 0.41916227 | 0.0051764 | 0.01930191 |
| sp\|Q9H1K0\|I | 3999093.16 | 6817322.27 | 0.58719842 | 0.0051764 | 0.01930191 |
| sp\|Q01518\|C | 128169390 | 164325388 | 0.31547588 | 0.0051764 | 0.01930191 |
| sp\|P56192\|S | 17403071.5 | 11369552 | -0.6901075 | 0.0051764 | 0.01930191 |
| sp\|O75340\|F | 18499020.8 | 13196751.4 | -0.5917507 | 0.0051764 | 0.01930191 |
| sp\|Q8NFQ8\|T | 81493.506 | 55468.3381 | -0.5656112 | 0.0051764 | 0.01930191 |
| sp\|Q9Y255\|F | 7955494.99 | 10765537.4 | 0.43369399 | 0.0051764 | 0.01930191 |
| sp\|Q9Y250\|L | 8253234.94 | 11120247 | 0.43077621 | 0.0051764 | 0.01930191 |
| sp\|P82663\|R | 1345775.76 | 826684.304 | -0.863295 | 0.0051764 | 0.01930191 |
| sp\|Q8WYJ6\|S | 647525.578 | 463813.792 | -0.5077905 | 0.0051764 | 0.01930191 |
| sp\|Q9Y5J6\|T | 703838.961 | 1008519.24 | 0.54384621 | 0.0051764 | 0.01930191 |

| | | | | | |
|---|---|---|---|---|---|
| sp\|P63173\|R | 9027841.8 | 5423352.37 | -0.7684833 | 0.0051764 | 0.01930191 |
| sp\|Q9C0D2\| | 54643119.8 | 70936425.1 | 0.32769902 | 0.00560315 | 0.02047945 |
| sp\|P80297\|M | 23011933.2 | 39701979.9 | 0.60817194 | 0.00560315 | 0.02047945 |
| sp\|P37840\|S | 172949560 | 307535420 | 0.6926197 | 0.00560315 | 0.02047945 |
| sp\|P06748\|N | 150950831 | 119711235 | -0.3805518 | 0.00560315 | 0.02047945 |
| sp\|Q6ZSJ8\|C | 743512.793 | 1370074.35 | 0.78407416 | 0.00560315 | 0.02047945 |
| sp\|Q92934\|E | 3862503.73 | 4672158.82 | 0.28552934 | 0.00560315 | 0.02047945 |
| sp\|Q6ZVC0\|N | 6348482.08 | 7806544.94 | 0.27120443 | 0.00560315 | 0.02047945 |
| sp\|Q6NXT6\|T | 3637027.07 | 2616243.01 | -0.495427 | 0.00560315 | 0.02047945 |
| sp\|P61457\|P | 231599272 | 287513677 | 0.30303344 | 0.00560315 | 0.02047945 |
| sp\|O15247\|C | 34533410 | 24418624 | -0.5558618 | 0.00560315 | 0.02047945 |
| sp\|Q9H3P7\| | 2693126.02 | 1935494.15 | -0.4628739 | 0.00560315 | 0.02047945 |
| sp\|Q9P2T1\|G | 106616388 | 133062411 | 0.29835065 | 0.00560315 | 0.02047945 |
| sp\|P55795\|H | 162050072 | 115149876 | -0.5779246 | 0.00560315 | 0.02047945 |
| sp\|O60232\|Z | 1841370.03 | 2582655.2 | 0.54387184 | 0.00560315 | 0.02047945 |
| sp\|Q14314\|F | 1807572.21 | 2997961.53 | 0.64742636 | 0.00560315 | 0.02047945 |
| sp\|Q16864\|V | 1866175.79 | 1954375.64 | 0.57504853 | 0.00560315 | 0.02047945 |
| sp\|Q04837\|S | 41728188.3 | 71850293.7 | 0.72500693 | 0.00560315 | 0.02047945 |
| sp\|O75594\|F | 1613848.3 | 6243378.69 | 0.86235573 | 0.00560315 | 0.02047945 |
| sp\|O75153\|C | 8619059.11 | 6343721.99 | -0.5797567 | 0.00560315 | 0.02047945 |
| sp\|P53816\|P | 4457194.13 | 5862686.45 | 0.4645025 | 0.00560315 | 0.02047945 |
| sp\|Q9Y4K1\|C | 14955708.3 | 17661978.5 | 0.22526539 | 0.00560315 | 0.02047945 |
| sp\|P41091\|IF | 25396244.2 | 19804772.6 | -0.3737223 | 0.00560315 | 0.02047945 |
| sp\|Q96GK7\|I | 65736625.9 | 94330600.3 | 0.32885285 | 0.00560315 | 0.02047945 |
| sp\|Q15836\|V | 10878364.3 | 13435582.1 | 0.26483834 | 0.00560315 | 0.02047945 |
| sp\|P55786\|P | 632446519 | 489212951 | -0.4222145 | 0.00560315 | 0.02047945 |
| sp\|P78386\|K | 877075555 | 1018787784 | 0.22103133 | 0.00560315 | 0.02047945 |
| sp\|Q13201\|N | 3603553.87 | 2408546.31 | -0.7747471 | 0.00606003 | 0.02170304 |
| sp\|P49368\|T | 267777336 | 207816132 | -0.4149241 | 0.00606003 | 0.02170304 |
| sp\|P80303\|N | 26256077.2 | 34166583 | 0.43263747 | 0.00606003 | 0.02170304 |
| sp\|P42167\|L | 33894483.3 | 23856989.5 | -0.5073951 | 0.00606003 | 0.02170304 |
| sp\|Q96KX1\|C | 36621063.5 | 65399877.1 | 0.73471853 | 0.00606003 | 0.02170304 |
| sp\|Q9UHQ4\| | 4773120.57 | 6650397.91 | 0.41622042 | 0.00606003 | 0.02170304 |
| sp\|Q14BN4\|S | 2401469.18 | 3251759.39 | 0.42079733 | 0.00606003 | 0.02170304 |
| sp\|O43617\|T | 9751582.44 | 8008378.79 | -0.3737297 | 0.00606003 | 0.02170304 |
| sp\|P47897\|S | 190186407 | 131800519 | -0.6293512 | 0.00606003 | 0.02170304 |
| sp\|Q9NQX5\| | 73900.9348 | 131966.716 | 1.06306645 | 0.00606003 | 0.02170304 |
| sp\|P08174\|D | 15606106.1 | 21725815.9 | 0.43955539 | 0.00606003 | 0.02170304 |
| sp\|P58557\|Y | 1973005.97 | 2750880.45 | 0.53015736 | 0.00606003 | 0.02170304 |

| | | | | | |
|---|---|---|---|---|---|
| sp\|P54886\|P | 8931714.55 | 6600999.4 | -0.5506711 | 0.00606003 | 0.02170304 |
| sp\|P51149\|R | 258428140 | 327379111 | 0.30862093 | 0.00606003 | 0.02170304 |
| sp\|P30048\|P | 978271450 | 1150708416 | 0.20149352 | 0.00606003 | 0.02170304 |
| sp\|Q9UJA5\|T | 637930.328 | 465533.096 | -0.5813615 | 0.00606003 | 0.02170304 |
| sp\|O75494\|S | 5341052.17 | 3672548.25 | -0.5039284 | 0.00606003 | 0.02170304 |
| sp\|P40189\|IL | 1608671.93 | 2300179.44 | 0.48861844 | 0.00606003 | 0.02170304 |
| sp\|P29466\|C | 37269231.2 | 28771749.3 | -0.35045 | 0.00606003 | 0.02170304 |
| sp\|Q9BYB0\|S | 12329970.8 | 17963401.7 | 0.4925633 | 0.00606003 | 0.02170304 |
| sp\|O95197\|F | 4570303.09 | 6015251.75 | 0.3199336 | 0.00606003 | 0.02170304 |
| sp\|Q9UBT2\|S | 166120100 | 130701990 | -0.4162951 | 0.00606003 | 0.02170304 |
| sp\|P21399\|A | 1322565958 | 1664645183 | 0.34650064 | 0.00606003 | 0.02170304 |
| sp\|P61024\|C | 3879663.56 | 5172147.35 | 0.43749943 | 0.00606003 | 0.02170304 |
| sp\|Q01130\|S | 37747791.1 | 25549925.1 | -0.5744917 | 0.00606003 | 0.02170304 |
| sp\|Q12923\|F | 792611.011 | 552837.685 | -0.5003732 | 0.00606003 | 0.02170304 |
| sp\|P08637\|F | 11298245.3 | 16970635.2 | 0.4947056 | 0.00606003 | 0.02170304 |
| sp\|Q9H3M0\| | 6058011.08 | 8505857.34 | 0.50771891 | 0.00654876 | 0.02310846 |
| sp\|P09651\|R | 834252104 | 658420701 | -0.3703504 | 0.00654876 | 0.02310846 |
| sp\|Q5XKE5\|K | 1444474600 | 646315613 | -0.9911696 | 0.00654876 | 0.02310846 |
| sp\|Q8IWL3\|H | 2431994.56 | 3217248.67 | 0.40168138 | 0.00654876 | 0.02310846 |
| sp\|O95810\|C | 592379083 | 688468240 | 0.22170527 | 0.00654876 | 0.02310846 |
| sp\|Q7Z4V5\|H | 32410430.1 | 23285666.7 | -0.598097 | 0.00654876 | 0.02310846 |
| sp\|Q6NVY1\|I | 45259461 | 28946880.7 | -0.8862839 | 0.00654876 | 0.02310846 |
| sp\|Q9UPP1\|I | 1931095.58 | 2708090.25 | 0.43731804 | 0.00654876 | 0.02310846 |
| sp\|Q7Z5R6\|A | 5661416.65 | 7326599.62 | 0.38047511 | 0.00654876 | 0.02310846 |
| sp\|Q9BTE6\|A | 6660597.15 | 8436000.53 | 0.34910119 | 0.00654876 | 0.02310846 |
| sp\|P09874\|P | 22352146.2 | 18106746.7 | -0.281531 | 0.00654876 | 0.02310846 |
| sp\|Q9NYC9\|I | 5202071.07 | 3801614.26 | -0.5007997 | 0.00654876 | 0.02310846 |
| sp\|Q9BRK5\|C | 2056018.18 | 2716511.36 | 0.40783719 | 0.00654876 | 0.02310846 |
| sp\|Q9Y2W1\|T | 8635144.4 | 6431146.49 | -0.4474941 | 0.00654876 | 0.02310846 |
| sp\|P52789\|H | 27665857 | 32609709.3 | 0.24344231 | 0.00654876 | 0.02310846 |
| sp\|Q9UBQ0\| | 87401636.5 | 110473967 | 0.23554283 | 0.00654876 | 0.02310846 |
| sp\|P53396\|A | 380444519 | 280834712 | -0.730282 | 0.00654876 | 0.02310846 |
| sp\|Q14318\|F | 8370429.31 | 7042193.97 | -0.2650267 | 0.00654876 | 0.02310846 |
| sp\|P63208\|S | 8502858.07 | 10644512.6 | 0.29097279 | 0.00654876 | 0.02310846 |
| sp\|Q13423\|N | 45861192 | 35223591.9 | -0.3741381 | 0.00654876 | 0.02310846 |
| sp\|O43414\|E | 11971857.9 | 9552236.38 | -0.4143125 | 0.0070712 | 0.02459034 |
| sp\|O75015\|F | 7952453.64 | 13655073.4 | 0.52083996 | 0.0070712 | 0.02459034 |
| sp\|Q9H223\|E | 42055145.7 | 53595024 | 0.31450637 | 0.0070712 | 0.02459034 |
| sp\|Q9P1T7\|N | 380498.589 | 529826.905 | 0.34438927 | 0.0070712 | 0.02459034 |

| | | | | | |
|---|---|---|---|---|---|
| sp\|Q9H4G0\| | 71549834.2 | 90542873.2 | 0.30255344 | 0.0070712 | 0.02459034 |
| sp\|P51798\|C | 14218911.2 | 17691118.5 | 0.29938753 | 0.0070712 | 0.02459034 |
| sp\|Q96C19\|E | 21169994 | 27442652.1 | 0.30031063 | 0.0070712 | 0.02459034 |
| sp\|P36551\|H | 17828036.7 | 24781616.4 | 0.34979758 | 0.0070712 | 0.02459034 |
| sp\|P43487\|R | 80716500.8 | 94824077.9 | 0.22190002 | 0.0070712 | 0.02459034 |
| sp\|Q9UFN0\|I | 163745102 | 206004769 | 0.32092528 | 0.0070712 | 0.02459034 |
| sp\|Q12802\|A | 1025396.42 | 738662.406 | -0.5861836 | 0.0070712 | 0.02459034 |
| sp\|Q96BJ3\|A | 103536553 | 79892704 | -0.4042406 | 0.0070712 | 0.02459034 |
| sp\|P04899\|G | 135211616 | 108024613 | -0.3642028 | 0.0070712 | 0.02459034 |
| sp\|Q9Y3Z3\|S | 55733376.6 | 36416844.9 | -0.5975238 | 0.0070712 | 0.02459034 |
| sp\|P41250\|G | 99987435.6 | 129203396 | 0.31550658 | 0.0070712 | 0.02459034 |
| sp\|Q8WYQ3\| | 1592186.77 | 1500177.87 | 1.48126288 | 0.0070712 | 0.02459034 |
| sp\|O15143\|A | 85880236.8 | 65446556.9 | -0.4799612 | 0.0070712 | 0.02459034 |
| sp\|Q8WVV4\| | 21752072.3 | 41306117.1 | 1.52806447 | 0.0070712 | 0.02459034 |
| sp\|P35222\|C | 63004854.9 | 50603066.3 | -0.3214141 | 0.0070712 | 0.02459034 |
| sp\|Q99471\|P | 20323857.1 | 25162674.8 | 0.23496039 | 0.0070712 | 0.02459034 |
| sp\|Q68EM7\|I | 949922.306 | 1182204.26 | 0.28846705 | 0.00762919 | 0.02615176 |
| sp\|P23528\|C | 899619727 | 1072198542 | 0.24752268 | 0.00762919 | 0.02615176 |
| sp\|O14494\|P | 2669361.46 | 3901826.93 | 0.24856754 | 0.00762919 | 0.02615176 |
| sp\|Q9Y5L4\|T | 13322931.4 | 16191221.8 | 0.31765649 | 0.00762919 | 0.02615176 |
| sp\|P62877\|R | 4583698.81 | 6120466.64 | 0.32327588 | 0.00762919 | 0.02615176 |
| sp\|Q9H2H8\| | 6682191.87 | 8480106.98 | 0.37851078 | 0.00762919 | 0.02615176 |
| sp\|Q9Y520\|P | 14306130.3 | 18645397.4 | 0.35035833 | 0.00762919 | 0.02615176 |
| sp\|Q9Y2E5\|N | 15857348.3 | 23406200.8 | 0.39765594 | 0.00762919 | 0.02615176 |
| sp\|O75874\|I | 2007961770 | 2499263785 | 0.24406137 | 0.00762919 | 0.02615176 |
| sp\|Q15436\|S | 33143808.4 | 24684832.7 | -0.4670883 | 0.00762919 | 0.02615176 |
| sp\|Q8IUZ5\|A | 88452539.9 | 138763011 | 0.40890975 | 0.00762919 | 0.02615176 |
| sp\|Q5JSZ5\|P | 1778172.45 | 3966754.68 | 1.31659078 | 0.00762919 | 0.02615176 |
| sp\|Q9NPI6\|D | 2002203.49 | 3211770.21 | 0.71855088 | 0.00762919 | 0.02615176 |
| sp\|Q02818\|N | 98776624.3 | 129644832 | 0.32748162 | 0.00762919 | 0.02615176 |
| sp\|P50895\|B | 285993663 | 233112584 | -0.2717748 | 0.00762919 | 0.02615176 |
| sp\|Q99538\|L | 18662015.1 | 14488066.5 | -0.4253776 | 0.00762919 | 0.02615176 |
| sp\|O75477\|E | 7571993.79 | 5591220.16 | -0.607874 | 0.00762919 | 0.02615176 |
| sp\|P12004\|P | 12534966.6 | 10372402.6 | -0.315482 | 0.00762919 | 0.02615176 |
| sp\|O76061\|S | 3581078.98 | 4497027.23 | 0.3079904 | 0.00762919 | 0.02615176 |
| sp\|Q93096\|T | 5146935.26 | 8492105.83 | 0.86416824 | 0.00762919 | 0.02615176 |
| sp\|P49789\|F | 1928195.94 | 2658385.3 | 0.37506612 | 0.00822467 | 0.02787442 |
| sp\|Q8IUI8\|CI | 3752331.95 | 5409540.54 | 0.49687408 | 0.00822467 | 0.02787442 |
| sp\|O43236\|S | 5789148.09 | 4429939.63 | -0.4331153 | 0.00822467 | 0.02787442 |

| | | | | | |
|---|---|---|---|---|---|
| sp\|P61326\|M | 485982.305 | 280985.675 | -1.4596493 | 0.00822467 | 0.02787442 |
| sp\|P08754\|G | 101795444 | 81520785.1 | -0.3588793 | 0.00822467 | 0.02787442 |
| sp\|P80188\|N | 13132259.1 | 48295914.1 | 0.94920456 | 0.00822467 | 0.02787442 |
| sp\|Q9NVE7\|I | 23274760.9 | 26185479.6 | 0.19586135 | 0.00822467 | 0.02787442 |
| sp\|P28070\|P | 147727701 | 211579741 | 0.42104596 | 0.00822467 | 0.02787442 |
| sp\|Q9BXR0\|T | 18803238.3 | 16437081.8 | -0.2020558 | 0.00822467 | 0.02787442 |
| sp\|Q8N0W3\| | 131225857 | 112483008 | -0.2576588 | 0.00822467 | 0.02787442 |
| sp\|P30040\|E | 149638855 | 129742254 | -0.2296711 | 0.00822467 | 0.02787442 |
| sp\|P14209\|C | 6374981.63 | 12534967.4 | 0.75246725 | 0.00822467 | 0.02787442 |
| sp\|Q8TAQ2\|S | 4236579.42 | 3123392.83 | -0.5072736 | 0.00822467 | 0.02787442 |
| sp\|Q5HYK7\|S | 41072941.1 | 32536699.7 | -0.3022254 | 0.00822467 | 0.02787442 |
| sp\|Q9NVM6\| | 2447290.59 | 3145227.74 | 0.32592085 | 0.00822467 | 0.02787442 |
| sp\|B3EWG6\| | 32088496.3 | 40135734.6 | 0.26543677 | 0.00822467 | 0.02787442 |
| sp\|Q9UIB8\|S | 494273.693 | 923012.136 | 0.67827755 | 0.00885972 | 0.02946485 |
| sp\|Q05397\|F | 4347500.06 | 5202582.59 | 0.21195353 | 0.00885972 | 0.02946485 |
| sp\|Q9P2J5\|S | 47952411.7 | 41733156.3 | -0.2265516 | 0.00885972 | 0.02946485 |
| sp\|O43747\|A | 947429.476 | 660467.985 | -0.8380663 | 0.00885972 | 0.02946485 |
| sp\|Q9H0P0\|5 | 3231618.09 | 2391703.45 | -0.542408 | 0.00885972 | 0.02946485 |
| sp\|P51580\|T | 2173159.74 | 1485227.45 | -0.8693498 | 0.00885972 | 0.02946485 |
| sp\|O94929\|A | 21417892.6 | 26332940.7 | 0.25107918 | 0.00885972 | 0.02946485 |
| sp\|Q8NFH8\|I | 2683426.67 | 4874088.1 | 0.76712109 | 0.00885972 | 0.02946485 |
| sp\|P14735\|ID | 79747574.3 | 94008541.6 | 0.23058918 | 0.00885972 | 0.02946485 |
| sp\|Q12805\|F | 182543163 | 253476378 | 0.38158125 | 0.00885972 | 0.02946485 |
| sp\|Q12904\|A | 116556180 | 87284726.5 | -0.4928879 | 0.00885972 | 0.02946485 |
| sp\|Q01970\|F | 21815340 | 17719362.7 | -0.3385208 | 0.00885972 | 0.02946485 |
| sp\|P16615\|A | 42343008.4 | 32114326.1 | -0.5704002 | 0.00885972 | 0.02946485 |
| sp\|Q8IXK0\|P | 1761713.76 | 2577538.23 | 0.37993296 | 0.00885972 | 0.02946485 |
| sp\|O95352\|A | 39252899.3 | 28896932 | -0.5832903 | 0.00885972 | 0.02946485 |
| sp\|Q9H3U5\| | 3194367.44 | 5085027.26 | 0.61523058 | 0.00885972 | 0.02946485 |
| sp\|O95793\|S | 6938216.51 | 5533857.51 | -0.3248746 | 0.00885972 | 0.02946485 |
| sp\|P57087\|JA | 692967.052 | 1168902.64 | 0.7929332 | 0.00885972 | 0.02946485 |
| sp\|P82673\|R | 2483277.16 | 1806158.93 | -0.6455226 | 0.00885972 | 0.02946485 |
| sp\|Q9BSL1\|U | 22517685.2 | 58574947 | 1.19754797 | 0.00885972 | 0.02946485 |
| sp\|Q16627\|C | 1004553.68 | 1780059.57 | 0.60716486 | 0.00885972 | 0.02946485 |
| sp\|Q15366\|F | 82738951.3 | 67033910.4 | -0.3570017 | 0.00885972 | 0.02946485 |
| sp\|Q13009\|T | 9123870.06 | 12802566.8 | 0.44705551 | 0.00885972 | 0.02946485 |
| sp\|A2VCK2\|D | 9123870.06 | 12802566.8 | 0.44705551 | 0.00885972 | 0.02946485 |
| sp\|Q71RC2\|I | 2775280.7 | 2668481.97 | -0.3713452 | 0.00885972 | 0.02946485 |
| sp\|P06858\|L | 3901548.1 | 4858828.37 | 0.31752619 | 0.00885972 | 0.02946485 |

| | | | | | |
|---|---|---|---|---|---|
| sp\|Q9UHD9\| | 40057961.2 | 48761474.1 | 0.26244703 | 0.00885972 | 0.02946485 |
| sp\|O43707\|A | 939641692 | 665806991 | -0.4992744 | 0.00953641 | 0.03119647 |
| sp\|P15502\|E | 2444533.97 | 3742944.11 | 0.73100493 | 0.00953641 | 0.03119647 |
| sp\|Q5TD97\|F | 18321192.8 | 13913864.9 | -0.3731906 | 0.00953641 | 0.03119647 |
| sp\|P16435\|N | 65423193.8 | 85837658 | 0.30824174 | 0.00953641 | 0.03119647 |
| sp\|Q9UBV8\|I | 10067139.3 | 8328680.3 | -0.3280732 | 0.00953641 | 0.03119647 |
| sp\|Q14012\|K | 1206036.12 | 1951998.12 | 0.44804165 | 0.00953641 | 0.03119647 |
| sp\|O15120\|F | 12905572.5 | 9500487.03 | -0.6343543 | 0.00953641 | 0.03119647 |
| sp\|Q92905\|C | 29164270.4 | 33210153.7 | 0.19275102 | 0.00953641 | 0.03119647 |
| sp\|P61160\|A | 172816684 | 129022982 | -0.497345 | 0.00953641 | 0.03119647 |
| sp\|Q13442\|H | 167042404 | 219350748 | 0.33967466 | 0.00953641 | 0.03119647 |
| sp\|P42773\|C | 26298545.6 | 36428650.3 | 0.35692927 | 0.00953641 | 0.03119647 |
| sp\|Q76M96\|( | 2234692.6 | 3078720.81 | 0.41212453 | 0.00953641 | 0.03119647 |
| sp\|Q9BY44\|E | 31944846.9 | 36823341.1 | 0.19965692 | 0.00953641 | 0.03119647 |
| sp\|O94925\|C | 7118575.58 | 5480245.03 | -0.3869024 | 0.00953641 | 0.03119647 |
| sp\|P53814\|S | 3875252.11 | 2496288.66 | -0.4885724 | 0.00953641 | 0.03119647 |
| sp\|P13591\|N | 9293780.26 | 6962220.34 | -0.6112711 | 0.00953641 | 0.03119647 |
| sp\|P63172\|D | 892109.131 | 692731.496 | -0.3863653 | 0.00953641 | 0.03119647 |
| sp\|Q15700\|D | 4511290.7 | 3666286.47 | -0.3228264 | 0.00953641 | 0.03119647 |
| sp\|P09960\|L | 972851894 | 782707781 | -0.3365332 | 0.00953641 | 0.03119647 |
| sp\|Q96B36\|A | 29753797.9 | 35353841 | 0.25142851 | 0.00953641 | 0.03119647 |
| sp\|P98095\|F | 30045946.5 | 44790987.3 | 0.45390757 | 0.00953641 | 0.03119647 |
| sp\|O15144\|A | 73335858.8 | 59260240.5 | -0.3809506 | 0.00953641 | 0.03119647 |
| sp\|P0C870\|J | 9976635.83 | 12829537.4 | 0.27967724 | 0.00953641 | 0.03119647 |
| sp\|P09603\|C | 236317.989 | 355784.185 | 0.75347385 | 0.00953641 | 0.03119647 |
| sp\|Q96S90\|L | 2114908.11 | 2848958.25 | 0.5706905 | 0.01025691 | 0.03299122 |
| sp\|Q8N9V3\|\ | 618347.972 | 1052335.92 | 0.51194247 | 0.01025691 | 0.03299122 |
| sp\|Q9UBF6\|F | 4348605.05 | 5975878.27 | 0.28079769 | 0.01025691 | 0.03299122 |
| sp\|P06132\|D | 58141403.9 | 75874000.4 | 0.31444265 | 0.01025691 | 0.03299122 |
| sp\|Q99575\|F | 1381337.44 | 1986786.27 | 0.36229011 | 0.01025691 | 0.03299122 |
| sp\|P59190\|R | 97704294.6 | 123053822 | 0.33321835 | 0.01025691 | 0.03299122 |
| sp\|Q9NP97\|I | 16920402.2 | 21143591.1 | 0.23074712 | 0.01025691 | 0.03299122 |
| sp\|P02671\|F | 3647470901 | 2978234725 | -0.4736045 | 0.01025691 | 0.03299122 |
| sp\|P49257\|L | 18745583.3 | 14639857.6 | -0.410115 | 0.01025691 | 0.03299122 |
| sp\|Q14677\|E | 11319298.9 | 13571802.5 | 0.23249467 | 0.01025691 | 0.03299122 |
| sp\|Q9Y4P3\|T | 4026144.38 | 6388830.88 | 0.49612376 | 0.01025691 | 0.03299122 |
| sp\|Q02040\|A | 2819815.02 | 3940383.23 | 0.37875622 | 0.01025691 | 0.03299122 |
| sp\|Q9BWM7\| | 36159177.4 | 27270366.1 | -0.4285553 | 0.01025691 | 0.03299122 |
| sp\|P29218\|II | 65852166.6 | 54961782.5 | -0.3567287 | 0.01025691 | 0.03299122 |

| | | | | | |
|---|---|---|---|---|---|
| sp\|P30520\|P | 20272068.7 | 17624426.2 | -0.3157946 | 0.01025691 | 0.03299122 |
| sp\|P51688\|S | 9867310.45 | 13783406.8 | 0.39541228 | 0.01025691 | 0.03299122 |
| sp\|Q6UXH1\| | 8712163.6 | 10535069.9 | 0.22096269 | 0.01025691 | 0.03299122 |
| sp\|O15533\|T | 5091789.01 | 3707275.25 | -0.55867 | 0.01025691 | 0.03299122 |
| sp\|Q9BV38\|\ | 995457.037 | 804312.151 | -0.4420675 | 0.01025691 | 0.03299122 |
| sp\|E9PAV3\|N | 42936641.3 | 50113340.3 | 0.2352429 | 0.01025691 | 0.03299122 |
| sp\|P25686\|D | 19374242.7 | 23187015.4 | 0.24025956 | 0.01025691 | 0.03299122 |
| sp\|Q9NTK5\|( | 99041078.4 | 71973892.4 | -0.6926079 | 0.01025691 | 0.03299122 |
| sp\|Q9H3Q1\| | 23327550.7 | 29958306.5 | 0.29148907 | 0.01025691 | 0.03299122 |
| sp\|P04271\|S | 128563164 | 160445570 | 0.36129829 | 0.01025691 | 0.03299122 |
| sp\|P22087\|F | 6727125.35 | 4929632.7 | -0.4315431 | 0.01025691 | 0.03299122 |
| sp\|P25815\|S | 2269160.04 | 6086269.64 | 0.67499155 | 0.01102353 | 0.03484975 |
| sp\|Q16222\|U | 67555168 | 90097435.8 | 0.3329944 | 0.01102353 | 0.03484975 |
| sp\|Q6ZSR9\|Y | 11954462.9 | 14712306.2 | 0.31016634 | 0.01102353 | 0.03484975 |
| sp\|Q99798\|A | 1194872225 | 855463266 | -0.7138405 | 0.01102353 | 0.03484975 |
| sp\|P00491\|P | 406761172 | 573186563 | 0.44115706 | 0.01102353 | 0.03484975 |
| sp\|Q12962\|T | 135276.19 | 240018.21 | 0.71417338 | 0.01102353 | 0.03484975 |
| sp\|Q96PU5\|I | 676901.57 | 483469.508 | -0.5757964 | 0.01102353 | 0.03484975 |
| sp\|P50225\|S | 70789104.3 | 52122248.1 | -0.5288315 | 0.01102353 | 0.03484975 |
| sp\|O95758\|F | 29757489 | 23308118.4 | -0.3831632 | 0.01102353 | 0.03484975 |
| sp\|O43583\|D | 6226435.85 | 8558902.32 | 0.38484471 | 0.01102353 | 0.03484975 |
| sp\|O94989\|A | 688439.583 | 1065178.47 | 0.46154652 | 0.01102353 | 0.03484975 |
| sp\|P10523\|A | 5046646.39 | 6623198.21 | 0.28385821 | 0.01102353 | 0.03484975 |
| sp\|Q9NW68\| | 12619070.6 | 14698819 | 0.22546596 | 0.01102353 | 0.03484975 |
| sp\|Q5HYW2\| | 1289377.71 | 1664582.32 | 0.42972452 | 0.01102353 | 0.03484975 |
| sp\|Q9Y305\|A | 13866303.6 | 10229945.3 | -0.4414303 | 0.01102353 | 0.03484975 |
| sp\|P14780\|M | 26880776.6 | 139869081 | 0.80341894 | 0.01102353 | 0.03484975 |
| sp\|O00391\|( | 18319370.5 | 22426557.6 | 0.25469792 | 0.01102353 | 0.03484975 |
| sp\|Q6P9B6\|I | 3753452.37 | 4993736.39 | 0.36516211 | 0.01102353 | 0.03484975 |
| sp\|P14543\|N | 508136605 | 675310516 | 0.34581337 | 0.01102353 | 0.03484975 |
| sp\|Q9Y657\|S | 1417014.55 | 1078240.54 | -0.4979962 | 0.01102353 | 0.03484975 |
| sp\|Q9BSU1\|I | 2094806.42 | 3135978.64 | 0.66420034 | 0.01102353 | 0.03484975 |
| sp\|Q9UHG3\| | 137596999 | 173491277 | 0.26053585 | 0.01102353 | 0.03484975 |
| sp\|P59998\|A | 117921169 | 88118143.5 | -0.4952237 | 0.01102353 | 0.03484975 |
| sp\|Q16740\|( | 26563022.5 | 32037754.7 | 0.28025 | 0.01102353 | 0.03484975 |
| sp\|Q6ZU35\|( | 412882.327 | 155064.845 | -2.480706 | 0.01102353 | 0.03484975 |
| sp\|Q9Y5S2\|M | 4639982.41 | 5822683.68 | 0.27730774 | 0.01102353 | 0.03484975 |
| sp\|Q9HD15\| | 3810225.9 | 5021440.64 | 0.34099017 | 0.01183857 | 0.03679618 |
| sp\|Q86TU7\|S | 18898680.8 | 21822654.5 | 0.18353798 | 0.01183857 | 0.03679618 |

| | | | | | |
|---|---|---|---|---|---|
| sp\|Q99715\|( | 27394142.6 | 21365194.5 | -0.3859329 | 0.01183857 | 0.03679618 |
| sp\|P49914\|M | 228436.402 | 549887.63 | 0.61921344 | 0.01183857 | 0.03679618 |
| sp\|Q10570\|( | 18451321.7 | 15890786.9 | -0.2061774 | 0.01183857 | 0.03679618 |
| sp\|P15848\|A | 6161997.64 | 8753254.9 | 0.42183928 | 0.01183857 | 0.03679618 |
| sp\|Q14517\|F | 411738.357 | 718404.415 | 0.82836063 | 0.01183857 | 0.03679618 |
| sp\|P17813\|E | 3812402.97 | 3210701.17 | -0.5580221 | 0.01183857 | 0.03679618 |
| sp\|Q8WW22 | 209685.247 | 556620.066 | 1.04182554 | 0.01183857 | 0.03679618 |
| sp\|Q86VX9\|N | 1413249.58 | 1895664.74 | 0.34189212 | 0.01183857 | 0.03679618 |
| sp\|Q9HCN8\| | 10103155.3 | 12846695 | 0.25214999 | 0.01183857 | 0.03679618 |
| sp\|P53990\|IS | 4599023.67 | 5353456.56 | 0.20527727 | 0.01183857 | 0.03679618 |
| sp\|O75023\|L | 9657797.07 | 11966840.4 | 0.2504974 | 0.01183857 | 0.03679618 |
| sp\|Q06265\|E | 3037970.58 | 2111544.66 | -0.55503 | 0.01183857 | 0.03679618 |
| sp\|Q9NWX5\| | 12238758.7 | 8775916.89 | -0.495292 | 0.01183857 | 0.03679618 |
| sp\|P30046\|D | 627317996 | 742049990 | 0.25927089 | 0.01183857 | 0.03679618 |
| sp\|P12883\|M | 73937760.7 | 87357555 | 0.26283564 | 0.01183857 | 0.03679618 |
| sp\|Q9Y2I6\|N | 720033.637 | 1154774.25 | 0.75541365 | 0.01183857 | 0.03679618 |
| sp\|P20336\|R | 66675836.2 | 79760947.2 | 0.25787284 | 0.01183857 | 0.03679618 |
| sp\|P08865\|R | 430588586 | 332546509 | -0.3447977 | 0.01183857 | 0.03679618 |
| sp\|O14791\|A | 3927665.78 | 6981225.06 | 0.58701258 | 0.01183857 | 0.03679618 |
| sp\|Q9BQG2\| | 12523513.3 | 17272040.3 | 0.45444118 | 0.01183857 | 0.03679618 |
| sp\|Q9GZM7\| | 8422917.71 | 10298114.7 | 0.29683929 | 0.01183857 | 0.03679618 |
| sp\|Q9NTM9\| | 22755850.3 | 29909999.8 | 0.40951649 | 0.01183857 | 0.03679618 |
| sp\|P17987\|T | 81911210.7 | 66185211.8 | -0.4221405 | 0.01183857 | 0.03679618 |
| sp\|Q8IZY2\|AI | 5528349.33 | 7560821.16 | 0.36238976 | 0.01183857 | 0.03679618 |
| sp\|O75323\|N | 37570167.1 | 26726876.3 | -0.5513708 | 0.01270445 | 0.03885828 |
| sp\|O75558\|S | 476783.224 | 634156.931 | 0.37421511 | 0.01270445 | 0.03885828 |
| sp\|Q15833\|S | 10289485.4 | 13952429.7 | 0.28243447 | 0.01270445 | 0.03885828 |
| sp\|P28340\|D | 66408699.1 | 47233328.3 | -0.8648605 | 0.01270445 | 0.03885828 |
| sp\|Q8NDH3\| | 26310808.3 | 41110706.3 | 0.51350837 | 0.01270445 | 0.03885828 |
| sp\|O43768\|E | 10648265.3 | 15490782.8 | 0.46082018 | 0.01270445 | 0.03885828 |
| sp\|O75947\|A | 62222912.3 | 76515466.4 | 0.30576515 | 0.01270445 | 0.03885828 |
| sp\|O43837\|I | 41451966.3 | 33581837.8 | -0.3295317 | 0.01270445 | 0.03885828 |
| sp\|Q9HC35\| | 10931192.8 | 8232560.26 | -0.4404196 | 0.01270445 | 0.03885828 |
| sp\|Q13432\|U | 7261619.97 | 5195573.95 | -0.6721347 | 0.01270445 | 0.03885828 |
| sp\|Q6FI13\|H | 243008205 | 144915200 | -0.8014461 | 0.01270445 | 0.03885828 |
| sp\|P35611\|A | 201826117 | 152627018 | -0.5136769 | 0.01270445 | 0.03885828 |
| sp\|Q9NRX2\|I | 11433180.5 | 8651099.03 | -0.6194382 | 0.01270445 | 0.03885828 |
| sp\|Q7KZF4\|S | 18075046.7 | 14466723.7 | -0.3420782 | 0.01270445 | 0.03885828 |
| sp\|Q16630\|( | 7821369.71 | 5909641.3 | -0.3968729 | 0.01270445 | 0.03885828 |

| | | | | | |
|---|---|---|---|---|---|
| sp\|Q969V3\|N | 5182077.77 | 4098067.65 | -0.3382306 | 0.01270445 | 0.03885828 |
| sp\|P62805\|H | 359354674 | 230678689 | -0.8351801 | 0.01270445 | 0.03885828 |
| sp\|P07357\|C | 102789137 | 71788803.3 | -0.7197724 | 0.01270445 | 0.03885828 |
| sp\|Q8WUF5\| | 1923379.37 | 1459533.87 | -0.4603488 | 0.01270445 | 0.03885828 |
| sp\|Q8IWZ3\|A | 2556731.76 | 1571330.9 | -0.7372701 | 0.01270445 | 0.03885828 |
| sp\|Q12974\|T | 5724778.94 | 8645848.7 | 0.70737528 | 0.01270445 | 0.03885828 |
| sp\|Q5VWN6\| | 1609689.4 | 2415022.19 | 0.48116032 | 0.01270445 | 0.03885828 |
| sp\|Q9P0J0\|N | 8639179.01 | 6510236.47 | -0.3823787 | 0.01270445 | 0.03885828 |
| sp\|P09669\|C | 14639858.3 | 10608717.2 | -0.4565793 | 0.01270445 | 0.03885828 |
| sp\|O43251\|F | 15680221.5 | 12388046.3 | -0.4978259 | 0.01270445 | 0.03885828 |
| sp\|A4D1P6\|V | 8441336.11 | 6901027.31 | -0.3550195 | 0.0136237 | 0.0410164 |
| sp\|P51884\|L | 4637096657 | 7111881168 | 0.41299058 | 0.0136237 | 0.0410164 |
| sp\|Q9Y388\|F | 2487369.04 | 3537215.65 | 0.35928323 | 0.0136237 | 0.0410164 |
| sp\|Q96H79\|Z | 44870160.1 | 29164354.4 | -0.7421056 | 0.0136237 | 0.0410164 |
| sp\|Q8NHH1\| | 2428565.47 | 3514323.63 | 0.46471983 | 0.0136237 | 0.0410164 |
| sp\|P28906\|C | 42626246.8 | 32050049 | -0.443297 | 0.0136237 | 0.0410164 |
| sp\|Q04721\|N | 10112209.2 | 12474438.4 | 0.24853222 | 0.0136237 | 0.0410164 |
| sp\|Q32MK0\|I | 7565197.09 | 6409622.63 | -0.2743969 | 0.0136237 | 0.0410164 |
| sp\|Q15006\|E | 1754594.56 | 1410210.63 | -0.2646152 | 0.0136237 | 0.0410164 |
| sp\|Q9BV44\|T | 2333612.68 | 1984955.84 | -0.2591888 | 0.0136237 | 0.0410164 |
| sp\|Q86T03\|F | 4744430.45 | 6921231.9 | 0.3672924 | 0.0136237 | 0.0410164 |
| sp\|Q13200\|F | 79980490.7 | 92393709.6 | 0.17324509 | 0.0136237 | 0.0410164 |
| sp\|O15127\|S | 6181192.96 | 7567524.13 | 0.28404143 | 0.0136237 | 0.0410164 |
| sp\|Q9Y3E7\|C | 10807666.4 | 13634597.4 | 0.34754038 | 0.0136237 | 0.0410164 |
| sp\|P19105\|M | 64917493.6 | 80426649.7 | 0.28801135 | 0.0136237 | 0.0410164 |
| sp\|P46109\|C | 69222690.9 | 59044215.8 | -0.2472457 | 0.0136237 | 0.0410164 |
| sp\|Q9BUN8\| | 6532362.31 | 4729752.13 | -0.5858668 | 0.0136237 | 0.0410164 |
| sp\|P84090\|E | 2430616.94 | 1561369.81 | -0.7310769 | 0.0136237 | 0.0410164 |
| sp\|Q9NXV6\|C | 409727.984 | 274301.927 | -0.6920112 | 0.0136237 | 0.0410164 |
| sp\|Q9H425\|C | 42599257.1 | 55864930.9 | 0.28995215 | 0.0136237 | 0.0410164 |
| sp\|O95210\|S | 34172382.2 | 39929222.2 | 0.21372932 | 0.0136237 | 0.0410164 |
| sp\|P60660\|M | 956276427 | 1062068930 | 0.14713362 | 0.0136237 | 0.0410164 |
| sp\|Q9NVI7\|A | 1799635.18 | 2480162.45 | 0.6873251 | 0.0136237 | 0.0410164 |
| sp\|Q96DE0\|N | 21807994.4 | 26624276.8 | 0.32234619 | 0.0136237 | 0.0410164 |
| sp\|P36542\|A | 157549144 | 201514583 | 0.36470258 | 0.0136237 | 0.0410164 |
| sp\|P34913\|H | 52224990.2 | 40393126.1 | -0.4075312 | 0.01459887 | 0.04351551 |
| sp\|Q9H3Y8\|F | 189754.949 | 273131.152 | 0.61201882 | 0.01459887 | 0.04351551 |
| sp\|Q5T9S5\|C | 2871632.15 | 4075864.47 | 0.26854933 | 0.01459887 | 0.04351551 |
| sp\|Q12857\|N | 960687.881 | 1496517.7 | 0.5313072 | 0.01459887 | 0.04351551 |

| | | | | | |
|---|---|---|---|---|---|
| sp\|P27918\|P | 6724603.95 | 8784807.16 | 0.42996794 | 0.01459887 | 0.04351551 |
| sp\|Q8TET4\|G | 11771440.6 | 15038963.7 | 0.35057675 | 0.01459887 | 0.04351551 |
| sp\|P53985\|M | 216762.473 | 213689.597 | -1.4218815 | 0.01459887 | 0.04351551 |
| sp\|Q6UUV9\| | 4671623.42 | 5753182.06 | 0.35233716 | 0.01459887 | 0.04351551 |
| sp\|P22626\|R | 455122525 | 366295605 | -0.3452038 | 0.01459887 | 0.04351551 |
| sp\|Q9Y6K1\|D | 219233.903 | 395385.577 | -0.0269064 | 0.01459887 | 0.04351551 |
| sp\|Q04446\|C | 323608779 | 475739894 | 0.34255222 | 0.01459887 | 0.04351551 |
| sp\|Q96T37\|F | 604041.158 | 380136.106 | -0.7275054 | 0.01459887 | 0.04351551 |
| sp\|Q92954\|F | 11037194 | 14866326.5 | 0.3652644 | 0.01459887 | 0.04351551 |
| sp\|Q96MP5\|. | 3492029.28 | 4234640.16 | 0.24758662 | 0.01459887 | 0.04351551 |
| sp\|Q9NY12\|C | 5815765 | 4174409.88 | -0.5307184 | 0.01459887 | 0.04351551 |
| sp\|P07998\|R | 36567870.5 | 51406455.5 | 0.28417382 | 0.01459887 | 0.04351551 |
| sp\|Q9HBK9\|A | 5978073.31 | 13190551.8 | 0.82146514 | 0.01563261 | 0.04574444 |
| sp\|A0A0B4J2 | 1874361.32 | 1411010.78 | -0.437539 | 0.01563261 | 0.04574444 |
| sp\|Q13310\|F | 77639164 | 61029127.2 | -0.4849404 | 0.01563261 | 0.04574444 |
| sp\|Q96HE7\|E | 201981689 | 165108802 | -0.3354488 | 0.01563261 | 0.04574444 |
| sp\|Q6VEQ5\|V | 3079183.18 | 3816687.66 | 0.29119533 | 0.01563261 | 0.04574444 |
| sp\|Q13838\|D | 49432312 | 38366988.4 | -0.3405949 | 0.01563261 | 0.04574444 |
| sp\|P07996\|T | 111578258 | 182545196 | 0.44465456 | 0.01563261 | 0.04574444 |
| sp\|Q9UKV3\|A | 16216721.5 | 13007318.6 | -0.3143597 | 0.01563261 | 0.04574444 |
| sp\|O75122\|C | 296762.926 | 227547.292 | -0.0593061 | 0.01563261 | 0.04574444 |
| sp\|P19397\|C | 1007213.77 | 1906060.98 | 0.6433899 | 0.01563261 | 0.04574444 |
| sp\|Q9H496\|I | 1000276.12 | 1451759 | 0.34058541 | 0.01563261 | 0.04574444 |
| sp\|P19087\|G | 89573462.6 | 72790033 | -0.3450727 | 0.01563261 | 0.04574444 |
| sp\|P35579\|M | 270290244 | 326980548 | 0.25418219 | 0.01563261 | 0.04574444 |
| sp\|P48509\|C | 27952564.9 | 34694358.6 | 0.28656668 | 0.01563261 | 0.04574444 |
| sp\|Q9BQ61\|T | 8472559.98 | 10461974.6 | 0.26338559 | 0.01563261 | 0.04574444 |
| sp\|P60228\|E | 3949540.43 | 2640244.6 | -0.56284 | 0.01563261 | 0.04574444 |
| sp\|P24593\|IE | 11334905.2 | 14053033.3 | 0.26649166 | 0.01563261 | 0.04574444 |
| sp\|Q9Y3D8\|K | 461366.606 | 2235022.72 | 0.73556523 | 0.01563261 | 0.04574444 |
| sp\|Q16695\|H | 149551828 | 100005304 | -0.7715164 | 0.01563261 | 0.04574444 |
| sp\|O43678\|N | 11942992.6 | 14690874 | 0.17944821 | 0.01563261 | 0.04574444 |
| sp\|Q16773\|K | 4633061.81 | 5807451.01 | 0.30251769 | 0.01563261 | 0.04574444 |
| sp\|Q8IWC1\|I | 3259065.98 | 4156104.98 | 0.58402575 | 0.01563261 | 0.04574444 |
| sp\|Q9UBS4\|I | 12039379 | 13880807.2 | 0.18169509 | 0.01563261 | 0.04574444 |
| sp\|Q15796\|S | 1398292.62 | 941125.433 | -0.6548917 | 0.01563261 | 0.04574444 |
| sp\|Q9Y512\|S | 1656769.21 | 1293236.48 | -0.3410764 | 0.01563261 | 0.04574444 |
| sp\|P07108\|A | 458394975 | 647705004 | 0.36018475 | 0.01563261 | 0.04574444 |
| sp\|Q15811\|IT | 44370212.2 | 51177132.8 | 0.19021727 | 0.01563261 | 0.04574444 |

| | | | | | |
|---|---|---|---|---|---|
| sp\|Q9NQ50\| | 3108127.54 | 2262552.77 | -0.5656408 | 0.01563261 | 0.04574444 |
| sp\|Q17R31\|T | 8694321.93 | 10508222.5 | 0.24262822 | 0.01563261 | 0.04574444 |
| sp\|O95415\|E | 522775.354 | 743559.158 | -0.1414731 | 0.01563261 | 0.04574444 |
| sp\|O75167\|F | 1091720.35 | 1444780.11 | 0.52751803 | 0.01672769 | 0.04835916 |
| sp\|P35609\|A | 153196182 | 110667346 | -0.4864198 | 0.01672769 | 0.04835916 |
| sp\|Q9H3Z7\|A | 10918250.8 | 8178948.02 | -0.7145868 | 0.01672769 | 0.04835916 |
| sp\|P13489\|R | 987938146 | 868906410 | -0.180491 | 0.01672769 | 0.04835916 |
| sp\|Q96NT1\|N | 33597453.1 | 38831466.3 | 0.22642452 | 0.01672769 | 0.04835916 |
| sp\|Q96F24\|N | 3310300.96 | 4749237.38 | 0.66330928 | 0.01672769 | 0.04835916 |
| sp\|P21912\|S | 162740451 | 190238511 | 0.2184263 | 0.01672769 | 0.04835916 |
| sp\|P62495\|E | 1595925.94 | 2353391.29 | 0.49044057 | 0.01672769 | 0.04835916 |
| sp\|P42025\|A | 111978750 | 94847812.5 | -0.2837709 | 0.01672769 | 0.04835916 |
| sp\|Q96RU3\|I | 21554232.3 | 25559246.3 | 0.2363195 | 0.01672769 | 0.04835916 |
| sp\|Q9HDC9\| | 140845623 | 111992748 | -0.4772618 | 0.01672769 | 0.04835916 |
| sp\|Q8ND56\| | 1557027.33 | 1080062.49 | -0.4100383 | 0.01672769 | 0.04835916 |
| sp\|Q96EL2\|F | 7885935.25 | 16075418.5 | 1.19007773 | 0.01672769 | 0.04835916 |
| sp\|Q14789\|C | 13437852.6 | 15944801.1 | 0.22545335 | 0.01672769 | 0.04835916 |
| sp\|O43169\|C | 5102972.24 | 6361683.52 | 0.27160857 | 0.01672769 | 0.04835916 |
| sp\|P21266\|G | 306258668 | 229731726 | -0.7472602 | 0.01672769 | 0.04835916 |
| sp\|P61586\|R | 181987397 | 221097382 | 0.25015989 | 0.01672769 | 0.04835916 |
| sp\|P26641\|E | 125666672 | 88806453.2 | -0.5925052 | 0.01672769 | 0.04835916 |
| sp\|P33316\|D | 34114130.1 | 29097386 | -0.2306026 | 0.01672769 | 0.04835916 |
| sp\|P32004\|L | 2273897.25 | 3175371.03 | 0.48322111 | 0.01672769 | 0.04835916 |
| sp\|P49006\|M | 4438573.42 | 2819992.44 | -0.6855198 | 0.01788691 | 0.05127796 |
| sp\|O43715\|T | 706983.826 | 949187.372 | 0.67087112 | 0.01788691 | 0.05127796 |
| sp\|Q9P2R3\|A | 2573472.55 | 4395490.92 | 0.43678531 | 0.01788691 | 0.05127796 |
| sp\|O14776\|T | 1321006.96 | 1002323.19 | -0.5874945 | 0.01788691 | 0.05127796 |
| sp\|Q86TE4\|L | 1946254.36 | 1505617.01 | -0.6490106 | 0.01788691 | 0.05127796 |
| sp\|Q4G0P3\|I | 4845643.95 | 3203967.06 | -0.5483094 | 0.01788691 | 0.05127796 |
| sp\|P18669\|P | 1717002548 | 2006486600 | 0.1741613 | 0.01788691 | 0.05127796 |
| sp\|O43292\|C | 790977.451 | 509057.937 | -0.6739969 | 0.01788691 | 0.05127796 |
| sp\|Q13283\|C | 26907359.7 | 23108778.1 | -0.2446159 | 0.01788691 | 0.05127796 |
| sp\|Q00169\|F | 14484946.4 | 11039070.9 | -0.4832675 | 0.01788691 | 0.05127796 |
| sp\|Q9P121\|N | 2445183.49 | 3220757.77 | 0.40021774 | 0.01788691 | 0.05127796 |
| sp\|O15455\|T | 1236537.81 | 1745788.61 | 0.42816393 | 0.01788691 | 0.05127796 |
| sp\|Q92526\|T | 17865374.8 | 14303394 | -0.4138872 | 0.01788691 | 0.05127796 |
| sp\|Q92692\|N | 12635367.2 | 15922370 | 0.37716823 | 0.01788691 | 0.05127796 |
| sp\|Q9NV35\|I | 3800646.65 | 5985566.92 | 0.51107562 | 0.01911315 | 0.05398704 |
| sp\|Q9UL25\|F | 28494540.3 | 34027539.4 | 0.2600162 | 0.01911315 | 0.05398704 |

| | | | | | |
|---|---|---|---|---|---|
| sp\|P45954\|A | 79608205.2 | 93380247.9 | 0.24250716 | 0.01911315 | 0.05398704 |
| sp\|Q9UL18\|/ | 33382513.4 | 27524015.1 | -0.3286941 | 0.01911315 | 0.05398704 |
| sp\|P51911\|C | 191199045 | 104612682 | -0.5454615 | 0.01911315 | 0.05398704 |
| sp\|P62314\|S | 26796798.2 | 19949098.1 | -0.5150604 | 0.01911315 | 0.05398704 |
| sp\|P02766\|T | 894793258 | 1241526341 | 0.40336747 | 0.01911315 | 0.05398704 |
| sp\|P25963\|II | 1452867 | 1770260.09 | -0.1955975 | 0.01911315 | 0.05398704 |
| sp\|P41439\|F | 2567696.42 | 11134202 | 0.70697935 | 0.01911315 | 0.05398704 |
| sp\|Q0VD83\|/ | 3349727.79 | 4588795.41 | 0.44976055 | 0.01911315 | 0.05398704 |
| sp\|O14639\|/ | 88638835.8 | 101598411 | 0.16947685 | 0.01911315 | 0.05398704 |
| sp\|O14558\|I | 1682370410 | 2077250605 | 0.36299751 | 0.01911315 | 0.05398704 |
| sp\|Q96LD4\|1 | 12476997.4 | 17008540.9 | 0.40377 | 0.01911315 | 0.05398704 |
| sp\|P35749\|N | 556212127 | 425313460 | -0.3131758 | 0.01911315 | 0.05398704 |
| sp\|P26196\|D | 8479654.54 | 7062889.5 | -0.3296946 | 0.01911315 | 0.05398704 |
| sp\|P09493\|T | 1997963305 | 1435483340 | -0.3358752 | 0.01911315 | 0.05398704 |
| sp\|P80723\|B | 22446429 | 33786972.8 | 0.44746232 | 0.01911315 | 0.05398704 |
| sp\|P22352\|G | 432202520 | 619850310 | 0.38637213 | 0.01911315 | 0.05398704 |
| sp\|Q9NZ09\|U | 1191025.84 | 1485469.27 | 0.27266249 | 0.01911315 | 0.05398704 |
| sp\|P60981\|D | 356007268 | 262430202 | -0.3872547 | 0.01911315 | 0.05398704 |
| sp\|P21589\|5 | 49670146.4 | 42164794.4 | -0.2839058 | 0.01911315 | 0.05398704 |
| sp\|Q8TBF2\|F | 3517681.52 | 4421575.12 | 0.39111063 | 0.01911315 | 0.05398704 |
| sp\|Q5TEC6\|I | 105700106 | 73847277.9 | -0.618597 | 0.01911315 | 0.05398704 |
| sp\|P07585\|P | 274205554 | 424916006 | 0.48934868 | 0.01911315 | 0.05398704 |
| sp\|O43772\|N | 24575665.9 | 21611116.4 | -0.1913741 | 0.01911315 | 0.05398704 |
| sp\|P05455\|L | 169081136 | 142024415 | -0.3366885 | 0.02040941 | 0.05661548 |
| sp\|P52815\|R | 14427959.9 | 18374085.8 | 0.33787373 | 0.02040941 | 0.05661548 |
| sp\|Q16181\|S | 184633814 | 134335392 | -0.4889289 | 0.02040941 | 0.05661548 |
| sp\|Q5SRE7\|F | 17968211.5 | 15610005.1 | -0.1987177 | 0.02040941 | 0.05661548 |
| sp\|P34096\|R | 15024924.8 | 17958236.4 | 0.21671958 | 0.02040941 | 0.05661548 |
| sp\|O14744\|/ | 148020905 | 114945416 | -0.5438261 | 0.02040941 | 0.05661548 |
| sp\|P49321\|N | 19843265.7 | 16887735.3 | -0.2623781 | 0.02040941 | 0.05661548 |
| sp\|Q9P258\|F | 4197629.09 | 3693663.68 | -0.2592706 | 0.02040941 | 0.05661548 |
| sp\|Q9HCE5\| | 1685321.15 | 2540999.69 | 0.63607607 | 0.02040941 | 0.05661548 |
| sp\|Q9UPT5\|E | 798711.403 | 1420748.82 | 0.56847194 | 0.02040941 | 0.05661548 |
| sp\|Q92599\|S | 198042892 | 155521629 | -0.3572829 | 0.02040941 | 0.05661548 |
| sp\|Q2LD37\|K | 675134.005 | 1039781.17 | 0.56162205 | 0.02040941 | 0.05661548 |
| sp\|O43504\|L | 3197014.17 | 4853627.21 | 0.50259725 | 0.02040941 | 0.05661548 |
| sp\|Q9Y4P8\|V | 74056528.2 | 104349096 | 0.75282971 | 0.02040941 | 0.05661548 |
| sp\|Q13045\|F | 8071754.08 | 6175053.08 | -0.4359813 | 0.02040941 | 0.05661548 |
| sp\|Q86WI1\|F | 212808.358 | 4033411.16 | -1.8514349 | 0.02040941 | 0.05661548 |

| | | | | | |
|---|---|---|---|---|---|
| sp\|Q7Z6K5\|A | 38732625.5 | 34595025.8 | -0.1903005 | 0.02040941 | 0.05661548 |
| sp\|Q9HC77\| | 5835610.62 | 7978809.78 | 0.45281554 | 0.02040941 | 0.05661548 |
| sp\|Q13084\|F | 4123479.52 | 3279561.26 | -0.4640683 | 0.02040941 | 0.05661548 |
| sp\|O00743\|F | 19002273.2 | 21856414 | 0.20064812 | 0.02040941 | 0.05661548 |
| sp\|Q8TDP1\|F | 5528725.2 | 4429869.8 | -0.2578725 | 0.02040941 | 0.05661548 |
| sp\|Q86YS6\|F | 53355190.6 | 66417168.3 | 0.33674862 | 0.02040941 | 0.05661548 |
| sp\|O43155\|F | 5429464.3 | 6628847.63 | 0.30738085 | 0.02040941 | 0.05661548 |
| sp\|O00182\|L | 6746613.31 | 4388380.26 | -0.6194461 | 0.02040941 | 0.05661548 |
| sp\|Q9BY49\|F | 2913360.15 | 3550716.51 | 0.31990971 | 0.02040941 | 0.05661548 |
| sp\|Q9BQE5\|A | 2378881.39 | 2946835.91 | 0.32078421 | 0.02040941 | 0.05661548 |
| sp\|Q9Y696\|C | 60060883.2 | 45755950.1 | -0.5265475 | 0.02040941 | 0.05661548 |
| sp\|Q8NCC3\| | 28371503.3 | 38192979.3 | 0.32921193 | 0.02040941 | 0.05661548 |
| sp\|Q9BZF9\|U | 19370874.2 | 15296223.3 | -0.2619679 | 0.02040941 | 0.05661548 |
| sp\|Q8WXU2\| | 1674815.42 | 1252616.31 | -0.4600737 | 0.02040941 | 0.05661548 |
| sp\|Q15813\|T | 798578.922 | 467963.633 | -0.6974502 | 0.02040941 | 0.05661548 |
| sp\|Q5BKZ1\|Z | 1398774.65 | 933118.196 | -0.6586187 | 0.02177873 | 0.05941793 |
| sp\|Q96E39\|F | 55575380.5 | 46697958 | -0.2823147 | 0.02177873 | 0.05941793 |
| sp\|P04406\|G | 3790219232 | 6272552450 | 0.6138876 | 0.02177873 | 0.05941793 |
| sp\|Q9HD40\| | 2318439.68 | 1668905.53 | -0.5292806 | 0.02177873 | 0.05941793 |
| sp\|P07451\|C | 836345248 | 470653666 | -0.6303185 | 0.02177873 | 0.05941793 |
| sp\|Q8NCN5\| | 77243524.9 | 66575066.3 | -0.2414726 | 0.02177873 | 0.05941793 |
| sp\|P78406\|R | 7875084.56 | 6531586.73 | -0.3654916 | 0.02177873 | 0.05941793 |
| sp\|Q6P6C2\|A | 507596.682 | 394394.341 | -0.3642934 | 0.02177873 | 0.05941793 |
| sp\|P62714\|P | 138632711 | 161971749 | 0.1759352 | 0.02177873 | 0.05941793 |
| sp\|P50579\|M | 10398964.1 | 8686203.89 | -0.2387321 | 0.02177873 | 0.05941793 |
| sp\|Q96BW5\| | 103562768 | 142897552 | 0.24127436 | 0.02177873 | 0.05941793 |
| sp\|Q0ZGT2\|N | 33144061.9 | 41233308.8 | 0.26347351 | 0.02177873 | 0.05941793 |
| sp\|Q92686\|N | 694042.396 | 677580.703 | -0.8736064 | 0.02177873 | 0.05941793 |
| sp\|Q14254\|F | 183933239 | 222340103 | 0.26132339 | 0.02177873 | 0.05941793 |
| sp\|Q8IYB5\|S | 3698610.73 | 5170031.51 | 0.23931866 | 0.02177873 | 0.05941793 |
| sp\|P50395\|G | 387769077 | 302294922 | -0.584295 | 0.02177873 | 0.05941793 |
| sp\|P62633\|C | 10749079.2 | 14630099.4 | 0.25495775 | 0.02177873 | 0.05941793 |
| sp\|Q96FW1\| | 15485599.7 | 11121989 | -0.5700893 | 0.02177873 | 0.05941793 |
| sp\|Q6SPF0\|S | 1557064.77 | 932557.185 | -0.8402131 | 0.02177873 | 0.05941793 |
| sp\|Q96GC5\| | 323572.777 | 213668.265 | -1.1935789 | 0.02177873 | 0.05941793 |
| sp\|Q99816\|T | 2492776.35 | 3167805.76 | 0.22148412 | 0.02177873 | 0.05941793 |
| sp\|O00748\|E | 11289963.7 | 16605117.1 | 0.41729255 | 0.02177873 | 0.05941793 |
| sp\|P61952\|G | 3719871.69 | 4391066.9 | 0.27721423 | 0.02177873 | 0.05941793 |
| sp\|Q14166\|T | 31836203.3 | 24457579.5 | -0.4967636 | 0.02177873 | 0.05941793 |

| | | | | | |
|---|---|---|---|---|---|
| sp\|P98155\|V | 5561107.42 | 7258925.44 | 0.34807507 | 0.02177873 | 0.05941793 |
| sp\|Q99417\|N | 4241313.04 | 5427212.44 | 0.40727234 | 0.02177873 | 0.05941793 |
| sp\|Q05209\|F | 6507547.62 | 7832909.91 | 0.2245707 | 0.02177873 | 0.05941793 |
| sp\|Q03426\|K | 4962682.35 | 5818847.85 | 0.1843401 | 0.02177873 | 0.05941793 |
| sp\|Q6NYC1\|. | 850883.785 | 1438479.63 | 0.72479457 | 0.02177873 | 0.05941793 |
| sp\|P46020\|K | 4734859.01 | 6191852.46 | 0.29217524 | 0.02322421 | 0.06233389 |
| sp\|Q6IA86\|E | 10522725.8 | 12076859.2 | 0.17519204 | 0.02322421 | 0.06233389 |
| sp\|O15145\|A | 78772624.7 | 64860295 | -0.356546 | 0.02322421 | 0.06233389 |
| sp\|Q13188\|S | 4112692.08 | 5725706.77 | 0.46838888 | 0.02322421 | 0.06233389 |
| sp\|P08236\|B | 68967486 | 93987130.3 | 0.33066428 | 0.02322421 | 0.06233389 |
| sp\|P04233\|H | 2938735.73 | 4226950.15 | 0.40629098 | 0.02322421 | 0.06233389 |
| sp\|P79483\|D | 7455091.02 | 10082060 | 0.35913079 | 0.02322421 | 0.06233389 |
| sp\|Q9UN86\| | 9286259.33 | 8016974.91 | -0.2132021 | 0.02322421 | 0.06233389 |
| sp\|Q13637\|F | 3860527.65 | 7071027.1 | 0.70857311 | 0.02322421 | 0.06233389 |
| sp\|P01911\|D | 23610473.3 | 30629104.9 | 0.26393334 | 0.02322421 | 0.06233389 |
| sp\|P00915\|C | 1.2164E+10 | 1.9391E+10 | 0.54389283 | 0.02322421 | 0.06233389 |
| sp\|Q96MW1\| | 657010.766 | 1072498.76 | 0.52680132 | 0.02322421 | 0.06233389 |
| sp\|Q12846\|S | 9781111.77 | 11440388 | 0.1836244 | 0.02322421 | 0.06233389 |
| sp\|Q12874\|S | 3275176.18 | 2688543.6 | -0.3271236 | 0.02322421 | 0.06233389 |
| sp\|O95816\|E | 3164228.36 | 2736563.79 | -0.2324272 | 0.02322421 | 0.06233389 |
| sp\|P28676\|G | 7271921.77 | 16841746.6 | 0.55709657 | 0.02322421 | 0.06233389 |
| sp\|P61966\|A | 3069677.94 | 2518675.1 | -0.3587306 | 0.02322421 | 0.06233389 |
| sp\|P55001\|M | 36991880.1 | 30234735.5 | -0.4435664 | 0.02322421 | 0.06233389 |
| sp\|Q8TBC4\|U | 26989073.6 | 22718260 | -0.3378385 | 0.02322421 | 0.06233389 |
| sp\|Q63HM2\| | 2563388.91 | 3382341.71 | 0.08879704 | 0.02322421 | 0.06233389 |
| sp\|P27708\|P | 19279334 | 14765395.3 | -0.3629715 | 0.02322421 | 0.06233389 |
| sp\|P30047\|G | 21162339.9 | 21840225.6 | 0.2016087 | 0.02322421 | 0.06233389 |
| sp\|Q14764\|M | 158981299 | 127825041 | -0.3467467 | 0.02322421 | 0.06233389 |
| sp\|Q9HCB6\| | 45951411.1 | 40852561 | -0.2911097 | 0.02322421 | 0.06233389 |
| sp\|Q9Y3I1\|FI | 3162239.76 | 3888812.88 | 0.28628209 | 0.02322421 | 0.06233389 |
| sp\|Q9NVS2\|I | 5572254.61 | 4345936.08 | -0.4099829 | 0.02322421 | 0.06233389 |
| sp\|P06276\|C | 29571292.3 | 24165245.7 | -0.4204175 | 0.02322421 | 0.06233389 |
| sp\|P62875\|R | 10244084.3 | 7183651.41 | -0.7779842 | 0.02322421 | 0.06233389 |
| sp\|Q15477\|S | 439378.606 | 580967.627 | 0.30477353 | 0.02322421 | 0.06233389 |
| sp\|Q16099\|G | 1540795.53 | 2133228.53 | 0.39550387 | 0.02474912 | 0.06511568 |
| sp\|Q15661\|T | 421032524 | 697654688 | 0.74461461 | 0.02474912 | 0.06511568 |
| sp\|Q6UXP7\|I | 3546654.82 | 7740733.4 | 1.3450828 | 0.02474912 | 0.06511568 |
| sp\|Q9NWV4\| | 127908319 | 168203352 | 0.33168039 | 0.02474912 | 0.06511568 |
| sp\|Q9Y5B9\|S | 406619.602 | 324455.682 | -1.141028 | 0.02474912 | 0.06511568 |

| | | | | | |
|---|---|---|---|---|---|
| sp\|Q9NW38\| | 5381781.42 | 7469608.74 | 0.45248832 | 0.02474912 | 0.06511568 |
| sp\|O15075\|D | 8334830.53 | 12493899.2 | 0.3071229 | 0.02474912 | 0.06511568 |
| sp\|Q8IV08\|P | 49536193.5 | 73335781.1 | 0.3286156 | 0.02474912 | 0.06511568 |
| sp\|P50416\|C | 9106623.92 | 6629584.92 | -0.5624444 | 0.02474912 | 0.06511568 |
| sp\|P32320\|C | 10437325.5 | 19792607.4 | 0.44631179 | 0.02474912 | 0.06511568 |
| sp\|Q9UFG5\| | 4249426.77 | 5452957.78 | 0.43678165 | 0.02474912 | 0.06511568 |
| sp\|P10909\|C | 251675116 | 365241996 | 0.41034589 | 0.02474912 | 0.06511568 |
| sp\|P24534\|E | 92757667.8 | 107192130 | 0.15994859 | 0.02474912 | 0.06511568 |
| sp\|Q8WU39\| | 30057347.6 | 6858123.83 | -0.8478691 | 0.02474912 | 0.06511568 |
| sp\|P24855\|D | 18200167.5 | 28492230 | 0.64885369 | 0.02474912 | 0.06511568 |
| sp\|Q8N2G6\| | 180648.444 | 128321.753 | -0.7962507 | 0.02474912 | 0.06511568 |
| sp\|Q9GZZ9\|U | 2583000.02 | 3731699.28 | 0.42846826 | 0.02474912 | 0.06511568 |
| sp\|Q9NZT1\|C | 9658969.14 | 14092496.7 | 0.54299682 | 0.02474912 | 0.06511568 |
| sp\|P61916\|N | 105569640 | 144210075 | 0.34891666 | 0.02474912 | 0.06511568 |
| sp\|Q9Y3E1\|H | 31597588.6 | 24205765.7 | -0.4869282 | 0.02474912 | 0.06511568 |
| sp\|Q96R05\|F | 7301435.61 | 10652051.1 | 0.36178213 | 0.02474912 | 0.06511568 |
| sp\|P15259\|P | 841936211 | 968167640 | 0.19013233 | 0.02474912 | 0.06511568 |
| sp\|Q96S66\|C | 14928024 | 17265457.6 | 0.18115964 | 0.02474912 | 0.06511568 |
| sp\|Q8N573\|C | 10341801.9 | 11908590.9 | 0.16084289 | 0.02474912 | 0.06511568 |
| sp\|Q969X5\|E | 2933666.28 | 2330015.62 | -0.4287749 | 0.02474912 | 0.06511568 |
| sp\|Q6NYC8\| | 11188044.5 | 13976595.3 | 0.2465887 | 0.02474912 | 0.06511568 |
| sp\|P42785\|P | 31171676.9 | 40155527.7 | 0.27036857 | 0.02474912 | 0.06511568 |
| sp\|P62191\|P | 34801839.5 | 42176915.1 | 0.23052598 | 0.02474912 | 0.06511568 |
| sp\|Q9P2N2\| | 3166907.49 | 2689493.22 | -0.3598895 | 0.02474912 | 0.06511568 |
| sp\|Q9HB90\| | 13255537.6 | 10999222.1 | -0.3302168 | 0.02474912 | 0.06511568 |
| sp\|P17050\|N | 27733476.2 | 39422439.2 | 0.40541857 | 0.02474912 | 0.06511568 |
| sp\|P16104\|H | 157394036 | 94603917.2 | -0.73154 | 0.02474912 | 0.06511568 |
| sp\|P57737\|C | 74472730.5 | 57033277.1 | -0.4177066 | 0.02474912 | 0.06511568 |
| sp\|Q16478\|C | 1540795.53 | 2133228.53 | 0.39550387 | 0.02474912 | 0.06511568 |
| sp\|P07360\|C | 52111008.5 | 41604589.7 | -0.5599228 | 0.02474912 | 0.06511568 |
| sp\|P13473\|L | 9480963.24 | 12241699.9 | 0.3358743 | 0.02474912 | 0.06511568 |
| sp\|Q9NP79\|V | 3809335.72 | 4788986.13 | 0.33402381 | 0.02635667 | 0.06818633 |
| sp\|Q9H444\|C | 71640063.2 | 81397139.1 | 0.2369266 | 0.02635667 | 0.06818633 |
| sp\|Q86SX6\|C | 4695512.96 | 6353645.92 | 0.24889561 | 0.02635667 | 0.06818633 |
| sp\|P09497\|C | 147276501 | 169014022 | 0.16963842 | 0.02635667 | 0.06818633 |
| sp\|Q3LXA3\|T | 62695597.4 | 54723181.5 | -0.3280764 | 0.02635667 | 0.06818633 |
| sp\|P42126\|E | 273448801 | 192673388 | -0.4514385 | 0.02635667 | 0.06818633 |
| sp\|P40121\|C | 139022843 | 107270362 | -0.4760346 | 0.02635667 | 0.06818633 |
| sp\|P06454\|P | 5031905.5 | 3583903.67 | -0.6040623 | 0.02635667 | 0.06818633 |

| | | | | | |
|---|---|---|---|---|---|
| sp\|P48739\|P | 10508605.1 | 7607234.6 | -0.6480921 | 0.02635667 | 0.06818633 |
| sp\|Q15370\|E | 22888323.9 | 28479455.9 | 0.21839112 | 0.02635667 | 0.06818633 |
| sp\|Q9UK76\|J | 5294364.01 | 7585108.26 | 0.46953293 | 0.02635667 | 0.06818633 |
| sp\|Q96DT5\|D | 6956434.75 | 13803566.6 | 1.13412421 | 0.02635667 | 0.06818633 |
| sp\|P62316\|S | 57130096.6 | 42935798.8 | -0.5658714 | 0.02635667 | 0.06818633 |
| sp\|P56211\|A | 10069467.6 | 13337934.7 | 0.34612879 | 0.02635667 | 0.06818633 |
| sp\|O00469\|F | 43846157 | 54091119.5 | 0.23903786 | 0.02635667 | 0.06818633 |
| sp\|Q969H8\|I | 111076158 | 146968976 | 0.23744283 | 0.02635667 | 0.06818633 |
| sp\|P54619\|A | 3705722.26 | 4514479.61 | 0.19062337 | 0.02635667 | 0.06818633 |
| sp\|P55196\|A | 31064632 | 34723994.9 | 0.15004551 | 0.02635667 | 0.06818633 |
| sp\|Q92542\|N | 171707.194 | 232677.286 | 0.38979503 | 0.02635667 | 0.06818633 |
| sp\|Q9UPX0\|T | 688502.583 | 931332.279 | 0.45067755 | 0.02635667 | 0.06818633 |
| sp\|P31937\|3 | 351319551 | 300135504 | -0.2696154 | 0.02635667 | 0.06818633 |
| sp\|P31944\|C | 507577.804 | 1572437.45 | 1.08264736 | 0.02635667 | 0.06818633 |
| sp\|P25774\|C | 30559587.2 | 44124819.6 | 0.41475263 | 0.02635667 | 0.06818633 |
| sp\|Q15005\|S | 265986.41 | 205548.515 | -1.3535565 | 0.02635667 | 0.06818633 |
| sp\|Q9NWU1\| | 34250257.9 | 35989470.6 | 0.12845706 | 0.02635667 | 0.06818633 |
| sp\|Q96DV4\|I | 7763644.78 | 6163551.64 | -0.6132515 | 0.02635667 | 0.06818633 |
| sp\|Q15750\|T | 8481727.55 | 10603075.2 | 0.35651368 | 0.02635667 | 0.06818633 |
| sp\|Q15843\|N | 21989714.4 | 27648998.4 | 0.21768718 | 0.02635667 | 0.06818633 |
| sp\|P78347\|G | 8662730.97 | 6469924.98 | -0.5166588 | 0.02635667 | 0.06818633 |
| sp\|Q9UKK3\|I | 10777195.8 | 14883137 | 0.43239217 | 0.02635667 | 0.06818633 |
| sp\|Q9UKK9\|I | 144442153 | 169616420 | 0.21358096 | 0.02635667 | 0.06818633 |
| sp\|O60260\|F | 858219.855 | 1059747.69 | 0.27753194 | 0.0280502 | 0.07137482 |
| sp\|Q99569\|F | 756798.571 | 1066234.63 | 0.36749919 | 0.0280502 | 0.07137482 |
| sp\|Q16775\|G | 118885651 | 150587551 | 0.3167728 | 0.0280502 | 0.07137482 |
| sp\|Q16666\|I | 19394350.2 | 16479649 | -0.2314316 | 0.0280502 | 0.07137482 |
| sp\|O43602\|D | 3585542.45 | 5417709.94 | 0.48700639 | 0.0280502 | 0.07137482 |
| sp\|P29323\|E | 1840388.95 | 2398287.16 | 0.3430891 | 0.0280502 | 0.07137482 |
| sp\|Q9NPE3\|I | 49677498.1 | 50083596.7 | -0.4188533 | 0.0280502 | 0.07137482 |
| sp\|P49366\|D | 669945.321 | 972136.305 | 0.56237241 | 0.0280502 | 0.07137482 |
| sp\|Q92947\|G | 110090950 | 117835033 | 0.22969307 | 0.0280502 | 0.07137482 |
| sp\|P41217\|C | 616929.86 | 2129938.6 | -0.3432253 | 0.0280502 | 0.07137482 |
| sp\|Q9P0R6\|G | 514440.806 | 623891.046 | 0.26957604 | 0.0280502 | 0.07137482 |
| sp\|O15061\|S | 259151541 | 308350624 | 0.29897625 | 0.0280502 | 0.07137482 |
| sp\|Q5JTJ3\|C | 12388464.6 | 14114916.3 | 0.27115791 | 0.0280502 | 0.07137482 |
| sp\|Q14978\|N | 174931.906 | 176962.625 | -0.692585 | 0.0280502 | 0.07137482 |
| sp\|Q14964\|F | 56101596.9 | 69635913.1 | 0.34361423 | 0.0280502 | 0.07137482 |
| sp\|P30405\|P | 32439111.3 | 41348017.2 | 0.26855814 | 0.0280502 | 0.07137482 |

| | | | | | |
|---|---|---|---|---|---|
| sp\|Q9BXJ4\|C | 2755457.97 | 3473586.22 | 0.27234039 | 0.0280502 | 0.07137482 |
| sp\|Q9UDY4\|I | 1571792.16 | 1179684.6 | -0.8324932 | 0.0280502 | 0.07137482 |
| sp\|P14649\|M | 551160357 | 607891282 | 0.14609622 | 0.0280502 | 0.07137482 |
| sp\|O43237\|D | 17834239 | 20178186.2 | 0.15297497 | 0.0280502 | 0.07137482 |
| sp\|Q149M9\|I | 1572616.04 | 2348304.4 | 0.55939976 | 0.0280502 | 0.07137482 |
| sp\|Q96AQ6\|I | 6779078.6 | 8162198.98 | 0.21762548 | 0.0280502 | 0.07137482 |
| sp\|O00418\|E | 9979086.41 | 8196469.93 | -0.3030565 | 0.0280502 | 0.07137482 |
| sp\|Q9H082\|F | 56101596.9 | 69635913.1 | 0.34361423 | 0.0280502 | 0.07137482 |
| sp\|P14625\|E | 408104713 | 328326690 | -0.3801451 | 0.0280502 | 0.07137482 |
| sp\|Q9UHR5\| | 686392.175 | 557581.456 | -0.5337828 | 0.0280502 | 0.07137482 |
| sp\|Q9UJC5\|S | 8480594.44 | 6653723.49 | -0.3964228 | 0.0280502 | 0.07137482 |
| sp\|P52566\|G | 211025342 | 250788688 | 0.20732515 | 0.0280502 | 0.07137482 |
| sp\|Q14714\|S | 293464.687 | 385490.365 | 0.79442056 | 0.0280502 | 0.07137482 |
| sp\|P61086\|U | 2810576.41 | 3290617.83 | 0.21092964 | 0.0280502 | 0.07137482 |
| sp\|Q9Y617\|S | 181163116 | 238125441 | 0.31714464 | 0.0280502 | 0.07137482 |
| sp\|O15259\|N | 6427271.63 | 8528590.25 | 0.40875957 | 0.02983319 | 0.07503642 |
| sp\|O15382\|E | 61186239.8 | 87782390.2 | 0.29478899 | 0.02983319 | 0.07503642 |
| sp\|Q99497\|F | 1307415050 | 1567561292 | 0.19421105 | 0.02983319 | 0.07503642 |
| sp\|P50452\|S | 65097532.6 | 79185168.9 | 0.2545974 | 0.02983319 | 0.07503642 |
| sp\|P28799\|G | 26477005.7 | 35379027 | 0.33183844 | 0.02983319 | 0.07503642 |
| sp\|P51649\|S | 254674311 | 217364582 | -0.2297564 | 0.02983319 | 0.07503642 |
| sp\|Q5JRX3\|P | 56085685.2 | 48349572.7 | -0.2199469 | 0.02983319 | 0.07503642 |
| sp\|Q15274\|N | 45482059.9 | 27799847.3 | -1.0918352 | 0.02983319 | 0.07503642 |
| sp\|O60331\|F | 4089305.61 | 3344590.49 | -0.3254772 | 0.02983319 | 0.07503642 |
| sp\|P54709\|A | 7054724.37 | 6151129.28 | -0.2061807 | 0.02983319 | 0.07503642 |
| sp\|Q9BWF3\| | 28179982.9 | 24003104.6 | -0.2066428 | 0.02983319 | 0.07503642 |
| sp\|O15067\|F | 94078248.1 | 76953310.4 | -0.396127 | 0.02983319 | 0.07503642 |
| sp\|Q9P0L0\|V | 15538635.2 | 13044835.8 | -0.269975 | 0.02983319 | 0.07503642 |
| sp\|Q9BYM8\|I | 2062554.16 | 1537651.6 | -0.6285117 | 0.02983319 | 0.07503642 |
| sp\|Q8MH63\| | 3134123.1 | 4175208.73 | 0.42209782 | 0.02983319 | 0.07503642 |
| sp\|Q6XQN6\| | 246886040 | 293961749 | 0.2123303 | 0.02983319 | 0.07503642 |
| sp\|Q96C90\|F | 5167397.11 | 6065158.62 | 0.22477718 | 0.02983319 | 0.07503642 |
| sp\|P45381\|A | 78984841 | 67021491.7 | -0.2407017 | 0.02983319 | 0.07503642 |
| sp\|Q9HBH5\| | 1374524.98 | 1971155.5 | 0.38931632 | 0.02983319 | 0.07503642 |
| sp\|Q9Y371\|S | 19998877.8 | 15703801.2 | -0.3817811 | 0.02983319 | 0.07503642 |
| sp\|Q6ZS17\|F | 489752.383 | 999638.105 | 0.80540646 | 0.02983319 | 0.07503642 |
| sp\|Q13435\|S | 42921580.7 | 37753047 | -0.1941596 | 0.02983319 | 0.07503642 |
| sp\|Q9BXL7\|C | 640802.818 | 374225.019 | -0.7556371 | 0.03170908 | 0.07860116 |
| sp\|Q15555\|N | 219541.126 | 153127.773 | -2.8360687 | 0.03170908 | 0.07860116 |

| | | | | | |
|---|---|---|---|---|---|
| sp\|Q8N4T8\|( | 21600518.7 | 17585033.7 | -0.3105845 | 0.03170908 | 0.07860116 |
| sp\|Q8N129\|( | 7691517.38 | 6286865.06 | -0.3970223 | 0.03170908 | 0.07860116 |
| sp\|O95359\|T | 158420203 | 127169358 | -0.3093707 | 0.03170908 | 0.07860116 |
| sp\|O95563\|N | 3348638.85 | 2666997.22 | -0.2424017 | 0.03170908 | 0.07860116 |
| sp\|P61960\|U | 9776371.51 | 14262138 | 0.43304826 | 0.03170908 | 0.07860116 |
| sp\|Q9UPV0\|( | 1067035.12 | 1532512.93 | 0.45560896 | 0.03170908 | 0.07860116 |
| sp\|P49747\|C | 12657399.7 | 16845686.4 | 0.32454694 | 0.03170908 | 0.07860116 |
| sp\|Q96HC4\| | 140608945 | 121194574 | -0.2055076 | 0.03170908 | 0.07860116 |
| sp\|Q99832\|T | 81848935.8 | 66899429.5 | -0.4216572 | 0.03170908 | 0.07860116 |
| sp\|Q5VYX0\|F | 187134.002 | 139287.877 | 0.05365518 | 0.03170908 | 0.07860116 |
| sp\|O00425\|I | 1795442.28 | 1248445.91 | -0.4862071 | 0.03170908 | 0.07860116 |
| sp\|P46937\|Y | 88855867.6 | 108223661 | 0.24576015 | 0.03170908 | 0.07860116 |
| sp\|Q13596\|S | 16820322 | 19295864.8 | 0.16620146 | 0.03170908 | 0.07860116 |
| sp\|Q9NZ32\|A | 10864955.6 | 9331324.48 | -0.2444999 | 0.03170908 | 0.07860116 |
| sp\|Q15382\|F | 28724612.8 | 34776222.3 | 0.26712673 | 0.03170908 | 0.07860116 |
| sp\|P07737\|P | 2102481063 | 2503803676 | 0.26534957 | 0.03170908 | 0.07860116 |
| sp\|P01889\|H | 53647749.8 | 43596776.6 | -0.2899654 | 0.03170908 | 0.07860116 |
| sp\|O75208\|( | 53426000.1 | 61279588.1 | 0.23316575 | 0.03170908 | 0.07860116 |
| sp\|Q9NX46\|A | 52821061.9 | 45005320.9 | -0.2618204 | 0.03170908 | 0.07860116 |
| sp\|Q9UJ70\|N | 163990755 | 145459106 | -0.1925396 | 0.03170908 | 0.07860116 |
| sp\|O15160\|F | 1533490.39 | 1246297.45 | -0.3249693 | 0.03170908 | 0.07860116 |
| sp\|Q9Y2Q5\|I | 9183724.33 | 11014839.2 | 0.21036987 | 0.03170908 | 0.07860116 |
| sp\|P11717\|M | 40697780.6 | 45466459.1 | 0.14955803 | 0.03170908 | 0.07860116 |
| sp\|Q9NZK5\|A | 4415707 | 5726437.25 | 0.28219767 | 0.03170908 | 0.07860116 |
| sp\|Q8TEA8\|D | 14858601.1 | 17585556.9 | 0.2570711 | 0.03170908 | 0.07860116 |
| sp\|Q9HA77\|S | 7504593.88 | 9343532.84 | 0.31934118 | 0.03170908 | 0.07860116 |
| sp\|Q9UIL1\|S | 24994752.1 | 29770169.5 | 0.2515151 | 0.03368139 | 0.08242581 |
| sp\|Q7Z739\|Y | 1616826.28 | 2762963.59 | 0.54188746 | 0.03368139 | 0.08242581 |
| sp\|Q9UQ35\| | 12910777.2 | 9563568.68 | -0.4237547 | 0.03368139 | 0.08242581 |
| sp\|Q15165\|F | 3075038.2 | 2226083.83 | -0.5240307 | 0.03368139 | 0.08242581 |
| sp\|Q9UQ13\| | 924630.755 | 1454957.46 | 0.5264138 | 0.03368139 | 0.08242581 |
| sp\|Q5T0F9\|C | 5807118.82 | 7368431.44 | 0.36247729 | 0.03368139 | 0.08242581 |
| sp\|O75828\|( | 387893803 | 330505873 | -0.2433855 | 0.03368139 | 0.08242581 |
| sp\|Q5EBL4\|F | 24136236.3 | 17532446.4 | -0.5401052 | 0.03368139 | 0.08242581 |
| sp\|P67775\|P | 140019462 | 163083064 | 0.17421806 | 0.03368139 | 0.08242581 |
| sp\|Q96HY7\|I | 31032529.5 | 38203146.2 | 0.23279347 | 0.03368139 | 0.08242581 |
| sp\|P51858\|H | 119751987 | 95891636.9 | -0.4144801 | 0.03368139 | 0.08242581 |
| sp\|Q9NPH2\| | 155171262 | 114598304 | -0.4652156 | 0.03368139 | 0.08242581 |
| sp\|Q9Y237\|F | 12336543.8 | 13647475 | 0.18461747 | 0.03368139 | 0.08242581 |

| | | | | | |
|---|---|---|---|---|---|
| sp\|Q6UXI9\|N | 1573559.8 | 1187106.53 | -0.6153267 | 0.03368139 | 0.08242581 |
| sp\|P01116\|R | 24982985 | 30876072.2 | 0.28808708 | 0.03368139 | 0.08242581 |
| sp\|P15374\|U | 26884379.7 | 35234011.4 | 0.29473544 | 0.03368139 | 0.08242581 |
| sp\|Q96DZ1\|E | 6378144.78 | 9445148.48 | 0.40923662 | 0.03368139 | 0.08242581 |
| sp\|P62308\|R | 46020759.1 | 36180992 | -0.4101458 | 0.03368139 | 0.08242581 |
| sp\|P01763\|H | 838833488 | 995786877 | 0.24272448 | 0.03368139 | 0.08242581 |
| sp\|Q9NRP0\|( | 2255085.19 | 1891859.65 | -0.3489762 | 0.03368139 | 0.08242581 |
| sp\|Q13683\|I | 74599875.5 | 84729614.1 | 0.19571327 | 0.03368139 | 0.08242581 |
| sp\|P14678\|R | 84463238.9 | 69319790.9 | -0.2929338 | 0.03368139 | 0.08242581 |
| sp\|Q9Y6C2\|E | 85218592.4 | 70403480 | -0.2496314 | 0.03368139 | 0.08242581 |
| sp\|Q6P1X6\|C | 11230514.3 | 9585375.97 | -0.2656444 | 0.03368139 | 0.08242581 |
| sp\|Q9H2W6\| | 7139731.61 | 5907817.73 | -0.3796258 | 0.03368139 | 0.08242581 |
| sp\|P05109\|S | 49540916.5 | 118988031 | 0.56767939 | 0.03575381 | 0.08665784 |
| sp\|O75251\|N | 6862667.68 | 5832460.83 | -0.2813882 | 0.03575381 | 0.08665784 |
| sp\|P12829\|M | 1570356.82 | 5305677.91 | 0.57243424 | 0.03575381 | 0.08665784 |
| sp\|Q9NVS9\|I | 65178303.2 | 54020242.1 | -0.3233377 | 0.03575381 | 0.08665784 |
| sp\|Q9H0J4\|C | 692540.494 | 1562832.67 | 0.64544066 | 0.03575381 | 0.08665784 |
| sp\|P23588\|IF | 125100600 | 153553891 | 0.25192471 | 0.03575381 | 0.08665784 |
| sp\|P20061\|T | 376063.253 | 1370544.37 | 1.02165427 | 0.03575381 | 0.08665784 |
| sp\|Q8TCD5\|I | 33722977.8 | 41955932.3 | 0.19913832 | 0.03575381 | 0.08665784 |
| sp\|P09622\|D | 674387509 | 754175510 | 0.15985928 | 0.03575381 | 0.08665784 |
| sp\|Q9Y3D6\|F | 24660509.9 | 28756809.3 | 0.24683253 | 0.03575381 | 0.08665784 |
| sp\|Q9H3K6\|I | 9852009.73 | 12020938.8 | 0.27909366 | 0.03575381 | 0.08665784 |
| sp\|P61421\|V | 2385067.35 | 3774324.77 | 0.43063305 | 0.03575381 | 0.08665784 |
| sp\|Q5JS54\|P | 344306.377 | 445546.565 | 0.38927807 | 0.03575381 | 0.08665784 |
| sp\|Q9H773\|I | 1893046.87 | 2915441.18 | 0.38845773 | 0.03575381 | 0.08665784 |
| sp\|Q96FQ6\|S | 3639106.73 | 4245726.54 | 0.24471647 | 0.03575381 | 0.08665784 |
| sp\|P26440\|IV | 167488309 | 155562769 | -0.2452062 | 0.03575381 | 0.08665784 |
| sp\|P27986\|P | 261931.399 | 178598.882 | -0.4685644 | 0.03575381 | 0.08665784 |
| sp\|P14868\|S | 124494612 | 108334085 | -0.2568739 | 0.03575381 | 0.08665784 |
| sp\|Q9Y383\|L | 23451225.2 | 19785810.3 | -0.2520795 | 0.03575381 | 0.08665784 |
| sp\|O60437\|F | 341030579 | 426241364 | 0.27528637 | 0.03792995 | 0.09042516 |
| sp\|O60493\|S | 38806365.9 | 43136909.7 | 0.14629763 | 0.03792995 | 0.09042516 |
| sp\|P56277\|C | 22155397 | 28893971.7 | 0.28561197 | 0.03792995 | 0.09042516 |
| sp\|Q04637\|I | 10722785.4 | 12379850.7 | 0.22013825 | 0.03792995 | 0.09042516 |
| sp\|Q15113\|F | 38029943.6 | 50642798.7 | 0.29634536 | 0.03792995 | 0.09042516 |
| sp\|Q9BZ67\|F | 2568347.95 | 3151381.85 | 0.176335 | 0.03792995 | 0.09042516 |
| sp\|P53041\|P | 16907941.1 | 14732810.5 | -0.2296838 | 0.03792995 | 0.09042516 |
| sp\|P54252\|A | 3986666.86 | 11101003.4 | 0.73719641 | 0.03792995 | 0.09042516 |

| | | | | | |
|---|---|---|---|---|---|
| sp\|Q8NBJ9\|S | 594561.414 | 850728.849 | 0.50691231 | 0.03792995 | 0.09042516 |
| sp\|P51854\|T | 4327740.02 | 6093355.88 | 0.40097953 | 0.03792995 | 0.09042516 |
| sp\|P51606\|R | 56354546.8 | 73187050.4 | 0.26187908 | 0.03792995 | 0.09042516 |
| sp\|Q9H446\|I | 499559.391 | 782299.431 | 0.59922976 | 0.03792995 | 0.09042516 |
| sp\|Q9NYB0\|T | 809075.258 | 719036.117 | -0.1255839 | 0.03792995 | 0.09042516 |
| sp\|P18850\|A | 11807772.2 | 16121290.3 | 0.47019946 | 0.03792995 | 0.09042516 |
| sp\|O95164\|U | 102383.923 | 212047.194 | 0.05850021 | 0.03792995 | 0.09042516 |
| sp\|Q9BWU0\| | 682990.449 | 495561.85 | -0.5805982 | 0.03792995 | 0.09042516 |
| sp\|Q9UPN7\| | 3149332.84 | 4548750.9 | 0.426317 | 0.03792995 | 0.09042516 |
| sp\|Q13131\|A | 5962980.58 | 4611163.14 | -0.4727706 | 0.03792995 | 0.09042516 |
| sp\|Q14185\|D | 984344.678 | 2032987.27 | 0.79369386 | 0.03792995 | 0.09042516 |
| sp\|Q92973\|T | 724053.671 | 939546.557 | 0.38221611 | 0.03792995 | 0.09042516 |
| sp\|Q9NUP7\|T | 4052509.76 | 5858532.98 | 0.43908993 | 0.03792995 | 0.09042516 |
| sp\|P47755\|C | 449939668 | 390272246 | -0.2703398 | 0.03792995 | 0.09042516 |
| sp\|P62328\|T | 222823071 | 322931729 | 0.34084357 | 0.03792995 | 0.09042516 |
| sp\|P19971\|T | 34783567.5 | 58356542.9 | 0.6715501 | 0.03792995 | 0.09042516 |
| sp\|O95497\|V | 3359814.59 | 3967359.2 | 0.39175723 | 0.03792995 | 0.09042516 |
| sp\|Q8IZT6\|AS | 2536632.21 | 3064649.37 | 0.21610607 | 0.03792995 | 0.09042516 |
| sp\|O95704\|A | 2776225.06 | 3469305.73 | 0.42633652 | 0.03792995 | 0.09042516 |
| sp\|Q9C0B7\|T | 2228465.11 | 3915320.51 | 0.80046545 | 0.03792995 | 0.09042516 |
| sp\|Q96I23\|P | 3681597.12 | 3573671.29 | 0.13208876 | 0.03792995 | 0.09042516 |
| sp\|Q9UHY7\|I | 13983935.7 | 11610978.7 | -0.4188007 | 0.03792995 | 0.09042516 |
| sp\|Q9ULC3\|I | 19690983.8 | 17327786.9 | -0.2080501 | 0.03792995 | 0.09042516 |
| sp\|Q7L523\|R | 15479984.3 | 12998942.8 | -0.3261409 | 0.03792995 | 0.09042516 |
| sp\|P42331\|R | 1034207.3 | 687071.423 | -0.5629776 | 0.03792995 | 0.09042516 |
| sp\|P10253\|L | 100207982 | 141216491 | 0.34545102 | 0.04021353 | 0.09446145 |
| sp\|Q9UQB8\| | 316570.58 | 247170.792 | -0.430573 | 0.04021353 | 0.09446145 |
| sp\|O76024\|W | 6024335.2 | 8172306.13 | 0.27282383 | 0.04021353 | 0.09446145 |
| sp\|O75334\|L | 318647.053 | 536273.382 | 0.60835517 | 0.04021353 | 0.09446145 |
| sp\|Q8WX93\| | 84463545.5 | 70154036.8 | -0.2387301 | 0.04021353 | 0.09446145 |
| sp\|Q9BX67\|J | 5433765.67 | 6849734.22 | 0.24283405 | 0.04021353 | 0.09446145 |
| sp\|Q9H000\|I | 6226809.41 | 11536311.8 | 0.49967266 | 0.04021353 | 0.09446145 |
| sp\|Q8TD16\|E | 1596312.9 | 1941013.21 | 0.33687393 | 0.04021353 | 0.09446145 |
| sp\|P62304\|R | 46980938 | 37571801.3 | -0.4320937 | 0.04021353 | 0.09446145 |
| sp\|Q99873\|A | 47844161.1 | 40278009.4 | -0.3423809 | 0.04021353 | 0.09446145 |
| sp\|P08397\|H | 35031464.6 | 52463426.6 | 0.43421193 | 0.04021353 | 0.09446145 |
| sp\|Q9NZN3\|I | 25508850.8 | 30095639.9 | 0.22697813 | 0.04021353 | 0.09446145 |
| sp\|Q7Z4W1\| | 320414683 | 397835320 | 0.17550998 | 0.04021353 | 0.09446145 |
| sp\|O95155\|U | 10624946.8 | 16097094.9 | 0.30659894 | 0.04021353 | 0.09446145 |

| | | | | | |
|---|---|---|---|---|---|
| sp\|P82930\|R | 9787122.9 | 7681211.68 | -0.4118983 | 0.04021353 | 0.09446145 |
| sp\|Q9H6Q4\| | 5405714.83 | 8092736.89 | 0.39421888 | 0.04021353 | 0.09446145 |
| sp\|Q9BSV6\|S | 1133699.99 | 1476526.87 | 0.42775678 | 0.04021353 | 0.09446145 |
| sp\|Q9UII2\|AT | 45205068.3 | 55921585.4 | 0.26480819 | 0.04021353 | 0.09446145 |
| sp\|A5D8V6\|V | 3013943.89 | 3460310.86 | 0.19758657 | 0.04021353 | 0.09446145 |
| sp\|Q9ULP9\|T | 3186778.36 | 4150103.07 | 0.27550561 | 0.04021353 | 0.09446145 |
| sp\|P07305\|H | 79960330.7 | 58435467.9 | -0.4073568 | 0.04021353 | 0.09446145 |
| sp\|Q6PL24\|T | 23637711 | 20028992.9 | -0.2608114 | 0.04021353 | 0.09446145 |
| sp\|P23511\|N | 12350838.6 | 16349760.9 | 0.32361639 | 0.04021353 | 0.09446145 |
| sp\|P04424\|A | 63677629.4 | 71826370.8 | 0.17473724 | 0.04021353 | 0.09446145 |
| sp\|Q8TB22\|S | 3754884.24 | 4855202.53 | 0.37623007 | 0.04021353 | 0.09446145 |
| sp\|Q96LD8\|S | 75354006.5 | 108902798 | 0.82459141 | 0.04021353 | 0.09446145 |
| sp\|Q15008\|F | 16811436.9 | 14039324 | -0.3173775 | 0.04021353 | 0.09446145 |
| sp\|Q96PU8\|C | 4247370.5 | 3292679.69 | -0.3795102 | 0.04021353 | 0.09446145 |
| sp\|L0R8F8\|M | 671149.059 | 1193381.44 | 1.21122252 | 0.04021353 | 0.09446145 |
| sp\|A0A075B6 | 22308512.2 | 34114218.2 | 0.57472655 | 0.04021353 | 0.09446145 |
| sp\|P28845\|D | 107167.086 | 64675.7783 | -4.5621402 | 0.04023868 | 0.09447427 |
| sp\|P01705\|L | 20091616.2 | 25960559.5 | 0.38940366 | 0.04260839 | 0.09873378 |
| sp\|Q9NR28\|I | 10161424.5 | 12305814.7 | 0.28485597 | 0.04260839 | 0.09873378 |
| sp\|Q99714\|H | 290832076 | 332223275 | 0.22007585 | 0.04260839 | 0.09873378 |
| sp\|O95721\|S | 7416715.12 | 8442796.96 | 0.20765762 | 0.04260839 | 0.09873378 |
| sp\|Q14511\|C | 3215674.84 | 2797832.59 | -0.2056903 | 0.04260839 | 0.09873378 |
| sp\|Q8IUD2\|F | 47140042.8 | 52214118 | 0.14154718 | 0.04260839 | 0.09873378 |
| sp\|P01903\|D | 12346799.5 | 17869763.1 | 0.3834027 | 0.04260839 | 0.09873378 |
| sp\|O00499\|B | 72033838.9 | 80697937.6 | 0.15220756 | 0.04260839 | 0.09873378 |
| sp\|Q9H1K1\|I | 11858122.1 | 10095013.7 | -0.2297659 | 0.04260839 | 0.09873378 |
| sp\|Q9Y277\|V | 19988734.9 | 17384775.8 | -0.1833367 | 0.04260839 | 0.09873378 |
| sp\|O75955\|F | 27590959.6 | 33558971.8 | 0.26424933 | 0.04260839 | 0.09873378 |
| sp\|O60678\|A | 3744514.26 | 3037891.19 | -0.4340537 | 0.04260839 | 0.09873378 |
| sp\|A4UGR9\|X | 1146122.34 | 1521446.41 | -0.5807249 | 0.04260839 | 0.09873378 |
| sp\|P62745\|R | 16069904 | 18826414.7 | 0.19381017 | 0.04260839 | 0.09873378 |
| sp\|O00187\|N | 5105482.62 | 4177307.05 | -0.4280372 | 0.04260839 | 0.09873378 |
| sp\|C9JLW8\|N | 5592954.5 | 7067508.26 | 0.30506074 | 0.04260839 | 0.09873378 |
| sp\|P20774\|M | 1053491880 | 911311965 | -0.6722908 | 0.04260839 | 0.09873378 |
| sp\|Q9C0B5\|Z | 3913208.65 | 5143132.16 | 0.22817234 | 0.04260839 | 0.09873378 |
| sp\|O75157\|T | 6797287.12 | 8256251.07 | 0.32911812 | 0.04260839 | 0.09873378 |
| sp\|O43164\|F | 3810703.3 | 4358769.78 | 0.35397448 | 0.04260839 | 0.09873378 |
| sp\|P17544\|A | 246633.616 | 429319.504 | 0.61504812 | 0.04260839 | 0.09873378 |
| sp\|Q92620\|F | 263753.428 | 187041.097 | -1.851601 | 0.04260839 | 0.09873378 |

| | | | | | |
|---|---|---|---|---|---|
| sp\|P00751\|C | 1416466348 | 1077680795 | -0.7012114 | 0.04260839 | 0.09873378 |
| sp\|Q9Y3B2\|E | 6404468.57 | 5564711.56 | -0.2035739 | 0.04260839 | 0.09873378 |
| sp\|Q3B8N2\|I | 6686508.9 | 4781654.1 | -0.4721154 | 0.04260839 | 0.09873378 |
| sp\|Q9NX40\|( | 226478.146 | 155320.894 | -0.7601949 | 0.04260839 | 0.09873378 |
| sp\|P27824\|C | 104413873 | 84976332.5 | -0.4110393 | 0.04260839 | 0.09873378 |
| sp\|Q9NR12\|I | 102919911 | 79142618.8 | -0.2790146 | 0.04511831 | 0.10320437 |
| sp\|Q99767\|A | 2552786.48 | 3225908.12 | 0.25910549 | 0.04511831 | 0.10320437 |
| sp\|P50570\|D | 652599.282 | 3166894.34 | 0.75534519 | 0.04511831 | 0.10320437 |
| sp\|P61758\|P | 13936220.3 | 16039326.4 | 0.22720153 | 0.04511831 | 0.10320437 |
| sp\|Q8IXB3\|T/ | 21003926.4 | 26403575.2 | 0.31894983 | 0.04511831 | 0.10320437 |
| sp\|Q9Y2Y0\|A | 10920203.7 | 21035808.6 | 0.62096094 | 0.04511831 | 0.10320437 |
| sp\|P13073\|C | 16198330.8 | 14156333.3 | -0.1826868 | 0.04511831 | 0.10320437 |
| sp\|Q9Y2U8\|M | 129471.382 | 205050.723 | 0.40908598 | 0.04511831 | 0.10320437 |
| sp\|P16885\|P | 673540.632 | 460609.056 | -0.709585 | 0.04511831 | 0.10320437 |
| sp\|P14854\|C | 60437769.9 | 71309219.3 | 0.32869948 | 0.04511831 | 0.10320437 |
| sp\|P36915\|G | 8647790.88 | 6718615.32 | -0.3789439 | 0.04511831 | 0.10320437 |
| sp\|O00399\|D | 43159535.2 | 49211985.1 | 0.17154323 | 0.04511831 | 0.10320437 |
| sp\|Q9UJU6\|D | 139703575 | 156761015 | 0.15898059 | 0.04511831 | 0.10320437 |
| sp\|Q9BTY2\|F | 10908442.9 | 14248189.3 | 0.25316567 | 0.04511831 | 0.10320437 |
| sp\|P07919\|Q | 18549341.8 | 16511675.3 | 0.63530331 | 0.04511831 | 0.10320437 |
| sp\|P82921\|R | 3991392.19 | 3519638.95 | -0.2587286 | 0.04511831 | 0.10320437 |
| sp\|Q6AI12\|A | 54419694.3 | 64312257.1 | 0.22184151 | 0.04511831 | 0.10320437 |
| sp\|Q13938\|( | 732603.377 | 758640.493 | 0.7297491 | 0.04511831 | 0.10320437 |
| sp\|Q7Z3F1\|G | 342935.952 | 518100.372 | 0.27606957 | 0.04511831 | 0.10320437 |
| sp\|Q68DK2\|Z | 3482275.7 | 4568186.82 | 0.44937518 | 0.04511831 | 0.10320437 |
| sp\|Q86WV6\| | 16149729.1 | 12588575.6 | -0.4503741 | 0.04511831 | 0.10320437 |
| sp\|P25311\|Z | 987150311 | 1371789141 | 0.46156492 | 0.04511831 | 0.10320437 |
| sp\|P29373\|R | 16803824.2 | 13511488.2 | -0.3677395 | 0.04511831 | 0.10320437 |
| sp\|P13984\|T | 1875954.3 | 3854828.63 | 1.2397055 | 0.04511831 | 0.10320437 |
| sp\|P55789\|A | 4060769.99 | 4075178.85 | 0.05950446 | 0.04511831 | 0.10320437 |
| sp\|Q8TDZ2\|M | 1658772.08 | 3522539.49 | 0.4786791 | 0.04511831 | 0.10320437 |
| sp\|Q9H9G7\|. | 34075715.4 | 29095417.5 | -0.3215008 | 0.04511831 | 0.10320437 |
| sp\|Q86WC4\| | 1577105.29 | 2054180.3 | 0.40987564 | 0.04774715 | 0.10762732 |
| sp\|P09132\|S | 504725.319 | 395150.056 | -0.718533 | 0.04774715 | 0.10762732 |
| sp\|Q7Z3E5\|A | 13137528.2 | 10823750.4 | -0.39609 | 0.04774715 | 0.10762732 |
| sp\|P35221\|C | 80395103.2 | 97920966.6 | 0.25846265 | 0.04774715 | 0.10762732 |
| sp\|Q9BVC4\|I | 7796351.79 | 10258414.1 | 0.22188481 | 0.04774715 | 0.10762732 |
| sp\|Q15007\|F | 364187.37 | 294349.944 | -0.3568323 | 0.04774715 | 0.10762732 |
| sp\|O60861\|( | 2032468.33 | 3300918.1 | 0.60575047 | 0.04774715 | 0.10762732 |

| | | | | | |
|---|---|---|---|---|---|
| sp\|P30050\|R | 67142198 | 54905670.3 | -0.3095184 | 0.04774715 | 0.10762732 |
| sp\|Q9UBF8\|I | 1968221.12 | 4133918.46 | 0.90331165 | 0.04774715 | 0.10762732 |
| sp\|P51178\|P | 99879039.8 | 134766011 | 0.10418947 | 0.04774715 | 0.10762732 |
| sp\|Q00722\|F | 321552.922 | 215223.434 | -1.0041278 | 0.04774715 | 0.10762732 |
| sp\|O95865\|D | 620834234 | 541151921 | -0.1857078 | 0.04774715 | 0.10762732 |
| sp\|Q99805\|T | 1126337.68 | 1085591.43 | 0.81295847 | 0.04774715 | 0.10762732 |
| sp\|P11142\|H | 3078293887 | 3351051128 | 0.17865754 | 0.04774715 | 0.10762732 |
| sp\|P34932\|H | 273192275 | 227236221 | -0.3743706 | 0.04774715 | 0.10762732 |
| sp\|Q14974\|I | 53476156.1 | 70769990.9 | 0.79914349 | 0.04774715 | 0.10762732 |
| sp\|Q9H6U6\| | 944579.5 | 1099104.78 | 0.19670402 | 0.04774715 | 0.10762732 |
| sp\|Q9NNW7\| | 61836310.6 | 78670643.4 | 0.22133174 | 0.04774715 | 0.10762732 |
| sp\|Q8N4V1\|I | 708265.078 | 992993.887 | 0.23821788 | 0.04774715 | 0.10762732 |
| sp\|Q9BX66\|S | 512272119 | 641743616 | 0.34200122 | 0.04774715 | 0.10762732 |
| sp\|O75569\|F | 614167.85 | 498853.064 | -0.5144517 | 0.04774715 | 0.10762732 |
| sp\|P82932\|R | 382061.618 | 262854.674 | -0.7754524 | 0.04774715 | 0.10762732 |
| sp\|P00734\|T | 950470165 | 790199513 | -0.3526201 | 0.04774715 | 0.10762732 |
| sp\|Q9H3P2\|I | 1023998.79 | 701261.536 | -0.4803469 | 0.04774715 | 0.10762732 |
| sp\|P02795\|M | 24409413.9 | 38197862.7 | 0.37734219 | 0.04774715 | 0.10762732 |
| sp\|O00478\|E | 2765535.26 | 2018256.31 | -0.5161407 | 0.04774715 | 0.10762732 |
| sp\|A8MXV4\|N | 10865078.4 | 13958811.8 | 0.23880813 | 0.04774715 | 0.10762732 |
| sp\|P41567\|E | 4014008.53 | 3340666.11 | -0.382879 | 0.04774715 | 0.10762732 |
| sp\|P42224\|S | 8052920.79 | 6467595.48 | -0.3572621 | 0.04774715 | 0.10762732 |
| sp\|P08134\|R | 147743054 | 177320047 | 0.23167348 | 0.04774715 | 0.10762732 |
| sp\|Q9BUH6\| | 14269504.3 | 12431149.1 | -0.2031346 | 0.04774715 | 0.10762732 |
| sp\|Q96A70\|A | 699455.566 | 955440.819 | 0.41742906 | 0.05049889 | 0.11319204 |
| sp\|Q5T013\|H | 958985247 | 1170632546 | 0.1556478 | 0.05049889 | 0.11319204 |
| sp\|P35659\|D | 6935997.97 | 5828754.42 | -0.2579995 | 0.05049889 | 0.11319204 |
| sp\|Q15031\|S | 4200703.72 | 3451309.85 | -0.3289145 | 0.05049889 | 0.11319204 |
| sp\|Q96PV0\|S | 5651682.51 | 3964712.76 | -1.3146845 | 0.05049889 | 0.11319204 |
| sp\|P22748\|C | 12295849.2 | 15120920.9 | 0.35277875 | 0.05049889 | 0.11319204 |
| sp\|Q92930\|F | 120591120 | 134439373 | 0.14625374 | 0.05049889 | 0.11319204 |
| sp\|Q8IUE1\|T | 35116501.2 | 42910942.2 | 0.40383337 | 0.05049889 | 0.11319204 |
| sp\|Q5TAQ9\|I | 4616923.05 | 3566818.15 | -0.6097614 | 0.05049889 | 0.11319204 |
| sp\|O15258\|F | 3092450.89 | 2648581.33 | -0.4712171 | 0.05049889 | 0.11319204 |
| sp\|O15260\|S | 21637709.3 | 17604148 | -0.4978638 | 0.05049889 | 0.11319204 |
| sp\|P41208\|C | 6181166.48 | 5154557.96 | -0.3196154 | 0.05049889 | 0.11319204 |
| sp\|Q9Y6Q9\|I | 5856525 | 7564892.45 | 0.2443931 | 0.05337744 | 0.11831793 |
| sp\|P78346\|R | 429730.458 | 327366.438 | -1.4128763 | 0.05337744 | 0.11831793 |
| sp\|O95834\|E | 160647152 | 136379913 | -0.2848902 | 0.05337744 | 0.11831793 |

| | | | | | |
|---|---|---|---|---|---|
| sp\|O95967\|F | 12104189 | 8269739.09 | -0.4219901 | 0.05337744 | 0.11831793 |
| sp\|Q14393\|( | 7658600.93 | 12750009.5 | 0.65384283 | 0.05337744 | 0.11831793 |
| sp\|O95831\|/ | 121982756 | 132797348 | 0.1863173 | 0.05337744 | 0.11831793 |
| sp\|P00738\|H | 1.004E+10 | 1.3122E+10 | 0.27493846 | 0.05337744 | 0.11831793 |
| sp\|O43765\|S | 14216267.9 | 15498519.4 | 0.11551012 | 0.05337744 | 0.11831793 |
| sp\|P57088\|T | 3209295.03 | 4095477.42 | 0.28231028 | 0.05337744 | 0.11831793 |
| sp\|Q9H792\|I | 304494.783 | 388431.74 | 0.2958733 | 0.05337744 | 0.11831793 |
| sp\|Q9UL46\|F | 20936627.4 | 28243503.9 | 0.3683388 | 0.05337744 | 0.11831793 |
| sp\|Q99700\|/ | 10424703.8 | 11739107.3 | 0.16806084 | 0.05337744 | 0.11831793 |
| sp\|Q6FIF0\|ZI | 1016606.48 | 1149176.78 | 0.15010296 | 0.05337744 | 0.11831793 |
| sp\|P24043\|L | 49343076.3 | 58678424.9 | 0.24361433 | 0.05337744 | 0.11831793 |
| sp\|Q86WA6\| | 18370164.1 | 14681729 | -0.4219608 | 0.05337744 | 0.11831793 |
| sp\|P52209\|6 | 1981330742 | 2538054831 | 0.39871936 | 0.05337744 | 0.11831793 |
| sp\|Q9NZJ7\|M | 3711819.79 | 3721116.75 | -0.3441873 | 0.05337744 | 0.11831793 |
| sp\|Q9BQ52\|I | 23375416.3 | 21859547.6 | -0.1586905 | 0.05337744 | 0.11831793 |
| sp\|P49441\|II | 279615.942 | 222838.331 | -0.543877 | 0.05337744 | 0.11831793 |
| sp\|Q96EE3\|S | 4811945.33 | 4257448.76 | -0.2157215 | 0.05337744 | 0.11831793 |
| sp\|P51513\|N | 20139057.9 | 23241991.3 | 0.21179966 | 0.05337744 | 0.11831793 |
| sp\|Q4G0J3\|L | 6098820.68 | 5150612.92 | -0.2860706 | 0.05337744 | 0.11831793 |
| sp\|Q9NT62\|/ | 1182106.04 | 1511717.8 | 0.34234681 | 0.05337744 | 0.11831793 |
| sp\|Q9Y6E0\|S | 5039161.79 | 4326017.53 | -0.2971164 | 0.05337744 | 0.11831793 |
| sp\|Q01459\|D | 5860255.26 | 6958082.23 | 0.18945863 | 0.05638675 | 0.12378776 |
| sp\|Q6PCB7\|S | 7780449.49 | 11329838.6 | 0.32399438 | 0.05638675 | 0.12378776 |
| sp\|Q9ULC0\|I | 378902.361 | 459993.978 | 0.35210599 | 0.05638675 | 0.12378776 |
| sp\|Q9UJ68\|M | 20302615.9 | 26257410.9 | 0.24373146 | 0.05638675 | 0.12378776 |
| sp\|Q8IYK4\|G | 3231661.07 | 2609536.8 | -0.4184758 | 0.05638675 | 0.12378776 |
| sp\|Q13573\|S | 1711712.7 | 2399659.63 | 0.42748024 | 0.05638675 | 0.12378776 |
| sp\|P35914\|H | 67298193.3 | 73472827.6 | 0.1138446 | 0.05638675 | 0.12378776 |
| sp\|Q9UPQ9\|T | 473631.899 | 367681.439 | -0.4884104 | 0.05638675 | 0.12378776 |
| sp\|Q8IZP9\|A | 410814.352 | 536903.943 | 0.42202975 | 0.05638675 | 0.12378776 |
| sp\|Q9NZB8\|I | 28303886.4 | 24230250.7 | -0.249484 | 0.05638675 | 0.12378776 |
| sp\|O14618\|( | 107485836 | 133150001 | 0.24792753 | 0.05638675 | 0.12378776 |
| sp\|Q9UMY4\| | 8398367.74 | 12745745.9 | 0.32570432 | 0.05638675 | 0.12378776 |
| sp\|P46940\|I( | 106392940 | 87590683.8 | -0.2875344 | 0.05638675 | 0.12378776 |
| sp\|Q96G03\|I | 298977680 | 272909191 | -0.1943419 | 0.05638675 | 0.12378776 |
| sp\|P50583\|A | 4606714.73 | 5742968.06 | 0.27616362 | 0.05638675 | 0.12378776 |
| sp\|Q13564\|U | 93780720.6 | 82491695.9 | -0.2248364 | 0.05638675 | 0.12378776 |
| sp\|O95870\|/ | 3652349.76 | 5590452.67 | 0.39076262 | 0.05638675 | 0.12378776 |
| sp\|Q8TF74\|V | 404205.812 | 1708546.11 | 0.89409653 | 0.05638675 | 0.12378776 |

| | | | | | |
|---|---|---|---|---|---|
| sp\|Q9NR30\|I | 989163.665 | 1305277.56 | 0.32812093 | 0.05638675 | 0.12378776 |
| sp\|P19784\|C | 19889564.9 | 16913991.4 | -0.3151803 | 0.05638675 | 0.12378776 |
| sp\|Q9Y6U3\|A | 107230322 | 75915875 | -0.6084243 | 0.05638675 | 0.12378776 |
| sp\|P07951\|T | 1833467786 | 1405410948 | -0.2742366 | 0.05953093 | 0.12915413 |
| sp\|P02790\|H | 1.1952E+10 | 1.55E+10 | 0.38265035 | 0.05953093 | 0.12915413 |
| sp\|Q96GS4\|I | 410945.136 | 582887.184 | 0.33982016 | 0.05953093 | 0.12915413 |
| sp\|Q7L4P6\|E | 1155825.01 | 1583129.04 | 0.3283725 | 0.05953093 | 0.12915413 |
| sp\|O60701\|U | 52551188.4 | 45496945.2 | -0.3646917 | 0.05953093 | 0.12915413 |
| sp\|Q9UBW5\| | 6154163.46 | 6908899.33 | 0.16367589 | 0.05953093 | 0.12915413 |
| sp\|Q8IX12\|C | 976100.195 | 854041.007 | -0.1516912 | 0.05953093 | 0.12915413 |
| sp\|O75376\|N | 2576631.21 | 3681896.21 | 0.48983693 | 0.05953093 | 0.12915413 |
| sp\|O95202\|L | 18552830.7 | 22024344.3 | 0.25629208 | 0.05953093 | 0.12915413 |
| sp\|Q9UI10\|E | 337190.481 | 498503.247 | 0.53570245 | 0.05953093 | 0.12915413 |
| sp\|P48960\|A | 828989.934 | 1133732.46 | 0.3282049 | 0.05953093 | 0.12915413 |
| sp\|Q9Y3D5\|I | 370546.087 | 394227.255 | -1.1656511 | 0.05953093 | 0.12915413 |
| sp\|P49023\|P | 28100777.1 | 31973065.8 | 0.16560888 | 0.05953093 | 0.12915413 |
| sp\|Q8IV33\|K | 253574956 | 315364267 | 0.35960952 | 0.05953093 | 0.12915413 |
| sp\|Q01469\|F | 2432909276 | 2035460299 | -0.3020297 | 0.05953093 | 0.12915413 |
| sp\|P28331\|N | 75591133.1 | 88088706.4 | 0.1616145 | 0.05953093 | 0.12915413 |
| sp\|Q14197\|I | 6415516.9 | 5359903.4 | -0.2910582 | 0.05953093 | 0.12915413 |
| sp\|O95785\|V | 1771142.6 | 2360650.19 | 0.35975654 | 0.05953093 | 0.12915413 |
| sp\|P08758\|A | 1097533602 | 1848288064 | 0.7220424 | 0.05953093 | 0.12915413 |
| sp\|Q96FE7\|F | 622465.358 | 986267.235 | 0.45732553 | 0.05953093 | 0.12915413 |
| sp\|P16035\|T | 1548022.72 | 1978245.44 | 0.3261744 | 0.05953093 | 0.12915413 |
| sp\|P15169\|C | 21234151 | 17414722.2 | -0.8221961 | 0.05953093 | 0.12915413 |
| sp\|Q9UKG9\| | 102690.496 | 133119.874 | 0.59720739 | 0.05953093 | 0.12915413 |
| sp\|Q7Z5H3\|I | 155118.485 | 233556.926 | 0.34680663 | 0.05953093 | 0.12915413 |
| sp\|O43310\|C | 2283325.06 | 3319594.88 | 0.63077824 | 0.05953093 | 0.12915413 |
| sp\|Q9NTJ4\|M | 369847594 | 339525752 | -0.2039784 | 0.05953093 | 0.12915413 |
| sp\|P27144\|K | 5952045.34 | 7295949.5 | 0.19170454 | 0.06281395 | 0.13469353 |
| sp\|Q9H553\|A | 588528.675 | 428140.246 | -0.7276943 | 0.06281395 | 0.13469353 |
| sp\|Q9GZS3\|V | 13432563.3 | 15463314.9 | 0.18690701 | 0.06281395 | 0.13469353 |
| sp\|Q9Y263\|F | 7446273.51 | 8929071.78 | 0.23549015 | 0.06281395 | 0.13469353 |
| sp\|Q05086\|U | 14863364.8 | 12236138.1 | -0.2735289 | 0.06281395 | 0.13469353 |
| sp\|Q13586\|S | 4980803.77 | 5777033.75 | 0.218457 | 0.06281395 | 0.13469353 |
| sp\|Q8WW59 | 39749315.6 | 47080546.6 | 0.20175688 | 0.06281395 | 0.13469353 |
| sp\|P60673\|P | 649259.946 | 824700.82 | 0.37186284 | 0.06281395 | 0.13469353 |
| sp\|Q13615\|N | 1092578.3 | 1553849.72 | 0.47177718 | 0.06281395 | 0.13469353 |
| sp\|P25325\|T | 207795663 | 164619558 | -0.6110404 | 0.06281395 | 0.13469353 |

| | | | | | |
|---|---|---|---|---|---|
| sp\|O43665\|F | 14796907.8 | 19166734.8 | 0.34156556 | 0.06281395 | 0.13469353 |
| sp\|Q96JD6\|A | 99341527.4 | 83018724.8 | -0.3368831 | 0.06281395 | 0.13469353 |
| sp\|Q9BV19\|C | 669641.195 | 817795.611 | -0.3586988 | 0.06281395 | 0.13469353 |
| sp\|P40926\|M | 1321771249 | 1493113824 | 0.11387606 | 0.06281395 | 0.13469353 |
| sp\|Q6IQ22\|R | 61948413.1 | 67036903.4 | 0.10553247 | 0.06281395 | 0.13469353 |
| sp\|Q9BS26\|E | 130329908 | 111016136 | -0.2699763 | 0.06281395 | 0.13469353 |
| sp\|Q96EL3\|F | 16386216.4 | 20930558.3 | 0.29053251 | 0.06281395 | 0.13469353 |
| sp\|Q7Z7N9\|T | 739523.903 | 1072582.71 | 0.59112004 | 0.06281395 | 0.13469353 |
| sp\|O14841\|C | 202572877 | 172814302 | -0.3079099 | 0.06281395 | 0.13469353 |
| sp\|P22692\|IE | 27623334.8 | 32790984.1 | 0.21266228 | 0.06281395 | 0.13469353 |
| sp\|Q96L73\|N | 19039243 | 27291917.8 | 0.17038024 | 0.06281395 | 0.13469353 |
| sp\|O75410\|T | 19318512.6 | 21081847 | 0.12782195 | 0.06281395 | 0.13469353 |
| sp\|P51148\|R | 44464914.3 | 51282246.4 | 0.17867073 | 0.06281395 | 0.13469353 |
| sp\|P43034\|L | 173770347 | 207985690 | 0.18703472 | 0.06281395 | 0.13469353 |
| sp\|Q9Y490\|T | 304746009 | 389760143 | 0.25246603 | 0.06281395 | 0.13469353 |
| sp\|O95571\|E | 79722515.7 | 66634315.3 | -0.2635456 | 0.06281395 | 0.13469353 |
| sp\|Q9H3H3\| | 61305427.5 | 54103564.3 | -0.2141024 | 0.06623983 | 0.14065706 |
| sp\|P20674\|C | 20668458.5 | 25332476.5 | 0.36516064 | 0.06623983 | 0.14065706 |
| sp\|Q9P2G1\|A | 3748275.5 | 3147623.86 | -0.230475 | 0.06623983 | 0.14065706 |
| sp\|P05161\|IS | 18774147.4 | 22542758.9 | 0.27605024 | 0.06623983 | 0.14065706 |
| sp\|P55072\|T | 2158514659 | 1804336148 | -0.3898596 | 0.06623983 | 0.14065706 |
| sp\|Q99704\|D | 1688377.31 | 2065316.13 | 0.23679341 | 0.06623983 | 0.14065706 |
| sp\|O95425\|S | 14934009.4 | 13267111.9 | -0.1683593 | 0.06623983 | 0.14065706 |
| sp\|P00390\|G | 284350908 | 340606986 | 0.2160654 | 0.06623983 | 0.14065706 |
| sp\|Q96T51\|F | 6721397.18 | 5480562.02 | -0.3141316 | 0.06623983 | 0.14065706 |
| sp\|Q8N556\|A | 3524334.21 | 3028168.48 | -0.2826253 | 0.06623983 | 0.14065706 |
| sp\|Q9NP80\|I | 18146878.4 | 22624851.1 | 0.31878223 | 0.06623983 | 0.14065706 |
| sp\|P21964\|C | 32085055 | 27251344.8 | -0.311052 | 0.06623983 | 0.14065706 |
| sp\|Q6UXV4\|I | 12431408.3 | 14991890.2 | 0.18381363 | 0.06623983 | 0.14065706 |
| sp\|Q8N465\|I | 8676869.89 | 10129133.5 | 0.17902986 | 0.06623983 | 0.14065706 |
| sp\|Q9P0M9\|I | 4121067.59 | 5572122.36 | 0.22506427 | 0.06623983 | 0.14065706 |
| sp\|Q5VUB5\|I | 1116984.97 | 1347511.58 | 0.34246978 | 0.06623983 | 0.14065706 |
| sp\|P35241\|R | 996660309 | 873994346 | -0.204825 | 0.06623983 | 0.14065706 |
| sp\|Q8IUW5\|I | 784860.28 | 1041997.62 | 0.45843466 | 0.06623983 | 0.14065706 |
| sp\|O95633\|F | 5435505.17 | 6598157.29 | 0.35620171 | 0.06623983 | 0.14065706 |
| sp\|P82675\|R | 4176803.74 | 3509140.65 | -0.275365 | 0.06623983 | 0.14065706 |
| sp\|Q5VW36\| | 1585625.66 | 2075225.84 | 0.27489628 | 0.06623983 | 0.14065706 |
| sp\|P62942\|F | 152106631 | 165806540 | 0.15600373 | 0.06623983 | 0.14065706 |
| sp\|O94916\|N | 614101.909 | 466284.41 | -0.440618 | 0.06981274 | 0.14617423 |

| | | | | | |
|---|---|---|---|---|---|
| sp\|A0MZ66\|S | 134050642 | 150635206 | 0.14813798 | 0.06981274 | 0.14617423 |
| sp\|Q99707\|N | 16013862.7 | 19120701.4 | 0.2103073 | 0.06981274 | 0.14617423 |
| sp\|P23975\|S | 4421019.82 | 2698465.52 | -0.9733606 | 0.06981274 | 0.14617423 |
| sp\|Q8TB45\|D | 15211788.6 | 26434144.9 | 2.7706889 | 0.06981274 | 0.14617423 |
| sp\|Q8WV99\| | 450437.134 | 623927.907 | 0.23451612 | 0.06981274 | 0.14617423 |
| sp\|Q9HCM4\| | 648448.458 | 851342.94 | 0.23825318 | 0.06981274 | 0.14617423 |
| sp\|Q96CN7\| | 138947982 | 122980433 | -0.1945453 | 0.06981274 | 0.14617423 |
| sp\|Q9BW71\| | 6685380.53 | 13641686.3 | 0.5838306 | 0.06981274 | 0.14617423 |
| sp\|P49354\|F | 22962112.1 | 28172329.3 | 0.25038701 | 0.06981274 | 0.14617423 |
| sp\|P47712\|P | 8335200.88 | 10134098.5 | 0.26644312 | 0.06981274 | 0.14617423 |
| sp\|Q9NZJ9\|N | 13178350.7 | 9749984.97 | -0.3230977 | 0.06981274 | 0.14617423 |
| sp\|Q9Y697\|N | 41434731.3 | 37990569.6 | -0.1422929 | 0.06981274 | 0.14617423 |
| sp\|Q96HY6\|D | 3338054.56 | 3993185.32 | 0.27315062 | 0.06981274 | 0.14617423 |
| sp\|P17612\|K | 48235292.4 | 39625820.8 | -0.3953657 | 0.06981274 | 0.14617423 |
| sp\|P20701\|IT | 2152946.81 | 2308517.83 | 0.29867054 | 0.06981274 | 0.14617423 |
| sp\|P02768\|A | 2.95E+11 | 3.61E+11 | 0.32885973 | 0.06981274 | 0.14617423 |
| sp\|P49755\|T | 9444663.69 | 8291050.54 | -0.2250438 | 0.06981274 | 0.14617423 |
| sp\|O94813\|S | 282662.178 | 544796.136 | 0.4835177 | 0.06981274 | 0.14617423 |
| sp\|Q99733\|N | 43003369 | 37321585 | -0.2908092 | 0.06981274 | 0.14617423 |
| sp\|Q93052\|L | 278392526 | 324008200 | 0.16981906 | 0.06981274 | 0.14617423 |
| sp\|P38159\|R | 89860390.9 | 78994700.1 | -0.2368637 | 0.06981274 | 0.14617423 |
| sp\|Q8WYP3\| | 3419422.24 | 3095397.62 | -1.8310323 | 0.06981274 | 0.14617423 |
| sp\|Q8WZ42\|T | 4424485.52 | 3460428.33 | -0.49925 | 0.06981274 | 0.14617423 |
| sp\|Q92839\|H | 7572958.31 | 6538160.1 | -0.3006253 | 0.06981274 | 0.14617423 |
| sp\|Q969X1\|L | 867844.876 | 1139556.28 | 0.27057259 | 0.06981274 | 0.14617423 |
| sp\|P26992\|C | 1366609.06 | 1587500.92 | 0.30314626 | 0.06981274 | 0.14617423 |
| sp\|P05546\|H | 286968901 | 231247567 | -0.683928 | 0.06981274 | 0.14617423 |
| sp\|O00422\|S | 3560831.88 | 2940635.35 | -0.3455856 | 0.06981274 | 0.14617423 |
| sp\|Q9UGT4\|S | 3481361.02 | 3077283.66 | -0.2534027 | 0.06981274 | 0.14617423 |
| sp\|P50479\|P | 26453903.7 | 22212198.4 | -0.2824908 | 0.06981274 | 0.14617423 |
| sp\|Q86VP6\|C | 2161307.47 | 1533652.07 | -0.4589738 | 0.06981274 | 0.14617423 |
| sp\|Q9NSC5\| | 961020.183 | 773095.48 | -0.3311956 | 0.07353667 | 0.15217874 |
| sp\|Q32MZ4\|L | 17240965.5 | 21845178 | 0.24535216 | 0.07353667 | 0.15217874 |
| sp\|Q13126\|N | 248349280 | 277595729 | 0.1528152 | 0.07353667 | 0.15217874 |
| sp\|Q92552\|F | 455007.994 | 286156.77 | -0.6480103 | 0.07353667 | 0.15217874 |
| sp\|Q8TE49\|C | 1520009.57 | 2273926.22 | 0.39818209 | 0.07353667 | 0.15217874 |
| sp\|Q9Y680\|F | 14024352.4 | 12540003.9 | -0.2463804 | 0.07353667 | 0.15217874 |
| sp\|Q9Y6N7\|F | 1455940.61 | 1271493.97 | -0.2913241 | 0.07353667 | 0.15217874 |
| sp\|P22894\|M | 10227097.9 | 34073544 | 0.65806069 | 0.07353667 | 0.15217874 |

| | | | | | |
|---|---|---|---|---|---|
| sp\|P49589\|S | 62623837.8 | 56582425.8 | -0.1805799 | 0.07353667 | 0.15217874 |
| sp\|O60879\|D | 35953.9827 | 58390.4394 | 4.4480697 | 0.07353667 | 0.15217874 |
| sp\|Q9H0R4\|I | 75826013.7 | 89671495.8 | 0.2090257 | 0.07353667 | 0.15217874 |
| sp\|P16455\|M | 3262064.1 | 3985526.43 | 0.20343066 | 0.07353667 | 0.15217874 |
| sp\|Q99536\|V | 305979387 | 371949790 | 0.21754501 | 0.07353667 | 0.15217874 |
| sp\|Q9UKU9\|A | 313408.892 | 453778.105 | 0.46638979 | 0.07353667 | 0.15217874 |
| sp\|Q02750\|M | 8235359.76 | 12373324.6 | 0.3832468 | 0.07353667 | 0.15217874 |
| sp\|O60716\|C | 28950271.3 | 33182570.8 | 0.15957671 | 0.07353667 | 0.15217874 |
| sp\|Q6PII3\|CC | 1827034.74 | 2538395.24 | 0.35209534 | 0.07353667 | 0.15217874 |
| sp\|Q99627\|C | 7213063.99 | 8579461.36 | 0.21148323 | 0.07353667 | 0.15217874 |
| sp\|P12830\|C | 4488816.72 | 6609913.91 | 0.45529429 | 0.07353667 | 0.15217874 |
| sp\|Q96SI9\|ST | 9137576.1 | 7461286.58 | -0.3128501 | 0.07353667 | 0.15217874 |
| sp\|Q9H0N0\| | 1873997.6 | 1546418.25 | -0.2888906 | 0.07353667 | 0.15217874 |
| sp\|P00740\|F | 13966469.7 | 19701577.5 | 0.39341235 | 0.07353667 | 0.15217874 |
| sp\|Q86VP3\|F | 399835.791 | 477332.367 | 0.30465518 | 0.07353667 | 0.15217874 |
| sp\|Q15155\|N | 20845503.6 | 18500425.5 | -0.1764337 | 0.07353667 | 0.15217874 |
| sp\|P61601\|N | 1060121.52 | 868333.329 | -0.4803774 | 0.07353667 | 0.15217874 |
| sp\|O14907\|T | 6439226.4 | 8885676.78 | 0.34681042 | 0.07353667 | 0.15217874 |
| sp\|Q13228\|S | 6585271436 | 6002117244 | -0.293557 | 0.07353667 | 0.15217874 |
| sp\|P09936\|U | 388299934 | 497701213 | 0.27679421 | 0.0774157 | 0.1578241 |
| sp\|Q86SZ2\|T | 9209773.31 | 13249522.7 | 0.6679163 | 0.0774157 | 0.1578241 |
| sp\|O95479\|G | 223650938 | 186469502 | -0.4868508 | 0.0774157 | 0.1578241 |
| sp\|Q8IVM0\|C | 45901440 | 51700680.4 | 0.16023907 | 0.0774157 | 0.1578241 |
| sp\|O60256\|K | 55280315.8 | 48944885.4 | -0.2293829 | 0.0774157 | 0.1578241 |
| sp\|P82933\|R | 12544508.9 | 18807070.8 | 0.34220306 | 0.0774157 | 0.1578241 |
| sp\|Q02338\|B | 2095440.32 | 1895561.89 | -0.2044722 | 0.0774157 | 0.1578241 |
| sp\|P01833\|P | 3606037.57 | 4404907.58 | 0.25479727 | 0.0774157 | 0.1578241 |
| sp\|P11441\|U | 6006947.94 | 6854389.91 | 0.24267813 | 0.0774157 | 0.1578241 |
| sp\|Q15286\|F | 113491870 | 123979941 | 0.13203948 | 0.0774157 | 0.1578241 |
| sp\|Q5K4L6\|S | 9031470.34 | 7708100.6 | -0.2456741 | 0.0774157 | 0.1578241 |
| sp\|O95218\|Z | 8298373.78 | 6990726.39 | -0.2806601 | 0.0774157 | 0.1578241 |
| sp\|Q6FI81\|C | 10597888.3 | 11687036.1 | 0.13632938 | 0.0774157 | 0.1578241 |
| sp\|Q8WWM9 | 1972602.53 | 2573451.58 | 0.27836703 | 0.0774157 | 0.1578241 |
| sp\|O15042\|S | 10399031.1 | 8431651.65 | -0.400645 | 0.0774157 | 0.1578241 |
| sp\|Q92738\|U | 1098450.37 | 1901904.98 | 0.65557161 | 0.0774157 | 0.1578241 |
| sp\|Q96DG6\| | 77841477.1 | 69801843.5 | -0.1585129 | 0.0774157 | 0.1578241 |
| sp\|Q8TD46\|M | 348047.047 | 480492.626 | 0.35083091 | 0.0774157 | 0.1578241 |
| sp\|P02763\|A | 1024144276 | 1327101172 | 0.3813913 | 0.0774157 | 0.1578241 |
| sp\|Q92609\|T | 5621843.95 | 6978339.59 | 0.31148638 | 0.0774157 | 0.1578241 |

| | | | | | |
|---|---|---|---|---|---|
| sp\|O15027\|S | 16276189.2 | 19085577.9 | 0.18626232 | 0.0774157 | 0.1578241 |
| sp\|Q9BRQ8\| | 23152346.4 | 27639924.2 | 0.17761631 | 0.0774157 | 0.1578241 |
| sp\|Q13443\|A | 269484.489 | 209114.68 | -0.9162849 | 0.0774157 | 0.1578241 |
| sp\|Q99598\|T | 68001440.3 | 61805526.4 | -0.1674916 | 0.0774157 | 0.1578241 |
| sp\|Q06323\|F | 21162240.1 | 28563209.8 | 0.26536409 | 0.0774157 | 0.1578241 |
| sp\|Q8IYT4\|KA | 231813476 | 253419953 | 0.13511237 | 0.0774157 | 0.1578241 |
| sp\|Q9Y296\|T | 3803394.09 | 4267479.92 | 0.1389663 | 0.0774157 | 0.1578241 |
| sp\|P51636\|C | 2230339.6 | 2612471.05 | 0.16724566 | 0.0774157 | 0.1578241 |
| sp\|Q9NRN7\| | 2663802.65 | 2216709.44 | -0.2806508 | 0.0774157 | 0.1578241 |
| sp\|Q9NPQ8\| | 940137.308 | 1446580.01 | 0.46063461 | 0.0774157 | 0.1578241 |
| sp\|Q9H061\|T | 935715.506 | 1344775.7 | 0.30877 | 0.0774157 | 0.1578241 |
| sp\|Q9BV73\|C | 4623134.94 | 5664430 | 0.26224963 | 0.0774157 | 0.1578241 |
| sp\|Q9Y4L1\|H | 101258965 | 88591576.3 | -0.2242045 | 0.0774157 | 0.1578241 |
| sp\|Q96A33\|C | 19975869.7 | 24346955.8 | 0.22070462 | 0.0774157 | 0.1578241 |
| sp\|Q9Y6N5\|S | 21224962.2 | 24204735.9 | 0.14922865 | 0.0774157 | 0.1578241 |
| sp\|Q9BWD1\| | 70506643.9 | 78081699.4 | 0.15188713 | 0.08145395 | 0.16369242 |
| sp\|Q14192\|F | 26383006.7 | 24062018.8 | -0.1528771 | 0.08145395 | 0.16369242 |
| sp\|P55809\|S | 454645250 | 561117763 | 0.22832031 | 0.08145395 | 0.16369242 |
| sp\|Q5JS13\|R | 3209201.29 | 4381814.93 | 0.57947615 | 0.08145395 | 0.16369242 |
| sp\|Q9HCL2\|( | 19904020.8 | 17490229.9 | -0.2398173 | 0.08145395 | 0.16369242 |
| sp\|Q9H8M7\| | 5360231.52 | 6791516.51 | 0.30595642 | 0.08145395 | 0.16369242 |
| sp\|Q8TC07\|T | 2079828.02 | 5449926.07 | 1.27017085 | 0.08145395 | 0.16369242 |
| sp\|Q00688\|F | 38793799.3 | 45896571.1 | 0.20335023 | 0.08145395 | 0.16369242 |
| sp\|O96000\|N | 3126717.05 | 2847892.67 | -0.3778525 | 0.08145395 | 0.16369242 |
| sp\|O43676\|N | 4705465.52 | 3916882.4 | -0.2701152 | 0.08145395 | 0.16369242 |
| sp\|O00115\|D | 22602331.9 | 25745564.3 | 0.14435812 | 0.08145395 | 0.16369242 |
| sp\|Q53FA7\|C | 32017003.3 | 25678252.7 | -0.361096 | 0.08145395 | 0.16369242 |
| sp\|O00763\|A | 7694328.79 | 6898962.22 | -0.2456638 | 0.08145395 | 0.16369242 |
| sp\|P54652\|H | 1567748062 | 1712169251 | 0.15487301 | 0.08145395 | 0.16369242 |
| sp\|Q9H6V9\|I | 741070.323 | 958638.81 | 0.12409031 | 0.08145395 | 0.16369242 |
| sp\|P35613\|B | 77878303.7 | 86480768.2 | 0.17733659 | 0.08145395 | 0.16369242 |
| sp\|Q8TAF3\|V | 204057.812 | 417456.385 | 0.54900332 | 0.08145395 | 0.16369242 |
| sp\|Q9UHL4\|I | 85578973.5 | 105919566 | 0.16024673 | 0.08145395 | 0.16369242 |
| sp\|Q9BY67\|C | 381489.998 | 560735.105 | 0.32725485 | 0.08145395 | 0.16369242 |
| sp\|P16520\|G | 75759394.2 | 67675460.7 | -0.1535383 | 0.08145395 | 0.16369242 |
| sp\|Q8N6C5\| | 2665133.35 | 3575022.09 | 0.31638808 | 0.08145395 | 0.16369242 |
| sp\|P21695\|G | 7057480892 | 8644281967 | 0.30519273 | 0.08145395 | 0.16369242 |
| sp\|P46063\|R | 6370672.96 | 5660864.51 | -0.2284039 | 0.08145395 | 0.16369242 |
| sp\|Q9NRS6\|S | 435674.135 | 593436.88 | 0.22606844 | 0.08145395 | 0.16369242 |

| | | | | | |
|---|---|---|---|---|---|
| sp\|Q9BRG1\|' | 937279.631 | 1244390.75 | 0.25722173 | 0.08145395 | 0.16369242 |
| sp\|O75534\|( | 56541361.9 | 45074788.2 | -0.3290582 | 0.08145395 | 0.16369242 |
| sp\|Q9NZD4\|/ | 22058784.1 | 38701001.4 | 0.62655854 | 0.08145395 | 0.16369242 |
| sp\|O14908\|( | 4864804.14 | 6778369.8 | 0.21997289 | 0.08145395 | 0.16369242 |
| sp\|P19532\|T | 315228.763 | 608259.679 | 2.29592323 | 0.08145395 | 0.16369242 |
| sp\|P07902\|G | 45499352.1 | 44385723.3 | -0.1216343 | 0.08145395 | 0.16369242 |
| sp\|P19652\|A | 1106714500 | 1475577587 | 0.396886 | 0.08145395 | 0.16369242 |
| sp\|Q12907\|L | 17877937.4 | 15702347.4 | -0.2487203 | 0.08145395 | 0.16369242 |
| sp\|Q13033\|S | 37964581.1 | 41090693.1 | 0.10408208 | 0.08145395 | 0.16369242 |
| sp\|Q7Z2W4\|: | 122968509 | 109198626 | -0.202213 | 0.08145395 | 0.16369242 |
| sp\|Q96H20\|S | 3137117.24 | 4166991.82 | 0.23541398 | 0.08565543 | 0.16957937 |
| sp\|P18084\|IT | 8632672.25 | 11050413.3 | 0.27407415 | 0.08565543 | 0.16957937 |
| sp\|P49913\|C | 2834244.48 | 23543363.3 | 1.13374102 | 0.08565543 | 0.16957937 |
| sp\|O14498\|I: | 14947781.5 | 12961884.9 | -0.3213357 | 0.08565543 | 0.16957937 |
| sp\|O94811\|T | 37090487.2 | 28283520.1 | -0.386567 | 0.08565543 | 0.16957937 |
| sp\|Q09028\|F | 30673918.7 | 26598532.9 | -0.280385 | 0.08565543 | 0.16957937 |
| sp\|Q0VDG4\| | 27451236.2 | 33380472.1 | 0.18592326 | 0.08565543 | 0.16957937 |
| sp\|Q86VE3\|S | 60446618.2 | 70502913.1 | 0.24414221 | 0.08565543 | 0.16957937 |
| sp\|Q8N436\|( | 868063.816 | 667906.657 | -0.3170821 | 0.08565543 | 0.16957937 |
| sp\|P16118\|F | 5274490.56 | 6124197.93 | 0.24702967 | 0.08565543 | 0.16957937 |
| sp\|Q4G176\|/ | 7562749.56 | 8947946.46 | 0.24265888 | 0.08565543 | 0.16957937 |
| sp\|P22694\|K | 40566593.1 | 35290909.5 | -0.3062262 | 0.08565543 | 0.16957937 |
| sp\|P22102\|P | 140694342 | 115561333 | -0.3870459 | 0.08565543 | 0.16957937 |
| sp\|Q93045\|S | 49539195.1 | 66092916.1 | 0.30529954 | 0.08565543 | 0.16957937 |
| sp\|O15514\|F | 1054984.07 | 1317361.08 | 0.33726165 | 0.08565543 | 0.16957937 |
| sp\|Q9NTJ5\|S | 20392460.6 | 17745794.3 | -0.3131966 | 0.08565543 | 0.16957937 |
| sp\|P08519\|A | 15276053.7 | 22036699 | 0.28519907 | 0.08565543 | 0.16957937 |
| sp\|P11678\|P | 17355097.4 | 21375076.4 | -0.2405372 | 0.08565543 | 0.16957937 |
| sp\|P11766\|A | 971864962 | 1364309171 | 0.2270118 | 0.08565543 | 0.16957937 |
| sp\|Q9GZT3\|S | 1452216.46 | 1262924.03 | -0.2569462 | 0.08565543 | 0.16957937 |
| sp\|Q8IXM3\|F | 4380822.76 | 3618059.61 | -0.364025 | 0.08565543 | 0.16957937 |
| sp\|O00602\|F | 895294.138 | 1467128.46 | 0.48392145 | 0.08565543 | 0.16957937 |
| sp\|Q01995\|T | 1323151905 | 910007100 | -0.3833114 | 0.08565543 | 0.16957937 |
| sp\|Q9UEE9\|( | 2625400.6 | 2149891.05 | -0.3622564 | 0.08565543 | 0.16957937 |
| sp\|Q5T011\|S | 8615415.26 | 7605893.46 | -0.4017536 | 0.08565543 | 0.16957937 |
| sp\|P10415\|B | 538432.931 | 588801.789 | 0.2454893 | 0.08565543 | 0.16957937 |
| sp\|O60237\|N | 28579278.1 | 24838645.6 | -0.2475154 | 0.08565543 | 0.16957937 |
| sp\|Q9H7M9\| | 781220.994 | 1006165.13 | 0.28045535 | 0.08565543 | 0.16957937 |
| sp\|Q9Y365\|S | 3008592.3 | 5184222.28 | 0.45698488 | 0.08565543 | 0.16957937 |

| | | | | | |
|---|---|---|---|---|---|
| sp\|Q9Y4E1\|V | 18046310 | 20581201.2 | 0.17434763 | 0.08565543 | 0.16957937 |
| sp\|Q00013\|E | 13058283 | 21422504.2 | 0.42347231 | 0.08565543 | 0.16957937 |
| sp\|O43516\|V | 12439675.6 | 19085700 | 0.3609792 | 0.08565543 | 0.16957937 |
| sp\|A6NC98\|C | 6013862.61 | 10721519 | 0.61101446 | 0.08565543 | 0.16957937 |
| sp\|P61970\|N | 196721652 | 174941980 | -0.2470657 | 0.08565543 | 0.16957937 |
| sp\|P29966\|M | 24038237.3 | 28309004.2 | 0.17062647 | 0.08565543 | 0.16957937 |
| sp\|Q99807\|C | 769768.432 | 988879.911 | 0.33199604 | 0.08565543 | 0.16957937 |
| sp\|O14617\|A | 21668906.5 | 19073833.2 | -0.2062085 | 0.09002415 | 0.17619327 |
| sp\|Q15654\|T | 26516170.4 | 30796144.4 | 0.18058527 | 0.09002415 | 0.17619327 |
| sp\|P49908\|S | 22724148.4 | 27404762.9 | 0.30600149 | 0.09002415 | 0.17619327 |
| sp\|P20645\|M | 3073979.62 | 2576904.78 | -0.3815986 | 0.09002415 | 0.17619327 |
| sp\|Q8NBJ5\|C | 61090047.4 | 50626886.8 | -0.3595346 | 0.09002415 | 0.17619327 |
| sp\|P07858\|C | 278640922 | 373907488 | 0.22152078 | 0.09002415 | 0.17619327 |
| sp\|Q9UJ72\|A | 1939594.08 | 2668935.05 | 0.42982996 | 0.09002415 | 0.17619327 |
| sp\|Q5TDH0\|I | 57079839.3 | 92112612.8 | 0.44723075 | 0.09002415 | 0.17619327 |
| sp\|Q9NUP9\| | 15066094.7 | 17455549.5 | 0.16601051 | 0.09002415 | 0.17619327 |
| sp\|Q9Y4W6\|. | 744942.395 | 554524.609 | -0.4875649 | 0.09002415 | 0.17619327 |
| sp\|O60936\|N | 406541875 | 435343496 | 0.10437439 | 0.09002415 | 0.17619327 |
| sp\|Q9HCU5\| | 1302341.92 | 1150464.71 | -0.357498 | 0.09002415 | 0.17619327 |
| sp\|Q30154\|D | 10946978.7 | 14653844.5 | 0.21528487 | 0.09002415 | 0.17619327 |
| sp\|Q9BT92\|T | 3564729.12 | 5156232.85 | 0.47003696 | 0.09002415 | 0.17619327 |
| sp\|Q86V21\|A | 15580144.3 | 19831467.7 | 0.19100646 | 0.09002415 | 0.17619327 |
| sp\|Q9HAT2\|S | 44690612.2 | 53531180.4 | 0.1586846 | 0.09002415 | 0.17619327 |
| sp\|Q01082\|S | 412682585 | 456746807 | 0.13797256 | 0.09002415 | 0.17619327 |
| sp\|Q15149\|F | 388104695 | 437091301 | 0.15791397 | 0.09002415 | 0.17619327 |
| sp\|Q9UBE0\|S | 39096788.7 | 34964857.2 | -0.2087187 | 0.09002415 | 0.17619327 |
| sp\|Q96PE2\|A | 457451.001 | 362216.093 | -0.5551815 | 0.09002415 | 0.17619327 |
| sp\|Q9H201\|E | 175966.219 | 237705.059 | 0.26840909 | 0.09002415 | 0.17619327 |
| sp\|Q05469\|L | 293394077 | 355133463 | 0.19528591 | 0.09002415 | 0.17619327 |
| sp\|Q16576\|F | 24078228.1 | 21639050.6 | -0.2031537 | 0.09002415 | 0.17619327 |
| sp\|Q96FN9\|D | 4138714.37 | 6215607.37 | 0.45435202 | 0.09002415 | 0.17619327 |
| sp\|O15031\|F | 48310921.5 | 41620536.5 | -0.2533693 | 0.09002415 | 0.17619327 |
| sp\|Q12797\|A | 182675305 | 145332638 | -0.5065704 | 0.09002415 | 0.17619327 |
| sp\|P63220\|R | 103346558 | 88504492.1 | -0.2731991 | 0.09002415 | 0.17619327 |
| sp\|Q9Y5Y2\|N | 26977787.6 | 26530766.4 | -0.2142337 | 0.09002415 | 0.17619327 |
| sp\|Q8IWW8\| | 67310384.8 | 76528048.3 | 0.09134706 | 0.0945642 | 0.18343314 |
| sp\|O95747\|C | 10529440.5 | 12907935.7 | 0.19642636 | 0.0945642 | 0.18343314 |
| sp\|Q9H5X1\|C | 3626777.61 | 3112946.34 | -0.2638745 | 0.0945642 | 0.18343314 |
| sp\|Q6AW86\| | 19297396.9 | 16350856.1 | -0.3318131 | 0.0945642 | 0.18343314 |

| | | | | | |
|---|---|---|---|---|---|
| sp\|Q96P67\|( | 2117159.03 | 2636114.8 | 0.16979478 | 0.0945642 | 0.18343314 |
| sp\|Q9NYK5\|I | 10329304 | 12611741.1 | 0.13176373 | 0.0945642 | 0.18343314 |
| sp\|Q99747\|S | 5261381.23 | 6238877.92 | 0.08501323 | 0.0945642 | 0.18343314 |
| sp\|Q9UQN3\| | 12371695.1 | 15099424.2 | 0.20444145 | 0.0945642 | 0.18343314 |
| sp\|P00736\|C | 354895743 | 301928307 | -0.3779785 | 0.0945642 | 0.18343314 |
| sp\|P21506\|Z | 1004116.36 | 1302339.01 | 0.30948996 | 0.0945642 | 0.18343314 |
| sp\|P04083\|A | 724114485 | 940099292 | 0.28612757 | 0.0945642 | 0.18343314 |
| sp\|P46459\|N | 28638057.7 | 31886652.8 | 0.15168206 | 0.0945642 | 0.18343314 |
| sp\|Q9Y5S1\|T | 211438.461 | 455573.371 | -0.1965151 | 0.0945642 | 0.18343314 |
| sp\|P02775\|C | 80514161.8 | 128460968 | 0.51792926 | 0.0945642 | 0.18343314 |
| sp\|A0A0U1RI | 1248912.87 | 1428240.24 | 0.39922071 | 0.0945642 | 0.18343314 |
| sp\|Q8WUI4\|I | 7561848.58 | 10093775.1 | 0.31168997 | 0.0945642 | 0.18343314 |
| sp\|O96007\|N | 2909633.35 | 3351294.84 | 0.23225782 | 0.0945642 | 0.18343314 |
| sp\|Q9BTZ2\|D | 9153299.62 | 15438186.2 | 0.46628407 | 0.0945642 | 0.18343314 |
| sp\|P27348\|1 | 1299212753 | 1166517347 | -0.1439344 | 0.0945642 | 0.18343314 |
| sp\|O14497\|A | 432302.675 | 334793.287 | -0.4727274 | 0.0945642 | 0.18343314 |
| sp\|Q9Y3D2\|I | 9533828.84 | 7615245.81 | -0.3525644 | 0.0945642 | 0.18343314 |
| sp\|Q96EP5\|D | 38083287 | 35171682.8 | -0.1260171 | 0.0945642 | 0.18343314 |
| sp\|P10515\|C | 96655120.7 | 82902022.2 | -0.2695202 | 0.09927951 | 0.19034853 |
| sp\|Q8NGJ5\|( | 4506966.85 | 2420097.72 | -0.6263948 | 0.09927951 | 0.19034853 |
| sp\|Q8WWP7 | 22106946.3 | 20624802.1 | -0.3853055 | 0.09927951 | 0.19034853 |
| sp\|O43324\|N | 2918458.22 | 2471502.52 | -0.3006524 | 0.09927951 | 0.19034853 |
| sp\|P14621\|A | 58993323.6 | 72886193.1 | 0.22408809 | 0.09927951 | 0.19034853 |
| sp\|O94851\|N | 450238.157 | 676056.617 | 0.51991612 | 0.09927951 | 0.19034853 |
| sp\|P51648\|A | 372356948 | 347674092 | -0.1136109 | 0.09927951 | 0.19034853 |
| sp\|Q8NEB9\|I | 4676316.93 | 4239993.79 | -0.1710384 | 0.09927951 | 0.19034853 |
| sp\|P24310\|C | 1460754.64 | 2137364.05 | 0.33365462 | 0.09927951 | 0.19034853 |
| sp\|P35612\|A | 54721771.9 | 94004773.4 | 0.42141582 | 0.09927951 | 0.19034853 |
| sp\|Q9Y2Q9\|I | 1529849.73 | 1037794.72 | -0.5800935 | 0.09927951 | 0.19034853 |
| sp\|Q92804\|F | 18470318.3 | 22752265.5 | 0.23036906 | 0.09927951 | 0.19034853 |
| sp\|Q8NBS9\|T | 432896937 | 505171322 | 0.17680556 | 0.09927951 | 0.19034853 |
| sp\|P46379\|B | 535836.589 | 674361.668 | 0.26309023 | 0.09927951 | 0.19034853 |
| sp\|P67809\|Y | 42239691.4 | 50574441.6 | 0.19303242 | 0.09927951 | 0.19034853 |
| sp\|P48735\|II | 570753580 | 520474912 | -0.2899377 | 0.09927951 | 0.19034853 |
| sp\|O15173\|F | 41545874.9 | 48368217.5 | 0.1504478 | 0.09927951 | 0.19034853 |
| sp\|Q6IBS0\|T | 90089402.8 | 78739940.8 | -0.2565694 | 0.09927951 | 0.19034853 |
| sp\|Q96PM5\|: | 1417187.92 | 1572716.54 | 0.16067676 | 0.09927951 | 0.19034853 |
| sp\|P98082\|D | 30313701 | 27660621.5 | -0.1616985 | 0.09927951 | 0.19034853 |
| sp\|P06681\|C | 139315160 | 117351660 | -0.522122 | 0.09927951 | 0.19034853 |

| | | | | | |
|---|---|---|---|---|---|
| sp\|Q14376\|( | 123161771 | 111102333 | -0.2204337 | 0.09927951 | 0.19034853 |
| sp\|Q9H009\|I | 11803372.7 | 14370952.8 | 0.17898862 | 0.09927951 | 0.19034853 |
| sp\|Q9H019\|I | 1160112.35 | 1047356.92 | -0.2373429 | 0.09927951 | 0.19034853 |
| sp\|Q15166\|F | 18231575.4 | 21592957.8 | 0.2237602 | 0.09927951 | 0.19034853 |
| sp\|Q9UBR2\|( | 136019811 | 172385507 | 0.1876639 | 0.09927951 | 0.19034853 |
| sp\|Q9BXS5\|/ | 3787856.35 | 3373741.69 | -0.3171359 | 0.09927951 | 0.19034853 |
| sp\|Q8N699\|I | 7291414.71 | 8780401.48 | 0.36002043 | 0.09927951 | 0.19034853 |
| sp\|Q7Z422\|S | 5265179.73 | 6927258.58 | 0.33467674 | 0.09927951 | 0.19034853 |
| sp\|O75688\|F | 50458258.5 | 58033763.7 | 0.07948134 | 0.10417399 | 0.19728925 |
| sp\|O43318\|N | 1692080.17 | 2238208.86 | 0.30875384 | 0.10417399 | 0.19728925 |
| sp\|O95171\|S | 726862.082 | 2215416.96 | 0.65088424 | 0.10417399 | 0.19728925 |
| sp\|Q8WXA9\| | 677610.892 | 491349.341 | -1.6174798 | 0.10417399 | 0.19728925 |
| sp\|Q07283\|T | 4337010.13 | 5633237.91 | 0.3049095 | 0.10417399 | 0.19728925 |
| sp\|P23921\|R | 1497994.58 | 2519772.78 | 0.64160089 | 0.10417399 | 0.19728925 |
| sp\|Q9NY33\|I | 151534224 | 175270262 | 0.12745075 | 0.10417399 | 0.19728925 |
| sp\|O00487\|F | 21377723.5 | 23890931.9 | 0.15145778 | 0.10417399 | 0.19728925 |
| sp\|O14974\|N | 61131225.1 | 56051743.9 | -0.1373924 | 0.10417399 | 0.19728925 |
| sp\|Q16563\|S | 16044391.4 | 14571087.2 | -0.2132297 | 0.10417399 | 0.19728925 |
| sp\|Q9NRG0\| | 2421885.96 | 4511070.58 | -2.9764988 | 0.10346928 | 0.19728925 |
| sp\|P63098\|C | 390159.55 | 498068.599 | 0.42748405 | 0.10417399 | 0.19728925 |
| sp\|Q92888\|/ | 382113.96 | 744981.868 | 0.51287381 | 0.10417399 | 0.19728925 |
| sp\|Q8TER5\|/ | 126392.292 | 200380.5 | 0.64390249 | 0.10346928 | 0.19728925 |
| sp\|Q9Y6H1\|( | 4117659.54 | 5075865.52 | 0.24432499 | 0.10417399 | 0.19728925 |
| sp\|P08779\|K | 333048803 | 137097167 | -0.6407512 | 0.10417399 | 0.19728925 |
| sp\|Q9NW64\| | 765143.496 | 1052683.66 | 2.24216322 | 0.10417399 | 0.19728925 |
| sp\|Q16774\|K | 44757085 | 38866917 | -0.1975383 | 0.10417399 | 0.19728925 |
| sp\|Q9Y6K0\|C | 11940401.6 | 10544202.8 | -0.2307438 | 0.10417399 | 0.19728925 |
| sp\|Q9NRX4\|I | 195164747 | 225633709 | 0.15738605 | 0.10417399 | 0.19728925 |
| sp\|O94769\|E | 7332262.19 | 5836492.54 | -0.3633212 | 0.10417399 | 0.19728925 |
| sp\|Q641Q2\|V | 20178632.9 | 22508458.7 | 0.15822902 | 0.10417399 | 0.19728925 |
| sp\|Q9UHK6\|/ | 16647625.6 | 20887882 | 0.27159695 | 0.10417399 | 0.19728925 |
| sp\|Q14240\|I | 25927532 | 22570162.7 | -0.269473 | 0.10417399 | 0.19728925 |
| sp\|Q8WU79\| | 9598798.05 | 8320961.13 | -0.2324765 | 0.10417399 | 0.19728925 |
| sp\|Q8WUN7\| | 2281519.89 | 3294648.56 | 0.61544868 | 0.10417399 | 0.19728925 |
| sp\|P09543\|C | 3895216.1 | 4664995.28 | 0.20037695 | 0.10417399 | 0.19728925 |
| sp\|P60002\|E | 883490.779 | 660032.068 | -0.4504468 | 0.10417399 | 0.19728925 |
| sp\|P61018\|R | 78728151.6 | 88205530.7 | 0.19362052 | 0.10417399 | 0.19728925 |
| sp\|P01854\|I( | 1076582.81 | 1658390.34 | 2.26044357 | 0.10417399 | 0.19728925 |
| sp\|Q8N4L2\|F | 1522579.27 | 1815777.79 | 0.15055867 | 0.10417399 | 0.19728925 |

| | | | | | |
|---|---|---|---|---|---|
| sp\|Q16531\|[ | 272492889 | 318131707 | 0.13948525 | 0.1092516 | 0.20496421 |
| sp\|Q9HAB8\|I | 68229049.5 | 62689955.8 | -0.2568967 | 0.1092516 | 0.20496421 |
| sp\|Q96K21\|A | 17447845 | 19470225.1 | 0.14202765 | 0.1092516 | 0.20496421 |
| sp\|O43172\|F | 6497015.37 | 5386700.1 | -0.2582767 | 0.1092516 | 0.20496421 |
| sp\|P22695\|Q | 38762718.3 | 32295672.3 | -0.2040488 | 0.1092516 | 0.20496421 |
| sp\|Q4V328\|C | 12439199.9 | 13260874.2 | 0.11658144 | 0.1092516 | 0.20496421 |
| sp\|Q9Y2R9\|F | 48597678.6 | 41387179.2 | -0.3582663 | 0.1092516 | 0.20496421 |
| sp\|Q99996\|A | 99086144.1 | 120542792 | 0.26730194 | 0.1092516 | 0.20496421 |
| sp\|Q86VS8\|H | 1165696.26 | 1311961.63 | 0.42163715 | 0.1092516 | 0.20496421 |
| sp\|P05204\|H | 21886814.1 | 29800837.4 | 0.36275844 | 0.1092516 | 0.20496421 |
| sp\|P41236\|IF | 11247291.8 | 14299007.1 | 0.27722574 | 0.1092516 | 0.20496421 |
| sp\|Q9NQ79\| | 4992243.42 | 6097140.34 | 0.35436368 | 0.1092516 | 0.20496421 |
| sp\|Q08722\|C | 23951181.3 | 21379398.4 | -0.1878614 | 0.1092516 | 0.20496421 |
| sp\|Q9BXP5\|S | 2036482.28 | 1471107.29 | -0.4028394 | 0.1092516 | 0.20496421 |
| sp\|O00160\|N | 3108672.88 | 2450779.86 | -0.3685058 | 0.1092516 | 0.20496421 |
| sp\|P0C0L4\|C | 4071922415 | 3454632651 | -0.3449141 | 0.1092516 | 0.20496421 |
| sp\|O95302\|F | 49833263.9 | 61203331.5 | 0.21450775 | 0.1092516 | 0.20496421 |
| sp\|P36575\|A | 7437781.97 | 6678453.31 | -0.2002716 | 0.1092516 | 0.20496421 |
| sp\|P31146\|C | 205812464 | 215322946 | -0.3106922 | 0.1092516 | 0.20496421 |
| sp\|A4D126\|IS | 864703.699 | 760703.922 | -0.2342471 | 0.1092516 | 0.20496421 |
| sp\|Q86T13\|C | 4570875.41 | 5317180.42 | 0.22849264 | 0.1092516 | 0.20496421 |
| sp\|P61009\|S | 1639547.32 | 1434506.3 | -0.3477105 | 0.1092516 | 0.20496421 |
| sp\|Q9Y2W6\|T | 45281118.3 | 55943152.4 | 0.37878924 | 0.1092516 | 0.20496421 |
| sp\|O43776\|S | 120530377 | 95960031.2 | -0.4569446 | 0.1092516 | 0.20496421 |
| sp\|Q14678\|K | 5521106.28 | 4934100.9 | -0.1670585 | 0.11451615 | 0.21153311 |
| sp\|O43813\|L | 189059821 | 243948582 | 0.21217247 | 0.11451615 | 0.21153311 |
| sp\|A6NED2\|F | 381286.616 | 443381.211 | 0.26530362 | 0.11451615 | 0.21153311 |
| sp\|Q96D15\|F | 1723684.75 | 2449689.35 | 0.38173659 | 0.11451615 | 0.21153311 |
| sp\|Q9BUE6\|I | 448890.38 | 720892.16 | 0.71270709 | 0.11451615 | 0.21153311 |
| sp\|P0C0L5\|C | 4109923221 | 3470827901 | -0.3511212 | 0.11451615 | 0.21153311 |
| sp\|Q5U5X0\|L | 1301410.2 | 1940937.32 | 0.42001111 | 0.11451615 | 0.21153311 |
| sp\|P61221\|A | 22623040.3 | 20407822.4 | -0.1678335 | 0.11451615 | 0.21153311 |
| sp\|P02749\|A | 163138564 | 242272616 | 0.55133477 | 0.11451615 | 0.21153311 |
| sp\|Q8IWV8\|U | 89394.0776 | 56186.5805 | -2.5391697 | 0.11451615 | 0.21153311 |
| sp\|O95278\|E | 5552381.38 | 6324732.01 | 0.08021562 | 0.11451615 | 0.21153311 |
| sp\|Q99459\|C | 2656292.61 | 2246530.87 | -0.3543716 | 0.11451615 | 0.21153311 |
| sp\|Q13464\|F | 214461.601 | 427949.945 | 0.07921997 | 0.11451615 | 0.21153311 |
| sp\|P35754\|G | 144193376 | 161913182 | 0.12300526 | 0.11451615 | 0.21153311 |
| sp\|O00560\|S | 11451038.1 | 12977245.8 | 0.17945407 | 0.11451615 | 0.21153311 |

| | | | | | |
|---|---|---|---|---|---|
| sp\|Q9UNS2\| | 8349964.34 | 8176029.6 | -0.1885833 | 0.11451615 | 0.21153311 |
| sp\|P40616\|A | 1311807.78 | 1593041.01 | 0.25655991 | 0.11451615 | 0.21153311 |
| sp\|P15153\|R | 114119320 | 143516460 | 0.22179308 | 0.11451615 | 0.21153311 |
| sp\|Q12899\|T | 685186.007 | 516769.305 | -1.2196427 | 0.11451615 | 0.21153311 |
| sp\|P16949\|S | 93874508.2 | 130582174 | 0.34789942 | 0.11451615 | 0.21153311 |
| sp\|Q6P179\|E | 29560462 | 23495585.9 | -0.2637495 | 0.11451615 | 0.21153311 |
| sp\|Q13363\|C | 47128825.5 | 40545386.2 | -0.2279184 | 0.11451615 | 0.21153311 |
| sp\|Q9NPD3\| | 1705057.21 | 1582560.58 | -0.2237953 | 0.11451615 | 0.21153311 |
| sp\|P50897\|P | 15062420.8 | 16960128.6 | 0.17514702 | 0.11451615 | 0.21153311 |
| sp\|Q8NG27\| | 351535.361 | 430926.925 | 0.09278629 | 0.11451615 | 0.21153311 |
| sp\|Q8NHZ8\| | 500296.445 | 756492.87 | 0.21890568 | 0.11451615 | 0.21153311 |
| sp\|O95232\|L | 491312.555 | 455898.001 | -0.4049392 | 0.11451615 | 0.21153311 |
| sp\|P55209\|N | 14613532.7 | 12636814 | -0.2351837 | 0.11451615 | 0.21153311 |
| sp\|Q8N6H7\| | 6981246.59 | 7785450.03 | 0.1315782 | 0.11451615 | 0.21153311 |
| sp\|Q9H3S4\|T | 961905.561 | 1330192.08 | 0.18587218 | 0.11451615 | 0.21153311 |
| sp\|Q9NR45\|S | 109023756 | 98135550.9 | -0.1044492 | 0.11451615 | 0.21153311 |
| sp\|O75879\|C | 320676.732 | 399674.287 | 0.22159459 | 0.11451615 | 0.21153311 |
| sp\|Q9Y2I1\|N | 6086321.76 | 5259825 | -0.3908862 | 0.11451615 | 0.21153311 |
| sp\|Q5VY43\|F | 1518741.58 | 1975468.63 | 0.35900399 | 0.11451615 | 0.21153311 |
| sp\|Q14653\|I | 1291225.56 | 998749.075 | -0.2895647 | 0.11451615 | 0.21153311 |
| sp\|Q7Z7C8\|T | 2019879.46 | 4567203.2 | 0.34630125 | 0.11451615 | 0.21153311 |
| sp\|Q63ZY3\|K | 15916311.6 | 14742034.4 | -0.1495292 | 0.11451615 | 0.21153311 |
| sp\|P08590\|M | 15548227.5 | 40705159.6 | 0.4254595 | 0.11451615 | 0.21153311 |
| sp\|A0A1B0G\ | 823200.196 | 1209928.92 | 0.38766161 | 0.11451615 | 0.21153311 |
| sp\|P82094\|T | 719409.115 | 1022285.57 | 0.2625171 | 0.11451615 | 0.21153311 |
| sp\|Q86Y39\|N | 7648278.36 | 6993108.92 | -0.2352775 | 0.11997137 | 0.21899681 |
| sp\|O14828\|S | 7195047.41 | 8002860.89 | 0.18716919 | 0.11997137 | 0.21899681 |
| sp\|P01031\|C | 279535326 | 237893981 | -0.6472818 | 0.11997137 | 0.21899681 |
| sp\|P12814\|A | 710868017 | 599003127 | -0.3121453 | 0.11997137 | 0.21899681 |
| sp\|P51553\|I | 17027374.1 | 15639334.5 | -0.1577867 | 0.11997137 | 0.21899681 |
| sp\|Q13488\|V | 4348928.15 | 5312574.2 | 0.22033995 | 0.11997137 | 0.21899681 |
| sp\|Q92900\|F | 38514984.2 | 34723039.5 | -0.1643032 | 0.11997137 | 0.21899681 |
| sp\|A0A0C4D | 21556218.5 | 25147651.9 | 0.3823684 | 0.11997137 | 0.21899681 |
| sp\|O60502\|C | 36164131.7 | 32736680.5 | -0.1903686 | 0.11997137 | 0.21899681 |
| sp\|P33121\|A | 292341872 | 246422123 | -0.3955927 | 0.11997137 | 0.21899681 |
| sp\|P02741\|C | 29327827.2 | 23917680 | -0.2213227 | 0.11997137 | 0.21899681 |
| sp\|P02654\|A | 15105431.9 | 22955156 | 0.34317317 | 0.11997137 | 0.21899681 |
| sp\|Q5BKX8\|C | 12622587.1 | 10129825.9 | -0.8177384 | 0.11997137 | 0.21899681 |
| sp\|Q8WWI5\| | 5355093.35 | 4498895.52 | -0.2419428 | 0.11997137 | 0.21899681 |

| | | | | | |
|---|---|---|---|---|---|
| sp\|Q13636\|F | 1292375.82 | 1519818.01 | 0.25605405 | 0.11997137 | 0.21899681 |
| sp\|Q9BRJ2\|R | 6188611.26 | 5182794.75 | -0.4685958 | 0.11997137 | 0.21899681 |
| sp\|Q3YEC7\|I | 1603154.92 | 1794210.76 | 0.14680503 | 0.11997137 | 0.21899681 |
| sp\|P15559\|N | 773080821 | 794636450 | 0.61041345 | 0.11997137 | 0.21899681 |
| sp\|Q07817\|E | 1793490.18 | 3035580.09 | 0.32045994 | 0.11997137 | 0.21899681 |
| sp\|Q06210\|C | 10642853.9 | 12020140 | 0.15879408 | 0.11997137 | 0.21899681 |
| sp\|P09429\|H | 155091212 | 129268556 | -0.413864 | 0.11997137 | 0.21899681 |
| sp\|Q14739\|L | 22397335.6 | 28357512.7 | 0.16291019 | 0.11997137 | 0.21899681 |
| sp\|Q96D96\|I | 666027.542 | 860081.744 | -0.4720106 | 0.11997137 | 0.21899681 |
| sp\|Q96D71\|I | 1025752.24 | 1185110.12 | 0.26986084 | 0.11997137 | 0.21899681 |
| sp\|Q8N4C8\| | 2109856.95 | 2440989.34 | 0.15983468 | 0.11997137 | 0.21899681 |
| sp\|Q99808\|S | 208993.272 | 263331.049 | 0.35846149 | 0.11997137 | 0.21899681 |
| sp\|Q5VWZ2\| | 28335850.5 | 32847057.2 | 0.1941875 | 0.11997137 | 0.21899681 |
| sp\|Q9BRP4\|I | 16089286.1 | 19320949.8 | 0.20645604 | 0.11997137 | 0.21899681 |
| sp\|Q9Y316\|N | 10246288.2 | 9026210.37 | -0.3077009 | 0.11997137 | 0.21899681 |
| sp\|Q9NZZ3\|C | 19707434.4 | 20765457.4 | 0.11296349 | 0.11997137 | 0.21899681 |
| sp\|P56537\|I | 37574700 | 42805081.3 | 0.18843461 | 0.11997137 | 0.21899681 |
| sp\|Q9NVH6\| | 53169361.2 | 49191197.5 | -0.1529482 | 0.12562107 | 0.22655223 |
| sp\|A0A0C4DI | 22467588.9 | 19677012.3 | -0.1668507 | 0.12562107 | 0.22655223 |
| sp\|P35555\|F | 3613975313 | 4655768295 | 0.34439233 | 0.12562107 | 0.22655223 |
| sp\|Q5K130\|C | 3600870.57 | 5995914.24 | 0.41668873 | 0.12562107 | 0.22655223 |
| sp\|Q9NWX6\| | 1961478.56 | 3096099.06 | 0.3693213 | 0.12562107 | 0.22655223 |
| sp\|O95292\|V | 10814336.7 | 9883167.02 | -0.1539929 | 0.12562107 | 0.22655223 |
| sp\|Q02083\|N | 1262440.08 | 998506.027 | -0.3557881 | 0.12562107 | 0.22655223 |
| sp\|Q86YP4\|F | 310525.518 | 456650.397 | 1.03033098 | 0.12562107 | 0.22655223 |
| sp\|Q86YS7\|C | 686622.915 | 500293.858 | -0.444409 | 0.12562107 | 0.22655223 |
| sp\|Q8N983\|I | 25227720.3 | 20480562.4 | -0.4768435 | 0.12562107 | 0.22655223 |
| sp\|Q96HQ2\| | 1637234.65 | 1265146.3 | -0.3093345 | 0.12562107 | 0.22655223 |
| sp\|O60828\|F | 1258064.61 | 1053053.67 | -0.2986779 | 0.12562107 | 0.22655223 |
| sp\|P55265\|D | 2779691.57 | 2277321.4 | -0.3408855 | 0.12562107 | 0.22655223 |
| sp\|Q969U7\|I | 779517.919 | 1019237.73 | 1.5872745 | 0.12562107 | 0.22655223 |
| sp\|P23142\|F | 109987091 | 95275094.1 | -0.3109361 | 0.12562107 | 0.22655223 |
| sp\|P00739\|H | 4825043782 | 6212483002 | 0.24000627 | 0.12562107 | 0.22655223 |
| sp\|P08962\|C | 23172254.1 | 26126031.2 | 0.17663376 | 0.12562107 | 0.22655223 |
| sp\|P35244\|R | 7675245.12 | 6742264.14 | -0.2016887 | 0.12562107 | 0.22655223 |
| sp\|Q8WTU0\| | 7213065.93 | 13383574.5 | 0.46330022 | 0.12562107 | 0.22655223 |
| sp\|O43716\|C | 2698211.17 | 3208084.06 | 0.23437849 | 0.12562107 | 0.22655223 |
| sp\|P62140\|P | 99131208.1 | 90760321.4 | -0.1409029 | 0.12562107 | 0.22655223 |
| sp\|Q9UHJ6\|S | 58027612.9 | 75473652.2 | 0.15759856 | 0.12562107 | 0.22655223 |

| | | | | | |
|---|---|---|---|---|---|
| sp\|P30044\|P | 594855173 | 658846967 | 0.08990884 | 0.12562107 | 0.22655223 |
| sp\|A8MVU1\|I | 1631078.16 | 1998048.66 | -0.0504067 | 0.12562107 | 0.22655223 |
| sp\|P07384\|C | 155428929 | 148821705 | -0.2477763 | 0.12562107 | 0.22655223 |
| sp\|P62873\|G | 174780717 | 189563544 | 0.11037928 | 0.12562107 | 0.22655223 |
| sp\|O75880\|S | 5991531.53 | 7068885.19 | 0.18163347 | 0.12562107 | 0.22655223 |
| sp\|Q8N6T3\|A | 1405087.47 | 1862031.44 | 0.92963686 | 0.12562107 | 0.22655223 |
| sp\|Q8N806\|U | 898376.727 | 731402.824 | -0.478291 | 0.12562107 | 0.22655223 |
| sp\|P78362\|S | 14008411.8 | 17681111.3 | 0.23067002 | 0.12562107 | 0.22655223 |
| sp\|O15230\|L | 155092139 | 167384461 | 0.14651171 | 0.12562107 | 0.22655223 |
| sp\|O60220\|T | 15263262.7 | 15580254.4 | 0.11590994 | 0.12562107 | 0.22655223 |
| sp\|O75934\|S | 942376.952 | 1002113.41 | -0.2234793 | 0.1314688 | 0.2345423 |
| sp\|Q9Y2G5\|( | 17429694.3 | 16056934.9 | -0.1611588 | 0.1314688 | 0.2345423 |
| sp\|P49748\|A | 224714449 | 251945728 | 0.16045519 | 0.1314688 | 0.2345423 |
| sp\|P21796\|V | 73614951.3 | 60833608.3 | -0.2577244 | 0.1314688 | 0.2345423 |
| sp\|P43155\|C | 13189994.9 | 11443129.2 | -0.2301176 | 0.1314688 | 0.2345423 |
| sp\|Q9HD89\| | 7628920.64 | 32387735.6 | 0.8011956 | 0.1314688 | 0.2345423 |
| sp\|Q12988\|H | 28819331.7 | 38312159.5 | 0.61532388 | 0.1314688 | 0.2345423 |
| sp\|Q13287\|N | 11831510.1 | 15072926.7 | 0.20801617 | 0.1314688 | 0.2345423 |
| sp\|P30740\|IL | 290150356 | 276670952 | -0.1498255 | 0.1314688 | 0.2345423 |
| sp\|P98171\|R | 142805.977 | 102949.208 | -1.7534248 | 0.1314688 | 0.2345423 |
| sp\|Q96I51\|R | 9890043.88 | 10505869.9 | 0.12512215 | 0.1314688 | 0.2345423 |
| sp\|A0A0C4DI | 574639394 | 644065768 | 0.20580971 | 0.1314688 | 0.2345423 |
| sp\|Q8IVW4\|( | 244875.016 | 162901.036 | -0.901414 | 0.1314688 | 0.2345423 |
| sp\|Q9BVK6\|T | 8045328.68 | 6690647.43 | -0.4304589 | 0.1314688 | 0.2345423 |
| sp\|Q8TCF1\|Z | 604689.913 | 995215.476 | 0.57893626 | 0.1314688 | 0.2345423 |
| sp\|O14495\|F | 21671602.9 | 18341233.7 | -0.270236 | 0.1314688 | 0.2345423 |
| sp\|P53634\|C | 62287269.3 | 52334005.4 | -0.3977254 | 0.1314688 | 0.2345423 |
| sp\|Q6P4F2\|F | 6293046.45 | 7377369.76 | 0.10475999 | 0.1314688 | 0.2345423 |
| sp\|Q9GZP8\|I | 2157587.53 | 1767682.88 | -0.3365265 | 0.1314688 | 0.2345423 |
| sp\|Q9BXJ9\|N | 1251103.42 | 1073175.83 | -0.322781 | 0.1314688 | 0.2345423 |
| sp\|P11940\|P | 153976093 | 132636025 | -0.3241784 | 0.1314688 | 0.2345423 |
| sp\|P14550\|A | 739604811 | 636454617 | -0.3273733 | 0.1314688 | 0.2345423 |
| sp\|Q8IYB1\|M | 705009.63 | 1026623.79 | 0.34245749 | 0.1314688 | 0.2345423 |
| sp\|Q9UN37\|' | 15564484.9 | 17901471 | 0.17762503 | 0.1314688 | 0.2345423 |
| sp\|Q9H190\|S | 3200099.7 | 3629182.82 | 0.16796233 | 0.1314688 | 0.2345423 |
| sp\|P18065\|IE | 3677539.42 | 3276732 | -0.3301273 | 0.1314688 | 0.2345423 |
| sp\|P17174\|A | 822245862 | 881134999 | 0.14531859 | 0.1314688 | 0.2345423 |
| sp\|Q7Z7L8\|C | 4113200.15 | 4997408.87 | 0.29840162 | 0.1314688 | 0.2345423 |
| sp\|Q7Z7K6\|( | 671756.174 | 516908.001 | -0.4425547 | 0.1314688 | 0.2345423 |

| | | | | | |
|---|---|---|---|---|---|
| sp\|P50990\|T | 114125835 | 102382530 | -0.2632966 | 0.1375181 | 0.24218325 |
| sp\|O76071\|( | 30424305 | 34019321.5 | 0.1123674 | 0.1375181 | 0.24218325 |
| sp\|Q92499\|D | 113402720 | 97587256.4 | -0.2937848 | 0.1375181 | 0.24218325 |
| sp\|Q9H5Y7\|S | 265501.297 | 333261.034 | 0.98222927 | 0.1375181 | 0.24218325 |
| sp\|Q86W42\| | 344412.565 | 314462.786 | -0.2578238 | 0.1375181 | 0.24218325 |
| sp\|P06241\|F | 69811677.1 | 61586845.1 | -0.1697158 | 0.1375181 | 0.24218325 |
| sp\|O60841\|I | 2235050.88 | 3358114.69 | 0.36378817 | 0.1375181 | 0.24218325 |
| sp\|O75439\|N | 54473710.1 | 46436443.9 | -0.3063598 | 0.1375181 | 0.24218325 |
| sp\|Q9BQD3\| | 26174907.2 | 29192056.4 | 0.18418133 | 0.1375181 | 0.24218325 |
| sp\|P40123\|C | 19252429.1 | 23158291.7 | 0.20821471 | 0.1375181 | 0.24218325 |
| sp\|Q04323\|U | 16692428.8 | 18371453.5 | 0.11685397 | 0.1375181 | 0.24218325 |
| sp\|Q96QZ7\|I | 17561875 | 19230043 | 0.03807599 | 0.1375181 | 0.24218325 |
| sp\|P62306\|R | 5235597.8 | 4595844.42 | -0.2623721 | 0.1375181 | 0.24218325 |
| sp\|P31327\|C | 175823680 | 224446025 | 0.23798922 | 0.1375181 | 0.24218325 |
| sp\|Q5VTR2\|E | 1241904.91 | 1968796.9 | 0.66719266 | 0.1375181 | 0.24218325 |
| sp\|Q96DI7\|S | 1261486.54 | 913079.017 | -0.4165245 | 0.1375181 | 0.24218325 |
| sp\|Q9H0D6\|: | 20389637.2 | 17490937 | -0.2343972 | 0.1375181 | 0.24218325 |
| sp\|Q14766\|L | 10048332 | 12865341.5 | 0.26846277 | 0.1375181 | 0.24218325 |
| sp\|Q14831\|( | 2444859.21 | 3069175.44 | 0.35810807 | 0.1375181 | 0.24218325 |
| sp\|P30530\|U | 1377376.35 | 1670875.89 | 0.23985898 | 0.1375181 | 0.24218325 |
| sp\|Q96DX5\|/ | 1424640.55 | 1280539.83 | -0.227967 | 0.1375181 | 0.24218325 |
| sp\|Q96BP3\|F | 2539501.22 | 2241856.67 | -0.2043601 | 0.1375181 | 0.24218325 |
| sp\|Q8IXJ6\|SI | 557650.447 | 453824.363 | -0.445993 | 0.1375181 | 0.24218325 |
| sp\|P55735\|S | 23650526.4 | 28426193.2 | 0.16969946 | 0.1375181 | 0.24218325 |
| sp\|Q86YB7\|E | 16931511.5 | 20992469.8 | 0.18890777 | 0.1375181 | 0.24218325 |
| sp\|Q96PE7\|N | 12322173.6 | 14064104.4 | 0.08859023 | 0.1375181 | 0.24218325 |
| sp\|Q8WUY1\| | 27969174.8 | 40663201.2 | 0.41257242 | 0.1375181 | 0.24218325 |
| sp\|P17540\|K | 92527931.8 | 47288540.6 | -0.5297372 | 0.1375181 | 0.24218325 |
| sp\|Q6ZQY2\|L | 419870.097 | 505766.441 | 0.16787878 | 0.1375181 | 0.24218325 |
| sp\|O60234\|( | 22984135.4 | 26347139.8 | 0.15359886 | 0.1375181 | 0.24218325 |
| sp\|P78318\|I( | 40302308.1 | 54752379.4 | 0.34688712 | 0.1375181 | 0.24218325 |
| sp\|Q7Z7F7\|F | 1191332.43 | 1022090.37 | -0.3525597 | 0.1375181 | 0.24218325 |
| sp\|Q9H2X3\|( | 3376423.44 | 2959944.64 | -0.2754917 | 0.1375181 | 0.24218325 |
| sp\|Q9Y6I9\|T | 491242.07 | 610123.83 | 0.20313984 | 0.1375181 | 0.24218325 |
| sp\|Q9BUR5\|I | 3041468.21 | 3988176.05 | 0.27576768 | 0.1375181 | 0.24218325 |
| sp\|Q9BV68\|F | 1470566.52 | 1668093.89 | 0.13835999 | 0.14377251 | 0.24962527 |
| sp\|Q9HA65\|T | 2336262.03 | 2886230.05 | 0.07606619 | 0.14377251 | 0.24962527 |
| sp\|P63165\|S | 25726012.5 | 23247058.4 | -0.1753569 | 0.14377251 | 0.24962527 |
| sp\|Q16543\|( | 29474316.5 | 26694615.6 | -0.1410668 | 0.14377251 | 0.24962527 |

| | | | | | |
|---|---|---|---|---|---|
| sp\|Q29RF7\|F | 1769229.18 | 2353119.81 | 0.25935557 | 0.14377251 | 0.24962527 |
| sp\|Q9UPA5\|I | 1888387.38 | 2438758.5 | 0.18969484 | 0.14377251 | 0.24962527 |
| sp\|P04921\|G | 640370.764 | 817706.301 | 0.24194058 | 0.14377251 | 0.24962527 |
| sp\|O43157\|F | 3502729.14 | 3127362.55 | -0.2065315 | 0.14377251 | 0.24962527 |
| sp\|Q9NQC3\| | 66512847.7 | 80210885.4 | 0.1478892 | 0.14377251 | 0.24962527 |
| sp\|O75882\|A | 105445085 | 125478861 | 0.26911494 | 0.14377251 | 0.24962527 |
| sp\|Q9NYJ1\|C | 656999.309 | 747001.101 | 0.18635012 | 0.14377251 | 0.24962527 |
| sp\|P16157\|A | 124870374 | 231254905 | 0.40641369 | 0.14377251 | 0.24962527 |
| sp\|P21953\|C | 12308451.7 | 11230022.5 | -0.1820423 | 0.14377251 | 0.24962527 |
| sp\|P00568\|K | 99197897.7 | 99407677.9 | -0.3028176 | 0.14377251 | 0.24962527 |
| sp\|Q7Z401\|N | 4723372.99 | 4165591.3 | -0.2052976 | 0.14377251 | 0.24962527 |
| sp\|Q2TB90\|H | 18081192.4 | 20095111 | 0.1649323 | 0.14377251 | 0.24962527 |
| sp\|P55058\|P | 2032222.05 | 2718710.15 | 0.20859805 | 0.14377251 | 0.24962527 |
| sp\|P54819\|K | 81365515.2 | 86036014.1 | 0.09732557 | 0.14377251 | 0.24962527 |
| sp\|Q9GZT9\|E | 12888582.9 | 11062218 | -0.2169076 | 0.14377251 | 0.24962527 |
| sp\|O75380\|N | 23567668.9 | 25947174.1 | 0.23819772 | 0.14377251 | 0.24962527 |
| sp\|P53609\|P | 28312620.6 | 35383273.7 | 0.20735863 | 0.14377251 | 0.24962527 |
| sp\|Q05481\|Z | 3164508.54 | 3428184.78 | 0.101935 | 0.14377251 | 0.24962527 |
| sp\|Q15257\|F | 70314410.6 | 60988998.8 | -0.5400092 | 0.14377251 | 0.24962527 |
| sp\|P11586\|C | 395589128 | 335440772 | -0.380802 | 0.14377251 | 0.24962527 |
| sp\|Q15036\|S | 2090880.94 | 2435962.19 | 0.20886117 | 0.14377251 | 0.24962527 |
| sp\|Q13952\|N | 17513912.5 | 18601457.9 | 0.0904803 | 0.14377251 | 0.24962527 |
| sp\|P0DP01\|H | 124673347 | 141653826 | 0.35940544 | 0.14377251 | 0.24962527 |
| sp\|Q5VT66\|N | 21759843.8 | 19140948.1 | -0.2108888 | 0.14377251 | 0.24962527 |
| sp\|Q9UL63\|N | 1917474.06 | 3221399.1 | 0.51021162 | 0.14377251 | 0.24962527 |
| sp\|Q7LG56\|F | 5500792.37 | 6323271.73 | 0.15999121 | 0.14377251 | 0.24962527 |
| sp\|Q5T440\|C | 2987319.14 | 3394856.81 | 0.11174251 | 0.14377251 | 0.24962527 |
| sp\|Q86U42\|F | 19652152 | 17754604.5 | -0.162162 | 0.14377251 | 0.24962527 |
| sp\|P61081\|U | 11217458.1 | 14094456.6 | 0.27556952 | 0.14377251 | 0.24962527 |
| sp\|P61962\|D | 5377678.09 | 4974019.67 | -0.1412313 | 0.14377251 | 0.24962527 |
| sp\|Q96GA3\|I | 600929.674 | 1934200.65 | 0.75540802 | 0.14377251 | 0.24962527 |
| sp\|P30419\|N | 13921921.4 | 12681316.1 | -0.156127 | 0.14377251 | 0.24962527 |
| sp\|Q14141\|S | 175572599 | 154822136 | -0.2016011 | 0.14377251 | 0.24962527 |
| sp\|P07738\|P | 704499286 | 1035605918 | 0.30789164 | 0.14377251 | 0.24962527 |
| sp\|O60504\|V | 78384013.6 | 86720219.2 | 0.12000122 | 0.14377251 | 0.24962527 |
| sp\|Q9NWB6\| | 350578.433 | 298992.294 | -2.0010311 | 0.14815501 | 0.25714137 |
| sp\|Q96E11\|F | 284537.643 | 338105.411 | 0.30874178 | 0.15023531 | 0.25804555 |
| sp\|Q9BU40\|( | 1109167.45 | 1288943.49 | 0.16894534 | 0.15023531 | 0.25804555 |
| sp\|Q5T4F7\|S | 737045.353 | 849602.013 | 0.09544923 | 0.15023531 | 0.25804555 |

| | | | | | |
|---|---|---|---|---|---|
| sp\|Q5JTV8\|T( | 4070573.52 | 3548892.09 | -0.3208654 | 0.15023531 | 0.25804555 |
| sp\|P09172\|D | 6587137.07 | 6019598.52 | -0.1730979 | 0.15023531 | 0.25804555 |
| sp\|P32856\|S | 642687.18 | 1042600.51 | 0.51541292 | 0.15023531 | 0.25804555 |
| sp\|Q9ULT8\|H | 4736503.2 | 5917156.3 | 0.42617682 | 0.15023531 | 0.25804555 |
| sp\|Q75N90\|I | 19174022.9 | 26869904.6 | 0.14773735 | 0.15023531 | 0.25804555 |
| sp\|P55061\|B | 913201.599 | 1142792.47 | 0.1455991 | 0.15023531 | 0.25804555 |
| sp\|Q13627\|D | 572050.152 | 1036081.82 | 0.61886625 | 0.15023531 | 0.25804555 |
| sp\|O95159\|Z | 2611260.22 | 2887620.55 | 0.11554379 | 0.15023531 | 0.25804555 |
| sp\|Q9Y3Q8\|T | 2349644.54 | 3552253.86 | 0.21433003 | 0.15023531 | 0.25804555 |
| sp\|Q8NHV4\| | 285557.693 | 354645.365 | 0.22595601 | 0.15023531 | 0.25804555 |
| sp\|Q6H8Q1\| | 15856130.8 | 17110166.7 | 0.08114087 | 0.15023531 | 0.25804555 |
| sp\|Q9H479\|I | 24704057.6 | 27256779.6 | 0.16907565 | 0.15023531 | 0.25804555 |
| sp\|P02787\|T | 3.8363E+10 | 4.5137E+10 | 0.23469054 | 0.15023531 | 0.25804555 |
| sp\|P18433\|P | 2304192.93 | 1949750.43 | -0.3385 | 0.15023531 | 0.25804555 |
| sp\|Q9NR56\|I | 35723094.3 | 39815237.2 | 0.10422352 | 0.15023531 | 0.25804555 |
| sp\|O15178\|T | 4178200.87 | 4855836.06 | 0.36995628 | 0.15023531 | 0.25804555 |
| sp\|Q9P0K7\|F | 8791558.27 | 12512533.6 | 0.24719097 | 0.15023531 | 0.25804555 |
| sp\|Q32M88\|I | 9482306.16 | 7521484.02 | 0.22641629 | 0.15023531 | 0.25804555 |
| sp\|Q92743\|H | 2305853.36 | 2876378.15 | 0.26971964 | 0.15023531 | 0.25804555 |
| sp\|A0A0J9YX: | 78841760.7 | 86556356.5 | 0.21098827 | 0.15023531 | 0.25804555 |
| sp\|Q9BQ04\|I | 27053872.7 | 24949675.9 | -0.1259443 | 0.15023531 | 0.25804555 |
| sp\|O60669\|M | 6573426.53 | 6999452.78 | 0.23447107 | 0.15023531 | 0.25804555 |
| sp\|P30086\|P | 1783395135 | 2155013824 | 0.13333036 | 0.15023531 | 0.25804555 |
| sp\|Q14515\|S | 15086417.1 | 15170157.1 | 0.0606898 | 0.15023531 | 0.25804555 |
| sp\|P06744\|G | 2558155123 | 3275106764 | 0.27771679 | 0.15023531 | 0.25804555 |
| sp\|P32119\|P | 6843452836 | 9516731403 | 0.26057005 | 0.15023531 | 0.25804555 |
| sp\|Q6PFW1\| | 1156318.8 | 1016375.33 | -0.8354616 | 0.15690974 | 0.26570566 |
| sp\|Q96AZ6\|I | 1072431.79 | 623083.448 | 0.30912046 | 0.15690974 | 0.26570566 |
| sp\|O96033\|M | 8142790.4 | 9021396.71 | 0.09119786 | 0.15690974 | 0.26570566 |
| sp\|Q9BQS8\|I | 2707427.48 | 3299708.61 | 0.21561079 | 0.15690974 | 0.26570566 |
| sp\|P55210\|C | 5016723.68 | 6225620.08 | 0.20016635 | 0.15690974 | 0.26570566 |
| sp\|P23919\|K | 14574351.1 | 18695143.7 | 0.38974652 | 0.15690974 | 0.26570566 |
| sp\|Q5EG05\|( | 11521782.4 | 12646631.5 | 0.08352587 | 0.15690974 | 0.26570566 |
| sp\|Q4KMP7\|T | 1092268.38 | 1273804.91 | 0.18661094 | 0.15690974 | 0.26570566 |
| sp\|Q53FT3\|H | 1268812.65 | 2103090.6 | 0.3309412 | 0.15690974 | 0.26570566 |
| sp\|Q9H0U3\| | 13279075.3 | 15454256 | 0.18621407 | 0.15690974 | 0.26570566 |
| sp\|O60934\|N | 227724.313 | 184938.489 | -0.4471726 | 0.15690974 | 0.26570566 |
| sp\|O43148\|N | 24112729.2 | 20771294.1 | -0.2664346 | 0.15690974 | 0.26570566 |
| sp\|Q9BXV9\|( | 1902691.44 | 2138593.97 | 0.25494616 | 0.15690974 | 0.26570566 |

| | | | | | |
|---|---|---|---|---|---|
| sp\|P49961\|E | 89884384.7 | 81880633.3 | -0.2714979 | 0.15690974 | 0.26570566 |
| sp\|Q9Y4D7\|F | 8479783.78 | 9916900.27 | 0.16436714 | 0.15690974 | 0.26570566 |
| sp\|P28715\|E | 728409.059 | 949384.543 | 0.07251706 | 0.15690974 | 0.26570566 |
| sp\|Q96PR1\|K | 12097969.1 | 14199964.9 | 0.15649857 | 0.15690974 | 0.26570566 |
| sp\|Q5SXM8\|I | 537436.704 | 731748.188 | 0.30185684 | 0.15690974 | 0.26570566 |
| sp\|P11177\|C | 198450238 | 165930409 | -0.4511375 | 0.15690974 | 0.26570566 |
| sp\|P61289\|P | 1889082.08 | 2538735.85 | 0.31885095 | 0.15690974 | 0.26570566 |
| sp\|TRFE_HUN | 3.834E+10 | 4.5177E+10 | 0.23750979 | 0.15690974 | 0.26570566 |
| sp\|Q9Y530\|C | 522939.43 | 412478.552 | 0.18820773 | 0.15690974 | 0.26570566 |
| sp\|Q9UH65\|S | 6502552.22 | 7806962.11 | 0.24973593 | 0.15690974 | 0.26570566 |
| sp\|O00291\|H | 1211682.52 | 1455978.36 | 0.22598285 | 0.15690974 | 0.26570566 |
| sp\|Q9UD71\|I | 38782360.2 | 39720455.9 | 0.24933886 | 0.15690974 | 0.26570566 |
| sp\|Q14152\|E | 156970334 | 134459243 | -0.1829865 | 0.15690974 | 0.26570566 |
| sp\|P02750\|A | 400067275 | 451874688 | 0.22426953 | 0.15690974 | 0.26570566 |
| sp\|CAH2_HU | 1832217149 | 2753687342 | 0.3292571 | 0.15690974 | 0.26570566 |
| sp\|Q9NP92\|I | 6196696.17 | 5696058.54 | -0.1643183 | 0.15690974 | 0.26570566 |
| sp\|P50552\|V | 53463961.7 | 63275830.6 | 0.10203636 | 0.15690974 | 0.26570566 |
| sp\|Q13347\|E | 69475942.2 | 59314299.9 | -0.3725588 | 0.15690974 | 0.26570566 |
| sp\|P53611\|P | 6388663.97 | 7426479.39 | 0.15506249 | 0.15690974 | 0.26570566 |
| sp\|Q9GZZ1\|N | 9046890.41 | 8339995.6 | -0.2255027 | 0.15690974 | 0.26570566 |
| sp\|Q15424\|S | 25610561.4 | 22276620.9 | -0.1987094 | 0.15690974 | 0.26570566 |
| sp\|Q9HD33\| | 4319317.41 | 3825258.54 | -0.3632773 | 0.15690974 | 0.26570566 |
| sp\|P48061\|S | 1265438.37 | 1426203.11 | 0.40294322 | 0.15690974 | 0.26570566 |
| sp\|P19474\|R | 15763954.2 | 14165757 | -0.1788025 | 0.15690974 | 0.26570566 |
| sp\|Q5VZ89\|D | 890234.644 | 996844.936 | -0.0302719 | 0.15690974 | 0.26570566 |
| sp\|Q9H9P8\|I | 2038813.19 | 1680475.02 | -0.3958515 | 0.15690974 | 0.26570566 |
| sp\|Q9H9B4\|S | 2992072.96 | 2258394.92 | -0.639239 | 0.15690974 | 0.26570566 |
| sp\|P23083\|H | 592513641 | 637822217 | 0.17341934 | 0.16379901 | 0.27465809 |
| sp\|P55008\|A | 22666848.2 | 27744012.5 | 0.20623289 | 0.16379901 | 0.27465809 |
| sp\|P08243\|A | 12031553.6 | 11618022.7 | 0.24543603 | 0.16379901 | 0.27465809 |
| sp\|Q15080\|N | 1262101.09 | 2648891.42 | 0.32538834 | 0.16379901 | 0.27465809 |
| sp\|P24844\|M | 56360079.9 | 61440044.3 | 0.08431665 | 0.16379901 | 0.27465809 |
| sp\|Q99829\|C | 54344987.3 | 44076367.9 | -0.3702787 | 0.16379901 | 0.27465809 |
| sp\|Q9Y4G8\|F | 8375706.12 | 7607464.01 | -0.1954666 | 0.16379901 | 0.27465809 |
| sp\|P54687\|B | 76755325.1 | 87632849.3 | 0.10581246 | 0.16379901 | 0.27465809 |
| sp\|Q5H9R7\|I | 3288514.95 | 3696561.99 | 0.09508134 | 0.16379901 | 0.27465809 |
| sp\|P23526\|S | 583046379 | 675420088 | 0.14393822 | 0.16379901 | 0.27465809 |
| sp\|Q9H1E3\|I | 37728148.1 | 30716344.5 | -0.5447882 | 0.16379901 | 0.27465809 |
| sp\|P36776\|L | 13155466.5 | 15325343.6 | 0.19607769 | 0.16379901 | 0.27465809 |

| | | | | | |
|---|---|---|---|---|---|
| sp\|Q15485\|F | 42278991.3 | 50498674.7 | 0.10431776 | 0.16379901 | 0.27465809 |
| sp\|Q9BRX8\|F | 73541401.6 | 87566011.5 | 0.18637901 | 0.16379901 | 0.27465809 |
| sp\|P02765\|F | 1277850777 | 1497838329 | 0.20674198 | 0.16379901 | 0.27465809 |
| sp\|Q8NB37\|( | 3298893.28 | 2886962.49 | -0.3764275 | 0.16379901 | 0.27465809 |
| sp\|Q9NXG6\| | 2840893.83 | 2474302.63 | -0.3012847 | 0.16379901 | 0.27465809 |
| sp\|Q13523\|F | 2239537.23 | 1983530.53 | -0.7281344 | 0.16379901 | 0.27465809 |
| sp\|Q9NSA3\|( | 883850.822 | 1130092.5 | 0.41442249 | 0.16379901 | 0.27465809 |
| sp\|Q9BU61\|I | 3144118.4 | 4000712.8 | 0.33887656 | 0.16379901 | 0.27465809 |
| sp\|Q9Y3C0\|\ | 11992296.7 | 10410881.9 | -0.1967168 | 0.16379901 | 0.27465809 |
| sp\|P61981\|1 | 2176153948 | 2357000037 | 0.10185785 | 0.16379901 | 0.27465809 |
| sp\|A0A0B4J1 | 370862980 | 407000132 | 0.20011517 | 0.16379901 | 0.27465809 |
| sp\|P0DP08\|H | 108532114 | 123735911 | 0.22592001 | 0.16379901 | 0.27465809 |
| sp\|P10321\|H | 27764530.2 | 26105026.5 | -0.1706786 | 0.16379901 | 0.27465809 |
| sp\|Q6ZU15\|S | 52673498.5 | 42364082.3 | -0.3951652 | 0.16379901 | 0.27465809 |
| sp\|Q01581\|H | 641327.396 | 747911.532 | 0.03621416 | 0.16379901 | 0.27465809 |
| sp\|Q9Y570\|F | 21281449.6 | 20977658.8 | -0.2044029 | 0.16379901 | 0.27465809 |
| sp\|Q12888\|T | 3363039.95 | 3111263.55 | -0.0814871 | 0.17090607 | 0.28438912 |
| sp\|P17812\|P | 26721815.8 | 23088722.5 | -0.2732264 | 0.17090607 | 0.28438912 |
| sp\|P02751\|F | 422717061 | 624971882 | 0.17488549 | 0.17090607 | 0.28438912 |
| sp\|Q15751\|H | 6052646.25 | 5247044.12 | -0.3605813 | 0.17090607 | 0.28438912 |
| sp\|Q6NUK1\|S | 32858414.4 | 27295773.7 | -0.2244834 | 0.17090607 | 0.28438912 |
| sp\|Q9Y2L5\|T | 6623637.09 | 5514238.22 | -0.4909634 | 0.17090607 | 0.28438912 |
| sp\|O15372\|E | 2518588.34 | 2331850.42 | -0.188356 | 0.17090607 | 0.28438912 |
| sp\|Q6PKG0\|I | 16990084.3 | 15715313.9 | -0.1332998 | 0.17090607 | 0.28438912 |
| sp\|Q93062\|F | 10337110.8 | 9227155.07 | -0.2505499 | 0.17090607 | 0.28438912 |
| sp\|P36405\|A | 19062643.5 | 17563679.7 | -0.1669205 | 0.17090607 | 0.28438912 |
| sp\|Q969T9\|V | 88136700.4 | 112316767 | 0.21140905 | 0.17090607 | 0.28438912 |
| sp\|Q9UMR2\| | 24791609 | 28369356.8 | 0.13605356 | 0.17090607 | 0.28438912 |
| sp\|Q8WVY7\| | 11661363.8 | 13965970.9 | 0.24664348 | 0.17090607 | 0.28438912 |
| sp\|Q9H2V7\|S | 1592722.41 | 1952805.28 | 0.17463286 | 0.17090607 | 0.28438912 |
| sp\|Q9NQA3\| | 3631992.58 | 4106938.41 | 0.21315401 | 0.17090607 | 0.28438912 |
| sp\|Q96MI6\|F | 1923798.88 | 2246268.95 | 0.48318029 | 0.17090607 | 0.28438912 |
| sp\|Q9UPN3\| | 4453251.66 | 3779959.88 | -0.2966367 | 0.17090607 | 0.28438912 |
| sp\|P61163\|A | 131574051 | 120379824 | -0.1824834 | 0.17090607 | 0.28438912 |
| sp\|P26583\|H | 21070342.7 | 19324138.4 | -0.1392881 | 0.17090607 | 0.28438912 |
| sp\|P02748\|C | 192066858 | 175162604 | -0.40401 | 0.17090607 | 0.28438912 |
| sp\|P57721\|P | 62091088.2 | 56496815.8 | -0.1841536 | 0.17090607 | 0.28438912 |
| sp\|Q96A26\|F | 3920742.46 | 3400861.65 | -0.2013691 | 0.17090607 | 0.28438912 |
| sp\|P15927\|R | 5976635.03 | 5111058.66 | -0.26709 | 0.17823377 | 0.29352911 |

| | | | | | |
|---|---|---|---|---|---|
| sp\|Q9HCC0\| | 100826523 | 107365553 | 0.1288376 | 0.17823377 | 0.29352911 |
| sp\|Q8TE77\|S | 7639283.6 | 6913031.58 | -0.1392679 | 0.17823377 | 0.29352911 |
| sp\|Q96EK7\|F | 3088846.5 | 3412035.71 | 0.1268391 | 0.17823377 | 0.29352911 |
| sp\|Q9NX62\|I | 255889.297 | 365071.158 | 0.39106401 | 0.17823377 | 0.29352911 |
| sp\|P35542\|S | 16276639.8 | 25250008.9 | 0.37877426 | 0.17823377 | 0.29352911 |
| sp\|Q14160\|S | 4065936.42 | 3781167.48 | -0.1552747 | 0.17823377 | 0.29352911 |
| sp\|Q00765\|F | 6556998.73 | 7678489.27 | 0.16196109 | 0.17823377 | 0.29352911 |
| sp\|O00299\|C | 466413399 | 423444923 | -0.1306082 | 0.17823377 | 0.29352911 |
| sp\|Q5T8P6\|F | 19278465.1 | 17298372.5 | -0.1668907 | 0.17823377 | 0.29352911 |
| sp\|P52907\|C | 264757123 | 242326055 | -0.2117752 | 0.17823377 | 0.29352911 |
| sp\|Q8N3E9\|I | 5123078.26 | 4443281.97 | -0.176318 | 0.17823377 | 0.29352911 |
| sp\|O14545\|T | 7642361.61 | 9421924.93 | 0.17836557 | 0.17823377 | 0.29352911 |
| sp\|Q9UKV8\|A | 49496750.8 | 45191757.3 | -0.2381298 | 0.17823377 | 0.29352911 |
| sp\|Q9P2X3\|I | 1858588.86 | 2063757.71 | 0.15362191 | 0.17823377 | 0.29352911 |
| sp\|Q96RF0\|S | 11481325.3 | 14372432.3 | 0.16977251 | 0.17823377 | 0.29352911 |
| sp\|P55010\|IF | 27475600.5 | 24377277.3 | -0.2332203 | 0.17823377 | 0.29352911 |
| sp\|Q9BXW6\| | 15916515.4 | 14091778.9 | -0.1987686 | 0.17823377 | 0.29352911 |
| sp\|P41222\|P | 39219213 | 30792491.5 | -0.3774522 | 0.17823377 | 0.29352911 |
| sp\|O14653\|C | 582492.239 | 826304.876 | 0.38040245 | 0.17823377 | 0.29352911 |
| sp\|P17980\|P | 67956675 | 81630842.5 | 0.20321124 | 0.17823377 | 0.29352911 |
| sp\|Q9NZT2\|C | 15506205.2 | 13857435.7 | -0.2240459 | 0.17823377 | 0.29352911 |
| sp\|P78344\|IF | 209525.512 | 192845.044 | -1.4047925 | 0.17823377 | 0.29352911 |
| sp\|Q9BRR6\|A | 641150.26 | 723333.403 | 0.00708197 | 0.17823377 | 0.29352911 |
| sp\|Q13011\|E | 947811719 | 1045586089 | 0.12820001 | 0.17823377 | 0.29352911 |
| sp\|P01762\|H | 868691723 | 962162832 | 0.1568391 | 0.17823377 | 0.29352911 |
| sp\|P54105\|IC | 10507816.8 | 9618459.65 | -0.1839045 | 0.17823377 | 0.29352911 |
| sp\|P40227\|T | 64678917.4 | 59072481.8 | -0.2128173 | 0.17823377 | 0.29352911 |
| sp\|Q96JQ0\|F | 1452307.7 | 3068767.43 | 0.60855272 | 0.17823377 | 0.29352911 |
| sp\|Q9H0W9\| | 161235005 | 149777351 | -0.2117594 | 0.17823377 | 0.29352911 |
| sp\|P50053\|K | 467982.619 | 1251762.86 | -0.2203744 | 0.18578492 | 0.30325913 |
| sp\|Q68DX3\|F | 7537125.35 | 8491831.04 | 0.10393923 | 0.18578492 | 0.30325913 |
| sp\|Q16513\|F | 8984638.51 | 10429158.7 | 0.18550402 | 0.18578492 | 0.30325913 |
| sp\|O60927\|F | 11725386.8 | 12672642.1 | 0.07440234 | 0.18578492 | 0.30325913 |
| sp\|P08253\|M | 1017404.1 | 1323867.06 | 0.40641352 | 0.18578492 | 0.30325913 |
| sp\|Q8N357\|S | 1651980.47 | 1725876.7 | 0.22304951 | 0.18578492 | 0.30325913 |
| sp\|Q06830\|F | 3215640902 | 3088128957 | -0.0628976 | 0.18578492 | 0.30325913 |
| sp\|P14770\|G | 1942201.39 | 3184421.12 | 0.44347765 | 0.18578492 | 0.30325913 |
| sp\|Q92765\|S | 4844792.13 | 5613119.01 | 0.1864837 | 0.18578492 | 0.30325913 |
| sp\|Q92496\|F | 35819931.3 | 44844729.4 | 0.24260722 | 0.18578492 | 0.30325913 |

| | | | | | |
|---|---|---|---|---|---|
| sp\|Q9H2U2\| | 112272729 | 121114295 | 0.0619953 | 0.18578492 | 0.30325913 |
| sp\|Q9NRW1\| | 62851318.5 | 70188061.3 | 0.15098426 | 0.18578492 | 0.30325913 |
| sp\|Q5JWF2\|( | 80025325.9 | 77179882.4 | -0.083139 | 0.18578492 | 0.30325913 |
| sp\|Q96JQ2\|( | 906480.487 | 1045864.64 | 0.19188176 | 0.18578492 | 0.30325913 |
| sp\|Q9NTZ6\|F | 4501108 | 4085182.35 | -0.1393136 | 0.18578492 | 0.30325913 |
| sp\|P55854\|S | 160329188 | 198343072 | 0.23176518 | 0.18578492 | 0.30325913 |
| sp\|Q5MNZ6\| | 1757454.4 | 1654652.14 | -0.742519 | 0.18578492 | 0.30325913 |
| sp\|O00264\|F | 57668808.7 | 53311473 | -0.1406278 | 0.18578492 | 0.30325913 |
| sp\|Q8IVD9\|N | 3597190.92 | 4345269.15 | 0.34728556 | 0.18578492 | 0.30325913 |
| sp\|Q00341\|V | 37477227.9 | 33832473.9 | -0.176859 | 0.18578492 | 0.30325913 |
| sp\|Q14005\|I | 106553826 | 98731005.7 | -0.1376738 | 0.18578492 | 0.30325913 |
| sp\|Q14103\|H | 340447420 | 315078926 | -0.2183497 | 0.18578492 | 0.30325913 |
| sp\|O43815\|S | 54042136.4 | 56811206.7 | 0.07068169 | 0.18578492 | 0.30325913 |
| sp\|P78560\|C | 660615.569 | 736322.659 | 0.22165513 | 0.18578492 | 0.30325913 |
| sp\|A0A0B4J1 | 73392677.5 | 80893265.2 | 0.14984444 | 0.18578492 | 0.30325913 |
| sp\|Q96EY8\|N | 2646102.23 | 3461969.58 | 0.44594519 | 0.18578492 | 0.30325913 |
| sp\|Q9Y676\|F | 2104082.82 | 2389454.09 | 0.06806309 | 0.19356208 | 0.31381905 |
| sp\|Q5TBB1\|F | 1173629.48 | 1467448.61 | 0.40099994 | 0.19356208 | 0.31381905 |
| sp\|Q5JVS0\|H | 737472.949 | 959138.716 | 0.33150425 | 0.19356208 | 0.31381905 |
| sp\|Q99622\|( | 1372901.43 | 1195472.78 | -0.1861923 | 0.19356208 | 0.31381905 |
| sp\|P11802\|C | 1460114.72 | 2284739 | -0.4149771 | 0.19356208 | 0.31381905 |
| sp\|P14618\|K | 2412978952 | 2303925499 | -0.0729616 | 0.19356208 | 0.31381905 |
| sp\|Q9Y2Z9\|( | 42460976.8 | 49993087.1 | 0.22012823 | 0.19356208 | 0.31381905 |
| sp\|Q14061\|( | 2052742.93 | 2565644.4 | 0.30724157 | 0.19356208 | 0.31381905 |
| sp\|O43508\|T | 6478199.22 | 7147222.97 | 0.13926382 | 0.19356208 | 0.31381905 |
| sp\|P26927\|H | 24986954.3 | 22204268.8 | -0.2401401 | 0.19356208 | 0.31381905 |
| sp\|Q15102\|F | 70358690.1 | 66598532.4 | -0.3416354 | 0.19356208 | 0.31381905 |
| sp\|A0AVT1\|U | 2628866.87 | 3323627.4 | 0.21059623 | 0.19356208 | 0.31381905 |
| sp\|Q9HAU0\| | 9829854.02 | 9044934.97 | -0.1953446 | 0.19356208 | 0.31381905 |
| sp\|P42892\|E | 1277447.82 | 1434257.98 | 0.15147271 | 0.19356208 | 0.31381905 |
| sp\|P41223\|B | 931048.606 | 791415.682 | -0.2910979 | 0.19356208 | 0.31381905 |
| sp\|P05976\|M | 62128729.6 | 315542639 | 0.53294748 | 0.19356208 | 0.31381905 |
| sp\|P53674\|C | 963176.938 | 1118825.36 | 0.2838658 | 0.19356208 | 0.31381905 |
| sp\|Q7LGC8\|( | 161797.314 | 142761.568 | -2.7517561 | 0.19356208 | 0.31381905 |
| sp\|Q02809\|F | 5193465.56 | 5553975.34 | 0.08678128 | 0.19356208 | 0.31381905 |
| sp\|P56159\|G | 1924390.96 | 1665947.28 | -0.2996695 | 0.19356208 | 0.31381905 |
| sp\|P20042\|IF | 4358192.82 | 3865351.23 | -0.2681336 | 0.20156763 | 0.32351941 |
| sp\|P35556\|F | 793145365 | 985684130 | 0.31822515 | 0.20156763 | 0.32351941 |
| sp\|P23469\|P | 945755.009 | 1985514.24 | 0.44486066 | 0.20156763 | 0.32351941 |

| | | | | | |
|---|---|---|---|---|---|
| sp\|Q9UHC6\| | 7715332.41 | 6721456.57 | -0.4560861 | 0.20156763 | 0.32351941 |
| sp\|Q9Y2B9\|I | 6197833.07 | 6543993.1 | 0.06317322 | 0.20156763 | 0.32351941 |
| sp\|O75663\|T | 1538391.36 | 1731195.3 | 0.11401084 | 0.20156763 | 0.32351941 |
| sp\|Q16718\|N | 6528448.02 | 7888553.73 | 0.18527539 | 0.20156763 | 0.32351941 |
| sp\|Q9H910\|J | 57817298 | 66319530.3 | 0.15726437 | 0.20156763 | 0.32351941 |
| sp\|P0DTE8\|A | 111353.255 | 196393.555 | 0.54865383 | 0.20156763 | 0.32351941 |
| sp\|Q5JTD0\|T | 343195.597 | 429644.953 | -0.3641966 | 0.20156763 | 0.32351941 |
| sp\|P00326\|A | 5237994349 | 4920336286 | -0.3542307 | 0.20156763 | 0.32351941 |
| sp\|Q96FV2\|S | 97325088 | 121736967 | 0.12414586 | 0.20156763 | 0.32351941 |
| sp\|Q9NR31\|S | 77290865.2 | 91306999.6 | 0.13819285 | 0.20156763 | 0.32351941 |
| sp\|Q9NUQ6\| | 429536.885 | 324706.846 | -0.2290734 | 0.20156763 | 0.32351941 |
| sp\|P40222\|T | 6353305.14 | 7637466.21 | 0.11265552 | 0.20156763 | 0.32351941 |
| sp\|O43290\|S | 2224678.21 | 2690193.79 | 0.29241525 | 0.20156763 | 0.32351941 |
| sp\|Q9Y394\|D | 8317099.99 | 7560762.46 | -0.1587193 | 0.20156763 | 0.32351941 |
| sp\|Q13162\|P | 874505644 | 894868422 | 0.01150271 | 0.20156763 | 0.32351941 |
| sp\|Q92733\|P | 13527817.8 | 21229711.2 | 0.31811536 | 0.20156763 | 0.32351941 |
| sp\|Q96RQ3\|I | 28822253 | 26572517.6 | -0.1796234 | 0.20156763 | 0.32351941 |
| sp\|P56377\|A | 708050.922 | 1023594.82 | 0.24149289 | 0.20156763 | 0.32351941 |
| sp\|Q96AE4\|F | 175742019 | 157361205 | -0.195379 | 0.20156763 | 0.32351941 |
| sp\|Q96B45\|E | 11799363.4 | 13283209.7 | -0.0052738 | 0.20156763 | 0.32351941 |
| sp\|Q9BT04\|F | 7507163.41 | 9051197.28 | 0.24734975 | 0.20156763 | 0.32351941 |
| sp\|Q9Y5S9\|F | 4390320.98 | 4075115.03 | -0.1803939 | 0.20156763 | 0.32351941 |
| sp\|Q96MM6\| | 40556031.1 | 38227144.5 | -0.0974616 | 0.20156763 | 0.32351941 |
| sp\|Q969E2\|S | 1383211.81 | 1115044.99 | -0.3266233 | 0.20156763 | 0.32351941 |
| sp\|Q9UN36\| | 91990359.4 | 83907565.1 | -0.1123387 | 0.20156763 | 0.32351941 |
| sp\|Q9Y279\|V | 16021576 | 13579484.4 | -0.3867669 | 0.20156763 | 0.32351941 |
| sp\|P78396\|C | 605079.552 | 785272.173 | 0.31913652 | 0.20156763 | 0.32351941 |
| sp\|Q68DU8\|I | 675130.818 | 1010483.78 | -0.4539359 | 0.20614972 | 0.33076312 |
| sp\|Q8TAC1\|F | 1687213.98 | 2092054.7 | 0.21666714 | 0.20980394 | 0.33328338 |
| sp\|P27169\|P | 56432279.1 | 70289077.7 | 0.17773875 | 0.20980394 | 0.33328338 |
| sp\|O60674\|J | 1901292.75 | 1738065.66 | -0.1731595 | 0.20980394 | 0.33328338 |
| sp\|Q92530\|P | 10630412.7 | 15937261.3 | 0.28414564 | 0.20980394 | 0.33328338 |
| sp\|Q9BY42\|F | 516720.911 | 848609.335 | -0.3449824 | 0.20980394 | 0.33328338 |
| sp\|Q14694\|U | 563767.583 | 482863.213 | -0.2724468 | 0.20980394 | 0.33328338 |
| sp\|P29144\|T | 168700308 | 159729832 | -0.1870654 | 0.20980394 | 0.33328338 |
| sp\|P08246\|E | 38776472.7 | 179761739 | 0.58114464 | 0.20980394 | 0.33328338 |
| sp\|Q9UPY6\|V | 4008909.47 | 5648043.7 | 0.30550646 | 0.20980394 | 0.33328338 |
| sp\|Q9P2X0\|D | 414032.663 | 495788.669 | 0.34853211 | 0.20980394 | 0.33328338 |
| sp\|P13987\|C | 17418160.9 | 20941797.7 | 0.1890686 | 0.20980394 | 0.33328338 |

| | | | | | |
|---|---|---|---|---|---|
| sp\|Q8IWT6\|L | 1685106.58 | 1979084.98 | 0.28889313 | 0.20980394 | 0.33328338 |
| sp\|Q92817\|E | 996488.053 | 800859.654 | -0.5908997 | 0.20980394 | 0.33328338 |
| sp\|P51812\|K | 10532309.5 | 11862746.1 | 0.19982145 | 0.20980394 | 0.33328338 |
| sp\|P60059\|S | 3822396.59 | 4433164.82 | 0.11572753 | 0.20980394 | 0.33328338 |
| sp\|Q04864\|F | 547712.922 | 430041.322 | -0.3760339 | 0.20980394 | 0.33328338 |
| sp\|Q96KS0\|E | 527923.413 | 437057.756 | -0.2908376 | 0.20980394 | 0.33328338 |
| sp\|Q9BSY4\|C | 474988.175 | 620652.509 | 0.28565798 | 0.20980394 | 0.33328338 |
| sp\|Q05519\|S | 50787721.3 | 47902168.7 | -0.085671 | 0.20980394 | 0.33328338 |
| sp\|Q08379\|C | 781776.236 | 683186.001 | -0.4005169 | 0.20980394 | 0.33328338 |
| sp\|P35858\|A | 107974726 | 96389634.9 | -0.4430344 | 0.20980394 | 0.33328338 |
| sp\|P01701\|L | 15113071.8 | 17205113.6 | 0.24959868 | 0.20980394 | 0.33328338 |
| sp\|Q13480\|C | 6889500.69 | 7628862.4 | 0.09773207 | 0.20980394 | 0.33328338 |
| sp\|P45974\|U | 215737381 | 244434156 | 0.13892027 | 0.20980394 | 0.33328338 |
| sp\|O14972\|V | 1473969.11 | 1812215.71 | 0.33572681 | 0.20980394 | 0.33328338 |
| sp\|O15018\|F | 887701.797 | 1297409.13 | 0.30191796 | 0.20980394 | 0.33328338 |
| sp\|P78417\|G | 171836820 | 223314427 | 0.36224468 | 0.20980394 | 0.33328338 |
| sp\|P78371\|T | 82427193.2 | 78330017.7 | -0.1638732 | 0.20980394 | 0.33328338 |
| sp\|A0A087W | 108716864 | 114572512 | 0.31502165 | 0.20980394 | 0.33328338 |
| sp\|A0A075B6 | 51220814.7 | 57067321.9 | 0.21039011 | 0.20980394 | 0.33328338 |
| sp\|P15336\|A | 109107.666 | 131638.385 | -0.1374383 | 0.21827306 | 0.34355277 |
| sp\|P12259\|F | 7541389.07 | 8945760.35 | 0.15530959 | 0.21827306 | 0.34355277 |
| sp\|Q15437\|S | 1494240.08 | 1270723.26 | -0.2621172 | 0.21827306 | 0.34355277 |
| sp\|O94830\|D | 9135781.42 | 10459841.4 | 0.22598735 | 0.21827306 | 0.34355277 |
| sp\|Q6DD87\|Z | 940870.494 | 1382925.33 | 0.37840163 | 0.21827306 | 0.34355277 |
| sp\|Q9BVT8\|T | 3146245.32 | 3817901.13 | 0.14421507 | 0.21827306 | 0.34355277 |
| sp\|O95372\|L | 30061278.4 | 45231402.5 | 0.74779281 | 0.21827306 | 0.34355277 |
| sp\|P20290\|B | 121138572 | 106311494 | -0.2166733 | 0.21827306 | 0.34355277 |
| sp\|Q9H074\|I | 50139666.7 | 34361531.2 | -0.4457156 | 0.21827306 | 0.34355277 |
| sp\|O15541\|F | 2775258.39 | 2577291.18 | -0.1299146 | 0.21827306 | 0.34355277 |
| sp\|O43150\|A | 4079769.23 | 5999204.58 | 0.24774921 | 0.21827306 | 0.34355277 |
| sp\|Q15404\|F | 138131539 | 149099309 | 0.06257621 | 0.21827306 | 0.34355277 |
| sp\|Q86SQ0\|I | 6859340.1 | 7762983.53 | 0.12935022 | 0.21827306 | 0.34355277 |
| sp\|Q9NRG7\| | 208205.484 | 189108.425 | -0.2764366 | 0.21827306 | 0.34355277 |
| sp\|P51157\|R | 204028797 | 227246122 | 0.13462381 | 0.21827306 | 0.34355277 |
| sp\|P98164\|L | 10301889.8 | 9513759.22 | -0.1178419 | 0.21827306 | 0.34355277 |
| sp\|P02788\|T | 179088636 | 636633648 | 0.40703427 | 0.21827306 | 0.34355277 |
| sp\|HBA_HUM | 2.32E+11 | 3.59E+11 | 0.08465408 | 0.21827306 | 0.34355277 |
| sp\|O00303\|E | 36124009.9 | 34379672.9 | -0.1432369 | 0.21827306 | 0.34355277 |
| sp\|Q6ZNL6\|F | 1130241.53 | 1274484.32 | 0.22157343 | 0.21827306 | 0.34355277 |

| | | | | | |
|---|---|---|---|---|---|
| sp\|Q96JY6\|P | 20158179.5 | 17847496.1 | -0.2505131 | 0.21827306 | 0.34355277 |
| sp\|P54577\|S | 66239634.3 | 74822214.4 | 0.1249066 | 0.21827306 | 0.34355277 |
| sp\|P48643\|T | 86513334.8 | 83597627.9 | -0.1858103 | 0.21827306 | 0.34355277 |
| sp\|Q13094\|L | 7609391.41 | 11825965.9 | 0.31958886 | 0.21827306 | 0.34355277 |
| sp\|Q9H694\|I | 839339.661 | 1033412.57 | 0.30109233 | 0.21827306 | 0.34355277 |
| sp\|Q9Y2C3\|I | 498930.532 | 700807.029 | 0.39188738 | 0.21827306 | 0.34355277 |
| sp\|P78504\|J | 356990.73 | 428369.193 | 0.26936943 | 0.21827306 | 0.34355277 |
| sp\|Q9Y2J4\|A | 378468.81 | 411225.558 | 0.0855635 | 0.21827306 | 0.34355277 |
| sp\|Q16540\|F | 2051877.87 | 2454729.68 | -0.2761539 | 0.22697687 | 0.35446209 |
| sp\|Q9UHW9\| | 957944.803 | 1020153.86 | 0.08423538 | 0.22697687 | 0.35446209 |
| sp\|P08559\|C | 214179839 | 187753472 | -0.3323727 | 0.22697687 | 0.35446209 |
| sp\|Q9H1H9\| | 2900310.26 | 2636588.54 | -0.1341848 | 0.22697687 | 0.35446209 |
| sp\|P33240\|C | 21483492.5 | 23438816.1 | 0.10732428 | 0.22697687 | 0.35446209 |
| sp\|Q8WU90\| | 9323720.16 | 10011179.8 | 0.11545605 | 0.22697687 | 0.35446209 |
| sp\|Q8TBX8\|F | 1420850.33 | 1391943.37 | -0.191476 | 0.22697687 | 0.35446209 |
| sp\|P28332\|A | 136872356 | 135999654 | -0.3335528 | 0.22697687 | 0.35446209 |
| sp\|O75431\|N | 549760.892 | 729486.49 | 0.31475132 | 0.22697687 | 0.35446209 |
| sp\|O43242\|F | 64847201.3 | 59058719.6 | -0.2546381 | 0.22697687 | 0.35446209 |
| sp\|P01023\|A | 9836895539 | 1.1482E+10 | 0.2139025 | 0.22697687 | 0.35446209 |
| sp\|P49959\|M | 22057393.3 | 23010595.4 | 0.05059623 | 0.22697687 | 0.35446209 |
| sp\|O14686\|K | 6281172.33 | 7507719.01 | 0.07602649 | 0.22697687 | 0.35446209 |
| sp\|P05154\|IF | 38262038.9 | 43974213.2 | 0.09044176 | 0.22697687 | 0.35446209 |
| sp\|Q9HA64\|I | 57002626.5 | 52031037.8 | -0.151758 | 0.22697687 | 0.35446209 |
| sp\|P11362\|F | 58491179.5 | 51163658.9 | -0.3845386 | 0.22697687 | 0.35446209 |
| sp\|P16989\|Y | 47808304.7 | 53052925.8 | 0.11680043 | 0.22697687 | 0.35446209 |
| sp\|Q9Y2R5\|F | 5073172.35 | 4325181.97 | -0.3690528 | 0.22697687 | 0.35446209 |
| sp\|Q8NFZ8\|C | 1432487.75 | 1592759.23 | 0.13995008 | 0.22697687 | 0.35446209 |
| sp\|P03952\|K | 287761470 | 262970280 | -0.5440169 | 0.22697687 | 0.35446209 |
| sp\|Q9Y3X0\|C | 2351276.79 | 2584220.3 | 0.21059602 | 0.22697687 | 0.35446209 |
| sp\|Q9HD42\| | 3474790.88 | 4014748.7 | 0.19643085 | 0.22697687 | 0.35446209 |
| sp\|O43447\|F | 26693857 | 34147779.2 | 0.10994597 | 0.22697687 | 0.35446209 |
| sp\|P43304\|G | 7940861.14 | 7367287.09 | -0.3047185 | 0.22697687 | 0.35446209 |
| sp\|A0A075B6 | 19777954 | 23120468.1 | 0.19880974 | 0.23591723 | 0.36392374 |
| sp\|Q9Y5Z4\|H | 382328651 | 471928434 | 0.09812567 | 0.23591723 | 0.36392374 |
| sp\|P01834\|IC | 1.1672E+10 | 1.2635E+10 | 0.16030977 | 0.23591723 | 0.36392374 |
| sp\|P31321\|K | 14701947.5 | 16543361.4 | 0.1791512 | 0.23591723 | 0.36392374 |
| sp\|Q9H845\|A | 32555851.4 | 29125529.8 | -0.1754495 | 0.23591723 | 0.36392374 |
| sp\|P62136\|P | 64844373.9 | 60713499.4 | -0.0838122 | 0.23591723 | 0.36392374 |
| sp\|P36873\|P | 62254206.5 | 58915990.7 | -0.0698183 | 0.23591723 | 0.36392374 |

| | | | | | |
|---|---|---|---|---|---|
| sp\|O43294\|T | 45713797.8 | 40257115.6 | -0.1334925 | 0.23591723 | 0.36392374 |
| sp\|P05386\|R | 138695383 | 135051683 | -0.0974787 | 0.23591723 | 0.36392374 |
| sp\|Q96IV0\|N | 12038590.6 | 14921877.5 | 0.22938729 | 0.23591723 | 0.36392374 |
| sp\|P01703\|L | 8470300.51 | 10555615.2 | 0.23755996 | 0.23591723 | 0.36392374 |
| sp\|Q86VN1\|V | 2314073.79 | 2594091.03 | 0.15009363 | 0.23591723 | 0.36392374 |
| sp\|P82970\|H | 1912313.6 | 2176049.65 | 0.09598014 | 0.23591723 | 0.36392374 |
| sp\|P00748\|F | 219455665 | 192298176 | -0.4022872 | 0.23591723 | 0.36392374 |
| sp\|P60953\|C | 134498961 | 156466667 | 0.16009358 | 0.23591723 | 0.36392374 |
| sp\|P60033\|C | 37174570.4 | 33298504.1 | -0.2498814 | 0.23591723 | 0.36392374 |
| sp\|P59665\|D | 46742621.2 | 195702693 | 0.44268199 | 0.23591723 | 0.36392374 |
| sp\|Q6NUJ5\|F | 699284.883 | 843266.98 | 0.21830553 | 0.23591723 | 0.36392374 |
| sp\|Q01546\|K | 1265830341 | 755958446 | -0.5508978 | 0.23591723 | 0.36392374 |
| sp\|Q13370\|F | 646773.194 | 946977.846 | 0.30959147 | 0.23591723 | 0.36392374 |
| sp\|O00533\|N | 29476729.7 | 32986554.5 | 0.12959901 | 0.23591723 | 0.36392374 |
| sp\|P0DJG4\|T | 5141134.58 | 4303959.85 | -0.3225117 | 0.23591723 | 0.36392374 |
| sp\|P09668\|C | 23576206.4 | 30857072.3 | 0.17086413 | 0.23591723 | 0.36392374 |
| sp\|P08651\|N | 20138681.1 | 29909974.4 | 0.03817137 | 0.23591723 | 0.36392374 |
| sp\|P24158\|P | 18435911.9 | 69692283.1 | 0.44292402 | 0.23591723 | 0.36392374 |
| sp\|Q9BXX0\|E | 50156867.8 | 45693540.9 | -0.176694 | 0.23591723 | 0.36392374 |
| sp\|O76003\|G | 8741223.2 | 10290683.4 | 0.14951007 | 0.23591723 | 0.36392374 |
| sp\|O14672\|A | 45937881.1 | 48689604.6 | 0.04721627 | 0.23591723 | 0.36392374 |
| sp\|Q92506\|D | 36574778.6 | 31999266.2 | -0.1411748 | 0.23591723 | 0.36392374 |
| sp\|Q16204\|C | 34680199 | 36309867.8 | 0.05193803 | 0.23591723 | 0.36392374 |
| sp\|O95183\|V | 14053651.7 | 15689011.8 | 0.13452447 | 0.23591723 | 0.36392374 |
| sp\|O75436\|V | 106887771 | 113480372 | 0.09198665 | 0.23591723 | 0.36392374 |
| sp\|P78527\|P | 36837318.5 | 34361284.5 | -0.1101323 | 0.23591723 | 0.36392374 |
| sp\|Q9BRQ6\| | 2234664.95 | 2559709.9 | 0.05758576 | 0.23591723 | 0.36392374 |
| sp\|P62330\|A | 4316116.79 | 4680757.69 | 0.01754773 | 0.23591723 | 0.36392374 |
| sp\|Q13425\|S | 11456152.2 | 12219309.4 | 0.08101784 | 0.23591723 | 0.36392374 |
| sp\|P15924\|D | 2472774.57 | 2175382.45 | -0.1839295 | 0.23591723 | 0.36392374 |
| sp\|Q9UKN8\|T | 5847214.45 | 8259121.71 | 0.29026827 | 0.23591723 | 0.36392374 |
| sp\|P98088\|M | 294902.706 | 34520.5606 | -2.2776121 | 0.23919146 | 0.36885598 |
| sp\|P0C7P4\|U | 8678659.69 | 9887993.36 | 0.16091726 | 0.2450956 | 0.37494861 |
| sp\|Q6N063\|C | 97290.5334 | 126860.585 | -0.3779628 | 0.2450956 | 0.37494861 |
| sp\|Q9H254\|S | 24877690.7 | 22811090.7 | -0.0982151 | 0.2450956 | 0.37494861 |
| sp\|Q9BRQ0\| | 748666.216 | 688497.254 | -0.358948 | 0.2450956 | 0.37494861 |
| sp\|Q6ZNA5\|F | 2426073.52 | 2970231 | 0.32855826 | 0.2450956 | 0.37494861 |
| sp\|Q7Z2Q7\|L | 383455.585 | 314380.819 | -1.0140798 | 0.2450956 | 0.37494861 |
| sp\|O14979\|H | 149868410 | 133346641 | -0.2894854 | 0.2450956 | 0.37494861 |

| | | | | | |
|---|---|---|---|---|---|
| sp\|Q9NR34\|I | 17072191.4 | 15748187.3 | -0.4144179 | 0.2450956 | 0.37494861 |
| sp\|Q13242\|S | 6810706.67 | 5726398.21 | -0.3184811 | 0.2450956 | 0.37494861 |
| sp\|Q9NYU2\| | 133641764 | 122708630 | -0.1756121 | 0.2450956 | 0.37494861 |
| sp\|P10644\|K | 46690251.4 | 40837219.7 | -0.3411988 | 0.2450956 | 0.37494861 |
| sp\|P48200\|IF | 4100663.79 | 3948541.69 | -0.133656 | 0.2450956 | 0.37494861 |
| sp\|O75608\|L | 44325593.4 | 49151887.8 | 0.14396252 | 0.2450956 | 0.37494861 |
| sp\|Q8NCM8\| | 4947918.14 | 4571966.95 | -0.0602749 | 0.2450956 | 0.37494861 |
| sp\|Q96HS1\|I | 254039.335 | 221442.507 | -0.2438016 | 0.2450956 | 0.37494861 |
| sp\|P67936\|T | 1831473183 | 1880279126 | 0.0159938 | 0.2450956 | 0.37494861 |
| sp\|A0A0A0MT | 8877778.09 | 10047211.4 | 0.20313821 | 0.2450956 | 0.37494861 |
| sp\|P13797\|P | 182792160 | 162014788 | -0.2986438 | 0.2450956 | 0.37494861 |
| sp\|Q13057\|C | 20218928.3 | 18901087.8 | -0.3456794 | 0.2450956 | 0.37494861 |
| sp\|P27797\|C | 402003129 | 357910513 | -0.279197 | 0.2450956 | 0.37494861 |
| sp\|Q86XP3\|D | 17760727.4 | 16031217.3 | -0.1108437 | 0.2450956 | 0.37494861 |
| sp\|Q6P4A8\|F | 4481973.24 | 6171440.64 | 0.22635541 | 0.2450956 | 0.37494861 |
| sp\|Q86V40\|T | 7471292.14 | 10180127.5 | 0.29846629 | 0.2450956 | 0.37494861 |
| sp\|O60518\|F | 725227.224 | 1149168.22 | 0.79189378 | 0.2450956 | 0.37494861 |
| sp\|Q8TC12\|F | 484987.663 | 547515.588 | 0.08855972 | 0.2450956 | 0.37494861 |
| sp\|Q92520\|F | 9164811.54 | 11621115.3 | -0.2428081 | 0.25451332 | 0.38542425 |
| sp\|P11137\|M | 4342692.89 | 5074162.54 | 0.15336552 | 0.25451332 | 0.38542425 |
| sp\|Q8IX05\|C | 848721.652 | 730628.063 | -0.3006391 | 0.25451332 | 0.38542425 |
| sp\|Q16555\|D | 1950515073 | 2097946446 | 0.06147682 | 0.25451332 | 0.38542425 |
| sp\|P53367\|A | 18278398.8 | 19950664.8 | 0.08440536 | 0.25451332 | 0.38542425 |
| sp\|Q9NX08\|C | 105695.633 | 129813.388 | -0.7698445 | 0.25451332 | 0.38542425 |
| sp\|Q13576\|I | 1070303.94 | 1213787.65 | 0.12352839 | 0.25451332 | 0.38542425 |
| sp\|Q9NWV8\| | 7873408.6 | 9056683.77 | 0.12619661 | 0.25451332 | 0.38542425 |
| sp\|Q96SW2\| | 4565438.27 | 5008233.27 | 0.11851234 | 0.25451332 | 0.38542425 |
| sp\|Q9NXR7\|I | 446873.182 | 588241.27 | 0.31697318 | 0.25451332 | 0.38542425 |
| sp\|Q9GZN8\| | 7144982.41 | 8916477.52 | 0.16901003 | 0.25451332 | 0.38542425 |
| sp\|O95544\|N | 2678365.37 | 2956725.03 | 0.14202338 | 0.25451332 | 0.38542425 |
| sp\|Q9UNW1\| | 5019331.79 | 6254941.21 | 0.1867591 | 0.25451332 | 0.38542425 |
| sp\|P07947\|Y | 76735675.7 | 70658441.9 | -0.1475292 | 0.25451332 | 0.38542425 |
| sp\|P01871\|IC | 2469874727 | 2788866599 | 0.09798518 | 0.25451332 | 0.38542425 |
| sp\|P07954\|F | 329210549 | 361861059 | 0.05818531 | 0.25451332 | 0.38542425 |
| sp\|Q8TCT9\|H | 1338450.37 | 1181108.03 | -0.3096992 | 0.25451332 | 0.38542425 |
| sp\|P61201\|C | 11246782.4 | 13455409.3 | 0.16750295 | 0.25451332 | 0.38542425 |
| sp\|PPIA_HUM | 2461534404 | 2919437980 | 0.1393228 | 0.25451332 | 0.38542425 |
| sp\|P06331\|H | 112293740 | 123488495 | 0.18390637 | 0.25451332 | 0.38542425 |
| sp\|P05089\|A | 17950812.2 | 24366514.8 | 0.14340256 | 0.25451332 | 0.38542425 |

| | | | | | |
|---|---|---|---|---|---|
| sp\|Q16769\|( | 4001166.45 | 5287021.33 | 0.44919166 | 0.25451332 | 0.38542425 |
| sp\|P45877\|P | 4916607.54 | 4481851.83 | -0.2956386 | 0.25451332 | 0.38542425 |
| sp\|O14773\|T | 169633277 | 154632314 | -0.2019162 | 0.25451332 | 0.38542425 |
| sp\|Q8NFC6\|I | 7120755.12 | 6431179.32 | -0.2233979 | 0.25451332 | 0.38542425 |
| sp\|Q8IYS1\|P: | 491590.013 | 411852.546 | -0.6459035 | 0.25451332 | 0.38542425 |
| sp\|Q9P1Z2\|( | 3583695.62 | 3368044.95 | -0.1837474 | 0.25451332 | 0.38542425 |
| sp\|Q6ZT62\|B | 4657297.81 | 5690868.43 | 0.22148699 | 0.25451332 | 0.38542425 |
| sp\|P14923\|P | 30654092.2 | 28729717.1 | -0.145004 | 0.25451332 | 0.38542425 |
| sp\|Q9Y2E6\|D | 1858026.62 | 1713386.92 | -0.2910169 | 0.25451332 | 0.38542425 |
| sp\|O95050\|I | 214034.797 | 137891.071 | -2.6758348 | 0.25451332 | 0.38542425 |
| sp\|O94874\|U | 7444745.17 | 5809398.05 | -0.7478631 | 0.25451332 | 0.38542425 |
| sp\|A0A0C4DI | 326749596 | 362674348 | 0.16050296 | 0.26417164 | 0.39667074 |
| sp\|Q99571\|F | 477730.19 | 588723.075 | 0.17793575 | 0.26417164 | 0.39667074 |
| sp\|Q8NAT2\|T | 1774984.71 | 1570005.39 | -0.1466979 | 0.26417164 | 0.39667074 |
| sp\|Q9Y3M8\|S | 697276.567 | 980912.77 | 0.40740323 | 0.26417164 | 0.39667074 |
| sp\|Q96TA1\|N | 3413809.77 | 3990201.76 | -0.1182055 | 0.26417164 | 0.39667074 |
| sp\|Q15435\|F | 90578327 | 84331117.9 | -0.0947666 | 0.26417164 | 0.39667074 |
| sp\|P52657\|T | 1461583.66 | 1226633.49 | -0.6301022 | 0.26417164 | 0.39667074 |
| sp\|Q08426\|E | 397879.269 | 336510.587 | -0.2962931 | 0.26417164 | 0.39667074 |
| sp\|Q8IZ83\|A | 65653341.5 | 73404016.3 | 0.09122417 | 0.26417164 | 0.39667074 |
| sp\|Q10588\|E | 40580309.2 | 36006086.7 | -0.187883 | 0.26417164 | 0.39667074 |
| sp\|Q8N8R5\|( | 569533.864 | 638492.866 | 0.11027208 | 0.26417164 | 0.39667074 |
| sp\|Q9Y426\|( | 137963.161 | 188263.685 | 0.31898409 | 0.26417164 | 0.39667074 |
| sp\|P48426\|P | 37870828.9 | 34782104.7 | -0.4827961 | 0.26417164 | 0.39667074 |
| sp\|P82664\|R | 864703.722 | 1261535.7 | 0.37546371 | 0.26417164 | 0.39667074 |
| sp\|A0A075B6 | 127433906 | 152765865 | 0.16287556 | 0.26417164 | 0.39667074 |
| sp\|Q6ZVF9\|G | 22503995.7 | 31767981.2 | 0.35031357 | 0.26417164 | 0.39667074 |
| sp\|P35625\|T | 4656488.2 | 4579852.22 | -0.0673657 | 0.26417164 | 0.39667074 |
| sp\|Q9NTX7\|F | 2568282.5 | 2914446.14 | 0.10104072 | 0.26417164 | 0.39667074 |
| sp\|Q96KN2\|( | 13311265.2 | 15855706.5 | 0.18192587 | 0.26417164 | 0.39667074 |
| sp\|P48634\|P | 9944776.41 | 11494293.2 | 0.13649196 | 0.26417164 | 0.39667074 |
| sp\|P20073\|A | 45066783.7 | 49585859.5 | 0.1066855 | 0.26417164 | 0.39667074 |
| sp\|Q9BUP0\|I | 6763030.85 | 6483917.23 | -0.1410574 | 0.26417164 | 0.39667074 |
| sp\|O96005\|( | 5291740.29 | 3940222.61 | -0.8508492 | 0.26417164 | 0.39667074 |
| sp\|Q86YW9\| | 1107617.99 | 1298896.85 | 0.38497323 | 0.26417164 | 0.39667074 |
| sp\|Q86UE4\|L | 22310103 | 24659946.2 | 0.11645049 | 0.26417164 | 0.39667074 |
| sp\|B5ME19\|E | 9956647.21 | 8896592.52 | -0.1483938 | 0.26417164 | 0.39667074 |
| sp\|P55327\|T | 6667592.13 | 7206437.77 | 0.08455591 | 0.26417164 | 0.39667074 |
| sp\|P09661\|R | 12199081.5 | 11541237.8 | -0.1101318 | 0.27407142 | 0.40834174 |

| | | | | | |
|---|---|---|---|---|---|
| sp\|Q15126\|F | 7836494.49 | 8705258.04 | 0.1367259 | 0.27407142 | 0.40834174 |
| sp\|Q9UNH7\| | 19945823.3 | 21545241.8 | 0.11598998 | 0.27407142 | 0.40834174 |
| sp\|Q9Y6G9\|[ | 2035147.57 | 2285417.97 | 0.17169306 | 0.27407142 | 0.40834174 |
| sp\|P62491\|R | 202667267 | 215981316 | 0.06973841 | 0.27407142 | 0.40834174 |
| sp\|P33176\|K | 41106829.3 | 44666900.5 | 0.10557327 | 0.27407142 | 0.40834174 |
| sp\|Q53H96\|I | 7334135.37 | 7201826.87 | -0.1776374 | 0.27407142 | 0.40834174 |
| sp\|O14613\|E | 4282770.42 | 4868687.45 | 0.18341263 | 0.27407142 | 0.40834174 |
| sp\|Q53T59\|H | 15510240.7 | 14591348.4 | -0.0994697 | 0.27407142 | 0.40834174 |
| sp\|Q9H307\|I | 792492.101 | 1384303.9 | -0.5575937 | 0.27407142 | 0.40834174 |
| sp\|O14579\|C | 13828324.8 | 15101545.2 | 0.08824031 | 0.27407142 | 0.40834174 |
| sp\|P35558\|P | 85827870.7 | 114331429 | 0.20682758 | 0.27407142 | 0.40834174 |
| sp\|Q14258\|T | 15826360.6 | 14327685.2 | -0.1558955 | 0.27407142 | 0.40834174 |
| sp\|P07205\|P | 741955845 | 692870196 | -0.0785769 | 0.27407142 | 0.40834174 |
| sp\|Q00G26\|I | 6872456.49 | 7961467.51 | 0.15098801 | 0.27407142 | 0.40834174 |
| sp\|O60884\|D | 17995145.1 | 16523490.6 | -0.1346329 | 0.27407142 | 0.40834174 |
| sp\|Q96AT9\|F | 42452060 | 39466309.5 | -0.1312278 | 0.27407142 | 0.40834174 |
| sp\|O00571\|D | 12272966.7 | 13920205.6 | 0.15052943 | 0.27407142 | 0.40834174 |
| sp\|O95336\|6 | 328045499 | 310904978 | -0.0990814 | 0.27407142 | 0.40834174 |
| sp\|Q9BZL1\|U | 7793574.54 | 6467635.05 | -0.3213437 | 0.27407142 | 0.40834174 |
| sp\|Q92575\|U | 741792.796 | 662599.55 | -0.1962495 | 0.27407142 | 0.40834174 |
| sp\|P28161\|G | 287605928 | 279524282 | -0.3062878 | 0.27407142 | 0.40834174 |
| sp\|Q8WWM7 | 4182611.81 | 4512646.52 | 0.08861309 | 0.27407142 | 0.40834174 |
| sp\|P62258\|1 | 1036025797 | 1089574762 | 0.05414122 | 0.27407142 | 0.40834174 |
| sp\|P59666\|D | 52705093.3 | 189997003 | 0.33777916 | 0.27407142 | 0.40834174 |
| sp\|Q9Y265\|F | 60993514.2 | 56435653.5 | -0.1932403 | 0.28421335 | 0.41890045 |
| sp\|P11908\|P | 62085517.5 | 60020848 | -0.1042488 | 0.28421335 | 0.41890045 |
| sp\|Q13217\|D | 4097093.66 | 4491886.6 | 0.13465881 | 0.28421335 | 0.41890045 |
| sp\|P78330\|S | 1037270.24 | 1008827.28 | -0.3531125 | 0.28421335 | 0.41890045 |
| sp\|Q9HB40\|I | 31568819.9 | 36540569.4 | 0.13248163 | 0.28421335 | 0.41890045 |
| sp\|Q9H4A3\|V | 3729969.99 | 3305601.64 | -0.2083015 | 0.28421335 | 0.41890045 |
| sp\|P48728\|G | 718210.658 | 644466.757 | -0.1952544 | 0.28421335 | 0.41890045 |
| sp\|O14936\|C | 52999920.5 | 64225197.9 | 0.37029341 | 0.28421335 | 0.41890045 |
| sp\|A0A0B4J2 | 17954825.2 | 19700136.5 | 0.48150053 | 0.28421335 | 0.41890045 |
| sp\|A0A0C4DI | 61508491.4 | 67585902.6 | 0.17913195 | 0.28421335 | 0.41890045 |
| sp\|A0A0B4J1 | 973020491 | 1071327372 | 0.15660583 | 0.28421335 | 0.41890045 |
| sp\|Q9Y3C6\|F | 212843594 | 188370094 | -0.232248 | 0.28421335 | 0.41890045 |
| sp\|Q9H6R3\|A | 71159338 | 76124906.9 | 0.04727888 | 0.28421335 | 0.41890045 |
| sp\|O75306\|N | 15854471.9 | 17858978.1 | 0.15241937 | 0.28421335 | 0.41890045 |
| sp\|Q9H892\|T | 18382056.5 | 17296344.5 | -0.1176127 | 0.28421335 | 0.41890045 |

| | | | | | |
|---|---|---|---|---|---|
| sp\|P24752\|T | 362489833 | 335186937 | -0.1109557 | 0.28421335 | 0.41890045 |
| sp\|Q9H772\|( | 1533817.96 | 2158439.42 | 0.4908056 | 0.28421335 | 0.41890045 |
| sp\|Q9HCJ6\|\ | 5380549.24 | 6154726.99 | 0.18000254 | 0.28421335 | 0.41890045 |
| sp\|Q01085\|T | 38835811.9 | 41487154.8 | 0.099129 | 0.28421335 | 0.41890045 |
| sp\|O60888\|( | 32489208.3 | 36927512.7 | 0.13375165 | 0.28421335 | 0.41890045 |
| sp\|P06310\|K | 135130184 | 159605584 | 0.18663437 | 0.28421335 | 0.41890045 |
| sp\|Q9UNF1\|I | 8623269.77 | 9392481.7 | 0.10887266 | 0.28421335 | 0.41890045 |
| sp\|Q96QC0\| | 6634993.83 | 4752725.24 | -0.5548143 | 0.28421335 | 0.41890045 |
| sp\|P30085\|K | 120376200 | 131214848 | 0.09206867 | 0.28421335 | 0.41890045 |
| sp\|P04732\|M | 13404147.8 | 17658576 | 0.15004873 | 0.28421335 | 0.41890045 |
| sp\|Q14019\|( | 81623057.8 | 92532521 | 0.11583861 | 0.28421335 | 0.41890045 |
| sp\|Q9H0R6\|( | 1234341.17 | 1333702.59 | -0.3624411 | 0.28421335 | 0.41890045 |
| sp\|Q9NX57\|I | 3928506.32 | 4275191.27 | 0.10055071 | 0.28421335 | 0.41890045 |
| sp\|P52298\|N | 40056339.6 | 31675468.3 | -0.3354023 | 0.28421335 | 0.41890045 |
| sp\|P08133\|A | 392314792 | 366968195 | -0.1977607 | 0.28421335 | 0.41890045 |
| sp\|Q9BZK3\|N | 9782538.43 | 9767644.59 | -0.0505039 | 0.28421335 | 0.41890045 |
| sp\|Q5D862\|I | 254422.98 | 263861.376 | -0.0040884 | 0.28421335 | 0.41890045 |
| sp\|O94856\|N | 1668771.54 | 1969349.14 | 0.18971891 | 0.28421335 | 0.41890045 |
| sp\|P22059\|C | 1871265.49 | 2274341.21 | 0.07151241 | 0.28421335 | 0.41890045 |
| sp\|Q9GZV5\|\ | 4503630.42 | 4887433.82 | 0.07294671 | 0.28421335 | 0.41890045 |
| sp\|P05155\|I( | 1375673064 | 1544074561 | 0.14780671 | 0.29459803 | 0.43050424 |
| sp\|O43847\|N | 37623894.3 | 40788742.9 | 0.07257029 | 0.29459803 | 0.43050424 |
| sp\|P01011\|A | 869484061 | 751173294 | -0.1829219 | 0.29459803 | 0.43050424 |
| sp\|Q9H6Z4\|I | 18086904.2 | 17313261.7 | -0.078398 | 0.29459803 | 0.43050424 |
| sp\|Q6ICL3\|T | 35333908.6 | 47202984.4 | 0.22778265 | 0.29459803 | 0.43050424 |
| sp\|P08582\|T | 2815884.51 | 3138661.28 | 0.14441527 | 0.29459803 | 0.43050424 |
| sp\|Q6DD88\|/ | 14665183 | 16376573.4 | 0.06083753 | 0.29459803 | 0.43050424 |
| sp\|O95630\|S | 21176480.9 | 23719194.7 | 0.12796012 | 0.29459803 | 0.43050424 |
| sp\|P61020\|R | 38400905.2 | 41246102.5 | 0.09347349 | 0.29459803 | 0.43050424 |
| sp\|O94966\|U | 484519.481 | 565674.642 | 0.20040588 | 0.29459803 | 0.43050424 |
| sp\|P80511\|S | 492510.715 | 1117828.96 | 0.38554133 | 0.29459803 | 0.43050424 |
| sp\|P82909\|R | 14849378.2 | 16368161.7 | 0.23096407 | 0.29459803 | 0.43050424 |
| sp\|Q9ULE6\|I | 7504003.88 | 6697615.34 | -0.2433005 | 0.29459803 | 0.43050424 |
| sp\|Q9BYD6\|I | 1466199.69 | 1744989.09 | 0.20886373 | 0.29459803 | 0.43050424 |
| sp\|P35573\|G | 328588521 | 364262067 | 0.04688829 | 0.29459803 | 0.43050424 |
| sp\|P53804\|T | 8587798.69 | 7375020.95 | -0.1280115 | 0.29459803 | 0.43050424 |
| sp\|Q16363\|L | 675972264 | 635598219 | -0.1101417 | 0.29459803 | 0.43050424 |
| sp\|O75489\|N | 17490233.2 | 16372764.9 | -0.0924021 | 0.29459803 | 0.43050424 |
| sp\|Q9NPH0\| | 3375100.08 | 7294284.29 | 0.68259374 | 0.29459803 | 0.43050424 |

| | | | | | |
|---|---|---|---|---|---|
| sp\|Q9NP98\|[illegible] | 22374615.1 | 30126566.8 | 0.24520908 | 0.29459803 | 0.43050424 |
| sp\|P46736\|B | 47970358 | 45463717.3 | -0.1357268 | 0.29459803 | 0.43050424 |
| sp\|P12268\|I[illegible] | 90650229.2 | 96970406.1 | 0.11208758 | 0.29459803 | 0.43050424 |
| sp\|P14136\|G | 1060728285 | 1139451795 | 0.08511758 | 0.29459803 | 0.43050424 |
| sp\|Q3KQV9\|[illegible] | 15074968.4 | 17064028.9 | 0.19231872 | 0.29459803 | 0.43050424 |
| sp\|Q13098\|[illegible] | 186590.126 | 203962.425 | 0.1946468 | 0.29459803 | 0.43050424 |
| sp\|Q9UK55\|[illegible] | 7178195.71 | 10260924.8 | 0.18179086 | 0.29459803 | 0.43050424 |
| sp\|Q86UQ4\|[illegible] | 5329593.96 | 6579938.28 | 0.33008711 | 0.29459803 | 0.43050424 |
| sp\|Q9UIQ6\|L | 25081942.2 | 23665508.7 | -0.0901759 | 0.29459803 | 0.43050424 |
| sp\|Q14699\|F | 8661204.98 | 7809484.43 | -0.1566355 | 0.30522566 | 0.44173039 |
| sp\|P05114\|H | 7089131.41 | 6418276.4 | -0.2033341 | 0.30522566 | 0.44173039 |
| sp\|Q02985\|F | 59841355 | 69687490.5 | 0.15701455 | 0.30522566 | 0.44173039 |
| sp\|Q9H2G2\|[illegible] | 4195041.72 | 3827616.68 | -0.1935684 | 0.30522566 | 0.44173039 |
| sp\|Q96AT1\|K | 3026833.74 | 3246373.24 | 0.12545189 | 0.30522566 | 0.44173039 |
| sp\|Q9BQA1\|[illegible] | 36584090.2 | 32780814.8 | -0.2516492 | 0.30522566 | 0.44173039 |
| sp\|P01718\|L | 29698661.6 | 33081108.8 | 0.17698123 | 0.30522566 | 0.44173039 |
| sp\|P02774\|V | 2167563745 | 2429446625 | 0.13621701 | 0.30522566 | 0.44173039 |
| sp\|P26038\|M | 2194735941 | 1950178405 | -0.291502 | 0.30522566 | 0.44173039 |
| sp\|P28838\|A | 702456470 | 660634585 | -0.1350924 | 0.30522566 | 0.44173039 |
| sp\|P05160\|F | 85905447.3 | 111186082 | 0.21001335 | 0.30522566 | 0.44173039 |
| sp\|Q86UP2\|[illegible] | 116819658 | 121566156 | 0.04461944 | 0.30522566 | 0.44173039 |
| sp\|P01593\|K | 35537318.7 | 36163827.9 | 0.13341879 | 0.30522566 | 0.44173039 |
| sp\|Q08380\|L | 192596778 | 196877453 | 0.04083364 | 0.30522566 | 0.44173039 |
| sp\|P09871\|C | 302708567 | 271100574 | -0.3324852 | 0.30522566 | 0.44173039 |
| sp\|Q15582\|E | 165206881 | 200091682 | 0.13274487 | 0.30522566 | 0.44173039 |
| sp\|Q86XN8\|[illegible] | 9659515.35 | 8296301.41 | -0.4225679 | 0.30522566 | 0.44173039 |
| sp\|Q96AB3\|[illegible] | 78057528.5 | 80841627.6 | 0.04378401 | 0.30522566 | 0.44173039 |
| sp\|Q8NE71\|[illegible] | 9355609.26 | 8668412.24 | -0.1369776 | 0.30522566 | 0.44173039 |
| sp\|Q9HD47\| | 1646576.63 | 1778962.34 | 0.10737947 | 0.30522566 | 0.44173039 |
| sp\|Q9BWS9\| | 9484732.47 | 8490730.57 | -0.2179245 | 0.30522566 | 0.44173039 |
| sp\|P49585\|P | 799578.558 | 888092.183 | 0.17622332 | 0.30522566 | 0.44173039 |
| sp\|Q8N0U8\|[illegible] | 22020414.4 | 20177039.2 | -0.2054098 | 0.30522566 | 0.44173039 |
| sp\|P13807\|G | 45981203.4 | 42426994.4 | -0.1213028 | 0.30522566 | 0.44173039 |
| sp\|Q9H492\|[illegible] | 9270019.49 | 8624015.54 | -0.155892 | 0.30522566 | 0.44173039 |
| sp\|O95260\|[illegible] | 11768903 | 10908371 | -0.1296064 | 0.30522566 | 0.44173039 |
| sp\|P19823\|I[illegible] | 972637236 | 920662175 | -0.4375597 | 0.30522566 | 0.44173039 |
| sp\|Q12860\|[illegible] | 486237.699 | 443060.844 | -0.1870909 | 0.30522566 | 0.44173039 |
| sp\|O14807\|F | 10356876.9 | 11591214.3 | 0.1082616 | 0.30522566 | 0.44173039 |
| sp\|Q8N635\|[illegible] | 11583314.2 | 9851473.6 | -0.235664 | 0.30522566 | 0.44173039 |

| | | | | | |
|---|---|---|---|---|---|
| sp\|O75914\|F | 8763468.23 | 9708805.87 | 0.13330666 | 0.30522566 | 0.44173039 |
| sp\|P50336\|P | 1624348.87 | 1443633.8 | -0.2570935 | 0.30522566 | 0.44173039 |
| sp\|P68363\|T | 538838885 | 597556452 | 0.07161636 | 0.31609626 | 0.45322556 |
| sp\|Q9UKM7\| | 1068298.08 | 930006.033 | -0.1405867 | 0.31609626 | 0.45322556 |
| sp\|Q9BSH4\|T | 29814031.8 | 25453275.8 | -0.4246375 | 0.31609626 | 0.45322556 |
| sp\|Q8N766\|E | 5857180.79 | 5533285.92 | -0.1049963 | 0.31609626 | 0.45322556 |
| sp\|Q8N9N7\| | 12792865 | 14421106.2 | 0.03255961 | 0.31609626 | 0.45322556 |
| sp\|P46439\|G | 129497873 | 139715904 | -0.1229935 | 0.31609626 | 0.45322556 |
| sp\|Q07075\|A | 3905404.92 | 5238055.82 | 0.29867586 | 0.31609626 | 0.45322556 |
| sp\|P0CG39\|F | 3646180518 | 4223412188 | 0.14780615 | 0.31609626 | 0.45322556 |
| sp\|P53992\|S | 4149933.17 | 3912557.41 | -0.1076907 | 0.31609626 | 0.45322556 |
| sp\|P20827\|E | 492910.925 | 416324.27 | -1.5889648 | 0.31609626 | 0.45322556 |
| sp\|Q8IWI9\|M | 60661961.1 | 82463420.7 | 0.47790571 | 0.31609626 | 0.45322556 |
| sp\|Q99426\|T | 46896055.8 | 41662157 | -0.2056645 | 0.31609626 | 0.45322556 |
| sp\|Q9BWP8\| | 778557.68 | 889348.458 | 0.17217059 | 0.31609626 | 0.45322556 |
| sp\|O14530\|T | 171818.82 | 134843.259 | 0.16051081 | 0.31609626 | 0.45322556 |
| sp\|Q9NXG0\| | 37642319.4 | 42825235.6 | 0.07243644 | 0.31609626 | 0.45322556 |
| sp\|P36959\|G | 62500898.6 | 84907948.4 | 0.26829413 | 0.31609626 | 0.45322556 |
| sp\|P32321\|D | 26153359.6 | 26981682 | 0.07142159 | 0.31609626 | 0.45322556 |
| sp\|Q96I24\|F | 114717857 | 105972748 | -0.1182122 | 0.31609626 | 0.45322556 |
| sp\|Q9Y4F1\|F | 4636743.43 | 5509092.59 | 0.20292795 | 0.31609626 | 0.45322556 |
| sp\|O43488\|A | 251592595 | 240295376 | -0.1023232 | 0.31609626 | 0.45322556 |
| sp\|Q9Y3B3\|T | 11679088.9 | 12782790.1 | 0.04482839 | 0.31609626 | 0.45322556 |
| sp\|O43633\|C | 4340778.26 | 4899209.3 | 0.14940097 | 0.31609626 | 0.45322556 |
| sp\|Q14697\|C | 921180127 | 811844665 | -0.3026884 | 0.31609626 | 0.45322556 |
| sp\|Q14696\|M | 22980709.2 | 25084181.9 | 0.06267487 | 0.31609626 | 0.45322556 |
| sp\|Q15262\|F | 2450396.68 | 2777757.98 | 0.13118366 | 0.31609626 | 0.45322556 |
| sp\|O00505\|I | 304799.549 | 336665.937 | -0.10003 | 0.31609626 | 0.45322556 |
| sp\|Q6P1N0\|C | 7432121.69 | 8119484.56 | 0.11486956 | 0.31609626 | 0.45322556 |
| sp\|Q99972\|M | 1171082.32 | 1649886.27 | 0.20171051 | 0.31609626 | 0.45322556 |
| sp\|Q9H8Y8\|C | 28636044.2 | 30452981.4 | 0.02498829 | 0.31609626 | 0.45322556 |
| sp\|Q70E73\|F | 4788336.91 | 5746610.36 | 0.30899184 | 0.31609626 | 0.45322556 |
| sp\|Q8WW12 | 24254826.4 | 23302342 | -0.1511771 | 0.31609626 | 0.45322556 |
| sp\|P48729\|K | 3076118.04 | 2828372.3 | -0.3302813 | 0.32720971 | 0.46540587 |
| sp\|P23470\|P | 4816212.32 | 5232428.62 | 0.14552778 | 0.32720971 | 0.46540587 |
| sp\|Q15413\|F | 1038404.96 | 1220983.52 | 0.27593418 | 0.32720971 | 0.46540587 |
| sp\|P48507\|G | 17919350.4 | 23157195.4 | 0.11753909 | 0.32720971 | 0.46540587 |
| sp\|O95428\|F | 17427404.2 | 15547955.1 | -0.1332418 | 0.32720971 | 0.46540587 |
| sp\|P20839\|II | 43063997.8 | 46060623.8 | 0.07905628 | 0.32720971 | 0.46540587 |

| | | | | | |
|---|---|---|---|---|---|
| sp\|Q9UPY8\|I | 1472840.45 | 1985156.98 | -0.0489942 | 0.32720971 | 0.46540587 |
| sp\|P53004\|B | 15544012.2 | 18921846.3 | 0.18583011 | 0.32720971 | 0.46540587 |
| sp\|P54108\|C | 4931822.45 | 7711436.18 | 0.25897339 | 0.32720971 | 0.46540587 |
| sp\|O75223\|( | 56795102.6 | 63216890 | 0.10455181 | 0.32720971 | 0.46540587 |
| sp\|P28065\|P | 51994343.5 | 47391673.4 | -0.2236743 | 0.32720971 | 0.46540587 |
| sp\|Q9UN74\| | 558468.597 | 504741.89 | -0.1926527 | 0.32720971 | 0.46540587 |
| sp\|Q05048\|( | 3012162.19 | 2755441.41 | -0.1497268 | 0.32720971 | 0.46540587 |
| sp\|P00558\|P | 3967550000 | 3752451622 | -0.0603552 | 0.32720971 | 0.46540587 |
| sp\|Q6GMV2\| | 7301954.12 | 6795138.98 | -0.1685971 | 0.32720971 | 0.46540587 |
| sp\|P09417\|D | 604111492 | 552871370 | -0.2337638 | 0.32720971 | 0.46540587 |
| sp\|Q68DQ2\| | 6567770.65 | 6040485.13 | -0.1336285 | 0.32720971 | 0.46540587 |
| sp\|Q16854\|D | 3965951.11 | 3432737.31 | -0.1986046 | 0.32720971 | 0.46540587 |
| sp\|A0A075B6 | 25875277.9 | 31662543.1 | 0.13836862 | 0.32720971 | 0.46540587 |
| sp\|A0A075B6 | 977859.984 | 1818638.63 | 0.42748593 | 0.32720971 | 0.46540587 |
| sp\|A0AV96\|R | 579717.193 | 482096.157 | -0.1908408 | 0.32720971 | 0.46540587 |
| sp\|P35813\|P | 53675119.7 | 54070989 | -0.1271188 | 0.32720971 | 0.46540587 |
| sp\|Q3V6T2\|( | 6130985.51 | 7792764.03 | 0.30978591 | 0.32720971 | 0.46540587 |
| sp\|Q9Y6X5\|E | 6081244.2 | 6154199.66 | -0.1523269 | 0.32720971 | 0.46540587 |
| sp\|Q9NR16\|( | 27446727.5 | 29671441.9 | 0.07150277 | 0.32720971 | 0.46540587 |
| sp\|Q9Y5X3\|S | 14300301.7 | 13338748.3 | -0.1049496 | 0.32720971 | 0.46540587 |
| sp\|P02545\|L | 1674172758 | 1730622307 | 0.13930106 | 0.32720971 | 0.46540587 |
| sp\|P46977\|S | 1629695.04 | 1744983.65 | -0.1909227 | 0.33856552 | 0.47773477 |
| sp\|Q53H47\|S | 406741.511 | 439241.251 | 0.12660595 | 0.33856552 | 0.47773477 |
| sp\|Q8IZP0\|A | 10502976.2 | 11436141.2 | 0.0782165 | 0.33856552 | 0.47773477 |
| sp\|P05166\|P | 322384110 | 295646748 | -0.1567259 | 0.33856552 | 0.47773477 |
| sp\|P09038\|F | 3699391.57 | 3329587.41 | -0.1426933 | 0.33856552 | 0.47773477 |
| sp\|O94903\|F | 88045400.1 | 81655168.7 | -0.1982889 | 0.33856552 | 0.47773477 |
| sp\|P34810\|C | 584796.889 | 1425124.08 | -0.2987218 | 0.33856552 | 0.47773477 |
| sp\|O43760\|S | 20548068.7 | 23267365.3 | 0.10847649 | 0.33856552 | 0.47773477 |
| sp\|Q6WKZ4\| | 8819209.91 | 7522951.05 | -0.2206703 | 0.33856552 | 0.47773477 |
| sp\|Q14165\|I | 6946703.06 | 6625731.69 | -0.1363689 | 0.33856552 | 0.47773477 |
| sp\|Q9UJC3\|H | 2774090.89 | 2742000.78 | 0.07352175 | 0.33856552 | 0.47773477 |
| sp\|Q9BUF5\|T | 495839119 | 567108603 | 0.11616361 | 0.33856552 | 0.47773477 |
| sp\|O75110\|A | 1181896.19 | 998843.3 | -0.2767678 | 0.33856552 | 0.47773477 |
| sp\|P36955\|P | 83443251.3 | 95024984.3 | 0.25461413 | 0.33856552 | 0.47773477 |
| sp\|O60238\|E | 1024822.47 | 898679.388 | 0.03919163 | 0.33856552 | 0.47773477 |
| sp\|P41732\|T | 1723608.64 | 1798535.09 | 0.06087184 | 0.33856552 | 0.47773477 |
| sp\|Q6XE24\|F | 13507527.4 | 12735293.5 | -0.114238 | 0.33856552 | 0.47773477 |
| sp\|P14317\|H | 79254682.6 | 91412673.5 | 0.0925486 | 0.33856552 | 0.47773477 |

| | | | | | |
|---|---|---|---|---|---|
| sp\|P18428\|L | 335600.708 | 494576.376 | 0.25960576 | 0.33856552 | 0.47773477 |
| sp\|Q6ZV29\|F | 5523057.15 | 11738419.9 | 0.55404795 | 0.33856552 | 0.47773477 |
| sp\|Q2KHR2\|I | 10591341 | 17355096.1 | -0.2849022 | 0.33856552 | 0.47773477 |
| sp\|P63000\|R | 183217136 | 217411851 | 0.17135117 | 0.33856552 | 0.47773477 |
| sp\|Q05639\|E | 667547175 | 621295558 | -0.1297095 | 0.33856552 | 0.47773477 |
| sp\|Q9P2E9\|F | 185346432 | 209082627 | 0.1355133 | 0.33856552 | 0.47773477 |
| sp\|P14207\|F | 8616152.29 | 12259940.7 | 0.14121702 | 0.33856552 | 0.47773477 |
| sp\|P04259\|K | 1307151944 | 876414797 | -0.3312743 | 0.33856552 | 0.47773477 |
| sp\|Q9NVT9\|A | 2020283.2 | 3067357.99 | -0.6810349 | 0.33856552 | 0.47773477 |
| sp\|O75821\|E | 10155861.7 | 9373117.41 | -0.2150288 | 0.35016296 | 0.48906637 |
| sp\|Q96GD0\| | 133534891 | 153233955 | 0.14606924 | 0.35016296 | 0.48906637 |
| sp\|P17931\|L | 348863066 | 332906672 | -0.0902735 | 0.35016296 | 0.48906637 |
| sp\|Q92619\|H | 982301.373 | 1186743.69 | 0.18501642 | 0.35016296 | 0.48906637 |
| sp\|P48740\|M | 22030491.2 | 21408358.9 | -0.1211169 | 0.35016296 | 0.48906637 |
| sp\|Q9BXR6\|F | 14895478.2 | 14422304.6 | -0.2010403 | 0.35016296 | 0.48906637 |
| sp\|Q07960\|F | 13022308.5 | 14884841.2 | 0.08229453 | 0.35016296 | 0.48906637 |
| sp\|Q16527\|C | 66898687.4 | 55862156 | -0.2152093 | 0.35016296 | 0.48906637 |
| sp\|Q93034\|C | 712078.104 | 578764.527 | 0.19052634 | 0.35016296 | 0.48906637 |
| sp\|Q86U44\|N | 754466.924 | 680431.678 | -0.0631913 | 0.35016296 | 0.48906637 |
| sp\|Q8ND94\| | 6468977.35 | 6052832.45 | -0.1618773 | 0.35016296 | 0.48906637 |
| sp\|Q14554\|F | 2708725.82 | 2453021.99 | -0.1961669 | 0.35016296 | 0.48906637 |
| sp\|P53999\|T | 32668004.2 | 28939502.9 | -0.1918509 | 0.35016296 | 0.48906637 |
| sp\|Q96K76\|U | 4472941.62 | 4200436 | -0.0856019 | 0.35016296 | 0.48906637 |
| sp\|Q8TAT6\|N | 25988600.8 | 27800723.3 | 0.05198899 | 0.35016296 | 0.48906637 |
| sp\|P26022\|P | 3669091.87 | 4568365.28 | 0.03930858 | 0.35016296 | 0.48906637 |
| sp\|O60749\|S | 18600998.6 | 17606083.7 | -0.1049063 | 0.35016296 | 0.48906637 |
| sp\|Q9Y5C1\|A | 4638814.13 | 3955042.68 | -0.1966691 | 0.35016296 | 0.48906637 |
| sp\|P60510\|P | 13367916.5 | 14733440.7 | 0.08510453 | 0.35016296 | 0.48906637 |
| sp\|Q96K17\|E | 23898235 | 25495777.3 | 0.08581089 | 0.35016296 | 0.48906637 |
| sp\|P01344\|I | 8775060.16 | 10254050.1 | 0.07106777 | 0.35016296 | 0.48906637 |
| sp\|Q16539\|N | 12142176.5 | 11344369.5 | -0.199387 | 0.35016296 | 0.48906637 |
| sp\|P51692\|S | 1977546.96 | 2301839.39 | 0.14222282 | 0.35016296 | 0.48906637 |
| sp\|P22891\|P | 3988854.58 | 4390107.37 | -0.1554223 | 0.35016296 | 0.48906637 |
| sp\|Q5UCC4\| | 1946212.88 | 2004778.09 | 0.30805187 | 0.35016296 | 0.48906637 |
| sp\|Q8NC56\| | 754179.426 | 646246.726 | -0.4702489 | 0.35016296 | 0.48906637 |
| sp\|Q9UQE7\| | 2491516.77 | 2205752.53 | -0.3500025 | 0.35016296 | 0.48906637 |
| sp\|P49406\|R | 12323973.3 | 12022262.1 | -0.1503928 | 0.35016296 | 0.48906637 |
| sp\|Q96MG8\| | 147131.195 | 126580.675 | -0.4443167 | 0.35016296 | 0.48906637 |
| sp\|P04217\|A | 1253557208 | 1146268168 | -0.331233 | 0.35016296 | 0.48906637 |

| | | | | | |
|---|---|---|---|---|---|
| sp\|P62979\|R | 713436361 | 682426537 | -0.1135079 | 0.35016296 | 0.48906637 |
| sp\|P04440\|D | 4535023.22 | 4606588.03 | 0.15125234 | 0.35016296 | 0.48906637 |
| sp\|Q9BQE3\|T | 501210006 | 556265538 | 0.07469841 | 0.35016296 | 0.48906637 |
| sp\|Q8N5M1\| | 2025520.8 | 2168258.6 | 0.07650366 | 0.35016296 | 0.48906637 |
| sp\|O14786\|N | 9903910.73 | 9299682.65 | -0.135281 | 0.35016296 | 0.48906637 |
| sp\|Q9UBY9\|I | 83393365 | 88395162.5 | 0.07197474 | 0.36200119 | 0.50165859 |
| sp\|O60760\|H | 12371142.8 | 11068100.9 | -0.3395841 | 0.36200119 | 0.50165859 |
| sp\|Q9NPH3\| | 59443006.8 | 65715576.3 | 0.03215146 | 0.36200119 | 0.50165859 |
| sp\|P13727\|P | 1562775.93 | 1905670.85 | 0.11142038 | 0.36200119 | 0.50165859 |
| sp\|Q8N8N7\| | 56991583.9 | 64627253.4 | 0.04556712 | 0.36200119 | 0.50165859 |
| sp\|O75937\|D | 148881760 | 142528467 | -0.087561 | 0.36200119 | 0.50165859 |
| sp\|Q08J23\|N | 2436335.94 | 2179978.97 | -0.2549128 | 0.36200119 | 0.50165859 |
| sp\|O75475\|P | 28211069.9 | 26328317.7 | -0.118564 | 0.36200119 | 0.50165859 |
| sp\|Q14444\|C | 23710328.9 | 24797254.3 | 0.04443289 | 0.36200119 | 0.50165859 |
| sp\|P31689\|D | 7499801.6 | 6922793.7 | -0.1378507 | 0.36200119 | 0.50165859 |
| sp\|Q9Y2W2\|T | 5190493.37 | 4732125.08 | -0.2173752 | 0.36200119 | 0.50165859 |
| sp\|Q8TC27\|A | 12067803.6 | 14789718.1 | 0.19146793 | 0.36200119 | 0.50165859 |
| sp\|O43866\|C | 158352008 | 187221243 | 0.13577504 | 0.36200119 | 0.50165859 |
| sp\|O43312\|N | 472860.43 | 625495.154 | 0.69814963 | 0.36200119 | 0.50165859 |
| sp\|P01706\|L | 81503281.3 | 86720395 | 0.15657297 | 0.36200119 | 0.50165859 |
| sp\|Q8WWV3 | 630574.726 | 558979.882 | -0.3243655 | 0.36200119 | 0.50165859 |
| sp\|Q15459\|S | 35112192.3 | 37532006.9 | 0.0688641 | 0.36200119 | 0.50165859 |
| sp\|Q8TAE8\|G | 4593686.66 | 4005999.85 | -0.2732676 | 0.36200119 | 0.50165859 |
| sp\|P37198\|N | 726717.332 | 840657.68 | -0.462719 | 0.36200119 | 0.50165859 |
| sp\|Q86X55\|C | 3771499.41 | 4553553.96 | 0.06765842 | 0.36200119 | 0.50165859 |
| sp\|Q13885\|T | 726363459 | 844709407 | 0.14173931 | 0.36200119 | 0.50165859 |
| sp\|Q13868\|E | 22823714.8 | 22065388.5 | -0.1190698 | 0.36200119 | 0.50165859 |
| sp\|Q9BQE9\|I | 937922.314 | 1302001.43 | 0.34526408 | 0.36200119 | 0.50165859 |
| sp\|Q9NUL5\|S | 11327546.9 | 10704511 | -0.1549648 | 0.36200119 | 0.50165859 |
| sp\|P62826\|R | 690694433 | 760785864 | 0.12126775 | 0.36200119 | 0.50165859 |
| sp\|P02511\|C | 3320624576 | 3733888302 | 0.39621292 | 0.36200119 | 0.50165859 |
| sp\|P01920\|D | 588222.753 | 769540.1 | 0.31821007 | 0.36200119 | 0.50165859 |
| sp\|Q96EQ0\|S | 1165742.71 | 1901043.69 | -0.6387452 | 0.37407896 | 0.51276919 |
| sp\|A0A0A0MS | 150249811 | 160749121 | 0.11545955 | 0.37407896 | 0.51276919 |
| sp\|O43852\|C | 31872034.5 | 38016010.9 | -0.1182092 | 0.37407896 | 0.51276919 |
| sp\|Q14839\|C | 6121709.61 | 6083215.28 | -0.1086069 | 0.37407896 | 0.51276919 |
| sp\|P31153\|M | 153164583 | 176265948 | 0.05731739 | 0.37407896 | 0.51276919 |
| sp\|Q14203\|D | 57574755.1 | 60069870.2 | 0.04830317 | 0.37407896 | 0.51276919 |
| sp\|Q9NVZ3\|N | 834715.665 | 722055.31 | -0.4532848 | 0.37407896 | 0.51276919 |

| | | | | | |
|---|---|---|---|---|---|
| sp\|P31751\|A | 6327991.31 | 5663757.72 | -0.3039967 | 0.37407896 | 0.51276919 |
| sp\|O15511\|A | 86367405.2 | 93496599.9 | 0.09720703 | 0.37407896 | 0.51276919 |
| sp\|P23634\|A | 60449459.1 | 57298432.3 | -0.0817669 | 0.37407896 | 0.51276919 |
| sp\|Q9NX20\|I | 4244922.82 | 3739146.46 | -0.2010906 | 0.37407896 | 0.51276919 |
| sp\|Q9NZP8\|C | 63769296.2 | 59528774.6 | -0.1823323 | 0.37407896 | 0.51276919 |
| sp\|P19878\|N | 60289664.4 | 72938031 | 0.1749489 | 0.37407896 | 0.51276919 |
| sp\|Q9Y2H0\|I | 9435919.69 | 8856643.15 | -0.1459534 | 0.37407896 | 0.51276919 |
| sp\|Q96SZ5\|A | 8212672.73 | 7797032.76 | -0.0620732 | 0.37407896 | 0.51276919 |
| sp\|Q92504\|S | 370232.08 | 383911.73 | -0.2605272 | 0.37407896 | 0.51276919 |
| sp\|P15151\|P | 258244.944 | 215335.315 | 0.3627172 | 0.37407896 | 0.51276919 |
| sp\|Q9P015\|F | 19188553.6 | 18044103.7 | -0.1873541 | 0.37407896 | 0.51276919 |
| sp\|P13929\|E | 986338696 | 851862016 | 0.11396059 | 0.37407896 | 0.51276919 |
| sp\|P48681\|N | 115147638 | 122647866 | 0.07959214 | 0.37407896 | 0.51276919 |
| sp\|O95208\|E | 640450.051 | 627939.435 | -0.0709121 | 0.37407896 | 0.51276919 |
| sp\|Q9UBW8\| | 5849230.27 | 6371309.43 | 0.10510423 | 0.37407896 | 0.51276919 |
| sp\|Q9UBX5\|I | 54710210.7 | 50989932.2 | -0.1006133 | 0.37407896 | 0.51276919 |
| sp\|Q5HYI8\|R | 21283716.4 | 22575287.4 | 0.06522813 | 0.37407896 | 0.51276919 |
| sp\|O94776\|N | 579770.568 | 567675.441 | -0.2173295 | 0.37407896 | 0.51276919 |
| sp\|P01714\|L | 96380041.7 | 101632751 | 0.19503684 | 0.37407896 | 0.51276919 |
| sp\|Q9UL12\|S | 47617075.3 | 52339396.7 | 0.17830085 | 0.37407896 | 0.51276919 |
| sp\|Q9UIJ7\|K | 12607854.9 | 13205430.2 | 0.09150598 | 0.37407896 | 0.51276919 |
| sp\|P07204\|T | 798706.083 | 923977.991 | 0.11716827 | 0.37407896 | 0.51276919 |
| sp\|P08185\|C | 101142299 | 92513896.8 | -0.3087702 | 0.37407896 | 0.51276919 |
| sp\|Q5VTE0\|E | 1065763565 | 996179988 | -0.1288553 | 0.37407896 | 0.51276919 |
| sp\|Q9UEW3\| | 754407.948 | 1115333.13 | 0.25548545 | 0.37407896 | 0.51276919 |
| sp\|P84157\|M | 142760.023 | 274409.517 | 0.21318299 | 0.37407896 | 0.51276919 |
| sp\|P62310\|L | 50475921 | 58752278.4 | -0.0222776 | 0.37407896 | 0.51276919 |
| sp\|Q9HB71\|C | 119828270 | 113524783 | -0.0585738 | 0.37407896 | 0.51276919 |
| sp\|Q7L2J0\|M | 274803.678 | 318579.193 | 0.11299706 | 0.37407896 | 0.51276919 |
| sp\|Q7LBR1\|C | 2270764.3 | 2405990.64 | -0.1094517 | 0.37407896 | 0.51276919 |
| sp\|P82650\|R | 10458069.9 | 10229461.2 | -0.0563685 | 0.37407896 | 0.51276919 |
| sp\|O76080\|Z | 408632.943 | 392896.609 | 0.00632877 | 0.38639478 | 0.5251511 |
| sp\|Q5EB52\|N | 14040740.1 | 15723020.3 | 0.07831978 | 0.38639478 | 0.5251511 |
| sp\|Q15904\|V | 983375.879 | 1044809 | -0.0231363 | 0.38639478 | 0.5251511 |
| sp\|P80108\|P | 69050840.9 | 77262581.5 | 0.0617167 | 0.38639478 | 0.5251511 |
| sp\|Q9Y646\|C | 54313168.8 | 63983305.6 | 0.00418927 | 0.38639478 | 0.5251511 |
| sp\|P62256\|U | 2739137.17 | 2955203.01 | 0.07961208 | 0.38639478 | 0.5251511 |
| sp\|Q9Y5J7\|TI | 10821094.4 | 9753601.03 | -0.2313218 | 0.38639478 | 0.5251511 |
| sp\|Q9BTE3\|N | 540821.037 | 514155.722 | -0.288207 | 0.38639478 | 0.5251511 |

| | | | | | |
|---|---|---|---|---|---|
| sp\|Q9BUK0\|( | 291312.243 | 255332.37 | -0.2357556 | 0.38639478 | 0.5251511 |
| sp\|Q9BUQ8\| | 8527655.99 | 10043643.8 | 0.10356456 | 0.38639478 | 0.5251511 |
| sp\|Q93050\|\ | 88391934.1 | 93404561.9 | 0.08416756 | 0.38639478 | 0.5251511 |
| sp\|Q9H6S3\|I | 660260.044 | 1078559.58 | 0.30622553 | 0.38639478 | 0.5251511 |
| sp\|Q5VZK9\|( | 1047253.66 | 863445.202 | -0.3625312 | 0.38639478 | 0.5251511 |
| sp\|Q92626\|F | 129020099 | 149338266 | 0.16074607 | 0.38639478 | 0.5251511 |
| sp\|O75935\|D | 8592365.1 | 8958409.12 | -0.0047176 | 0.38639478 | 0.5251511 |
| sp\|Q16890\|T | 4793328.29 | 5248005.32 | 0.10391135 | 0.38639478 | 0.5251511 |
| sp\|Q9BYD2\|I | 842015.093 | 797711.443 | -0.2006356 | 0.38639478 | 0.5251511 |
| sp\|O75949\|F | 2554364.82 | 2720974.29 | 0.14601223 | 0.38639478 | 0.5251511 |
| sp\|A2RUB1\|N | 8429686.52 | 8012531.2 | -0.1200045 | 0.38639478 | 0.5251511 |
| sp\|O75915\|F | 18915507.2 | 18093499.6 | -0.1445396 | 0.38639478 | 0.5251511 |
| sp\|Q13620\|( | 794946.225 | 991336.866 | 0.25457602 | 0.38639478 | 0.5251511 |
| sp\|Q9HAT1\|L | 7588799.36 | 8164580.45 | -0.0606251 | 0.38639478 | 0.5251511 |
| sp\|O43306\|A | 287555.599 | 516456.777 | 0.463271 | 0.38639478 | 0.5251511 |
| sp\|Q14657\|L | 4912487.19 | 5847077.14 | -0.0388194 | 0.38639478 | 0.5251511 |
| sp\|Q9NS69\|T | 580435.321 | 666038.931 | 0.05176237 | 0.38639478 | 0.5251511 |
| sp\|Q9NY27\|I | 7238995.23 | 6867487.25 | -0.17377 | 0.38639478 | 0.5251511 |
| sp\|Q6QNY0\| | 1189524.21 | 1274806.84 | 0.08725418 | 0.38639478 | 0.5251511 |
| sp\|P01601\|K | 8760656.7 | 9080780.79 | 0.15405567 | 0.38639478 | 0.5251511 |
| sp\|Q03169\|T | 4839609.93 | 5576466.91 | 0.19614358 | 0.38639478 | 0.5251511 |
| sp\|Q9UNM6\| | 10997062.1 | 12197248.1 | 0.12140263 | 0.38639478 | 0.5251511 |
| sp\|Q5R3I4\|T | 53406389 | 60314518.7 | -0.0012882 | 0.39894705 | 0.53855046 |
| sp\|O95881\|T | 33596536 | 37868511.9 | 0.05041108 | 0.39894705 | 0.53855046 |
| sp\|Q9UGU5\| | 63562238 | 70129642.6 | 0.19064702 | 0.39894705 | 0.53855046 |
| sp\|P08311\|C | 127692284 | 229078285 | 0.09372941 | 0.39894705 | 0.53855046 |
| sp\|P11166\|G | 26907680.2 | 42947547.8 | 0.17798765 | 0.39894705 | 0.53855046 |
| sp\|Q15759\|N | 31943386.7 | 33599025.6 | 0.0973797 | 0.39894705 | 0.53855046 |
| sp\|Q16134\|E | 10619592 | 10082155.7 | -0.0836325 | 0.39894705 | 0.53855046 |
| sp\|Q92882\|( | 39206734.7 | 38870938.1 | -0.0416997 | 0.39894705 | 0.53855046 |
| sp\|Q9Y230\|F | 28927515.2 | 27573186.9 | -0.089045 | 0.39894705 | 0.53855046 |
| sp\|Q9Y262\|E | 4258841.47 | 4001829.56 | -0.1333152 | 0.39894705 | 0.53855046 |
| sp\|Q16831\|U | 881780.367 | 899812.073 | -0.2817404 | 0.39894705 | 0.53855046 |
| sp\|P18615\|N | 1383211.78 | 1810128.02 | 0.24361992 | 0.39894705 | 0.53855046 |
| sp\|Q9Y6B6\|S | 42473338.3 | 49600011.4 | 0.07413965 | 0.39894705 | 0.53855046 |
| sp\|Q15084\|F | 94948984.6 | 87647935.6 | -0.165297 | 0.39894705 | 0.53855046 |
| sp\|Q9H1E5\|T | 969413.772 | 1130046.74 | -0.8185852 | 0.39894705 | 0.53855046 |
| sp\|P69891\|H | 732321854 | 1221012423 | -0.0079593 | 0.39894705 | 0.53855046 |
| sp\|P41240\|C | 3932001.54 | 5847127.75 | 0.23196973 | 0.39894705 | 0.53855046 |

| | | | | | |
|---|---|---|---|---|---|
| sp\|P23434\|G | 9563385.76 | 8911756.29 | -0.0507671 | 0.39894705 | 0.53855046 |
| sp\|Q8NFU3\|T | 37555879 | 39065094.2 | 0.21381062 | 0.39894705 | 0.53855046 |
| sp\|P05026\|A | 1074225.26 | 941823.996 | -0.2649222 | 0.39894705 | 0.53855046 |
| sp\|Q96I15\|S | 17035416 | 16518244.9 | -0.0818991 | 0.39894705 | 0.53855046 |
| sp\|P01024\|C | 9812822743 | 9070912616 | -0.199541 | 0.39894705 | 0.53855046 |
| sp\|P01772\|H | 804913341 | 850342398 | 0.10921845 | 0.39894705 | 0.53855046 |
| sp\|Q9UPN6\| | 1080005.53 | 939837.715 | 0.46938936 | 0.39894705 | 0.53855046 |
| sp\|P07949\|R | 2110911.5 | 2414089.78 | 0.216357 | 0.41173374 | 0.54978025 |
| sp\|P55290\|C | 40872797.5 | 43156195.9 | 0.12074908 | 0.41173374 | 0.54978025 |
| sp\|P07148\|F | 16105272.3 | 17467162.6 | -0.1521357 | 0.41173374 | 0.54978025 |
| sp\|Q676U5\|A | 2663743.78 | 3007464.25 | 0.06093826 | 0.41173374 | 0.54978025 |
| sp\|Q99720\|S | 434151.261 | 528234.245 | 0.44234318 | 0.41173374 | 0.54978025 |
| sp\|O94880\|F | 3704981.2 | 4176050.83 | 0.02938289 | 0.41173374 | 0.54978025 |
| sp\|Q9Y266\|N | 7913944.52 | 7337100.13 | -0.135976 | 0.41173374 | 0.54978025 |
| sp\|Q92870\|A | 891808.162 | 991738.09 | 0.05328047 | 0.41173374 | 0.54978025 |
| sp\|P19320\|V | 1536781.94 | 1769986.86 | 0.09962363 | 0.41173374 | 0.54978025 |
| sp\|Q9NR46\|S | 7501433.57 | 7265053.89 | -0.1680583 | 0.41173374 | 0.54978025 |
| sp\|P13861\|K | 316318256 | 298380750 | -0.1267651 | 0.41173374 | 0.54978025 |
| sp\|Q92785\|F | 1679099.53 | 1595126.92 | -0.1187705 | 0.41173374 | 0.54978025 |
| sp\|Q9Y2A7\|N | 2010707.14 | 2735559.9 | 0.19349476 | 0.41173374 | 0.54978025 |
| sp\|Q9H1Z4\|V | 21102670.4 | 20237842.2 | -0.1512324 | 0.41173374 | 0.54978025 |
| sp\|Q8TB52\|F | 1193470.67 | 991503.748 | -0.5671708 | 0.41173374 | 0.54978025 |
| sp\|Q96LI6\|H | 21031007.8 | 14324632.1 | -0.4121433 | 0.41173374 | 0.54978025 |
| sp\|P07711\|C | 19187466.1 | 17332089.9 | -0.303369 | 0.41173374 | 0.54978025 |
| sp\|Q01484\|A | 68229039.8 | 74046084.4 | 0.07690945 | 0.41173374 | 0.54978025 |
| sp\|P01597\|K | 25399419 | 26652132.1 | 0.09275666 | 0.41173374 | 0.54978025 |
| sp\|Q01081\|U | 11185391.8 | 8552734.27 | -0.178539 | 0.41173374 | 0.54978025 |
| sp\|Q9Y5K8\|V | 331506.899 | 291142.694 | -1.4409431 | 0.41173374 | 0.54978025 |
| sp\|Q9BRA2\|T | 128872244 | 148533963 | 0.06859238 | 0.41173374 | 0.54978025 |
| sp\|P10606\|C | 62928361 | 78029588.7 | 0.25211157 | 0.41173374 | 0.54978025 |
| sp\|Q5SQN1\| | 6108827.98 | 6450295.02 | -0.4798742 | 0.41173374 | 0.54978025 |
| sp\|P52943\|C | 102225433 | 89697087.2 | -0.1783172 | 0.41173374 | 0.54978025 |
| sp\|P52565\|G | 95354105.1 | 106077256 | 0.1397851 | 0.41173374 | 0.54978025 |
| sp\|O00461\|C | 3108639 | 2895764.36 | -0.0944817 | 0.41173374 | 0.54978025 |
| sp\|P40197\|G | 2648480.31 | 3105437.6 | 0.08670366 | 0.41173374 | 0.54978025 |
| sp\|P31994\|F | 5648064.32 | 6317951.77 | 0.03480042 | 0.41173374 | 0.54978025 |
| sp\|P35580\|M | 112908664 | 106944604 | -0.0621665 | 0.41173374 | 0.54978025 |
| sp\|Q9Y3A3\|F | 4987780.98 | 4562038.03 | -0.1255005 | 0.41173374 | 0.54978025 |
| sp\|Q13643\|F | 2108996.26 | 1958616.37 | -0.1378573 | 0.41173374 | 0.54978025 |

| | | | | | |
|---|---|---|---|---|---|
| sp\|Q9NY65\|T | 201253707 | 220444501 | 0.06598527 | 0.41173374 | 0.54978025 |
| sp\|O43598\|D | 117808273 | 110968959 | -0.0925624 | 0.41173374 | 0.54978025 |
| sp\|A0A0B4J1 | 7476094.34 | 7605794.97 | 0.24786405 | 0.41173374 | 0.54978025 |
| sp\|P68366\|T | 356543301 | 387214182 | 0.04815337 | 0.41173374 | 0.54978025 |
| sp\|Q01844\|E | 34117024.3 | 37575488.4 | 0.03422282 | 0.41173374 | 0.54978025 |
| sp\|Q5SY16\|N | 2639034.26 | 3393728.86 | 0.21303517 | 0.41173374 | 0.54978025 |
| sp\|Q5SW79\| | 7532088.79 | 7940176.12 | 0.06472402 | 0.41173374 | 0.54978025 |
| sp\|Q8TBZ3\|V | 3504848.96 | 3100219.69 | -0.2117411 | 0.42475258 | 0.56340178 |
| sp\|Q9HAV0\|( | 100190068 | 95653124 | -0.0585178 | 0.42475258 | 0.56340178 |
| sp\|Q96A57\|T | 2635962.48 | 3175749.41 | 0.84658963 | 0.42475258 | 0.56340178 |
| sp\|Q9UJZ1\|S | 9772685.24 | 8681324.47 | -0.1605193 | 0.42475258 | 0.56340178 |
| sp\|Q6NUK4\| | 1470495.1 | 5938720.7 | 0.14332703 | 0.42475258 | 0.56340178 |
| sp\|P34897\|G | 70913026 | 66216868.3 | -0.1534305 | 0.42475258 | 0.56340178 |
| sp\|P36404\|A | 12765352.3 | 15634452.1 | 0.37843377 | 0.42475258 | 0.56340178 |
| sp\|Q14669\|T | 284096.947 | 350973.933 | 0.19126167 | 0.42475258 | 0.56340178 |
| sp\|O60462\|N | 20674723.6 | 22941324 | 0.12232728 | 0.42475258 | 0.56340178 |
| sp\|Q9H0Q0\| | 1799171.18 | 2038664.97 | 0.15788377 | 0.42475258 | 0.56340178 |
| sp\|O43396\|T | 85589246.4 | 78133338.2 | -0.2535592 | 0.42475258 | 0.56340178 |
| sp\|P22792\|C | 127881410 | 122562505 | -0.3094559 | 0.42475258 | 0.56340178 |
| sp\|Q9NTG7\|S | 203752.043 | 210550.276 | -0.4807629 | 0.42475258 | 0.56340178 |
| sp\|Q96N87\|S | 9248897.62 | 11362829.7 | -0.1673081 | 0.42475258 | 0.56340178 |
| sp\|Q9Y2Q3\|( | 121852637 | 156011217 | 0.24911831 | 0.42475258 | 0.56340178 |
| sp\|Q3YBR2\|T | 2153305.75 | 2247620.85 | 0.51563747 | 0.42475258 | 0.56340178 |
| sp\|P54578\|U | 150079443 | 145551393 | -0.2547693 | 0.42475258 | 0.56340178 |
| sp\|Q9BW91\| | 21046495.5 | 23571507.9 | 0.09965588 | 0.42475258 | 0.56340178 |
| sp\|O00339\|N | 15503506.9 | 14282098.2 | -0.0727081 | 0.42475258 | 0.56340178 |
| sp\|Q96IX5\|A | 2947915.4 | 3205297.61 | 0.03886983 | 0.42475258 | 0.56340178 |
| sp\|Q6DKJ4\|N | 303909.846 | 203927.882 | -0.03303 | 0.42475258 | 0.56340178 |
| sp\|L0R6Q1\|S | 1520255.31 | 1638119.15 | 0.1086802 | 0.42475258 | 0.56340178 |
| sp\|P00450\|C | 823628321 | 916213789 | 0.17804243 | 0.42475258 | 0.56340178 |
| sp\|Q8IX03\|K | 15811958.8 | 16205583 | -0.1803653 | 0.42475258 | 0.56340178 |
| sp\|P0DP04\|H | 388629929 | 408437293 | 0.12329723 | 0.43800119 | 0.57588157 |
| sp\|P61923\|C | 1394396.33 | 1609551.22 | 0.12497088 | 0.43800119 | 0.57588157 |
| sp\|P01859\|I( | 2.4531E+10 | 2.6822E+10 | 0.15913142 | 0.43800119 | 0.57588157 |
| sp\|Q7Z4G1\|( | 3669794.17 | 4528883.37 | 0.76930144 | 0.43800119 | 0.57588157 |
| sp\|O95487\|S | 5990308.84 | 5526274.96 | -0.1454572 | 0.43800119 | 0.57588157 |
| sp\|O75352\|N | 2377620.28 | 3134140.5 | 0.04622683 | 0.43800119 | 0.57588157 |
| sp\|P24386\|R | 1091178.37 | 995119.762 | -0.1921986 | 0.43800119 | 0.57588157 |
| sp\|Q15147\|P | 574305.072 | 535676.883 | -0.1788786 | 0.43800119 | 0.57588157 |

| | | | | | |
|---|---|---|---|---|---|
| sp\|P36871\|P | 1238451954 | 1337814512 | 0.05211855 | 0.43800119 | 0.57588157 |
| sp\|A0A0C4DI | 29519499.1 | 31867124 | 0.2595204 | 0.43800119 | 0.57588157 |
| sp\|Q14257\|F | 2078530.27 | 2205919.06 | 0.06620527 | 0.43800119 | 0.57588157 |
| sp\|Q96EV8\|D | 2531188.89 | 3022859.32 | 1.30213964 | 0.43800119 | 0.57588157 |
| sp\|Q96FX7\|T | 2742035.76 | 3020632.02 | 0.15365074 | 0.43800119 | 0.57588157 |
| sp\|Q9BV40\|V | 1638829.77 | 1854436.45 | 0.0742397 | 0.43800119 | 0.57588157 |
| sp\|Q9H7N4\| | 2684654.88 | 3575236.17 | 0.26740465 | 0.43800119 | 0.57588157 |
| sp\|Q5XPI4\|R | 1819019.46 | 2026282.67 | 0.00689685 | 0.43800119 | 0.57588157 |
| sp\|Q9BVA1\|T | 727914419 | 833872451 | 0.12411293 | 0.43800119 | 0.57588157 |
| sp\|Q6JBY9\|C | 15728341.6 | 17704499.8 | 0.15152206 | 0.43800119 | 0.57588157 |
| sp\|P13726\|T | 4368299.4 | 4521431.65 | -0.0290011 | 0.43800119 | 0.57588157 |
| sp\|Q5JSH3\|W | 14384245.2 | 13842087.3 | -0.0734 | 0.43800119 | 0.57588157 |
| sp\|P51398\|R | 3896484.21 | 3778000.59 | -0.1529351 | 0.43800119 | 0.57588157 |
| sp\|Q0VGL1\|L | 5057566.08 | 4794923.34 | -0.1195791 | 0.43800119 | 0.57588157 |
| sp\|O94913\|F | 1748612.24 | 1962673.59 | 0.01304322 | 0.43800119 | 0.57588157 |
| sp\|Q16787\|L | 4320281.37 | 4392529.03 | 0.07840065 | 0.43800119 | 0.57588157 |
| sp\|Q9NZJ6\|C | 1280542.9 | 1358675.47 | 0.0654567 | 0.43800119 | 0.57588157 |
| sp\|O60443\|C | 1712894.82 | 1647857.46 | -0.0243587 | 0.43800119 | 0.57588157 |
| sp\|Q13613\|M | 4192155.34 | 3595518.95 | -0.4019111 | 0.43800119 | 0.57588157 |
| sp\|O75636\|F | 38101745.9 | 35749001 | -0.2030758 | 0.43800119 | 0.57588157 |
| sp\|P19827\|IT | 721620002 | 707250728 | -0.2423634 | 0.43800119 | 0.57588157 |
| sp\|Q8N4P3\|I | 6811540.64 | 7249736.43 | 0.0956455 | 0.43800119 | 0.57588157 |
| sp\|Q96MH2\| | 113732.379 | 142770.17 | 0.12664109 | 0.43800119 | 0.57588157 |
| sp\|O95182\|N | 13114394.1 | 13261730.6 | 0.08509588 | 0.43800119 | 0.57588157 |
| sp\|A0A0C4DI | 86682211.1 | 90118934.5 | 0.09770281 | 0.45147675 | 0.58812078 |
| sp\|A0A0A0MT | 19369971.7 | 24635259.6 | 0.20222931 | 0.45147675 | 0.58812078 |
| sp\|Q8WWZ4\| | 2274304.01 | 2353536.16 | -0.0087915 | 0.45147675 | 0.58812078 |
| sp\|Q9H2P9\|I | 1009664.76 | 1702605 | 0.12461996 | 0.45147675 | 0.58812078 |
| sp\|P23381\|S | 192004259 | 176072562 | -0.1984688 | 0.45147675 | 0.58812078 |
| sp\|Q99719\|S | 10487613.6 | 9719712.66 | -0.1447481 | 0.45147675 | 0.58812078 |
| sp\|Q8IY21\|D | 4417421.21 | 3420206.54 | -0.2126063 | 0.45147675 | 0.58812078 |
| sp\|Q8N3X1\|I | 3279596.31 | 3612642.81 | 0.16890851 | 0.45147675 | 0.58812078 |
| sp\|Q4V9L6\|T | 5373272.36 | 4725932.88 | -0.2321043 | 0.45147675 | 0.58812078 |
| sp\|Q9P278\|F | 1666552.69 | 1648128.59 | -0.0490126 | 0.45147675 | 0.58812078 |
| sp\|Q96NC0\| | 360872.852 | 475720.171 | 0.20804317 | 0.45147675 | 0.58812078 |
| sp\|Q12972\|F | 212524682 | 194359405 | -0.1738217 | 0.45147675 | 0.58812078 |
| sp\|Q96M27\|I | 16973531 | 17800911.5 | 0.03444874 | 0.45147675 | 0.58812078 |
| sp\|A0A075B6 | 68569126.3 | 74918917.1 | 0.17448162 | 0.45147675 | 0.58812078 |
| sp\|A0A0J9YX | 107012671 | 113359949 | 0.22537711 | 0.45147675 | 0.58812078 |

| | | | | | |
|---|---|---|---|---|---|
| sp\|Q14767\|L | 25385989.4 | 24128803.1 | -0.1287863 | 0.45147675 | 0.58812078 |
| sp\|Q9Y303\|N | 46058177.8 | 53548843.1 | 0.09088723 | 0.45147675 | 0.58812078 |
| sp\|O60711\|L | 438645.156 | 574568.77 | 0.29301596 | 0.45147675 | 0.58812078 |
| sp\|O95394\|A | 83712019.7 | 89078570.5 | -0.0348233 | 0.45147675 | 0.58812078 |
| sp\|Q5T7N3\|K | 38768693.7 | 35033665.1 | -0.1439288 | 0.45147675 | 0.58812078 |
| sp\|O00442\|F | 65165099.6 | 74600816.8 | 0.08255637 | 0.45147675 | 0.58812078 |
| sp\|Q86UX7\|U | 13721273.3 | 15253039.9 | -0.0749793 | 0.45147675 | 0.58812078 |
| sp\|P01019\|A | 225402481 | 210206820 | -0.1946324 | 0.45147675 | 0.58812078 |
| sp\|Q9Y4E8\|U | 68679963.9 | 78346446.9 | 0.08997068 | 0.45147675 | 0.58812078 |
| sp\|P01700\|L | 84240243 | 87474552.4 | 0.08709141 | 0.45147675 | 0.58812078 |
| sp\|Q02952\|A | 1100364873 | 1143016354 | -0.0190801 | 0.45147675 | 0.58812078 |
| sp\|Q93091\|F | 427883.104 | 369867.497 | -0.2993967 | 0.45147675 | 0.58812078 |
| sp\|Q04941\|F | 6237714.26 | 6051932.95 | -0.1033287 | 0.45147675 | 0.58812078 |
| sp\|Q9UIC8\|L | 46944627.3 | 50563637.7 | 0.10561866 | 0.45147675 | 0.58812078 |
| sp\|Q9BTT0\|A | 6963395.09 | 6660186.65 | -0.2978338 | 0.45147675 | 0.58812078 |
| sp\|P01764\|H | 293518539 | 304969321 | 0.12394349 | 0.45147675 | 0.58812078 |
| sp\|P01782\|H | 387751661 | 410882550 | 0.12514729 | 0.45147675 | 0.58812078 |
| sp\|P01780\|H | 980531498 | 1037497778 | 0.14387101 | 0.45147675 | 0.58812078 |
| sp\|P02794\|F | 33679365.4 | 19428998.9 | -0.4833432 | 0.45147675 | 0.58812078 |
| sp\|P51608\|M | 19704584.1 | 16691577.6 | -0.1871167 | 0.46517621 | 0.60010232 |
| sp\|P01717\|L | 91400962.5 | 93885793.9 | 0.18441022 | 0.46517621 | 0.60010232 |
| sp\|P01009\|A | 1.044E+10 | 1.1408E+10 | 0.10297555 | 0.46517621 | 0.60010232 |
| sp\|P16333\|N | 29516176.1 | 31336190.9 | 0.08119499 | 0.46517621 | 0.60010232 |
| sp\|Q9Y6Y8\|S | 6291654.48 | 6892230.04 | 0.13387699 | 0.46517621 | 0.60010232 |
| sp\|P53621\|C | 8812415.99 | 8448597.12 | -0.1083681 | 0.46517621 | 0.60010232 |
| sp\|Q53TN4\|C | 26430707.3 | 24648933.6 | -0.0962963 | 0.46517621 | 0.60010232 |
| sp\|P01857\|IC | 3.6322E+10 | 3.8878E+10 | 0.11557495 | 0.46517621 | 0.60010232 |
| sp\|Q3MHD2\| | 8518236.21 | 9369980.85 | 0.05407045 | 0.46517621 | 0.60010232 |
| sp\|Q5HYK3\|C | 113555.99 | 151256.559 | 0.51346829 | 0.46517621 | 0.60010232 |
| sp\|P55145\|M | 3037268.65 | 3428836.34 | 0.1792845 | 0.46517621 | 0.60010232 |
| sp\|O95573\|A | 17812884.5 | 16874111.4 | -0.0836702 | 0.46517621 | 0.60010232 |
| sp\|Q10589\|E | 23123490.5 | 23209996.7 | 0.06209349 | 0.46517621 | 0.60010232 |
| sp\|P21810\|P | 71208151.3 | 104222520 | 0.10789532 | 0.46517621 | 0.60010232 |
| sp\|P21397\|A | 172323995 | 183269112 | 0.06781088 | 0.46517621 | 0.60010232 |
| sp\|Q9UK99\|F | 576843.585 | 1449110.31 | -0.4413785 | 0.46517621 | 0.60010232 |
| sp\|Q86X76\|N | 144176338 | 151781051 | -0.0221068 | 0.46517621 | 0.60010232 |
| sp\|Q9UI08\|E | 5611772.83 | 5855122.21 | -0.0909012 | 0.46517621 | 0.60010232 |
| sp\|Q9BPX5\|A | 6672118.47 | 6249359.77 | -0.104398 | 0.46517621 | 0.60010232 |
| sp\|HBB_HUM | 4.9486E+10 | 7.527E+10 | -0.0594716 | 0.46517621 | 0.60010232 |

| | | | | | |
|---|---|---|---|---|---|
| sp\|Q5VW32\| | 33185214.6 | 32427616.6 | -0.0577249 | 0.46517621 | 0.60010232 |
| sp\|P50747\|B | 1095059.74 | 1224541.91 | 0.15600338 | 0.46517621 | 0.60010232 |
| sp\|P51397\|D | 5734687 | 7768586.44 | 0.22026826 | 0.46517621 | 0.60010232 |
| sp\|P98175\|R | 5245126.93 | 4822283.42 | -0.1333717 | 0.46517621 | 0.60010232 |
| sp\|Q02218\|( | 453254352 | 475171480 | -0.0590843 | 0.46517621 | 0.60010232 |
| sp\|Q01415\|( | 1056256.4 | 1027098.91 | -0.4715746 | 0.46517621 | 0.60010232 |
| sp\|Q71U36\|T | 485132159 | 519781901 | 0.03190358 | 0.46517621 | 0.60010232 |
| sp\|P18887\|X | 357816.034 | 340927.382 | -0.3794393 | 0.46517621 | 0.60010232 |
| sp\|Q7L5N1\|( | 4158458.19 | 4101491.48 | -0.0193188 | 0.46517621 | 0.60010232 |
| sp\|Q7L2H7\|E | 1155753.38 | 1083752.95 | -0.4068909 | 0.46517621 | 0.60010232 |
| sp\|Q7Z3T8\|Z | 4415647.48 | 4719473 | 0.10410962 | 0.46517621 | 0.60010232 |
| sp\|Q7RTV0\|F | 22721138.3 | 21945845.5 | -0.1384749 | 0.46517621 | 0.60010232 |
| sp\|O14520\|A | 998961.761 | 1135517.05 | -0.0050812 | 0.46517621 | 0.60010232 |
| sp\|Q9H2C2\|. | 12093670.6 | 16512103.8 | 0.44607504 | 0.46517621 | 0.60010232 |
| sp\|Q96IY4\|C | 64010697.3 | 71256529.8 | 0.09209466 | 0.46517621 | 0.60010232 |
| sp\|Q96GX9\|I | 39058055 | 40179263.8 | -0.0223877 | 0.46517621 | 0.60010232 |
| sp\|Q9BZF1\|( | 949970.847 | 982654.75 | 0.05851416 | 0.47909638 | 0.61148498 |
| sp\|Q5T5U3\|F | 6640848.34 | 6921951.05 | 0.04232625 | 0.47909638 | 0.61148498 |
| sp\|Q9P1F3\|A | 10062846.1 | 10867512.5 | 0.06655106 | 0.47909638 | 0.61148498 |
| sp\|Q9Y5K5\|U | 187688.158 | 177061.648 | -0.6839036 | 0.47909638 | 0.61148498 |
| sp\|Q9ULA0\|D | 277212894 | 257316016 | -0.2298248 | 0.47909638 | 0.61148498 |
| sp\|Q9Y6W5\|' | 22623034.4 | 21575029.6 | -0.0509372 | 0.47909638 | 0.61148498 |
| sp\|Q7Z6Z7\|H | 6704212.94 | 6423865.73 | -0.1258198 | 0.47909638 | 0.61148498 |
| sp\|A8MT70\|Z | 31853039 | 28469610.8 | -0.5068876 | 0.47909638 | 0.61148498 |
| sp\|P63261\|A | 1.7193E+10 | 1.9959E+10 | 0.13301716 | 0.47909638 | 0.61148498 |
| sp\|Q86XP1\|D | 2498647.94 | 2667229.25 | -0.0330759 | 0.47909638 | 0.61148498 |
| sp\|Q9NW15\| | 392634.204 | 438653.812 | -1.5027003 | 0.47909638 | 0.61148498 |
| sp\|Q9P225\|D | 5126130.7 | 4774892.31 | -0.1906684 | 0.47909638 | 0.61148498 |
| sp\|Q8N201\|I | 6060848.19 | 6654652.1 | 0.12804489 | 0.47909638 | 0.61148498 |
| sp\|B9A064\|I( | 7154118910 | 7585792409 | 0.10380105 | 0.47909638 | 0.61148498 |
| sp\|P48506\|G | 205913161 | 188644635 | -0.2925739 | 0.47909638 | 0.61148498 |
| sp\|O00479\|H | 11821753.5 | 12189839.1 | -0.18086 | 0.47909638 | 0.61148498 |
| sp\|Q8TCA0\|L | 5253986.02 | 5124328.74 | -0.1407594 | 0.47909638 | 0.61148498 |
| sp\|P13671\|C | 235929200 | 224457214 | -0.4062774 | 0.47909638 | 0.61148498 |
| sp\|Q9Y315\|D | 31933592.3 | 30067527.8 | -0.0832053 | 0.47909638 | 0.61148498 |
| sp\|Q8NDX5\| | 402267.09 | 360170.245 | -0.8189934 | 0.47909638 | 0.61148498 |
| sp\|O60229\|K | 9805659.08 | 11348494 | -0.0339856 | 0.47909638 | 0.61148498 |
| sp\|P49459\|U | 571123.266 | 534808.579 | -0.1570365 | 0.47909638 | 0.61148498 |
| sp\|Q10713\|M | 38289477 | 34450970.4 | -0.196031 | 0.47909638 | 0.61148498 |

| | | | | | |
|---|---|---|---|---|---|
| sp\|Q12913\|F | 1050294.08 | 1049680.94 | 0.0800992 | 0.47909638 | 0.61148498 |
| sp\|Q12965\|N | 4564193.39 | 4081698.68 | 0.26828209 | 0.47909638 | 0.61148498 |
| sp\|Q9H3N8\| | 260352.162 | 173480.917 | -0.5772412 | 0.47909638 | 0.61148498 |
| sp\|P51665\|P | 21028794 | 22164911.2 | 0.08672692 | 0.47909638 | 0.61148498 |
| sp\|O43708\|N | 18635538.3 | 20037963.3 | -0.005852 | 0.47909638 | 0.61148498 |
| sp\|Q12800\|T | 7046514.16 | 6344855.45 | -0.0901745 | 0.47909638 | 0.61148498 |
| sp\|Q687X5\|S | 5771881.38 | 4950826.47 | -0.1724369 | 0.47909638 | 0.61148498 |
| sp\|Q02127\|F | 505304.67 | 552360.742 | 0.03430963 | 0.47909638 | 0.61148498 |
| sp\|O15344\|T | 1263411.57 | 1397119.53 | 0.07752866 | 0.47909638 | 0.61148498 |
| sp\|Q9Y4K3\|T | 1012438.85 | 1257934.23 | 0.19473212 | 0.47909638 | 0.61148498 |
| sp\|Q13610\|F | 9027628.24 | 9917039.17 | 0.09675885 | 0.47909638 | 0.61148498 |
| sp\|O14562\|U | 4266464.4 | 4442514.27 | 0.05075346 | 0.47909638 | 0.61148498 |
| sp\|Q96BY7\|A | 3117690.01 | 2808763.25 | -0.2860835 | 0.47909638 | 0.61148498 |
| sp\|Q63HN8\| | 335195.412 | 2577457.94 | 0.38040125 | 0.47909638 | 0.61148498 |
| sp\|Q9UDR5\|. | 6261568.9 | 5941215.26 | -0.1008246 | 0.47909638 | 0.61148498 |
| sp\|P10916\|M | 1163177.25 | 2926610.91 | 0.31758013 | 0.47909638 | 0.61148498 |
| sp\|Q15121\|F | 105504425 | 113143798 | 0.01331411 | 0.47909638 | 0.61148498 |
| sp\|Q9BZV1\|U | 7106783.23 | 7842246.05 | 0.13101813 | 0.49323367 | 0.62339435 |
| sp\|P21926\|C | 28706617.9 | 31983768.7 | 0.0983408 | 0.49323367 | 0.62339435 |
| sp\|Q8IZ21\|PI | 735305.117 | 988124.241 | -0.3476354 | 0.49323367 | 0.62339435 |
| sp\|Q8IYB3\|S | 4011469.32 | 8320132.27 | 0.03046819 | 0.49323367 | 0.62339435 |
| sp\|Q86TX2\|A | 679325413 | 724143468 | -0.0076912 | 0.49323367 | 0.62339435 |
| sp\|Q9NVA2\|S | 235792324 | 215655748 | -0.1904585 | 0.49323367 | 0.62339435 |
| sp\|Q562R1\|A | 6065053259 | 7367538740 | 0.18635414 | 0.49323367 | 0.62339435 |
| sp\|Q9UHD1\| | 2152014.76 | 2143176.71 | -0.1940025 | 0.49323367 | 0.62339435 |
| sp\|P18754\|R | 1884976.15 | 1954767.63 | 0.07432205 | 0.49323367 | 0.62339435 |
| sp\|Q9P013\|C | 16983717.4 | 15815964.9 | -0.1360678 | 0.49323367 | 0.62339435 |
| sp\|Q9BV20\|N | 759695.429 | 827068.961 | -0.0323438 | 0.49323367 | 0.62339435 |
| sp\|A0A0C4DI | 56262653.2 | 60048030 | 0.10062389 | 0.49323367 | 0.62339435 |
| sp\|O14531\|D | 82719450.1 | 80186062.9 | -0.0845145 | 0.49323367 | 0.62339435 |
| sp\|O00401\|V | 5282972.99 | 5404754.25 | 0.0180821 | 0.49323367 | 0.62339435 |
| sp\|Q6ZNX1\|S | 7465131.1 | 7211782.25 | -0.081291 | 0.49323367 | 0.62339435 |
| sp\|Q9BZZ2\|S | 12123926.3 | 12491477 | 0.02847695 | 0.49323367 | 0.62339435 |
| sp\|O14867\|E | 3635972.74 | 3123813.55 | -0.1005055 | 0.49323367 | 0.62339435 |
| sp\|Q9UBX3\|I | 3692795.13 | 3425832.41 | -0.2257681 | 0.49323367 | 0.62339435 |
| sp\|P22830\|H | 30126811.3 | 31472362.7 | 0.05230482 | 0.49323367 | 0.62339435 |
| sp\|Q9GZM8\| | 396996.296 | 430504.903 | 0.0564318 | 0.49323367 | 0.62339435 |
| sp\|O00232\|F | 9509313.58 | 9824769.36 | -0.1781595 | 0.49323367 | 0.62339435 |
| sp\|Q9UBQ7\| | 253358511 | 269478554 | 0.06129481 | 0.49323367 | 0.62339435 |

| | | | | | |
|---|---|---|---|---|---|
| sp\|O76054\|S | 1860707.89 | 2345932.03 | -0.1302172 | 0.49323367 | 0.62339435 |
| sp\|P17936\|IE | 18333621.9 | 20606343.5 | 0.08547214 | 0.49323367 | 0.62339435 |
| sp\|Q8IV36\|H | 1470677.52 | 1312812.78 | -0.503665 | 0.49323367 | 0.62339435 |
| sp\|Q9BVG4\|I | 7905998.94 | 8229094.91 | 0.0599891 | 0.49323367 | 0.62339435 |
| sp\|P15170\|E | 35321798.8 | 32790414.2 | -0.1441428 | 0.49323367 | 0.62339435 |
| sp\|Q14558\|K | 40041786.3 | 38282018.6 | -0.1286658 | 0.49323367 | 0.62339435 |
| sp\|P63241\|IF | 191896257 | 201961516 | -0.003395 | 0.49323367 | 0.62339435 |
| sp\|Q8IUR5\|T | 100367417 | 122664027 | 0.42894092 | 0.49323367 | 0.62339435 |
| sp\|P55212\|C | 10136596 | 10850661.5 | 0.04122631 | 0.49323367 | 0.62339435 |
| sp\|Q8TCG1\|( | 7465131.1 | 7211782.25 | -0.081291 | 0.49323367 | 0.62339435 |
| sp\|O43896\|K | 5693947.56 | 5273338.59 | -0.0549387 | 0.49323367 | 0.62339435 |
| sp\|P16220\|C | 3527877.8 | 3741053.28 | -0.0751663 | 0.49323367 | 0.62339435 |
| sp\|Q9Y5P4\|C | 500874.212 | 593000.972 | 0.07918889 | 0.49323367 | 0.62339435 |
| sp\|Q9UNP9\| | 11734114.4 | 13124036.2 | 0.08096059 | 0.49323367 | 0.62339435 |
| sp\|P49915\|G | 8553817.96 | 9248744.11 | 0.04996296 | 0.49323367 | 0.62339435 |
| sp\|P02008\|H | 6321269.66 | 5496458.16 | 0.25413931 | 0.50758421 | 0.63550656 |
| sp\|P60983\|G | 40799124.7 | 39474332.7 | -0.0759528 | 0.50758421 | 0.63550656 |
| sp\|Q8TD06\|A | 2641137.49 | 1937837.16 | -1.6120918 | 0.50758421 | 0.63550656 |
| sp\|Q9NYW2\| | 11590098.9 | 9123284.08 | -0.0742588 | 0.50758421 | 0.63550656 |
| sp\|P62879\|G | 150261019 | 143940250 | -0.0688915 | 0.50758421 | 0.63550656 |
| sp\|P07355\|A | 845154612 | 787996452 | -0.1206684 | 0.50758421 | 0.63550656 |
| sp\|P82912\|R | 1342476.19 | 1605407.58 | -0.1391191 | 0.50758421 | 0.63550656 |
| sp\|A2RUC4\|T | 5019439.6 | 17202974.3 | 0.4843471 | 0.50758421 | 0.63550656 |
| sp\|Q8IZL2\|M | 875966.348 | 1126227.55 | 0.1393903 | 0.50758421 | 0.63550656 |
| sp\|Q16698\|D | 149396879 | 159568955 | 0.09341243 | 0.50758421 | 0.63550656 |
| sp\|Q8N5L8\|F | 606832.108 | 569792.956 | -0.2368388 | 0.50758421 | 0.63550656 |
| sp\|P05543\|T | 69888587.3 | 65986478 | -0.1137036 | 0.50758421 | 0.63550656 |
| sp\|O75746\|C | 915656.579 | 852081.632 | -0.2145594 | 0.50758421 | 0.63550656 |
| sp\|P30153\|2 | 58692123.2 | 69093879.3 | 0.16535194 | 0.50758421 | 0.63550656 |
| sp\|Q9Y4A5\|T | 8992403.09 | 8582338.08 | -0.1368285 | 0.50758421 | 0.63550656 |
| sp\|O95139\|N | 10824471.4 | 13635766.2 | 0.20116252 | 0.50758421 | 0.63550656 |
| sp\|P18583\|S | 4670036.61 | 4109919.58 | -0.1715731 | 0.50758421 | 0.63550656 |
| sp\|Q7L5Y1\|E | 28508313.6 | 35015567.5 | 0.16038378 | 0.50758421 | 0.63550656 |
| sp\|P08575\|P | 47562447.4 | 50108831 | 0.05685809 | 0.50758421 | 0.63550656 |
| sp\|Q96HJ9\|F | 1813366.98 | 1994360.77 | 0.12101761 | 0.50758421 | 0.63550656 |
| sp\|Q86W50\| | 30501635.8 | 32841216.5 | 0.06495902 | 0.50758421 | 0.63550656 |
| sp\|Q9H008\|L | 88544447.7 | 88820455.8 | -0.0816779 | 0.50758421 | 0.63550656 |
| sp\|Q9BXW7\| | 3291190.62 | 3386816.78 | -0.0344569 | 0.50758421 | 0.63550656 |
| sp\|Q12955\|A | 11210771.6 | 18483375.2 | 0.20426972 | 0.50758421 | 0.63550656 |

| | | | | | |
|---|---|---|---|---|---|
| sp\|P02689\|M | 436662936 | 477382371 | 0.07547374 | 0.50758421 | 0.63550656 |
| sp\|P60174\|T | 5423295288 | 6610663237 | 0.11353597 | 0.50758421 | 0.63550656 |
| sp\|Q96B54\|Z | 5174933.17 | 4552101.66 | -0.3413117 | 0.50758421 | 0.63550656 |
| sp\|O00534\|V | 55207295 | 51298326.2 | -0.1607277 | 0.50758421 | 0.63550656 |
| sp\|Q15819\|U | 80303428.3 | 77117431 | -0.1549612 | 0.50758421 | 0.63550656 |
| sp\|Q9BTE1\|D | 4202844.38 | 3920512.98 | -0.1439911 | 0.50758421 | 0.63550656 |
| sp\|P0DP03\|H | 323999228 | 333026584 | 0.09403531 | 0.50758421 | 0.63550656 |
| sp\|Q9UFE4\|C | 3015966.24 | 3042896.61 | -0.1349237 | 0.50758421 | 0.63550656 |
| sp\|Q6IA69\|N | 29845448.1 | 28369129.6 | -0.0873817 | 0.50758421 | 0.63550656 |
| sp\|Q969K4\|A | 2558245.99 | 2119708.51 | -0.1932711 | 0.50758421 | 0.63550656 |
| sp\|Q6UX71\|P | 27620495.6 | 25743357.7 | -0.1314202 | 0.50758421 | 0.63550656 |
| sp\|A0A0B4J1 | 23937593 | 23242351.6 | 0.0906774 | 0.50758421 | 0.63550656 |
| sp\|Q92896\|G | 1233605.91 | 1143027.28 | 0.19571555 | 0.52214407 | 0.64882688 |
| sp\|Q6YHK3\|C | 36170737.5 | 39627459.6 | 0.04852202 | 0.52214407 | 0.64882688 |
| sp\|O75144\|I | 1232177.41 | 1096196.65 | 1.26047526 | 0.52214407 | 0.64882688 |
| sp\|P0DJD7\|P | 3366412.67 | 1564712.96 | 0.2324583 | 0.52214407 | 0.64882688 |
| sp\|Q9BZL4\|P | 18601624.9 | 17848881.1 | -0.0868 | 0.52214407 | 0.64882688 |
| sp\|Q2TAA2\|I | 92167492.8 | 89591275 | -0.0429989 | 0.52214407 | 0.64882688 |
| sp\|Q8IXI2\|MI | 1265117.7 | 1437590.63 | 0.13343681 | 0.52214407 | 0.64882688 |
| sp\|Q9UPQ0\| | 36525003 | 35218908 | -0.0460075 | 0.52214407 | 0.64882688 |
| sp\|Q9Y3M2\|C | 555895.978 | 492314.065 | -0.2067982 | 0.52214407 | 0.64882688 |
| sp\|Q86VQ1\|C | 4191962.09 | 4919778.32 | 0.1226991 | 0.52214407 | 0.64882688 |
| sp\|P68871\|H | 4.9589E+10 | 7.4933E+10 | -0.0646359 | 0.52214407 | 0.64882688 |
| sp\|P04070\|P | 6337537.94 | 6737201.02 | -0.1284628 | 0.52214407 | 0.64882688 |
| sp\|Q8WZ82\| | 5158473.65 | 8110341.34 | 0.18883306 | 0.52214407 | 0.64882688 |
| sp\|P29692\|E | 322249597 | 330099043 | 0.02014173 | 0.52214407 | 0.64882688 |
| sp\|P02042\|H | 4.2178E+10 | 6.3842E+10 | -0.0639255 | 0.52214407 | 0.64882688 |
| sp\|P01614\|K | 384960260 | 425408801 | 0.11879082 | 0.52214407 | 0.64882688 |
| sp\|P31948\|S | 242993221 | 260923733 | 0.08822914 | 0.52214407 | 0.64882688 |
| sp\|P31939\|P | 758742517 | 800241441 | 0.11365803 | 0.52214407 | 0.64882688 |
| sp\|Q9UQ80\| | 86786935.3 | 85785062.1 | -0.0236549 | 0.52214407 | 0.64882688 |
| sp\|P30043\|B | 2417222604 | 2951245844 | 0.10117603 | 0.52214407 | 0.64882688 |
| sp\|Q99611\|S | 825168.419 | 810023.175 | -0.2082599 | 0.52214407 | 0.64882688 |
| sp\|Q9BT09\|C | 20205955.9 | 19548858.5 | -0.1170617 | 0.52214407 | 0.64882688 |
| sp\|Q9P0P8\|M | 389931.675 | 374863.594 | -0.4071012 | 0.52214407 | 0.64882688 |
| sp\|O75995\|S | 3232499.29 | 3689729.03 | 0.07104901 | 0.52214407 | 0.64882688 |
| sp\|O94763\|R | 1092122.87 | 1298009.95 | 0.43875369 | 0.52214407 | 0.64882688 |
| sp\|Q9Y2L1\|R | 357267.277 | 387575.426 | 0.12646612 | 0.52214407 | 0.64882688 |
| sp\|P06312\|K | 446738301 | 473352934 | 0.20049318 | 0.52214407 | 0.64882688 |

| | | | | | |
|---|---|---|---|---|---|
| sp\|Q14624\|I | 974724471 | 944463075 | -0.2865563 | 0.52214407 | 0.64882688 |
| sp\|Q9UKY7\|( | 1883235.47 | 2193699.89 | 0.17280047 | 0.52214407 | 0.64882688 |
| sp\|P01599\|K | 30785843.7 | 32877779.6 | 0.1183942 | 0.53690887 | 0.6622014 |
| sp\|P01709\|L | 94696444.9 | 99914735.6 | 0.14068913 | 0.53690887 | 0.6622014 |
| sp\|Q9Y5X1\|S | 2632204.18 | 2629891.2 | -0.1845167 | 0.53690887 | 0.6622014 |
| sp\|O75674\|T | 3866358.38 | 4657671.78 | 0.10386105 | 0.53690887 | 0.6622014 |
| sp\|Q9NY15\|S | 44893971.9 | 44712092.6 | -0.0488081 | 0.53690887 | 0.6622014 |
| sp\|Q96PD5\|I | 118072885 | 115586351 | -0.5238783 | 0.53690887 | 0.6622014 |
| sp\|Q96A00\|F | 26621947.9 | 29364700.6 | 0.09362127 | 0.53690887 | 0.6622014 |
| sp\|P02730\|B | 198175180 | 390562335 | 0.22435431 | 0.53690887 | 0.6622014 |
| sp\|Q92561\|F | 6385461.18 | 6701863.17 | 0.05589449 | 0.53690887 | 0.6622014 |
| sp\|O43924\|F | 10516212.3 | 10227636.4 | -0.0610445 | 0.53690887 | 0.6622014 |
| sp\|P04181\|C | 56973218.9 | 56821301.1 | -0.1211642 | 0.53690887 | 0.6622014 |
| sp\|P49902\|5 | 11469265.1 | 11794166.6 | 0.03102356 | 0.53690887 | 0.6622014 |
| sp\|Q8TDS4\|I | 1498906.05 | 1594292.47 | 0.21482945 | 0.53690887 | 0.6622014 |
| sp\|P16452\|E | 37716939 | 67934654.2 | 0.20819284 | 0.53690887 | 0.6622014 |
| sp\|Q86U28\|I | 29795013.5 | 31166346.1 | 0.0337723 | 0.53690887 | 0.6622014 |
| sp\|Q15019\|S | 321388912 | 286888035 | -0.1970579 | 0.53690887 | 0.6622014 |
| sp\|Q99933\|E | 3308280.9 | 3613040.99 | 0.11442728 | 0.53690887 | 0.6622014 |
| sp\|Q9BYK8\|I | 5946170.94 | 5666782.81 | -0.6702073 | 0.53690887 | 0.6622014 |
| sp\|P23786\|C | 12960831.7 | 13807001.6 | 0.02097009 | 0.53690887 | 0.6622014 |
| sp\|P07311\|A | 20852062.9 | 20174719.9 | -0.0752047 | 0.53690887 | 0.6622014 |
| sp\|Q03154\|A | 295863603 | 264182473 | -0.1318109 | 0.53690887 | 0.6622014 |
| sp\|P98170\|X | 3143068.61 | 2678489.09 | -0.4312397 | 0.53690887 | 0.6622014 |
| sp\|Q9UNF0\|I | 3485424 | 5729499.87 | 0.28015806 | 0.53690887 | 0.6622014 |
| sp\|P51153\|R | 56972839.4 | 57450179.6 | -0.0203125 | 0.53690887 | 0.6622014 |
| sp\|Q9BTV5\|F | 7033158.97 | 6848973.18 | -0.0110074 | 0.53690887 | 0.6622014 |
| sp\|Q9BSJ8\|E | 50879641.6 | 48959962.4 | -0.1583488 | 0.53690887 | 0.6622014 |
| sp\|Q9UDY2\|Z | 51575855.1 | 49245534.2 | -0.1234689 | 0.53690887 | 0.6622014 |
| sp\|Q14126\|D | 55475319.2 | 58651559.3 | 0.05119467 | 0.53690887 | 0.6622014 |
| sp\|Q96IZ0\|PA | 11269193.4 | 10039862.2 | -0.2036665 | 0.53690887 | 0.6622014 |
| sp\|A0A075B6 | 162384114 | 173524181 | 0.15082488 | 0.55187397 | 0.67441894 |
| sp\|Q92917\|( | 2576663.35 | 2371415.61 | -0.1907539 | 0.55187397 | 0.67441894 |
| sp\|Q96EY5\|N | 1784234.62 | 1886086.69 | 0.04141711 | 0.55187397 | 0.67441894 |
| sp\|P10301\|R | 155321511 | 159384217 | 0.00348299 | 0.55187397 | 0.67441894 |
| sp\|P11021\|B | 2008702521 | 1950006332 | -0.0504181 | 0.55187397 | 0.67441894 |
| sp\|Q9NPA0\|I | 1392492.57 | 1492027.66 | -0.0527677 | 0.55187397 | 0.67441894 |
| sp\|P50238\|C | 482824529 | 478484358 | -0.3385049 | 0.55187397 | 0.67441894 |
| sp\|Q9Y6I3\|EI | 7512452.74 | 8212101.56 | 0.10871758 | 0.55187397 | 0.67441894 |

| | | | | | |
|---|---|---|---|---|---|
| sp\|P0DP02\|H | 817170004 | 836361892 | 0.08222843 | 0.55187397 | 0.67441894 |
| sp\|P07327\|A | 8266122207 | 8540406242 | -0.2342663 | 0.55187397 | 0.67441894 |
| sp\|P30613\|K | 297394820 | 425543180 | 0.2411511 | 0.55187397 | 0.67441894 |
| sp\|Q9UHW5\| | 1830108.49 | 1883372.72 | 0.01924427 | 0.55187397 | 0.67441894 |
| sp\|Q7Z2W9\| | 4807315.96 | 4557634.65 | -0.2622293 | 0.55187397 | 0.67441894 |
| sp\|O43286\|E | 304872.857 | 374725.808 | -0.1768561 | 0.55187397 | 0.67441894 |
| sp\|Q02161\|F | 751118.006 | 932195.261 | 0.02668035 | 0.55187397 | 0.67441894 |
| sp\|Q15431\|S | 4153524.09 | 4122514.42 | -0.1677402 | 0.55187397 | 0.67441894 |
| sp\|P16402\|H | 301868663 | 291397202 | -0.0893207 | 0.55187397 | 0.67441894 |
| sp\|Q04917\|1 | 1314545130 | 1269517296 | -0.0578311 | 0.55187397 | 0.67441894 |
| sp\|P35658\|N | 7932293.26 | 8105871.41 | 0.02057021 | 0.55187397 | 0.67441894 |
| sp\|Q06033\|I | 58567160.1 | 68093465.1 | 0.073076 | 0.55187397 | 0.67441894 |
| sp\|O60832\|D | 11701144.9 | 13328917.3 | 0.14625355 | 0.55187397 | 0.67441894 |
| sp\|Q8NBQ5\| | 94533725.6 | 101638269 | 0.06075918 | 0.55187397 | 0.67441894 |
| sp\|P53675\|C | 20729884.8 | 20880923.9 | -0.0638724 | 0.55187397 | 0.67441894 |
| sp\|P52803\|E | 2174485.97 | 1864632.44 | -0.3626746 | 0.55187397 | 0.67441894 |
| sp\|P21281\|V | 74987380.2 | 74382333.3 | -0.1041974 | 0.55187397 | 0.67441894 |
| sp\|P61964\|W | 2749372.18 | 2913069.45 | 0.03336477 | 0.55187397 | 0.67441894 |
| sp\|O95168\|N | 14234941.5 | 13257566.4 | -0.1768151 | 0.55187397 | 0.67441894 |
| sp\|Q8IXM2\|E | 4919195.77 | 4875668.25 | -0.0265716 | 0.55187397 | 0.67441894 |
| sp\|Q5VTE6\|A | 3462238.88 | 3753628.95 | -0.0021698 | 0.55187397 | 0.67441894 |
| sp\|Q8IW45\|N | 70194707 | 61866861.2 | -0.1288046 | 0.55187397 | 0.67441894 |
| sp\|Q13404\|U | 84508674.7 | 92978263.5 | -0.0340697 | 0.55187397 | 0.67441894 |
| sp\|P30837\|A | 953125792 | 910103338 | -0.0656005 | 0.55187397 | 0.67441894 |
| sp\|P49448\|D | 421681183 | 422936515 | -0.0248226 | 0.55187397 | 0.67441894 |
| sp\|Q8WUP2\| | 20169177.7 | 18499777.5 | -0.1025213 | 0.55187397 | 0.67441894 |
| sp\|Q12931\|T | 41921082.9 | 39654934.8 | -0.1132336 | 0.55187397 | 0.67441894 |
| sp\|P15291\|B | 304872.857 | 374725.808 | -0.1768561 | 0.55187397 | 0.67441894 |
| sp\|Q9H7Z7\|F | 3632257.26 | 4027990.93 | 0.11746767 | 0.56703466 | 0.68665136 |
| sp\|Q9UHY1\| | 1511280.75 | 1458571.44 | -0.1878971 | 0.56703466 | 0.68665136 |
| sp\|Q9Y2S7\|F | 9387087.23 | 8392979.05 | -0.1129946 | 0.56703466 | 0.68665136 |
| sp\|Q7Z3B1\|N | 7248622.99 | 7092344.48 | -0.1294348 | 0.56703466 | 0.68665136 |
| sp\|O00273\|D | 21622434 | 21140397.1 | -0.0699487 | 0.56703466 | 0.68665136 |
| sp\|P01721\|L | 584607.587 | 562639.393 | -0.0630154 | 0.56703466 | 0.68665136 |
| sp\|Q96EC8\|Y | 2741015.41 | 3198724.6 | 0.25415115 | 0.56703466 | 0.68665136 |
| sp\|Q96FN4\|C | 3480097.88 | 3288533.85 | -0.1658218 | 0.56703466 | 0.68665136 |
| sp\|A0A0B4J1 | 101865506 | 101851148 | 0.08889631 | 0.56703466 | 0.68665136 |
| sp\|Q9BV86\|N | 2306066.18 | 2384556.78 | -0.009442 | 0.56703466 | 0.68665136 |
| sp\|Q9H853\|T | 47188312.7 | 55281540.2 | 0.08995338 | 0.56703466 | 0.68665136 |

| | | | | | |
|---|---|---|---|---|---|
| sp\|P20742\|P | 2186519476 | 2076924731 | -0.0984254 | 0.56703466 | 0.68665136 |
| sp\|P30038\|A | 650415537 | 585834607 | -0.2432169 | 0.56703466 | 0.68665136 |
| sp\|P61764\|S | 27440341.7 | 28278022.9 | -0.0027481 | 0.56703466 | 0.68665136 |
| sp\|P46013\|K | 38298987.6 | 42360604.4 | -0.006551 | 0.56703466 | 0.68665136 |
| sp\|P46108\|C | 110864493 | 122285721 | 0.0733817 | 0.56703466 | 0.68665136 |
| sp\|Q8IZF2\|A | 826032.03 | 942872.032 | 0.10656924 | 0.56703466 | 0.68665136 |
| sp\|Q9BYZ2\|L | 19330530.7 | 15397122.5 | -0.3082524 | 0.56703466 | 0.68665136 |
| sp\|Q9BVS4\|F | 1521213.26 | 1445184.94 | -0.0856811 | 0.56703466 | 0.68665136 |
| sp\|P23508\|C | 482359.857 | 533266.181 | 0.10458488 | 0.56703466 | 0.68665136 |
| sp\|Q8N3D4\| | 7884538 | 8165207.17 | 0.03069002 | 0.56703466 | 0.68665136 |
| sp\|Q99460\|F | 197410764 | 203551653 | -0.0104775 | 0.56703466 | 0.68665136 |
| sp\|Q07352\|T | 696149.884 | 849446.161 | 0.24351846 | 0.56703466 | 0.68665136 |
| sp\|P53602\|M | 26775451.8 | 25708830.4 | -0.1050582 | 0.56703466 | 0.68665136 |
| sp\|P56545\|C | 9178453.16 | 8872226.01 | -0.0770775 | 0.56703466 | 0.68665136 |
| sp\|Q9Y605\|N | 1429900.33 | 1410499.47 | -0.0313602 | 0.56703466 | 0.68665136 |
| sp\|Q6S8J3\|P | 8800033402 | 9995286014 | 0.12142236 | 0.56703466 | 0.68665136 |
| sp\|P0CG29\|C | 12907014.1 | 11788844.5 | -0.0758607 | 0.56703466 | 0.68665136 |
| sp\|P04196\|H | 618741086 | 647358817 | -0.0630559 | 0.56703466 | 0.68665136 |
| sp\|Q9UL26\|F | 3333812.94 | 3316980.12 | -0.0072986 | 0.56703466 | 0.68665136 |
| sp\|P04003\|C | 406833103 | 446036732 | -0.5182095 | 0.56703466 | 0.68665136 |
| sp\|Q9Y6Q2\|S | 959211.335 | 969993.338 | -0.0378143 | 0.56703466 | 0.68665136 |
| sp\|P49841\|G | 535299.253 | 499934.582 | -0.1278699 | 0.56703466 | 0.68665136 |
| sp\|P00747\|P | 540394129 | 561325448 | -0.227933 | 0.56703466 | 0.68665136 |
| sp\|Q9Y5M8\|S | 3726229.3 | 3618402.41 | -0.0731137 | 0.56703466 | 0.68665136 |
| sp\|Q9Y5V0\|Z | 20304825.8 | 21791062.1 | -0.1413244 | 0.56703466 | 0.68665136 |
| sp\|Q96EI5\|T | 5408047.37 | 5253122.21 | -0.0926008 | 0.58238583 | 0.69767089 |
| sp\|Q96JB5\|C | 2219388.75 | 1723512.02 | -0.4216788 | 0.58238583 | 0.69767089 |
| sp\|Q92696\|F | 1719493874 | 1821301032 | 0.03118235 | 0.58238583 | 0.69767089 |
| sp\|Q96GG9\| | 9667088.46 | 11760843.5 | 0.19598517 | 0.58238583 | 0.69767089 |
| sp\|Q969M7\|U | 4244368.79 | 4252344.62 | -0.0424938 | 0.58238583 | 0.69767089 |
| sp\|Q9Y5Y7\|L | 65975243 | 77044952.7 | -0.1991834 | 0.58238583 | 0.69767089 |
| sp\|P13284\|G | 5045085.54 | 6618554.2 | 0.14674862 | 0.58238583 | 0.69767089 |
| sp\|P12270\|T | 106321341 | 102959163 | -0.0444602 | 0.58238583 | 0.69767089 |
| sp\|Q6DN14\| | 215463.503 | 198670.279 | -0.1593178 | 0.58238583 | 0.69767089 |
| sp\|O94826\|T | 7047018.58 | 7758044.83 | 0.06983729 | 0.58238583 | 0.69767089 |
| sp\|Q9UQL6\|I | 275882.341 | 284450.888 | 0.34498309 | 0.58238583 | 0.69767089 |
| sp\|Q00610\|C | 87146603.8 | 95458224.7 | 0.10976494 | 0.58238583 | 0.69767089 |
| sp\|P15090\|F | 1.0895E+10 | 1.1464E+10 | -0.0035638 | 0.58238583 | 0.69767089 |
| sp\|Q9H4B7\|T | 301930737 | 328140308 | 0.08411949 | 0.58238583 | 0.69767089 |

| | | | | | |
|---|---|---|---|---|---|
| sp\|Q9BW30\| | 210992427 | 215852259 | -0.175316 | 0.58238583 | 0.69767089 |
| sp\|Q2TAY7\|S | 58565564.5 | 61253929.5 | -0.0155441 | 0.58238583 | 0.69767089 |
| sp\|Q9UQR1\| | 20525680.2 | 18348482.8 | -0.6236619 | 0.58238583 | 0.69767089 |
| sp\|P40306\|P | 29071322.1 | 27819724.1 | -0.2067763 | 0.58238583 | 0.69767089 |
| sp\|Q86UU1\|I | 101061868 | 104755508 | 0.04474188 | 0.58238583 | 0.69767089 |
| sp\|P07225\|P | 75711473.7 | 90742836.9 | 0.1033594 | 0.58238583 | 0.69767089 |
| sp\|Q9H9J2\|F | 23255361.3 | 25687412.7 | 0.06672263 | 0.58238583 | 0.69767089 |
| sp\|P30519\|H | 1268744.38 | 1217900.32 | -0.1735739 | 0.58238583 | 0.69767089 |
| sp\|P25189\|M | 527309.91 | 1403239.88 | 0.1609801 | 0.58238583 | 0.69767089 |
| sp\|P24821\|T | 2250743.9 | 1762685.1 | -0.2759941 | 0.58238583 | 0.69767089 |
| sp\|Q6YN16\|I | 96682763.7 | 102545430 | 0.0168627 | 0.58238583 | 0.69767089 |
| sp\|P09488\|G | 255955131 | 320027305 | -0.1249067 | 0.58238583 | 0.69767089 |
| sp\|O00116\|A | 525480.887 | 585958.154 | 0.16252356 | 0.58238583 | 0.69767089 |
| sp\|P12724\|E | 26012322 | 44530723.6 | -0.2590626 | 0.58238583 | 0.69767089 |
| sp\|Q6UUV7\| | 3288982.52 | 4780841.24 | 0.82673519 | 0.58238583 | 0.69767089 |
| sp\|P21291\|C | 499979726 | 477593621 | -0.0930844 | 0.58238583 | 0.69767089 |
| sp\|Q14651\|F | 26593518.4 | 28105987.4 | -0.2605146 | 0.58238583 | 0.69767089 |
| sp\|O15116\|L | 6253635.92 | 5606912.81 | -0.0553979 | 0.58238583 | 0.69767089 |
| sp\|P04439\|H | 79233996 | 81827308.4 | 0.07073285 | 0.58238583 | 0.69767089 |
| sp\|P48444\|C | 83112680 | 80090774.8 | -0.0750126 | 0.58238583 | 0.69767089 |
| sp\|Q9NRF8\|I | 36076278.1 | 34124335.9 | -0.0802075 | 0.58238583 | 0.69767089 |
| sp\|P23368\|M | 93064022.3 | 89058534.8 | -0.1314385 | 0.58238583 | 0.69767089 |
| sp\|Q8WXF1\| | 27767220.9 | 27855123.1 | -0.0070861 | 0.58238583 | 0.69767089 |
| sp\|P49356\|F | 22178036.5 | 23684906.4 | 0.02503501 | 0.58238583 | 0.69767089 |
| sp\|O95477\|A | 4979207.55 | 5234909.36 | 0.14453289 | 0.58238583 | 0.69767089 |
| sp\|Q8NI60\|C | 1001561.86 | 930888.852 | -0.1456238 | 0.58238583 | 0.69767089 |
| sp\|A1L0T0\|H | 37646672.6 | 36750275.1 | -0.1020445 | 0.58238583 | 0.69767089 |
| sp\|P46531\|N | 1222937.1 | 1269474.67 | 0.02069335 | 0.58238583 | 0.69767089 |
| sp\|Q9NS37\|Z | 13203513.8 | 12635469 | -0.0704942 | 0.58238583 | 0.69767089 |
| sp\|P05106\|IT | 9803713.6 | 16890576.3 | 0.31810807 | 0.5979221 | 0.71254735 |
| sp\|O95749\|G | 10192782.4 | 12939559.2 | 0.15875034 | 0.5979221 | 0.71254735 |
| sp\|Q8N4Q0\| | 9038837.62 | 8509692.63 | -0.0441693 | 0.5979221 | 0.71254735 |
| sp\|P01042\|K | 1442867152 | 1560264700 | -0.0239439 | 0.5979221 | 0.71254735 |
| sp\|Q9H270\|V | 2211285.08 | 2247620.85 | -0.2211699 | 0.5979221 | 0.71254735 |
| sp\|O15151\|M | 12548108.3 | 11813778 | -0.1087207 | 0.5979221 | 0.71254735 |
| sp\|P60842\|IF | 27121906.4 | 26049423.9 | -0.0875635 | 0.5979221 | 0.71254735 |
| sp\|Q9C0E8\|L | 5212284.16 | 5106330.84 | -0.07348 | 0.5979221 | 0.71254735 |
| sp\|Q96AC1\|F | 138929338 | 130576414 | -0.0536406 | 0.5979221 | 0.71254735 |
| sp\|O75112\|L | 12067407.1 | 10924646.4 | -0.1864843 | 0.5979221 | 0.71254735 |

| | | | | | |
|---|---|---|---|---|---|
| sp\|O15488\|( | 12289573 | 12127431.5 | 0.00715945 | 0.5979221 | 0.71254735 |
| sp\|P14555\|P | 168542.588 | 137709.013 | -0.767164 | 0.5979221 | 0.71254735 |
| sp\|Q5VYS8\|T | 54969893.7 | 59579431.5 | -0.1462093 | 0.5979221 | 0.71254735 |
| sp\|Q4LDE5\|S | 5179449.23 | 5166069.11 | -0.0483228 | 0.5979221 | 0.71254735 |
| sp\|Q96GQ5\| | 1171374.32 | 1056415.54 | -0.3412197 | 0.5979221 | 0.71254735 |
| sp\|Q12889\|( | 1068147.15 | 652545.587 | 0.03969013 | 0.5979221 | 0.71254735 |
| sp\|P11047\|L | 942938923 | 905562740 | -0.0933163 | 0.5979221 | 0.71254735 |
| sp\|P68371\|T | 898421672 | 974398804 | 0.05412207 | 0.5979221 | 0.71254735 |
| sp\|Q96JJ7\|TN | 680807.666 | 705708.081 | 0.00402714 | 0.5979221 | 0.71254735 |
| sp\|Q5VVQ6\|( | 20300913.6 | 28526096.8 | 0.05477771 | 0.5979221 | 0.71254735 |
| sp\|P16066\|A | 1730370.01 | 1716473.73 | -0.0577057 | 0.5979221 | 0.71254735 |
| sp\|P24666\|P | 70138676.1 | 76959799 | -0.3353941 | 0.61363805 | 0.72676432 |
| sp\|Q86YJ6\|TI | 94437.9926 | 120749.445 | 0.26771305 | 0.61363805 | 0.72676432 |
| sp\|Q9UKG1\| | 5628944.9 | 6064118.62 | 0.10246868 | 0.61363805 | 0.72676432 |
| sp\|P05091\|A | 5460903111 | 5906924615 | 0.0358127 | 0.61363805 | 0.72676432 |
| sp\|O95825\|( | 25103953.8 | 25607368 | -0.0143139 | 0.61363805 | 0.72676432 |
| sp\|Q9H8W4\| | 1721867.08 | 1744255.81 | -0.0344772 | 0.61363805 | 0.72676432 |
| sp\|P69892\|H | 683797184 | 1160698514 | 0.0809188 | 0.61363805 | 0.72676432 |
| sp\|Q15369\|E | 28963645.2 | 30248574.6 | 0.02633245 | 0.61363805 | 0.72676432 |
| sp\|P82979\|S | 31736142.4 | 31138949.4 | -0.106685 | 0.61363805 | 0.72676432 |
| sp\|Q6Y7W6\| | 992829.431 | 1029535.75 | 0.02884367 | 0.61363805 | 0.72676432 |
| sp\|Q9Y508\|F | 14237743.6 | 14074416.3 | -0.0641737 | 0.61363805 | 0.72676432 |
| sp\|P09105\|H | 9402073.91 | 17082236.2 | 0.30313721 | 0.61363805 | 0.72676432 |
| sp\|Q16270\|I | 28588007.6 | 28352676.6 | -0.0584945 | 0.61363805 | 0.72676432 |
| sp\|Q9BY89\|K | 3682893.14 | 3562407.65 | -0.0544948 | 0.61363805 | 0.72676432 |
| sp\|P13647\|K | 1429999673 | 976226784 | -0.3263947 | 0.61363805 | 0.72676432 |
| sp\|O43402\|E | 34033055.6 | 33168537.6 | -0.8970986 | 0.61363805 | 0.72676432 |
| sp\|Q5EBM0\|( | 14028703.9 | 13968526 | -0.2163681 | 0.61363805 | 0.72676432 |
| sp\|Q96KP4\|( | 679140768 | 664535074 | -0.1320324 | 0.61363805 | 0.72676432 |
| sp\|A0A0A0MS | 4167747.29 | 3901698.6 | -0.1114945 | 0.61363805 | 0.72676432 |
| sp\|A5A3E0\|P | 8293664751 | 9462647983 | 0.11120271 | 0.61363805 | 0.72676432 |
| sp\|Q99442\|S | 392719.13 | 358169.321 | -0.3427227 | 0.61363805 | 0.72676432 |
| sp\|Q8TF40\|F | 1012334.74 | 18502740.3 | 0.64956903 | 0.61363805 | 0.72676432 |
| sp\|O60313\|( | 579939.214 | 561687.798 | -0.2503285 | 0.61363805 | 0.72676432 |
| sp\|Q9ULD0\|( | 109827418 | 114193788 | -0.0861786 | 0.61363805 | 0.72676432 |
| sp\|P00367\|D | 610884617 | 613033274 | -0.0119172 | 0.61363805 | 0.72676432 |
| sp\|A0A087W | 9924157.26 | 10942860 | 0.17025673 | 0.62952783 | 0.74010388 |
| sp\|Q9NYU1\| | 7950530.59 | 7798476.28 | -0.0605073 | 0.62952783 | 0.74010388 |
| sp\|Q4G0N4\| | 66198912.9 | 67682066 | 0.04448507 | 0.62952783 | 0.74010388 |

| | | | | | |
|---|---|---|---|---|---|
| sp\|Q0JRZ9\|F | 42584788.7 | 40734020 | -0.1228958 | 0.62952783 | 0.74010388 |
| sp\|O00712\|N | 1956071.21 | 2151774.6 | 0.16157882 | 0.62952783 | 0.74010388 |
| sp\|Q16881\|T | 167761977 | 166182218 | -0.0677494 | 0.62952783 | 0.74010388 |
| sp\|Q9C005\|[ | 4341277.51 | 3919846.76 | -0.1178447 | 0.62952783 | 0.74010388 |
| sp\|Q13356\|F | 5246953.6 | 5494164.36 | 0.14192398 | 0.62952783 | 0.74010388 |
| sp\|P15121\|A | 483587015 | 477842186 | -0.1292845 | 0.62952783 | 0.74010388 |
| sp\|Q9UHN6\| | 4262055.66 | 7541969.49 | 0.34584627 | 0.62952783 | 0.74010388 |
| sp\|Q9BR61\|A | 93275.4574 | 117037.727 | 0.3432588 | 0.62952783 | 0.74010388 |
| sp\|Q9Y4Y9\|L | 6169111.81 | 6435116.2 | 0.03722257 | 0.62952783 | 0.74010388 |
| sp\|Q9UBC2\| | 61559375.9 | 61824564.3 | -0.0020518 | 0.62952783 | 0.74010388 |
| sp\|Q6IAN0\|D | 2487756.3 | 2574607.55 | -0.0198864 | 0.62952783 | 0.74010388 |
| sp\|Q9P2A4\|A | 12043745.5 | 12897843.6 | 0.98538191 | 0.62952783 | 0.74010388 |
| sp\|Q5QJ74\|T | 2031367.59 | 2837441.05 | 0.20055329 | 0.62952783 | 0.74010388 |
| sp\|O75390\|C | 255889518 | 278648053 | 0.09659415 | 0.62952783 | 0.74010388 |
| sp\|P46060\|R | 4054190.19 | 3850118.65 | -0.1043447 | 0.62952783 | 0.74010388 |
| sp\|P80404\|G | 20795453.2 | 19642710.5 | -0.0304358 | 0.62952783 | 0.74010388 |
| sp\|Q86W25\| | 2221989.89 | 2605000.97 | -0.0956036 | 0.62952783 | 0.74010388 |
| sp\|A6NMZ7\|C | 2393412.76 | 2497362.55 | 0.01300552 | 0.62952783 | 0.74010388 |
| sp\|Q99584\|S | 22474461.3 | 24852554 | 0.06330021 | 0.62952783 | 0.74010388 |
| sp\|Q96PE1\|A | 681575.375 | 754267.22 | 0.24233843 | 0.62952783 | 0.74010388 |
| sp\|Q8WZ64\|A | 380661.523 | 403632.06 | 0.18908403 | 0.62952783 | 0.74010388 |
| sp\|Q99590\|S | 177326.881 | 176297.968 | -0.3178621 | 0.62952783 | 0.74010388 |
| sp\|P04062\|G | 11904115.8 | 11744523.3 | -0.0995762 | 0.62952783 | 0.74010388 |
| sp\|Q9NQ88\| | 5121781.41 | 5524888.52 | -0.0023331 | 0.62952783 | 0.74010388 |
| sp\|Q8NHV1\| | 1049340.91 | 1520891.23 | 0.65872095 | 0.62952783 | 0.74010388 |
| sp\|P04278\|S | 45336731.8 | 42245522.5 | -0.2078906 | 0.62952783 | 0.74010388 |
| sp\|Q9NQX3\| | 25127186.8 | 24158900.4 | -0.0536416 | 0.62952783 | 0.74010388 |
| sp\|P11413\|G | 416314586 | 414162632 | 0.00115815 | 0.64558538 | 0.75509731 |
| sp\|Q5T653\|F | 5499848.97 | 5525368.57 | -0.0631086 | 0.64558538 | 0.75509731 |
| sp\|Q6NUM9\| | 45270062.6 | 46379212.5 | -0.0547744 | 0.64558538 | 0.75509731 |
| sp\|Q03591\|F | 167810138 | 167110659 | -0.1203323 | 0.64558538 | 0.75509731 |
| sp\|Q8WVJ2\|N | 25000049.5 | 25280083.7 | -0.0338346 | 0.64558538 | 0.75509731 |
| sp\|O60496\|D | 199923.024 | 196659.995 | -0.5982842 | 0.64558538 | 0.75509731 |
| sp\|P04211\|L | 122301289 | 136794953 | 0.09990336 | 0.64558538 | 0.75509731 |
| sp\|O95340\|F | 527921.762 | 483689.263 | -0.363266 | 0.64558538 | 0.75509731 |
| sp\|O43567\|F | 3653445.48 | 3570674.11 | -0.0653477 | 0.64558538 | 0.75509731 |
| sp\|O94919\|E | 2338959.68 | 2645533.73 | 0.04490229 | 0.64558538 | 0.75509731 |
| sp\|Q5KU26\|C | 1354413.84 | 1355619.14 | -0.1362702 | 0.64558538 | 0.75509731 |
| sp\|Q9BZE1\|F | 14923383.6 | 15311626.9 | -0.0148965 | 0.64558538 | 0.75509731 |

| | | | | | |
|---|---|---|---|---|---|
| sp\|O15078\|( | 1570193.96 | 1762782.02 | 0.0580678 | 0.64558538 | 0.75509731 |
| sp\|Q96C03\|I | 8804273.91 | 9428184.78 | -0.0521709 | 0.64558538 | 0.75509731 |
| sp\|P30084\|E | 431390502 | 399794502 | -0.1242717 | 0.64558538 | 0.75509731 |
| sp\|Q14966\|Z | 311202.739 | 316202.406 | -0.1149052 | 0.64558538 | 0.75509731 |
| sp\|A1X283\|S | 2024213.24 | 2018930.79 | -0.0454281 | 0.64558538 | 0.75509731 |
| sp\|P01893\|H | 45704519.7 | 46073047.4 | 0.07791546 | 0.64558538 | 0.75509731 |
| sp\|P01008\|A | 1466880603 | 1424264594 | -0.1268387 | 0.64558538 | 0.75509731 |
| sp\|P49756\|R | 5590780.12 | 5478233.95 | -0.0219759 | 0.64558538 | 0.75509731 |
| sp\|P38646\|G | 914665029 | 901272276 | -0.0250994 | 0.64558538 | 0.75509731 |
| sp\|P17096\|H | 5936046.98 | 6301076.74 | 0.04969187 | 0.66180459 | 0.76789174 |
| sp\|O75506\|H | 1728221.41 | 2100927.93 | 0.06242903 | 0.66180459 | 0.76789174 |
| sp\|P25685\|D | 15545636.8 | 15153385.7 | -0.0984165 | 0.66180459 | 0.76789174 |
| sp\|P14406\|C | 8415492.63 | 8592978.8 | -0.2010759 | 0.66180459 | 0.76789174 |
| sp\|Q15417\|( | 86148140.5 | 80705907.8 | -0.0856092 | 0.66180459 | 0.76789174 |
| sp\|P36543\|V | 14022083.4 | 13005748.8 | -0.1940995 | 0.66180459 | 0.76789174 |
| sp\|P63302\|S | 91132.8632 | 126034.29 | 0.00485927 | 0.66180459 | 0.76789174 |
| sp\|Q8IU80\|T | 2430988.6 | 2400284.26 | -0.1710162 | 0.66180459 | 0.76789174 |
| sp\|Q9HB07\|I | 31324583.1 | 32136303.7 | -0.1315186 | 0.66180459 | 0.76789174 |
| sp\|P32519\|E | 2524920.81 | 3059762.15 | 0.11936876 | 0.66180459 | 0.76789174 |
| sp\|Q7Z2K6\|E | 2615560.06 | 1689900.03 | 0.01634558 | 0.66180459 | 0.76789174 |
| sp\|P00325\|A | 1.9864E+10 | 2.0621E+10 | -0.2386768 | 0.66180459 | 0.76789174 |
| sp\|P36269\|G | 21031207.7 | 19949394.3 | -0.138173 | 0.66180459 | 0.76789174 |
| sp\|P22612\|K | 15929203.3 | 18446779 | 0.05553216 | 0.66180459 | 0.76789174 |
| sp\|O14880\|M | 17927835.8 | 16758569.5 | -0.0912502 | 0.66180459 | 0.76789174 |
| sp\|Q14155\|A | 145459.78 | 120912.296 | -0.7088457 | 0.66180459 | 0.76789174 |
| sp\|Q14353\|( | 16859049 | 18705537.9 | -0.0519488 | 0.66180459 | 0.76789174 |
| sp\|O75695\|X | 857915.068 | 1089132 | 0.19915818 | 0.66180459 | 0.76789174 |
| sp\|P61956\|S | 174711853 | 183355469 | 0.0053976 | 0.66180459 | 0.76789174 |
| sp\|P61626\|L | 11920582.8 | 41546766.2 | 0.12992552 | 0.66180459 | 0.76789174 |
| sp\|Q8WWQ0 | 1300269.59 | 1210290.5 | -0.2074052 | 0.66180459 | 0.76789174 |
| sp\|P01817\|H | 9586297.7 | 10171228.6 | 0.07271144 | 0.66180459 | 0.76789174 |
| sp\|Q92539\|L | 516846.845 | 525179.921 | -0.1696128 | 0.66180459 | 0.76789174 |
| sp\|A6ZKI3\|RT | 2956403.09 | 2873515.26 | -0.1231271 | 0.66180459 | 0.76789174 |
| sp\|P52799\|E | 785469.811 | 845094.087 | -0.1277423 | 0.66180459 | 0.76789174 |
| sp\|P54826\|G | 276132.339 | 284650.19 | -0.2392108 | 0.66180459 | 0.76789174 |
| sp\|P53618\|C | 3190989.16 | 3074303.12 | -0.1084095 | 0.66180459 | 0.76789174 |
| sp\|Q96QG7\| | 1353341.36 | 1305503.27 | -0.202987 | 0.66180459 | 0.76789174 |
| sp\|Q9H1C4\| | 95549.214 | 86040.8302 | -0.1783751 | 0.66180459 | 0.76789174 |
| sp\|Q14520\|H | 37386524.6 | 40281711.3 | -0.1556106 | 0.66180459 | 0.76789174 |

| | | | | | |
|---|---|---|---|---|---|
| sp\|P32929\|C | 18699161.1 | 15531205 | -0.1518943 | 0.66180459 | 0.76789174 |
| sp\|Q13303\|K | 6429032.84 | 6269139.89 | -0.0736575 | 0.66180459 | 0.76789174 |
| sp\|Q9C0B0\|U | 3593105.29 | 3577713.28 | -0.0285942 | 0.66180459 | 0.76789174 |
| sp\|P08648\|IT | 30390028.1 | 30465841.6 | -0.0097424 | 0.67817897 | 0.78010088 |
| sp\|Q9H993\|A | 26119819.6 | 25747694.5 | -0.0365485 | 0.67817897 | 0.78010088 |
| sp\|Q9UH99\|S | 442659.31 | 523535.167 | 0.12487809 | 0.67817897 | 0.78010088 |
| sp\|P43652\|A | 481113344 | 471173888 | -0.1181738 | 0.67817897 | 0.78010088 |
| sp\|P46926\|G | 199679004 | 201696834 | -0.1124177 | 0.67817897 | 0.78010088 |
| sp\|Q13177\|P | 16208680.4 | 17275978.8 | 0.08196008 | 0.67817897 | 0.78010088 |
| sp\|Q9ULN7\|I | 311018.382 | 319833.981 | -0.2241218 | 0.67817897 | 0.78010088 |
| sp\|Q93009\|U | 21391708 | 20634117.9 | -0.1503892 | 0.67817897 | 0.78010088 |
| sp\|Q8WXH0\| | 7980169.63 | 7787821.24 | 0.19004401 | 0.67817897 | 0.78010088 |
| sp\|Q92781\|P | 13289151.1 | 14267394.5 | 0.0528011 | 0.67817897 | 0.78010088 |
| sp\|Q9Y6C9\|M | 12847019 | 13349252 | 0.06658463 | 0.67817897 | 0.78010088 |
| sp\|Q96AB6\|N | 24849870 | 27545075.3 | 0.00048173 | 0.67817897 | 0.78010088 |
| sp\|Q8N126\|C | 3703023.66 | 4242842.93 | -0.0378832 | 0.67817897 | 0.78010088 |
| sp\|P49662\|C | 25834732.2 | 24903941.4 | -0.0894873 | 0.67817897 | 0.78010088 |
| sp\|Q9ULC4\|I | 25659011 | 24795377.5 | -0.0321369 | 0.67817897 | 0.78010088 |
| sp\|Q8IZQ5\|S | 13726127.4 | 6783300.88 | -0.1300619 | 0.67817897 | 0.78010088 |
| sp\|Q8TAQ9\|S | 56867041.4 | 56517230.7 | -0.1479989 | 0.67817897 | 0.78010088 |
| sp\|P62195\|P | 18898462.6 | 20607048.8 | -0.0051053 | 0.67817897 | 0.78010088 |
| sp\|Q13724\|M | 2330125.75 | 2353426.7 | -0.1300729 | 0.67817897 | 0.78010088 |
| sp\|Q9Y3D3\|R | 1815616 | 1667087.2 | -0.3897073 | 0.67817897 | 0.78010088 |
| sp\|Q9BYN0\|S | 78586231.1 | 84448684.5 | 0.10661134 | 0.67817897 | 0.78010088 |
| sp\|P00505\|A | 277647573 | 305822515 | -0.0562595 | 0.67817897 | 0.78010088 |
| sp\|Q96CX2\|K | 167682701 | 159295114 | -0.0594022 | 0.67817897 | 0.78010088 |
| sp\|Q9BTW9\|T | 516877.059 | 529092.253 | -0.235675 | 0.67817897 | 0.78010088 |
| sp\|Q6Y1H2\|H | 410416.202 | 391125.513 | -0.2226285 | 0.67817897 | 0.78010088 |
| sp\|O00154\|B | 4222906.03 | 4208488.21 | -0.1124253 | 0.67817897 | 0.78010088 |
| sp\|Q68D91\|M | 431531.883 | 424514.669 | 0.09046313 | 0.67817897 | 0.78010088 |
| sp\|P13645\|K | 309984993 | 246493092 | -0.2104751 | 0.67817897 | 0.78010088 |
| sp\|O15126\|S | 129630.77 | 138230.234 | -0.9746027 | 0.67817897 | 0.78010088 |
| sp\|Q14112\|N | 110444107 | 103779679 | -0.0628469 | 0.67817897 | 0.78010088 |
| sp\|O00567\|N | 5567480.44 | 5686984.38 | -0.0101915 | 0.67817897 | 0.78010088 |
| sp\|P29972\|A | 1762137.85 | 1648487.48 | -0.4048486 | 0.67817897 | 0.78010088 |
| sp\|P26885\|F | 94730062.3 | 92747085.5 | -0.0423295 | 0.67817897 | 0.78010088 |
| sp\|P35908\|K | 285359068 | 230318240 | -0.1817922 | 0.67817897 | 0.78010088 |
| sp\|A0A0B4J1 | 143424864 | 149061005 | 0.12489537 | 0.67817897 | 0.78010088 |
| sp\|Q9Y6V7\|D | 1110532.22 | 1258283.49 | 0.07279617 | 0.67817897 | 0.78010088 |

| | | | | | |
|---|---|---|---|---|---|
| sp\|Q13123\|F | 57030492.1 | 40575605.6 | 0.06667348 | 0.69470181 | 0.79227044 |
| sp\|P16403\|H | 295063755 | 317355421 | 0.05267247 | 0.69470181 | 0.79227044 |
| sp\|Q9NZ43\|U | 759825.322 | 826110.857 | 0.09032881 | 0.69470181 | 0.79227044 |
| sp\|P00374\|D | 4006283.35 | 5080656.21 | 0.16890402 | 0.69470181 | 0.79227044 |
| sp\|A0A075B6 | 85227619.4 | 98389628.9 | 0.03204552 | 0.69470181 | 0.79227044 |
| sp\|O14657\|T | 825543.543 | 16247179.1 | -0.2256038 | 0.69470181 | 0.79227044 |
| sp\|P78563\|R | 21258332.8 | 20459153.8 | 0.01810913 | 0.69470181 | 0.79227044 |
| sp\|P46976\|G | 64168084.9 | 71123705.6 | 0.11496039 | 0.69470181 | 0.79227044 |
| sp\|Q53SF7\|C | 29968348.8 | 30533870.5 | -0.0545104 | 0.69470181 | 0.79227044 |
| sp\|Q04756\|H | 24302796 | 23763364.5 | -0.119066 | 0.69470181 | 0.79227044 |
| sp\|Q05655\|K | 27078765.3 | 50777164.3 | 0.11171572 | 0.69470181 | 0.79227044 |
| sp\|Q15018\|A | 12933648.7 | 13841587.2 | 0.0417309 | 0.69470181 | 0.79227044 |
| sp\|Q13825\|A | 11875339.8 | 11770661.2 | -0.0862896 | 0.69470181 | 0.79227044 |
| sp\|Q9NX63\|N | 8033776.67 | 9152807.04 | 0.15320766 | 0.69470181 | 0.79227044 |
| sp\|Q9Y2W7\| | 3476009.84 | 5608817.38 | 0.09787048 | 0.69470181 | 0.79227044 |
| sp\|Q969G3\|S | 2001241.72 | 2258849.54 | -0.0916977 | 0.69470181 | 0.79227044 |
| sp\|Q9UJW0\|I | 37017640.2 | 37434999.8 | -0.0135242 | 0.69470181 | 0.79227044 |
| sp\|A1Z1Q3\|N | 4216483.75 | 4321630.08 | -0.0292278 | 0.69470181 | 0.79227044 |
| sp\|P48556\|P | 3514083.58 | 3983307.07 | 0.09115215 | 0.69470181 | 0.79227044 |
| sp\|A0A0B4J1 | 8150562.73 | 7184453.05 | -0.1110312 | 0.69470181 | 0.79227044 |
| sp\|P61604\|C | 1100647265 | 1067915925 | -0.0027448 | 0.69470181 | 0.79227044 |
| sp\|O95456\|F | 1653768.91 | 1418518.07 | -0.8046509 | 0.69470181 | 0.79227044 |
| sp\|Q8N1F7\|N | 318913.837 | 324547.234 | -0.2818006 | 0.69470181 | 0.79227044 |
| sp\|P04150\|G | 7800418.97 | 7606716.86 | -0.0212373 | 0.69470181 | 0.79227044 |
| sp\|Q9Y6A4\|C | 367469.944 | 369731.845 | -0.0604917 | 0.69470181 | 0.79227044 |
| sp\|P51452\|D | 61313903.5 | 56598399.3 | -0.1117119 | 0.69470181 | 0.79227044 |
| sp\|Q6NUN0\| | 15845598.4 | 15915040.5 | -0.0100488 | 0.69470181 | 0.79227044 |
| sp\|Q9Y2I8\|W | 12867570.4 | 22428668.6 | 0.241326 | 0.69470181 | 0.79227044 |
| sp\|Q08431\|N | 1962772.98 | 2195600.53 | 0.03880069 | 0.69470181 | 0.79227044 |
| sp\|Q07021\|C | 5258051.55 | 5302016.66 | -0.2436755 | 0.69470181 | 0.79227044 |
| sp\|Q01658\|N | 1373275.37 | 1399330.23 | -0.2054979 | 0.69470181 | 0.79227044 |
| sp\|O75116\|R | 1367472.17 | 1355413.07 | -0.075331 | 0.69470181 | 0.79227044 |
| sp\|Q9Y2L6\|F | 7622296.63 | 7700259.02 | 0.0226877 | 0.69470181 | 0.79227044 |
| sp\|Q6UW68\| | 2565716.27 | 2444605.41 | -0.1693398 | 0.69470181 | 0.79227044 |
| sp\|P38606\|V | 138393263 | 138861733 | -0.1032819 | 0.69470181 | 0.79227044 |
| sp\|A0M8Q6\|I | 7099944494 | 7123036071 | 0.0231505 | 0.69470181 | 0.79227044 |
| sp\|Q9Y484\|W | 8596196.24 | 10264546.8 | -0.32124 | 0.7113664 | 0.80553265 |
| sp\|Q16570\|A | 1282792.61 | 1487072.83 | -0.0362877 | 0.7113664 | 0.80553265 |
| sp\|Q13976\|K | 400159.95 | 375166.154 | -0.5944836 | 0.7113664 | 0.80553265 |

| | | | | | |
|---|---|---|---|---|---|
| sp\|Q15223\|N | 1653480.46 | 1583529.26 | -0.0698659 | 0.7113664 | 0.80553265 |
| sp\|P45973\|C | 22852305.2 | 22706634.7 | -0.0849515 | 0.7113664 | 0.80553265 |
| sp\|Q99729\|F | 81567008 | 82660934 | -0.0483442 | 0.7113664 | 0.80553265 |
| sp\|Q99983\|( | 3352629.13 | 2935542.75 | -0.0056311 | 0.7113664 | 0.80553265 |
| sp\|O95777\|L | 13130831.2 | 14638709.8 | 0.09263282 | 0.7113664 | 0.80553265 |
| sp\|P08603\|C | 1859180792 | 1858101152 | -0.1635381 | 0.7113664 | 0.80553265 |
| sp\|Q709C8\|V | 30423425.2 | 30509562.3 | -0.0901768 | 0.7113664 | 0.80553265 |
| sp\|Q7L2E3\|D | 1345704.23 | 1440054.42 | -0.1275243 | 0.7113664 | 0.80553265 |
| sp\|Q86TI2\|D | 128185752 | 125164559 | -0.0974763 | 0.7113664 | 0.80553265 |
| sp\|O75083\|V | 732637611 | 757028565 | 0.00907776 | 0.7113664 | 0.80553265 |
| sp\|Q5VU97\|( | 205485.379 | 198609.389 | -0.0846314 | 0.70710065 | 0.80553265 |
| sp\|Q00059\|T | 971000.186 | 1002506.77 | 0.12846496 | 0.7113664 | 0.80553265 |
| sp\|Q06124\|F | 138578865 | 129106136 | -0.1958803 | 0.7113664 | 0.80553265 |
| sp\|Q69YN2\|( | 2777722.44 | 2721558.68 | 0.05944321 | 0.7113664 | 0.80553265 |
| sp\|Q9BTV6\|D | 4181187.17 | 4160950.03 | 0.13091383 | 0.7113664 | 0.80553265 |
| sp\|P20292\|A | 872826.566 | 1365788.5 | -2.0774613 | 0.7113664 | 0.80553265 |
| sp\|Q9NZU5\|I | 202463048 | 202521322 | -0.1787678 | 0.7113664 | 0.80553265 |
| sp\|Q13641\|T | 5209416.18 | 5261294.09 | -0.0487779 | 0.7113664 | 0.80553265 |
| sp\|A0A0C4DI | 52407931.8 | 53120427.8 | 0.14179589 | 0.7113664 | 0.80553265 |
| sp\|Q8NFH3\|I | 9563831.25 | 10137807.6 | 0.07444777 | 0.7113664 | 0.80553265 |
| sp\|A8MQB3\| | 463207.924 | 434661.958 | -0.3210399 | 0.7113664 | 0.80553265 |
| sp\|Q9NQ48\| | 14148867.6 | 14011805.6 | -0.0470231 | 0.7113664 | 0.80553265 |
| sp\|P48668\|K | 1341486959 | 820311923 | -0.2604444 | 0.7113664 | 0.80553265 |
| sp\|Q9Y6M1\|I | 15424514.5 | 16656564.9 | 0.05196308 | 0.7113664 | 0.80553265 |
| sp\|Q9NQT8\|I | 6649136.03 | 6570122.5 | -0.0280718 | 0.7113664 | 0.80553265 |
| sp\|P51114\|F | 7444933.14 | 8087675.56 | -0.0421322 | 0.7113664 | 0.80553265 |
| sp\|Q96AY3\|F | 39926954.2 | 38942827.2 | -0.1763964 | 0.7113664 | 0.80553265 |
| sp\|P52790\|H | 40627892.7 | 53287431 | 0.14777323 | 0.72816563 | 0.81895169 |
| sp\|Q9HD26\|( | 4799448.06 | 5348758.87 | 0.22493885 | 0.72816563 | 0.81895169 |
| sp\|P62993\|G | 68628857.4 | 67349266.5 | -0.0855954 | 0.72816563 | 0.81895169 |
| sp\|Q8WV41\| | 3277176.32 | 3783911.65 | 0.05992378 | 0.72816563 | 0.81895169 |
| sp\|O15305\|F | 37785673.2 | 38525390.1 | 0.01522032 | 0.72816563 | 0.81895169 |
| sp\|P35527\|K | 216042157 | 145188370 | 0.05278852 | 0.72816563 | 0.81895169 |
| sp\|Q9Y5F6\|F | 572836.587 | 558340.983 | 0.00284621 | 0.72816563 | 0.81895169 |
| sp\|P36980\|F | 78426859.9 | 83035686.7 | -0.0020455 | 0.72816563 | 0.81895169 |
| sp\|O43301\|H | 44730086.5 | 46022828.3 | -0.0735711 | 0.72816563 | 0.81895169 |
| sp\|Q13541\|4 | 5275620.9 | 4567390.23 | -0.0618197 | 0.72816563 | 0.81895169 |
| sp\|Q13428\|T | 41894980 | 41640059.3 | -0.0675302 | 0.72816563 | 0.81895169 |
| sp\|Q9NWH9\| | 120259.299 | 141377.302 | 0.01217162 | 0.72816563 | 0.81895169 |

| | | | | | |
|---|---|---|---|---|---|
| sp\|Q8IYM1\|S | 1804266.84 | 1809509.76 | 0.03296238 | 0.72816563 | 0.81895169 |
| sp\|Q08752\|F | 35925031.7 | 36517821.2 | 0.02376213 | 0.72816563 | 0.81895169 |
| sp\|Q9UM00\| | 4671384.22 | 4530103.97 | -0.1011029 | 0.72816563 | 0.81895169 |
| sp\|Q96SM3\|( | 7563863.57 | 7582836.42 | -0.0202304 | 0.72816563 | 0.81895169 |
| sp\|Q14204\|D | 59014490 | 64025464.8 | 0.06853622 | 0.72816563 | 0.81895169 |
| sp\|Q9UHQ9\| | 17844564.5 | 18225282.5 | -0.0015502 | 0.72816563 | 0.81895169 |
| sp\|Q9H098\|I | 6180814.16 | 6711633.33 | -0.0106739 | 0.72816563 | 0.81895169 |
| sp\|Q14573\|I | 7459575.88 | 7697371.17 | 0.01300447 | 0.72816563 | 0.81895169 |
| sp\|P20160\|C | 49480391.6 | 156545814 | -0.0661965 | 0.72816563 | 0.81895169 |
| sp\|Q9Y3P9\|F | 4828574.55 | 1270622.31 | -0.6404335 | 0.72816563 | 0.81895169 |
| sp\|Q9BYN8\|I | 2340375.98 | 2318076.28 | 0.05368691 | 0.72816563 | 0.81895169 |
| sp\|A0A0C4DI | 10681088.3 | 10729597.6 | 0.04659019 | 0.72816563 | 0.81895169 |
| sp\|Q9NPJ3\|A | 56432557 | 55361431 | -0.0029055 | 0.72816563 | 0.81895169 |
| sp\|Q92820\|( | 12366775.7 | 12920753.3 | -0.0034255 | 0.72816563 | 0.81895169 |
| sp\|O00161\|S | 3518181.36 | 3626603.6 | -0.0350288 | 0.72816563 | 0.81895169 |
| sp\|P0CAP2\|( | 769179.997 | 803069.116 | -0.1067188 | 0.72816563 | 0.81895169 |
| sp\|Q9UGP8\|. | 1213321.37 | 1253952.81 | -0.0575597 | 0.72816563 | 0.81895169 |
| sp\|Q96P48\|A | 175174.135 | 214177.017 | 1.26335408 | 0.74509226 | 0.8300134 |
| sp\|Q8TCJ2\|S | 2418129.66 | 2500219.84 | -0.015565 | 0.74509226 | 0.8300134 |
| sp\|P38571\|L | 2252803.18 | 2612814.85 | 0.07312623 | 0.74509226 | 0.8300134 |
| sp\|Q8NEL9\|D | 27570700.3 | 31652170.6 | 0.05482247 | 0.74509226 | 0.8300134 |
| sp\|Q13158\|F | 459685.286 | 446116.855 | -0.2134007 | 0.74509226 | 0.8300134 |
| sp\|Q8IWB7\|\ | 29412295.8 | 29241130.8 | -0.0051074 | 0.74509226 | 0.8300134 |
| sp\|P27338\|A | 88227118.1 | 85713869.4 | -0.1212094 | 0.74509226 | 0.8300134 |
| sp\|Q9Y2B0\|( | 79665387.7 | 78300130.1 | -0.0972864 | 0.74509226 | 0.8300134 |
| sp\|Q9NRL3\|S | 4746463.26 | 4768044.66 | 0.03083404 | 0.74509226 | 0.8300134 |
| sp\|Q9HBL8\|I | 9581996.32 | 8789596.32 | -0.2747636 | 0.74509226 | 0.8300134 |
| sp\|Q9Y2J8\|P | 596141.358 | 1296550.46 | -0.0538906 | 0.74509226 | 0.8300134 |
| sp\|Q7L9B9\|E | 800030.504 | 563745.19 | 0.31268847 | 0.74509226 | 0.8300134 |
| sp\|O75521\|E | 11731146.3 | 11276729.6 | -0.0463689 | 0.74509226 | 0.8300134 |
| sp\|Q13459\|N | 512246.77 | 645697.721 | 0.15136415 | 0.74509226 | 0.8300134 |
| sp\|Q15024\|E | 1948847.08 | 1882585.03 | -0.0882778 | 0.74509226 | 0.8300134 |
| sp\|O60516\|4 | 858974.017 | 844889.138 | -0.0750225 | 0.74509226 | 0.8300134 |
| sp\|Q92995\|U | 16939841.7 | 17433473.6 | 0.02379645 | 0.74509226 | 0.8300134 |
| sp\|GSTP1_HU | 1854903301 | 2034066708 | 0.01205439 | 0.74509226 | 0.8300134 |
| sp\|P14324\|F | 14502790.9 | 15151318.5 | -0.007003 | 0.74509226 | 0.8300134 |
| sp\|Q9BT78\|( | 37733355.8 | 40118650.5 | 0.04005079 | 0.74509226 | 0.8300134 |
| sp\|P20851\|C | 16789746.6 | 21242790.2 | -0.0842862 | 0.74509226 | 0.8300134 |
| sp\|P19256\|L | 5487974.43 | 5801257.81 | 0.05488672 | 0.74509226 | 0.8300134 |

| | | | | | |
|---|---|---|---|---|---|
| sp\|P14927\|Ç | 16373545.7 | 13958516.4 | -0.0163854 | 0.74509226 | 0.8300134 |
| sp\|Q5T6F2\|U | 2100179.67 | 2275159.24 | 0.00340949 | 0.74509226 | 0.8300134 |
| sp\|Q5VZ66\|J | 27570700.3 | 31652170.6 | 0.05482247 | 0.74509226 | 0.8300134 |
| sp\|Q32P28\|F | 4931450.48 | 5052673.58 | -0.1498524 | 0.74509226 | 0.8300134 |
| sp\|Q8N2S1\|I | 49410999.7 | 46866936.4 | -0.1694561 | 0.74509226 | 0.8300134 |
| sp\|Q9H4M9\| | 31804673.4 | 31017533.6 | -0.0347999 | 0.74509226 | 0.8300134 |
| sp\|Q96M83\|( | 2101217.05 | 2302493.68 | -0.2419178 | 0.74509226 | 0.8300134 |
| sp\|P62333\|P | 32985052.1 | 37219048.5 | 0.09152532 | 0.74509226 | 0.8300134 |
| sp\|O75351\|\ | 2645130.05 | 2780190.06 | 0.03214704 | 0.74509226 | 0.8300134 |
| sp\|P42768\|V | 571990.348 | 910799.106 | 0.04113137 | 0.74509226 | 0.8300134 |
| sp\|Q9NR19\|/ | 87061066.3 | 101601861 | -0.1271713 | 0.74509226 | 0.8300134 |
| sp\|P56524\|H | 2316653.73 | 2216209.19 | -0.2137867 | 0.74509226 | 0.8300134 |
| sp\|P52948\|N | 15705959.4 | 15883035 | -0.0387972 | 0.74509226 | 0.8300134 |
| sp\|Q9BYD1\|I | 4895290.17 | 5128162.34 | -0.0606613 | 0.74509226 | 0.8300134 |
| sp\|Q9BZV2\|S | 1364751.34 | 1356613.88 | -0.051108 | 0.74509226 | 0.8300134 |
| sp\|Q5JNZ3\|Z | 182745.625 | 174389.565 | -0.4040605 | 0.74509226 | 0.8300134 |
| sp\|Q9NQ29\| | 20157666.4 | 20978714.6 | 0.03785794 | 0.74509226 | 0.8300134 |
| sp\|Q96JM3\|( | 826835.739 | 581483.184 | -0.1565194 | 0.74509226 | 0.8300134 |
| sp\|Q9Y693\|L | 4177394.06 | 4205081.3 | 0.04885161 | 0.74509226 | 0.8300134 |
| sp\|P10646\|T | 1543378.69 | 1489688.61 | -0.0747107 | 0.762139 | 0.84157963 |
| sp\|Q9NZL9\|N | 73428675.1 | 77174921.2 | -0.0765679 | 0.762139 | 0.84157963 |
| sp\|Q14914\|F | 169148079 | 187508779 | -0.0227816 | 0.762139 | 0.84157963 |
| sp\|P02792\|F | 145913103 | 116030805 | -0.2683451 | 0.762139 | 0.84157963 |
| sp\|P02760\|A | 936489837 | 971472140 | 0.00038886 | 0.762139 | 0.84157963 |
| sp\|P10768\|E | 615282491 | 713216165 | 0.10088352 | 0.762139 | 0.84157963 |
| sp\|P24557\|T | 450338.115 | 552334.188 | 0.0420821 | 0.762139 | 0.84157963 |
| sp\|P09211\|G | 1505070866 | 1633026237 | -0.016322 | 0.762139 | 0.84157963 |
| sp\|P22061\|P | 141157805 | 137505672 | -0.2795346 | 0.762139 | 0.84157963 |
| sp\|Q99963\|S | 12829041.7 | 14682445.5 | -0.0672609 | 0.762139 | 0.84157963 |
| sp\|P01860\|I( | 3.1497E+10 | 3.2493E+10 | 0.06402717 | 0.762139 | 0.84157963 |
| sp\|Q9NTX5\|E | 53783759.2 | 59749230.1 | -0.0093172 | 0.762139 | 0.84157963 |
| sp\|P02776\|P | 6506858.18 | 12945622.2 | -0.3852619 | 0.762139 | 0.84157963 |
| sp\|P29536\|L | 32655341.1 | 31739551.4 | -0.071313 | 0.762139 | 0.84157963 |
| sp\|P04433\|K | 511374695 | 502102619 | 0.06484073 | 0.762139 | 0.84157963 |
| sp\|O95399\|U | 187681215 | 181521429 | -0.0524228 | 0.762139 | 0.84157963 |
| sp\|Q9Y281\|( | 443190608 | 451915825 | 0.04645787 | 0.762139 | 0.84157963 |
| sp\|Q9BZE9\|/ | 40708273.3 | 41215868.3 | -0.0026311 | 0.762139 | 0.84157963 |
| sp\|A0A1B0GU | 2014506.97 | 1947949.28 | -0.8176487 | 0.762139 | 0.84157963 |
| sp\|P49757\|N | 970702.596 | 1123867.79 | 0.06851447 | 0.762139 | 0.84157963 |

| | | | | | |
|---|---|---|---|---|---|
| sp\|Q96C23\|( | 54956073.3 | 64915917.9 | 0.02509855 | 0.762139 | 0.84157963 |
| sp\|O94979\|S | 22764683.9 | 23496571.9 | 0.05625272 | 0.762139 | 0.84157963 |
| sp\|Q9UNW9\| | 6606768.95 | 6994280.7 | 0.00117646 | 0.762139 | 0.84157963 |
| sp\|Q15800\|M | 649032.485 | 724873.489 | 0.07947882 | 0.762139 | 0.84157963 |
| sp\|Q9GZY8\|M | 2881939.59 | 2887601.32 | -0.041091 | 0.762139 | 0.84157963 |
| sp\|Q15365\|F | 107228356 | 112324509 | 0.01579021 | 0.762139 | 0.84157963 |
| sp\|P0CF74\|I( | 8693027128 | 9018661658 | 0.08848189 | 0.762139 | 0.84157963 |
| sp\|Q5RKV6\|E | 941883.813 | 1178255.26 | 0.09668728 | 0.762139 | 0.84157963 |
| sp\|Q9UBN7\| | 6301576.81 | 6289599.57 | 0.04190741 | 0.762139 | 0.84157963 |
| sp\|Q96A73\|F | 3299757.85 | 2759205 | -0.1467907 | 0.762139 | 0.84157963 |
| sp\|P01861\|I( | 1.9519E+10 | 1.994E+10 | 0.04880717 | 0.762139 | 0.84157963 |
| sp\|Q9Y3B7\|F | 25360721.1 | 24819343.7 | -0.0421526 | 0.762139 | 0.84157963 |
| sp\|Q02224\|( | 1152879.88 | 1181800.41 | 0.10016904 | 0.762139 | 0.84157963 |
| sp\|P28289\|T | 28736148.1 | 27967976.2 | -0.211207 | 0.762139 | 0.84157963 |
| sp\|Q9Y6W3\| | 1965178.56 | 1798128.23 | -0.1522201 | 0.762139 | 0.84157963 |
| sp\|Q9UNE7\|( | 7704947.33 | 7448340.72 | -0.0629459 | 0.762139 | 0.84157963 |
| sp\|P47914\|R | 40917274.9 | 38923606.7 | -0.0818798 | 0.762139 | 0.84157963 |
| sp\|Q8TEB1\|D | 768602.201 | 771189.563 | 0.04965964 | 0.762139 | 0.84157963 |
| sp\|Q8IZ73\|R | 854931.137 | 858892.043 | 0.04050375 | 0.77929825 | 0.85521434 |
| sp\|P28062\|P | 86597908.9 | 88913768.7 | -0.1795873 | 0.77929825 | 0.85521434 |
| sp\|Q8N7H5\| | 856954.798 | 1094497.94 | 0.21189488 | 0.77929825 | 0.85521434 |
| sp\|P07437\|T | 864996055 | 905841287 | -0.0060222 | 0.77929825 | 0.85521434 |
| sp\|P04180\|L | 9394679.79 | 9949714.08 | -0.0750065 | 0.77929825 | 0.85521434 |
| sp\|Q9HAV7\|( | 2068752.11 | 2059556.54 | -0.042992 | 0.77929825 | 0.85521434 |
| sp\|Q9Y2S6\|T | 6036047.53 | 6907325.24 | -0.0012541 | 0.77929825 | 0.85521434 |
| sp\|Q8TF27\|A | 2204909.72 | 10891449.5 | 0.36248442 | 0.77929825 | 0.85521434 |
| sp\|A0A0C4DI | 321682642 | 334486030 | 0.11030976 | 0.77929825 | 0.85521434 |
| sp\|P01704\|L | 32870464.4 | 32269289.8 | 0.1040027 | 0.77929825 | 0.85521434 |
| sp\|P14672\|G | 4262114.26 | 4628857.85 | 0.04342374 | 0.77929825 | 0.85521434 |
| sp\|Q9H2S9\|I | 302094.681 | 311837.799 | -0.0818522 | 0.77929825 | 0.85521434 |
| sp\|P04350\|T | 795681501 | 862446163 | 0.03889356 | 0.77929825 | 0.85521434 |
| sp\|Q9NS71\|( | 22190910.6 | 3211012.02 | -0.3459555 | 0.77929825 | 0.85521434 |
| sp\|Q8NFI3\|E | 16413989.7 | 16475253.2 | 0.00844094 | 0.77929825 | 0.85521434 |
| sp\|P36406\|T | 45319512.5 | 48962095.4 | -0.0081469 | 0.77929825 | 0.85521434 |
| sp\|Q9Y3A5\|S | 14947121.3 | 14878588.8 | -0.0422346 | 0.77929825 | 0.85521434 |
| sp\|P43003\|E | 392777.779 | 423658.412 | -0.0622226 | 0.77929825 | 0.85521434 |
| sp\|P06753\|T | 1637771945 | 1564734306 | -0.021478 | 0.77929825 | 0.85521434 |
| sp\|Q86V81\|T | 52836929.9 | 55498857.5 | -0.0469086 | 0.77929825 | 0.85521434 |
| sp\|O60637\|T | 4728883.47 | 3778501.89 | -0.0708567 | 0.77929825 | 0.85521434 |

| | | | | | |
|---|---|---|---|---|---|
| sp\|P20339\|R | 29835026.3 | 29626879.7 | -0.0259425 | 0.77929825 | 0.85521434 |
| sp\|Q6ZMQ8\| | 5062025.29 | 5784297.7 | 0.60541263 | 0.77929825 | 0.85521434 |
| sp\|Q16658\|F | 537268153 | 544830088 | -0.0363604 | 0.77929825 | 0.85521434 |
| sp\|Q9BTU6\|I | 6348783.05 | 6690146.3 | 0.07179419 | 0.77929825 | 0.85521434 |
| sp\|P20340\|R | 75542115.9 | 79305891.2 | 0.04766311 | 0.77929825 | 0.85521434 |
| sp\|Q9NZM1\| | 64835414.8 | 63611057.3 | -0.0232726 | 0.77929825 | 0.85521434 |
| sp\|P32121\|A | 6473124.08 | 6896632.18 | 0.0524651 | 0.79656226 | 0.86721921 |
| sp\|P56385\|A | 13373361.9 | 11839111.7 | -0.259418 | 0.79656226 | 0.86721921 |
| sp\|Q9Y5N5\|I | 1000378.27 | 1064990.07 | -0.130683 | 0.79656226 | 0.86721921 |
| sp\|P35998\|P | 35683227.6 | 37966783.1 | -0.0118106 | 0.79656226 | 0.86721921 |
| sp\|P37802\|T | 1163801991 | 1148771468 | -0.075504 | 0.79656226 | 0.86721921 |
| sp\|Q13813\|S | 757086638 | 788382366 | 0.04570885 | 0.79656226 | 0.86721921 |
| sp\|Q9BPY8\|H | 7019889.84 | 7365757.24 | -0.0570022 | 0.79656226 | 0.86721921 |
| sp\|P06280\|A | 8496771.56 | 8842265.35 | -0.1214589 | 0.79656226 | 0.86721921 |
| sp\|P53582\|M | 6750334.07 | 6723735.59 | 0.07805975 | 0.79656226 | 0.86721921 |
| sp\|Q9Y333\|L | 36046182.9 | 37335955.8 | 0.00043889 | 0.79656226 | 0.86721921 |
| sp\|P11217\|P | 1022379187 | 1044260762 | -0.022807 | 0.79656226 | 0.86721921 |
| sp\|Q92859\|N | 3088563.35 | 3108787.18 | -0.0519509 | 0.79656226 | 0.86721921 |
| sp\|Q6TDP4\|K | 18857806.1 | 16482531.9 | -0.0343269 | 0.79656226 | 0.86721921 |
| sp\|Q5TEU4\|N | 22096713.9 | 20912704.7 | -0.3701114 | 0.79656226 | 0.86721921 |
| sp\|A6NNZ2\|T | 398270428 | 423672914 | 0.04234115 | 0.79656226 | 0.86721921 |
| sp\|Q9H3K2\|G | 380150.898 | 354412.325 | -0.1940597 | 0.79656226 | 0.86721921 |
| sp\|Q8NBF2\|I | 45036951 | 46643783.7 | 0.03315534 | 0.79656226 | 0.86721921 |
| sp\|Q96SU4\|C | 1671535.78 | 1689419.58 | 0.02342607 | 0.79656226 | 0.86721921 |
| sp\|Q9BQ39\|I | 777637.496 | 847307.833 | -0.0426451 | 0.79656226 | 0.86721921 |
| sp\|P05362\|IC | 18437860.5 | 18564078.9 | -0.0246705 | 0.79656226 | 0.86721921 |
| sp\|Q13685\|A | 8653080.31 | 8843053.55 | -0.184777 | 0.79656226 | 0.86721921 |
| sp\|Q13951\|F | 966719.087 | 1125034.62 | 0.15611035 | 0.79656226 | 0.86721921 |
| sp\|Q7Z406\|M | 58435160.7 | 60600686.6 | 0.04460907 | 0.79656226 | 0.86721921 |
| sp\|Q9NYL9\|T | 60475249.6 | 61971759.5 | 0.00703686 | 0.79656226 | 0.86721921 |
| sp\|P09210\|G | 15200890.7 | 15207994 | -0.0876022 | 0.79656226 | 0.86721921 |
| sp\|P08493\|M | 2002498.92 | 2322393.08 | -0.3325302 | 0.79656226 | 0.86721921 |
| sp\|P08697\|A | 277240839 | 280364798 | -0.1773911 | 0.79656226 | 0.86721921 |
| sp\|P13796\|P | 249514671 | 278847153 | -0.1957695 | 0.79656226 | 0.86721921 |
| sp\|P11310\|A | 615394025 | 752877580 | -0.0695628 | 0.79656226 | 0.86721921 |
| sp\|O94804\|S | 5645917.87 | 6115785.41 | 0.07579557 | 0.79656226 | 0.86721921 |
| sp\|Q9UGM5\| | 3104634.29 | 3334748.87 | -0.0753199 | 0.79656226 | 0.86721921 |
| sp\|Q9BYD3\|I | 11487966.2 | 11812215.7 | 0.01300701 | 0.79656226 | 0.86721921 |
| sp\|Q9BS92\|N | 90646662.8 | 89242420 | -0.0652263 | 0.79656226 | 0.86721921 |

| | | | | | |
|---|---|---|---|---|---|
| sp\|Q38SD2\|[illegible] | 3752882.54 | 3927258.03 | -0.4790245 | 0.79656226 | 0.86721921 |
| sp\|O00231\|[illegible] | 22024415.7 | 23921655.7 | 0.06439202 | 0.79656226 | 0.86721921 |
| sp\|O75531\|[illegible] | 64404538.4 | 66538471.4 | -0.0629033 | 0.81392329 | 0.87795412 |
| sp\|O00767\|[illegible] | 140344.97 | 149246.418 | 0.85976172 | 0.81392329 | 0.87795412 |
| sp\|Q86VH2\|[illegible] | 665839.558 | 680841.844 | -0.0534874 | 0.81392329 | 0.87795412 |
| sp\|P07359\|[illegible] | 5969676.33 | 7520858.95 | -0.0573395 | 0.81392329 | 0.87795412 |
| sp\|Q13630\|[illegible] | 115559242 | 123721503 | 0.02614797 | 0.81392329 | 0.87795412 |
| sp\|P35606\|[illegible] | 45310104.7 | 45457884.6 | -0.0171648 | 0.81392329 | 0.87795412 |
| sp\|P80217\|[illegible] | 10612388.8 | 10482072.5 | 0.00792541 | 0.81392329 | 0.87795412 |
| sp\|P01602\|[illegible] | 143646909 | 138042577 | 0.04243398 | 0.81392329 | 0.87795412 |
| sp\|Q92604\|[illegible] | 540704.616 | 472835.665 | 0.40936402 | 0.81392329 | 0.87795412 |
| sp\|Q92508\|[illegible] | 1698207.46 | 1915307.32 | -0.3309933 | 0.81392329 | 0.87795412 |
| sp\|O60885\|[illegible] | 559825.928 | 588447.633 | -1.5690409 | 0.81392329 | 0.87795412 |
| sp\|Q9ULV4\|[illegible] | 157251757 | 165987219 | 0.01923509 | 0.81392329 | 0.87795412 |
| sp\|Q9UK45\|[illegible] | 31093362.8 | 32064025.1 | -0.0767358 | 0.81392329 | 0.87795412 |
| sp\|Q99653\|[illegible] | 21233234.3 | 19876229.4 | -0.09891 | 0.81392329 | 0.87795412 |
| sp\|Q8NCA5\| | 3939006.67 | 3798932.46 | -0.2034029 | 0.81392329 | 0.87795412 |
| sp\|P36954\|[illegible] | 1369799.34 | 1364561.75 | -0.0502981 | 0.81392329 | 0.87795412 |
| sp\|Q8TBE9\|[illegible] | 901800.938 | 939595.076 | 0.09080986 | 0.81392329 | 0.87795412 |
| sp\|P36957\|[illegible] | 272316246 | 241807190 | -0.0627724 | 0.81392329 | 0.87795412 |
| sp\|Q00587\|[illegible] | 9401812.34 | 8736378.08 | -0.1365326 | 0.81392329 | 0.87795412 |
| sp\|O43681\|[illegible] | 11201319.6 | 13027097.3 | 0.04196457 | 0.81392329 | 0.87795412 |
| sp\|O43432\|[illegible] | 4253043.97 | 2007550.27 | -0.0157386 | 0.81392329 | 0.87795412 |
| sp\|P08319\|A | 1915428250 | 1749000751 | -0.0577825 | 0.81392329 | 0.87795412 |
| sp\|Q68CZ2\|[illegible] | 16890096.7 | 17129479.3 | -0.0235008 | 0.81392329 | 0.87795412 |
| sp\|Q66GS9\|[illegible] | 66570941.7 | 68774249.6 | -0.0692551 | 0.81392329 | 0.87795412 |
| sp\|Q9BW83\| | 7685945.26 | 7918895.19 | 0.03801017 | 0.81392329 | 0.87795412 |
| sp\|MYG_HUM | 96614668.3 | 92500920.1 | -0.4116831 | 0.81392329 | 0.87795412 |
| sp\|Q9UBF2\|[illegible] | 1195809.27 | 1269167.9 | -0.0181711 | 0.81392329 | 0.87795412 |
| sp\|P14151\|L | 11883394 | 11725644.7 | 0.02208698 | 0.81392329 | 0.87795412 |
| sp\|Q5JUK3\|[illegible] | 1083148.43 | 1061713.68 | -0.2483642 | 0.81392329 | 0.87795412 |
| sp\|P13598\|[illegible] | 2308469 | 2414878.9 | -0.032333 | 0.81392329 | 0.87795412 |
| sp\|Q3ZCM7\|[illegible] | 474731818 | 512410480 | 0.04690318 | 0.81392329 | 0.87795412 |
| sp\|Q6UW78\| | 466301.822 | 474658.797 | 0.11950476 | 0.81392329 | 0.87795412 |
| sp\|Q6UN15\|[illegible] | 1184247.48 | 1376236.85 | 0.0811006 | 0.81392329 | 0.87795412 |
| sp\|O00214\|L | 17908801.8 | 14755272 | -0.1414563 | 0.81392329 | 0.87795412 |
| sp\|Q15637\|[illegible] | 27600792.8 | 28050603 | 0.02949823 | 0.81392329 | 0.87795412 |
| sp\|P09104\|E | 649189891 | 644366785 | -0.0210259 | 0.81392329 | 0.87795412 |
| sp\|Q16719\|[illegible] | 24060361.6 | 24624889.2 | -0.0627992 | 0.81392329 | 0.87795412 |

| | | | | | |
|---|---|---|---|---|---|
| sp\|Q9Y3S2\|Z | 156400.252 | 170322.88 | 0.59789517 | 0.81392329 | 0.87795412 |
| sp\|Q9Y217\|M | 990001.332 | 1039706.87 | 0.1119397 | 0.81392329 | 0.87795412 |
| sp\|Q96EK6\|G | 20069999.5 | 20470204.6 | 0.02504031 | 0.81392329 | 0.87795412 |
| sp\|A0A0C4DI | 63549340.6 | 64402815.8 | 0.05937548 | 0.81392329 | 0.87795412 |
| sp\|O95989\|N | 20827923.9 | 20458431.8 | -0.1131406 | 0.83137332 | 0.88957872 |
| sp\|Q96L93\|K | 3806636.9 | 4168203.49 | -0.022187 | 0.83137332 | 0.88957872 |
| sp\|P50995\|A | 115273926 | 111896534 | -0.0941646 | 0.83137332 | 0.88957872 |
| sp\|O94760\|D | 59926756.8 | 59998562 | -0.075163 | 0.83137332 | 0.88957872 |
| sp\|O00159\|M | 153126177 | 159347233 | -0.0924629 | 0.83137332 | 0.88957872 |
| sp\|P09234\|R | 887301.012 | 1028681.23 | -0.0269923 | 0.83137332 | 0.88957872 |
| sp\|Q4G0F5\|V | 22593125.5 | 22706232.1 | 0.00732332 | 0.83137332 | 0.88957872 |
| sp\|Q5JRA6\|T | 1746353.08 | 1735472.36 | -0.0227257 | 0.83137332 | 0.88957872 |
| sp\|Q9GZX9\|T | 311931.895 | 311653.094 | 0.01272602 | 0.83137332 | 0.88957872 |
| sp\|Q9NYL2\|M | 216459.401 | 204063.201 | -0.0279367 | 0.83137332 | 0.88957872 |
| sp\|P22234\|P | 73937477.2 | 82693852.3 | 0.0012688 | 0.83137332 | 0.88957872 |
| sp\|Q9BSJ2\|G | 143705.818 | 144299.899 | 0.37367895 | 0.83137332 | 0.88957872 |
| sp\|Q9UI30\|T | 13807393.6 | 13848718 | 0.00856215 | 0.83137332 | 0.88957872 |
| sp\|P07099\|H | 407463802 | 415089085 | -0.0243554 | 0.83137332 | 0.88957872 |
| sp\|Q7L5N7\|P | 294134.002 | 334634.074 | -0.03264 | 0.83137332 | 0.88957872 |
| sp\|Q9NUV9\| | 4937933.54 | 4898290.08 | -0.0689199 | 0.83137332 | 0.88957872 |
| sp\|Q9UKA9\|P | 8144061.35 | 7513375.64 | 0.26544786 | 0.83137332 | 0.88957872 |
| sp\|P04920\|B | 532584.42 | 642027.44 | 0.0756188 | 0.83137332 | 0.88957872 |
| sp\|A6NES4\|M | 1217682.79 | 1217019.29 | -0.6328356 | 0.83137332 | 0.88957872 |
| sp\|Q05682\|C | 504976152 | 508364170 | -0.0223044 | 0.83137332 | 0.88957872 |
| sp\|Q8NCW5\| | 220414911 | 227917162 | -0.0519625 | 0.83137332 | 0.88957872 |
| sp\|Q08257\|C | 339464364 | 416258380 | 0.04921333 | 0.83137332 | 0.88957872 |
| sp\|Q06203\|P | 12810987.2 | 13084901.8 | -0.0310688 | 0.83137332 | 0.88957872 |
| sp\|Q06587\|R | 1782952.06 | 1816468.81 | -0.0014991 | 0.83137332 | 0.88957872 |
| sp\|P23497\|S | 33344216.7 | 34952296.1 | 0.0323783 | 0.83137332 | 0.88957872 |
| sp\|Q14242\|S | 4389391.09 | 4612820.93 | -0.1541221 | 0.83137332 | 0.88957872 |
| sp\|Q13509\|T | 623650172 | 659206269 | 0.01085476 | 0.83137332 | 0.88957872 |
| sp\|Q86VB7\|C | 295382940 | 292331409 | -0.0998968 | 0.83137332 | 0.88957872 |
| sp\|P20962\|P | 110429033 | 111671841 | -0.0440732 | 0.83137332 | 0.88957872 |
| sp\|Q8TDQ7\| | 103088024 | 113615867 | -0.0193377 | 0.83137332 | 0.88957872 |
| sp\|P02686\|M | 1779944.28 | 1740256.71 | -0.3204363 | 0.83137332 | 0.88957872 |
| sp\|P37837\|T | 1306243724 | 1330456881 | 0.00380464 | 0.83137332 | 0.88957872 |
| sp\|Q96LJ7\|D | 3816932.64 | 3944622.17 | -0.1017894 | 0.83137332 | 0.88957872 |
| sp\|Q8TF65\|G | 2061519.69 | 2265045.42 | -0.0510847 | 0.83137332 | 0.88957872 |
| sp\|Q96PZ0\|P | 274701.182 | 383742.211 | -1.1045335 | 0.83137332 | 0.88957872 |

| | | | | | |
|---|---|---|---|---|---|
| sp\|Q8WXD9\| | 1135457.81 | 1184416.83 | 0.03718024 | 0.83137332 | 0.88957872 |
| sp\|O94819\|K | 11975031.2 | 11544670.4 | -0.1059015 | 0.84890418 | 0.90350214 |
| sp\|Q8IWE4\|D | 549545.688 | 495309.995 | -0.0597081 | 0.84890418 | 0.90350214 |
| sp\|P27694\|R | 29944214.1 | 31563462.6 | 0.01654708 | 0.84890418 | 0.90350214 |
| sp\|Q13325\|I | 854824.525 | 904279.1 | 0.00891126 | 0.84890418 | 0.90350214 |
| sp\|Q13232\|N | 47734469.3 | 46510734.2 | -0.0800425 | 0.84890418 | 0.90350214 |
| sp\|Q6P2E9\|E | 836972.67 | 1252806.24 | 0.21510678 | 0.84890418 | 0.90350214 |
| sp\|Q969L2\|N | 2960032.73 | 3137563.03 | -0.0223816 | 0.84890418 | 0.90350214 |
| sp\|Q15651\|H | 604537.237 | 752705.066 | -0.0582699 | 0.84890418 | 0.90350214 |
| sp\|P04632\|C | 89494311.2 | 91407691.4 | -0.0281081 | 0.84890418 | 0.90350214 |
| sp\|Q96KR1\|Z | 9034916.11 | 8853681.12 | -0.0285362 | 0.84890418 | 0.90350214 |
| sp\|Q96EM0\|T | 66564565.1 | 66012200.1 | -0.082229 | 0.84890418 | 0.90350214 |
| sp\|A0A075B6 | 70745958.7 | 66824503.9 | 0.08825487 | 0.84890418 | 0.90350214 |
| sp\|Q01432\|A | 7627470.29 | 8107320.22 | -0.0112725 | 0.84890418 | 0.90350214 |
| sp\|Q9UKZ9\|P | 1165547.55 | 1206655.2 | -0.1639092 | 0.84890418 | 0.90350214 |
| sp\|Q00534\|C | 59076644.8 | 112071776 | -0.0921277 | 0.84890418 | 0.90350214 |
| sp\|P37173\|T | 1006208.31 | 1090565.78 | -0.6581183 | 0.84890418 | 0.90350214 |
| sp\|P01743\|H | 150575835 | 150900502 | 0.01446635 | 0.84890418 | 0.90350214 |
| sp\|O43181\|N | 927848.958 | 836392.02 | 0.04170387 | 0.84890418 | 0.90350214 |
| sp\|Q8NFH4\| | 12759091.5 | 13102774.5 | 0.04695528 | 0.84890418 | 0.90350214 |
| sp\|Q8NHH9\| | 911140.817 | 988282.682 | -0.0626727 | 0.84890418 | 0.90350214 |
| sp\|P08195\|4 | 14399675.1 | 14367131.6 | 0.01320632 | 0.84890418 | 0.90350214 |
| sp\|Q5T1M5\|F | 2045505.65 | 2041823.33 | 0.06968075 | 0.84890418 | 0.90350214 |
| sp\|Q03468\|E | 14413799.2 | 14961146.2 | -0.2357068 | 0.84890418 | 0.90350214 |
| sp\|O75886\|S | 989240.798 | 996202.698 | -0.0562544 | 0.84890418 | 0.90350214 |
| sp\|Q16401\|F | 8294359.21 | 8967052.37 | -0.00326 | 0.86650778 | 0.91533587 |
| sp\|Q9UGP4\| | 972074.033 | 1090526.2 | -0.0333833 | 0.86650778 | 0.91533587 |
| sp\|P14314\|G | 363059696 | 357327294 | -0.0522063 | 0.86650778 | 0.91533587 |
| sp\|Q9NXH9\|T | 19238995.7 | 19119317 | -0.0793701 | 0.86650778 | 0.91533587 |
| sp\|Q13107\|U | 5985138.89 | 6485441.67 | -0.1546039 | 0.86650778 | 0.91533587 |
| sp\|P31930\|Q | 105997814 | 105453932 | 0.01750714 | 0.86650778 | 0.91533587 |
| sp\|Q12929\|E | 36259799.3 | 35188748.9 | -0.0979342 | 0.86650778 | 0.91533587 |
| sp\|P29558\|R | 15093201.4 | 15218053.7 | -0.0498628 | 0.86650778 | 0.91533587 |
| sp\|O95400\|C | 1668083.88 | 1548463.31 | -0.1444202 | 0.86650778 | 0.91533587 |
| sp\|P21741\|M | 4411442.19 | 4338957.94 | -0.0048223 | 0.86650778 | 0.91533587 |
| sp\|Q14331\|F | 2967455.94 | 3032929.33 | -0.1139099 | 0.86650778 | 0.91533587 |
| sp\|O15235\|F | 14275136.3 | 14411880.6 | -0.0273456 | 0.86650778 | 0.91533587 |
| sp\|Q7L576\|C | 32919350.9 | 32153283.6 | -0.0636358 | 0.86650778 | 0.91533587 |
| sp\|Q99961\|S | 29735026.8 | 29991445.2 | 0.02189335 | 0.86650778 | 0.91533587 |

| | | | | | |
|---|---|---|---|---|---|
| sp\|Q53HC9\| | 5919613.19 | 5867313.34 | 0.01924607 | 0.86650778 | 0.91533587 |
| sp\|O94832\|M | 2270346.02 | 2712156.01 | -0.0210802 | 0.86650778 | 0.91533587 |
| sp\|O60684\|I | 1533403.63 | 1910426.66 | 0.08208892 | 0.86650778 | 0.91533587 |
| sp\|Q9HD34\| | 1220465.93 | 1438943.92 | -0.0054935 | 0.86650778 | 0.91533587 |
| sp\|O75367\|H | 9887198.19 | 12869012.7 | 0.11879885 | 0.86650778 | 0.91533587 |
| sp\|P51809\|V | 90337.4498 | 82090.5874 | -0.1751598 | 0.86650778 | 0.91533587 |
| sp\|Q9Y399\|F | 1299505 | 1405043.43 | 0.02450961 | 0.86650778 | 0.91533587 |
| sp\|Q8TDN6\|I | 1721619.96 | 1665621.68 | -0.2207802 | 0.86650778 | 0.91533587 |
| sp\|O00757\|F | 23938593.4 | 23876369.2 | -0.2172352 | 0.86650778 | 0.91533587 |
| sp\|O00584\|F | 9598750.57 | 10014696.6 | 0.00964644 | 0.86650778 | 0.91533587 |
| sp\|P09467\|F | 163776584 | 160629284 | -0.2921804 | 0.86650778 | 0.91533587 |
| sp\|Q15631\|T | 68556088.9 | 70996436.7 | 0.00441988 | 0.86650778 | 0.91533587 |
| sp\|O00178\|C | 10102186.7 | 11452561.8 | 0.09739563 | 0.86650778 | 0.91533587 |
| sp\|P16383\|G | 1479058.21 | 1551007.22 | 0.00236295 | 0.86650778 | 0.91533587 |
| sp\|Q9BW61\| | 305024.295 | 360098.71 | -0.0553862 | 0.86650778 | 0.91533587 |
| sp\|P12931\|S | 9219557.52 | 9566056.92 | -0.0121244 | 0.86650778 | 0.91533587 |
| sp\|Q9UBS5\|( | 827969.017 | 884276.952 | 0.04089965 | 0.86650778 | 0.91533587 |
| sp\|Q969E8\|T | 1596546.16 | 1547040.88 | -0.0686691 | 0.86650778 | 0.91533587 |
| sp\|O43805\|S | 150033.002 | 157251.18 | -2.068725 | 0.86438359 | 0.91533587 |
| sp\|Q92956\|T | 735832.751 | 765270.921 | -0.1470786 | 0.86650778 | 0.91533587 |
| sp\|P10412\|H | 304932878 | 311788896 | -0.0347796 | 0.88417576 | 0.92544373 |
| sp\|P10153\|R | 5372599.67 | 8058238.2 | -0.0468176 | 0.88417576 | 0.92544373 |
| sp\|Q8N2G8\| | 2311353.19 | 3178181.97 | 0.20139619 | 0.88417576 | 0.92544373 |
| sp\|Q9H3N1\| | 4645176.9 | 4464067.58 | -0.1248414 | 0.88417576 | 0.92544373 |
| sp\|Q13153\|F | 6598170.62 | 7156215.93 | 0.08133103 | 0.88417576 | 0.92544373 |
| sp\|Q7Z7H5\|T | 3113719.3 | 3116049.14 | -0.1195372 | 0.88417576 | 0.92544373 |
| sp\|O95671\|A | 3901576.64 | 3667820.74 | -0.0691041 | 0.88417576 | 0.92544373 |
| sp\|P05387\|R | 158454655 | 157357099 | -0.0410407 | 0.88417576 | 0.92544373 |
| sp\|Q13439\|C | 7728418.81 | 7381938.76 | -0.064879 | 0.88417576 | 0.92544373 |
| sp\|Q9Y295\|D | 832400.574 | 789153.919 | -0.1115107 | 0.88417576 | 0.92544373 |
| sp\|Q86YW5\| | 3186952.56 | 3190182.74 | -0.0550499 | 0.88417576 | 0.92544373 |
| sp\|O75396\|S | 14435218.2 | 14677021.7 | -0.0845761 | 0.88417576 | 0.92544373 |
| sp\|O75665\|C | 39834583.6 | 40058785 | -0.0048031 | 0.88417576 | 0.92544373 |
| sp\|Q99797\|M | 45521847.5 | 51312672.5 | -0.1122074 | 0.88417576 | 0.92544373 |
| sp\|O00194\|F | 6465901.37 | 8231500.44 | -0.0612147 | 0.88417576 | 0.92544373 |
| sp\|O75822\|E | 2964608.87 | 2842669.34 | -0.0637953 | 0.88417576 | 0.92544373 |
| sp\|Q9NVF9\|E | 6142297.6 | 5455827.32 | -0.1115849 | 0.88417576 | 0.92544373 |
| sp\|P08571\|C | 39581005.6 | 40846156.4 | -0.1657276 | 0.88417576 | 0.92544373 |
| sp\|Q6NY19\|I | 775418.47 | 645998.454 | -0.1284653 | 0.88417576 | 0.92544373 |

| | | | | | |
|---|---|---|---|---|---|
| sp\|Q5T6V5\| | 10560748.4 | 11110723.3 | 0.02796355 | 0.88417576 | 0.92544373 |
| sp\|Q5U651\| | 736205.421 | 761857.626 | 0.00368065 | 0.88417576 | 0.92544373 |
| sp\|P00480\| | 1004428.56 | 1313442.6 | 0.08723495 | 0.88417576 | 0.92544373 |
| sp\|A0A0C4D | 18261583.2 | 18175856.1 | 0.08532232 | 0.88417576 | 0.92544373 |
| sp\|Q96HD1\| | 5036870.04 | 5465648.57 | -0.0745227 | 0.88417576 | 0.92544373 |
| sp\|A0A075B | 3985411.49 | 3383810.12 | 0.05540108 | 0.88417576 | 0.92544373 |
| sp\|Q69YN4\| | 15745293.3 | 17001655.2 | 0.01965507 | 0.88417576 | 0.92544373 |
| sp\|Q6BCY4\| | 29721331.3 | 30199018.4 | -0.0700117 | 0.88417576 | 0.92544373 |
| sp\|Q9UBS8\| | 672979.104 | 665412.488 | -0.067344 | 0.88417576 | 0.92544373 |
| sp\|Q9P2E2\| | 3534326.52 | 3548915.48 | 0.00288487 | 0.88417576 | 0.92544373 |
| sp\|Q6Y288\| | 2648290.85 | 2671175.64 | -0.0365442 | 0.88417576 | 0.92544373 |
| sp\|P15814\| | 6133085.87 | 6617352.4 | -0.0254431 | 0.88417576 | 0.92544373 |
| sp\|Q16186\| | 6381462.94 | 6345227.97 | -0.0513739 | 0.88417576 | 0.92544373 |
| sp\|Q6V1X1\| | 5515729.92 | 5471333.41 | -0.0210364 | 0.88417576 | 0.92544373 |
| sp\|Q6ZVK8\| | 14624512.6 | 14957557.2 | 0.02943644 | 0.88417576 | 0.92544373 |
| sp\|Q8WZA9\| | 230991.931 | 249593.222 | 0.01279662 | 0.88417576 | 0.92544373 |
| sp\|Q9Y639\| | 4951561.26 | 4813276.67 | -0.1138303 | 0.88417576 | 0.92544373 |
| sp\|Q9Y678\| | 35393719.5 | 36134248.5 | 0.01969402 | 0.88417576 | 0.92544373 |
| sp\|O75391\| | 980890.216 | 912451.692 | -0.0518691 | 0.88417576 | 0.92544373 |
| sp\|P40763\|S | 293503.493 | 314887.46 | 0.52888823 | 0.88417576 | 0.92544373 |
| sp\|Q9H788\| | 205293.494 | 217483.202 | -0.0763152 | 0.88417576 | 0.92544373 |
| sp\|P82914\|R | 4052771.17 | 5265855.54 | -0.0193888 | 0.88417576 | 0.92544373 |
| sp\|P48039\| | 1835777.27 | 1787817.49 | -0.1235828 | 0.88417576 | 0.92544373 |
| sp\|P35219\| | 1915293.64 | 1976925.46 | -0.0497096 | 0.90189965 | 0.938265 |
| sp\|Q9H3S7\| | 118342234 | 129662071 | 0.98537344 | 0.90189965 | 0.938265 |
| sp\|P30711\|G | 110092283 | 112381355 | -0.0178486 | 0.90189965 | 0.938265 |
| sp\|P29353\|S | 1463743.67 | 1470455.11 | -0.0446851 | 0.90189965 | 0.938265 |
| sp\|Q9Y3E5\| | 553086.694 | 563738.444 | -0.0255755 | 0.90189965 | 0.938265 |
| sp\|Q9UBM7\| | 2302875.77 | 2357489.98 | -0.0045376 | 0.90189965 | 0.938265 |
| sp\|P16152\| | 1055641857 | 1128271500 | 0.0733682 | 0.90189965 | 0.938265 |
| sp\|Q9NZ08\| | 271164385 | 261397316 | -0.1646721 | 0.90189965 | 0.938265 |
| sp\|P49750\|Y | 685207.498 | 772818.814 | 0.07050494 | 0.90189965 | 0.938265 |
| sp\|Q9BU02\| | 5460431.33 | 5464668.09 | -0.0102433 | 0.90189965 | 0.938265 |
| sp\|Q9UBQ5\| | 895145.331 | 892502.398 | -0.0575999 | 0.90189965 | 0.938265 |
| sp\|P50542\|P | 1395684.33 | 1392820.57 | -0.0400195 | 0.90189965 | 0.938265 |
| sp\|O75923\| | 3922098.63 | 4207032.21 | 0.04661004 | 0.90189965 | 0.938265 |
| sp\|Q86WQ0\| | 59811613.8 | 64434659.6 | -0.0146196 | 0.90189965 | 0.938265 |
| sp\|P05062\| | 26470710.6 | 25278284.9 | -0.0894348 | 0.90189965 | 0.938265 |
| sp\|Q9UPT8\| | 7687439.46 | 7800724.24 | -0.0464628 | 0.90189965 | 0.938265 |

| | | | | | |
|---|---|---|---|---|---|
| sp\|Q7L622\|( | 3360883.06 | 3563996.63 | -0.0431463 | 0.90189965 | 0.938265 |
| sp\|Q7Z460\|( | 1268509.78 | 1693262.26 | 0.14229059 | 0.90189965 | 0.938265 |
| sp\|Q9Y3D9\|F | 3045498.89 | 3039805.13 | -0.2057617 | 0.90189965 | 0.938265 |
| sp\|Q7Z6B0\|( | 563552.989 | 508585.559 | -0.0597204 | 0.90189965 | 0.938265 |
| sp\|P68032\|A | 1.4707E+10 | 1.6266E+10 | 0.04993494 | 0.90189965 | 0.938265 |
| sp\|P01877\|I( | 6463479265 | 5952496353 | -0.0037991 | 0.90189965 | 0.938265 |
| sp\|Q9ULI3\|H | 243678.75 | 249256.423 | -0.042011 | 0.90189965 | 0.938265 |
| sp\|Q9ULH1\|/ | 10256361.8 | 14507045.6 | 0.03056464 | 0.90189965 | 0.938265 |
| sp\|P01591\|I( | 155300434 | 149890991 | -0.0718283 | 0.90189965 | 0.938265 |
| sp\|Q8WUA7\| | 3336436.96 | 3382366 | 0.01700096 | 0.90189965 | 0.938265 |
| sp\|Q9Y2R0\|( | 5762060.14 | 13799974 | -0.2526221 | 0.90189965 | 0.938265 |
| sp\|A0A0C4DI | 3893490.93 | 3424031.35 | -0.0859389 | 0.90189965 | 0.938265 |
| sp\|A0A075B6 | 13569163 | 14098245.3 | 0.04505459 | 0.9196711 | 0.94934429 |
| sp\|A0A075B6 | 78969239.1 | 78663612.5 | 0.12145924 | 0.9196711 | 0.94934429 |
| sp\|Q96FS4\|S | 4896287.64 | 5218372.16 | 0.03670723 | 0.9196711 | 0.94934429 |
| sp\|P01742\|H | 141055441 | 133106640 | -0.0065717 | 0.9196711 | 0.94934429 |
| sp\|P01619\|K | 560571886 | 560542184 | 0.06794118 | 0.9196711 | 0.94934429 |
| sp\|P07948\|L | 62667993.9 | 67383804 | 0.04041508 | 0.9196711 | 0.94934429 |
| sp\|P05198\|IF | 29851990.9 | 29386083.2 | -0.0228631 | 0.9196711 | 0.94934429 |
| sp\|P04004\|V | 426966225 | 467848939 | -0.0690667 | 0.9196711 | 0.94934429 |
| sp\|Q9NZM3\| | 1129933.3 | 1109877.9 | 0.50758517 | 0.9196711 | 0.94934429 |
| sp\|Q9NZN5\|/ | 295306.906 | 298607.05 | 0.00465009 | 0.9196711 | 0.94934429 |
| sp\|P35251\|R | 57464951.2 | 54408609.1 | -0.1782426 | 0.9196711 | 0.94934429 |
| sp\|P24928\|R | 14228274 | 15275950.1 | 0.01534493 | 0.9196711 | 0.94934429 |
| sp\|P11308\|E | 145936.25 | 157178.402 | -0.5056749 | 0.9196711 | 0.94934429 |
| sp\|Q9P265\|[ | 5265726.23 | 5443525.69 | 0.00911494 | 0.9196711 | 0.94934429 |
| sp\|P16298\|P | 54057761.6 | 54608367.4 | 0.02147046 | 0.9196711 | 0.94934429 |
| sp\|Q9Y224\|F | 53942669 | 53285390.7 | 0.00427856 | 0.9196711 | 0.94934429 |
| sp\|P30566\|P | 22921944.8 | 30557038.5 | 0.1280036 | 0.9196711 | 0.94934429 |
| sp\|P31997\|C | 1460703.67 | 1850809.63 | 0.05813997 | 0.9196711 | 0.94934429 |
| sp\|Q06136\|K | 6992663.48 | 7874997.16 | 0.00921367 | 0.9196711 | 0.94934429 |
| sp\|Q9Y3C1\|I | 3490927.51 | 3033965.55 | -0.0278841 | 0.9196711 | 0.94934429 |
| sp\|P61244\|M | 3144341.98 | 1202140.22 | -0.2045816 | 0.9196711 | 0.94934429 |
| sp\|P51797\|C | 5962243.58 | 8087585.1 | 0.20559868 | 0.9196711 | 0.94934429 |
| sp\|P58335\|A | 493871.589 | 470649.152 | -0.0467116 | 0.9196711 | 0.94934429 |
| sp\|P57105\|S | 745746.272 | 753362.059 | -0.1093515 | 0.9196711 | 0.94934429 |
| sp\|Q99836\|I | 8154045.42 | 7324461.55 | -0.4328039 | 0.9196711 | 0.94934429 |
| sp\|Q8NI08\|I | 210099.919 | 220210.021 | -0.1076051 | 0.9196711 | 0.94934429 |
| sp\|Q8NC51\| | 53149121.3 | 56872734.1 | -0.0565492 | 0.9196711 | 0.94934429 |

| | | | | | |
|---|---|---|---|---|---|
| sp\|Q6P4I2\|W | 553149.803 | 549524.455 | -0.10925 | 0.9196711 | 0.94934429 |
| sp\|Q66K74\|M | 16657630.9 | 16550066.5 | -0.0043516 | 0.9196711 | 0.94934429 |
| sp\|Q16799\|F | 6077309.92 | 5919299.54 | -0.1181315 | 0.9196711 | 0.94934429 |
| sp\|Q16626\|M | 10922832.3 | 11201789.6 | 0.04515481 | 0.9196711 | 0.94934429 |
| sp\|Q9C0C9\| | 46136962.5 | 68416250.3 | 0.13845143 | 0.9196711 | 0.94934429 |
| sp\|Q96CV9\|( | 55357988.1 | 55455581.1 | 0.00689945 | 0.9196711 | 0.94934429 |
| sp\|Q96A35\|F | 3946072.48 | 3945212.08 | -0.0742133 | 0.9196711 | 0.94934429 |
| sp\|Q92636\|F | 365592.048 | 405586.154 | -0.065798 | 0.9196711 | 0.94934429 |
| sp\|Q8TBN0\|F | 862954.494 | 751812.949 | -0.2257842 | 0.9196711 | 0.94934429 |
| sp\|P35637\|F | 46696570.1 | 50835034.3 | -0.0359896 | 0.93748149 | 0.96296525 |
| sp\|Q9NP58\|/ | 46895671.1 | 50773029.4 | -0.0412244 | 0.93748149 | 0.96296525 |
| sp\|Q9NQE9\| | 16198445.6 | 16760735.4 | 0.04170972 | 0.93748149 | 0.96296525 |
| sp\|P58546\|M | 16407382.5 | 17036331 | 0.05045521 | 0.93748149 | 0.96296525 |
| sp\|Q9UK41\|V | 369426.639 | 353325.163 | 0.00107138 | 0.93748149 | 0.96296525 |
| sp\|P50502\|F | 358303921 | 346433539 | -0.0885328 | 0.93748149 | 0.96296525 |
| sp\|P04264\|K | 537382102 | 381430055 | -0.0923345 | 0.93748149 | 0.96296525 |
| sp\|Q9ULZ3\|/ | 13993921.9 | 14375106.8 | 0.02233834 | 0.93748149 | 0.96296525 |
| sp\|O75629\|( | 24174960.9 | 24260230.7 | -0.0892375 | 0.93748149 | 0.96296525 |
| sp\|Q8N8E3\|( | 25165227.1 | 25210594.9 | 0.8249703 | 0.93748149 | 0.96296525 |
| sp\|Q8IYN0\|Z | 275018.115 | 267167.133 | -0.5798385 | 0.93748149 | 0.96296525 |
| sp\|Q8NBX0\|S | 17026907.9 | 18610759.4 | 0.00472061 | 0.93748149 | 0.96296525 |
| sp\|Q16832\|D | 2773721.29 | 2733922.02 | -0.0734303 | 0.93748149 | 0.96296525 |
| sp\|Q5TFE4\|N | 85587013.9 | 85490205.8 | -0.0168735 | 0.93748149 | 0.96296525 |
| sp\|Q5TC12\|/ | 2122570.85 | 1997981.26 | -0.3705808 | 0.93748149 | 0.96296525 |
| sp\|Q8IYB7\|D | 3426518.97 | 3558518.14 | -0.0311567 | 0.93748149 | 0.96296525 |
| sp\|Q05315\|L | 8949562.04 | 13462666.5 | -0.0427743 | 0.93748149 | 0.96296525 |
| sp\|Q08209\|F | 68577874.6 | 69719159.5 | 0.0161462 | 0.93748149 | 0.96296525 |
| sp\|Q9H0K1\|S | 439204.242 | 496319.707 | 0.06220708 | 0.93748149 | 0.96296525 |
| sp\|P68133\|A | 1.457E+10 | 1.6304E+10 | 0.05616721 | 0.93748149 | 0.96296525 |
| sp\|A0A0C4DI | 144896709 | 127789911 | -0.0789094 | 0.93748149 | 0.96296525 |
| sp\|P17693\|H | 36032372.9 | 34756667.1 | -0.0271161 | 0.93748149 | 0.96296525 |
| sp\|P0DTE1\|H | 43465162.4 | 42451897.9 | 0.0208811 | 0.93748149 | 0.96296525 |
| sp\|P29350\|P | 22798033.7 | 25390055.3 | 0.0409709 | 0.95532217 | 0.9737874 |
| sp\|Q16082\|H | 5849967.06 | 6212499.7 | 0.12001947 | 0.95532217 | 0.9737874 |
| sp\|Q04760\|L | 325075640 | 327123827 | -0.0304129 | 0.95532217 | 0.9737874 |
| sp\|Q9H0U6\| | 5986695.88 | 6177154.19 | 0.00345758 | 0.95532217 | 0.9737874 |
| sp\|Q15642\|( | 8873797.85 | 8990323.53 | -0.0041652 | 0.95532217 | 0.9737874 |
| sp\|P60891\|P | 88735364.1 | 100715442 | 0.0480644 | 0.95532217 | 0.9737874 |
| sp\|P50454\|S | 83643089.4 | 84768180.2 | 0.0001537 | 0.95532217 | 0.9737874 |

| | | | | | |
|---|---|---|---|---|---|
| sp\|P17480\|U | 1082021.13 | 1266850.26 | -0.0849484 | 0.95532217 | 0.9737874 |
| sp\|Q2TAL8\|Q | 3306493.36 | 3557810.86 | 0.01454195 | 0.95532217 | 0.9737874 |
| sp\|Q6PCE3\|I | 37453341.6 | 42641603.4 | 0.0410653 | 0.95532217 | 0.9737874 |
| sp\|Q6ZV73\|F | 1320848.57 | 1473605.78 | -0.1141115 | 0.95532217 | 0.9737874 |
| sp\|Q6EEV4\|C | 146321.335 | 133770.444 | -2.3705285 | 0.95532217 | 0.9737874 |
| sp\|Q5EBL8\|F | 3435174.89 | 3827707.25 | -0.137389 | 0.95532217 | 0.9737874 |
| sp\|Q93088\|E | 11791160.3 | 8576663.3 | -0.0123012 | 0.95532217 | 0.9737874 |
| sp\|A0A087W | 5276469.11 | 4938357.13 | -0.1054421 | 0.95532217 | 0.9737874 |
| sp\|Q99489\|C | 1902635.18 | 1928365.62 | 0.05876819 | 0.95532217 | 0.9737874 |
| sp\|A0A0C4DI | 55964162.9 | 54080705.4 | 0.0244708 | 0.95532217 | 0.9737874 |
| sp\|Q9BVL4\|S | 10667241.4 | 10959968.2 | 0.03220798 | 0.95532217 | 0.9737874 |
| sp\|Q9GZQ8\| | 11931421.2 | 11719147.2 | -0.0358296 | 0.95532217 | 0.9737874 |
| sp\|P62312\|L | 48002209.2 | 49230709.1 | -0.0845273 | 0.95532217 | 0.9737874 |
| sp\|P83111\|L | 6277045.49 | 6685014.86 | 0.03939542 | 0.95532217 | 0.9737874 |
| sp\|P57772\|S | 1100148.48 | 1185438.85 | -0.1006229 | 0.95532217 | 0.9737874 |
| sp\|P30626\|S | 36419713 | 40548365.9 | 0.05252279 | 0.95532217 | 0.9737874 |
| sp\|P13798\|A | 476402560 | 504615317 | -0.0504628 | 0.95532217 | 0.9737874 |
| sp\|P24298\|A | 294843627 | 301708814 | -0.0325574 | 0.95532217 | 0.9737874 |
| sp\|Q9UG63\|A | 750832.558 | 757990.83 | 0.02175885 | 0.95532217 | 0.9737874 |
| sp\|P09601\|H | 70853743.4 | 70470490.9 | -0.0218158 | 0.95532217 | 0.9737874 |
| sp\|Q9UGV2\| | 1574056.77 | 2381064.69 | 0.03974894 | 0.95532217 | 0.9737874 |
| sp\|P50213\|ID | 50436431.8 | 49719460.9 | -0.0174942 | 0.95532217 | 0.9737874 |
| sp\|P62736\|A | 1.4719E+10 | 1.6165E+10 | 0.03442092 | 0.95532217 | 0.9737874 |
| sp\|P05164\|P | 64197573.8 | 181876205 | 0.12761015 | 0.95532217 | 0.9737874 |
| sp\|P00533\|E | 17285676.1 | 17324501.3 | -0.0059268 | 0.95532217 | 0.9737874 |
| sp\|P06732\|K | 122824912 | 95118019.1 | -0.2780141 | 0.95532217 | 0.9737874 |
| sp\|A0A075B6 | 1974245.31 | 2277684.67 | 0.00815106 | 0.95532217 | 0.9737874 |
| sp\|Q96HR8\|I | 188435.448 | 178752.955 | -0.0760792 | 0.95532217 | 0.9737874 |
| sp\|A0A0B4J1 | 125626425 | 127462105 | 0.01938575 | 0.95532217 | 0.9737874 |
| sp\|Q15643\|T | 9025024.17 | 8733196.39 | -0.1093782 | 0.97318462 | 0.98384517 |
| sp\|Q9Y4D1\|D | 2159806.15 | 2160237.39 | -0.1152304 | 0.97318462 | 0.98384517 |
| sp\|Q2M1P5\|I | 26833700.7 | 26283664.1 | -0.7498474 | 0.97318462 | 0.98384517 |
| sp\|Q86WR0\| | 2468667.02 | 2734911.75 | -0.091672 | 0.97318462 | 0.98384517 |
| sp\|Q13418\|I | 33172372.2 | 33658859.9 | -0.0502858 | 0.97318462 | 0.98384517 |
| sp\|P63027\|V | 8159059.03 | 8235639.12 | -0.0082577 | 0.97318462 | 0.98384517 |
| sp\|Q08378\|C | 9061442.18 | 8349014.8 | -0.0794896 | 0.97318462 | 0.98384517 |
| sp\|Q13409\|D | 50668943.5 | 51137656.6 | 0.01214742 | 0.97318462 | 0.98384517 |
| sp\|Q13155\|A | 19499194.6 | 19142766.3 | -0.0675965 | 0.97318462 | 0.98384517 |
| sp\|P54922\|A | 230519367 | 226368372 | -0.0284777 | 0.97318462 | 0.98384517 |

| | | | | | |
|---|---|---|---|---|---|
| sp\|Q9C040\|T | 2059207.79 | 2058449.31 | 0.11970649 | 0.97318462 | 0.98384517 |
| sp\|Q14498\|F | 3076756.87 | 3301161.86 | 0.03017826 | 0.97318462 | 0.98384517 |
| sp\|P40394\|A | 920653039 | 976338965 | -0.189424 | 0.97318462 | 0.98384517 |
| sp\|P48147\|P | 114886759 | 118749874 | -0.1179477 | 0.97318462 | 0.98384517 |
| sp\|P46527\|C | 3816705.91 | 3528940.16 | -0.0354466 | 0.97318462 | 0.98384517 |
| sp\|P28300\|L | 21172470.3 | 23101357.1 | 0.20703598 | 0.97318462 | 0.98384517 |
| sp\|P29622\|K | 86868533.2 | 88579602.3 | -0.089156 | 0.97318462 | 0.98384517 |
| sp\|P22735\|T | 13455471.9 | 13847834.2 | 0.01943676 | 0.97318462 | 0.98384517 |
| sp\|O75832\|F | 37554025.1 | 36082521 | -0.2827822 | 0.97318462 | 0.98384517 |
| sp\|P08514\|IT | 23533857.2 | 41639886.9 | 0.32216716 | 0.97318462 | 0.98384517 |
| sp\|P01876\|I( | 1.0197E+10 | 9507686017 | 0.01419204 | 0.97318462 | 0.98384517 |
| sp\|P01880\|I( | 80434932 | 97322739.7 | -0.0702646 | 0.97318462 | 0.98384517 |
| sp\|P05413\|F | 66099051.4 | 68249638.6 | 0.03610179 | 0.97318462 | 0.98384517 |
| sp\|P04839\|C | 8742106.77 | 10036423.2 | 0.04877826 | 0.97318462 | 0.98384517 |
| sp\|P04216\|T | 11227415.1 | 11055792.8 | -0.0643164 | 0.97318462 | 0.98384517 |
| sp\|P11498\|P | 456740306 | 494312005 | -0.1603409 | 0.97318462 | 0.98384517 |
| sp\|P49790\|N | 2305559.47 | 2200970.77 | -0.0684561 | 0.97318462 | 0.98384517 |
| sp\|Q03013\|( | 93990119.8 | 104006282 | -0.1447393 | 0.97318462 | 0.98384517 |
| sp\|Q9NSK0\|I | 9938891.28 | 10624492.5 | -0.0300972 | 0.97318462 | 0.98384517 |
| sp\|P42226\|S | 249574.843 | 259490.949 | -0.5648551 | 0.97318462 | 0.98384517 |
| sp\|Q96TC7\|F | 1752243.49 | 1738226.42 | -0.0802632 | 0.97318462 | 0.98384517 |
| sp\|Q99952\|F | 435680.416 | 444018.482 | -0.6184419 | 0.97318462 | 0.98384517 |
| sp\|Q8IYQ7\|T | 1829559.63 | 2033553.19 | -0.1267976 | 0.97318462 | 0.98384517 |
| sp\|Q16822\|F | 282158090 | 263946979 | -0.1749559 | 0.97318462 | 0.98384517 |
| sp\|Q6UW02\| | 150416.651 | 883766.049 | -0.4914059 | 0.97318462 | 0.98384517 |
| sp\|Q9BSH5\|I | 34529046.9 | 33622945.8 | 0.00069777 | 0.97318462 | 0.98384517 |
| sp\|Q92783\|S | 9080997.42 | 8910058.63 | -0.0277169 | 0.97318462 | 0.98384517 |
| sp\|Q9BRK3\|I | 3117796.41 | 3121018.55 | 0.03399071 | 0.97318462 | 0.98384517 |
| sp\|Q86YR5\|( | 24127531.6 | 23433009.5 | -0.9064817 | 0.97318462 | 0.98384517 |
| sp\|P08567\|P | 14454512.5 | 16772643.9 | -0.0233771 | 0.99106011 | 0.99416753 |
| sp\|P05156\|C | 345250393 | 348824323 | -0.0264234 | 0.99106011 | 0.99416753 |
| sp\|O95486\|S | 468799.258 | 541549.891 | -0.0915089 | 0.99106011 | 0.99416753 |
| sp\|Q16851\|U | 802766329 | 807956232 | -0.1220508 | 0.99106011 | 0.99416753 |
| sp\|P52434\|R | 2036747.91 | 2215868.81 | 0.0422292 | 0.99106011 | 0.99416753 |
| sp\|Q9NPF4\|( | 2192403.79 | 2227669.96 | 0.02102005 | 0.99106011 | 0.99416753 |
| sp\|Q07954\|L | 535917517 | 531083350 | -0.0397921 | 0.99106011 | 0.99416753 |
| sp\|P52788\|S | 162495757 | 162451812 | -0.0403772 | 0.99106011 | 0.99416753 |
| sp\|Q9NQH7\| | 47601192.1 | 46397798.2 | -0.0312927 | 0.99106011 | 0.99416753 |
| sp\|Q15173\|2 | 22428912.9 | 34317552.6 | -0.1401198 | 0.99106011 | 0.99416753 |

| | | | | | |
|---|---|---|---|---|---|
| sp\|P42566\|E | 25917501.4 | 26205976.7 | 0.02495686 | 0.99106011 | 0.99416753 |
| sp\|P43897\|E | 2768326.72 | 3112992.36 | 0.08762055 | 0.99106011 | 0.99416753 |
| sp\|Q9NQW7\| | 133354736 | 137093354 | -0.0738518 | 0.99106011 | 0.99416753 |
| sp\|Q9NQG5\| | 6679692 | 6658776.66 | -0.0004452 | 0.99106011 | 0.99416753 |
| sp\|P49419\|A | 285430600 | 290151191 | -0.0183569 | 0.99106011 | 0.99416753 |
| sp\|Q96EY7\|F | 148113.517 | 163422.622 | -1.1829068 | 0.99106011 | 0.99416753 |
| sp\|P00813\|A | 15288991.4 | 15512431.9 | -0.0323526 | 0.99106011 | 0.99416753 |
| sp\|P01766\|H | 378874249 | 372382282 | -0.0294301 | 0.99106011 | 0.99416753 |
| sp\|P21757\|M | 10202971.5 | 13024119.6 | 0.01916815 | 0.99106011 | 0.99416753 |
| sp\|Q9UBI1\|C | 309854.953 | 329091.021 | -0.7710247 | 0.99106011 | 0.99416753 |
| sp\|P10746\|H | 3233442.15 | 3969834.66 | 0.14439321 | 0.99106011 | 0.99416753 |
| sp\|P0DOY3\|I | 9158394876 | 9481754656 | 0.05009963 | 0.99106011 | 0.99416753 |
| sp\|P22033\|M | 109186906 | 111865334 | -0.0105925 | 0.99106011 | 0.99416753 |
| sp\|P98198\|A | 3231844 | 3637674.54 | 0.04218073 | 0.99106011 | 0.99416753 |
| sp\|Q13438\|C | 20136241 | 20602770.1 | 0.06346481 | 0.99106011 | 0.99416753 |
| sp\|Q15363\|T | 287215.29 | 315972.299 | -0.0654412 | 0.99106011 | 0.99416753 |
| sp\|Q96CP2\|I | 1732694.27 | 1770519 | -0.1002075 | 0.99106011 | 0.99416753 |
| sp\|Q7L014\|D | 3912708.6 | 4428239.48 | -0.0481071 | 0.99106011 | 0.99416753 |
| sp\|Q6UXG8\| | 3577686.59 | 3622682.04 | -0.063579 | 0.99106011 | 0.99416753 |
| sp\|Q6VY07\|F | 92488042.1 | 92273720.6 | -0.0428819 | 0.99106011 | 0.99416753 |
| sp\|Q86YZ3\|H | 4583697.43 | 3545769.73 | 0.08266888 | 0.99106011 | 0.99416753 |
| sp\|Q6FHJ7\|S | 3034852.78 | 3166754.81 | -0.0111499 | 0.99106011 | 0.99416753 |
| sp\|Q3ZCW2\| | 2066328.5 | 2402536.21 | -0.0243449 | 0.99106011 | 0.99416753 |
| sp\|Q6ULP2\|A | 312900.207 | 288625.965 | -0.019288 | 0.99106011 | 0.99416753 |
| sp\|Q9Y4Z0\|L | 58949764.5 | 60372020.5 | -0.121294 | 0.99106011 | 0.99416753 |
| sp\|Q16891\|M | 10643261.6 | 10540021 | -0.0363971 | 0.99106011 | 0.99416753 |
| sp\|Q9Y478\|A | 35073112.4 | 38452094.1 | -0.2851436 | 0.99106011 | 0.99416753 |
| sp\|O95714\|H | 3662566.3 | 4272944.21 | 0.03227495 | 1 | 1 |
| sp\|P04430\|K | 22570692.4 | 22008231.7 | 0.04769698 | 1 | 1 |
| sp\|Q9NV96\|C | 702072.748 | 720689.087 | -0.1970028 | 1 | 1 |
| sp\|O60613\|S | 5767575.6 | 5973170.29 | 0.01934167 | 1 | 1 |
| sp\|O94992\|H | 1173428.45 | 1128427.94 | -0.0280293 | 1 | 1 |
| sp\|Q9Y3E2\|E | 1152272.67 | 1197249.76 | -0.1104 | 1 | 1 |
| sp\|Q9H8G2\| | 6687332.63 | 6819068.91 | -0.0044043 | 1 | 1 |
| sp\|O43674\|N | 1180395.19 | 1239629.15 | -0.0452395 | 1 | 1 |
| sp\|O60869\|E | 6396468.22 | 7374765.84 | 0.0888923 | 1 | 1 |
| sp\|O75312\|Z | 7133198.36 | 7073238.72 | -0.035965 | 1 | 1 |
| sp\|O00330\|C | 35101013.1 | 34690119.7 | -0.0059582 | 1 | 1 |
| sp\|A6NHX0\|C | 600551.429 | 559216.159 | -0.2587443 | 1 | 1 |

| | | | | | |
|---|---|---|---|---|---|
| sp\|Q8N5N7\| | 4142212.73 | 4426513.62 | -0.0066848 | 1 | 1 |
| sp\|Q8NDH2\| | 1073987.5 | 1055950.39 | -0.0322574 | 1 | 1 |
| sp\|Q969P0\|I | 1013803.91 | 1030504.01 | 0.0816307 | 1 | 1 |
