## Supplemental Tables 1 and 2 for "Proteomic Profiling of Human Omental and Subcutaneous Adipose Tissue in Individuals with a Broad Range of BMI"

Supplementary Table S1: Sample order after randomization for proteomic analysis

| Randomized Sample Order | Multiwell plate location | Sample Name |
|---|---|---|
| **1** | A1 | 51-3528-SQ-1-lean |
| **2** | A2 | 51-3525-SQ-1-obese |
| **3** | A3 | 51-3521-OM-2-lean |
| **4** | A4 | 51-3523-SQ-4-obese |
| **5** | A5 | 51-3518-SQ-2-obese |
| **6** | A6 | 51-3530-SQ-2-lean |
| **7** | A7 | 51-3505-SQ-2-obese |
| **8** | A8 | 51-3506-OM-2-obese |
| **9** | A9 | 51-3507-OM-1-obese |
| **10** | A10 | 51-3532-OM-1-obese |
| **11** | A11 | 51-3512-OM-2-obese |
| **12** | A12 | 51-3525-OM-1-obese |
| **13** | B1 | 51-3506-SQ-4-obese |
| **14** | B2 | 51-3529-OM-4-obese |
| **15** | B3 | 51-3531-OM-1-obese |
| **16** | B4 | 51-3519-OM-2-obese |
| **17** | B5 | 51-3522-OM-2-obese |
| **18** | B6 | 51-3503-SQ-1-lean |
| **19** | B7 | 51-3523-OM-4-obese |
| **20** | B8 | 51-3533-SQ-2-lean |
| **21** | B9 | 51-3514-OM-4-obese |
| **22** | B10 | 51-3508-OM-3-obese |
| **23** | B11 | 51-3502-OM-1-obese |
| **24** | B12 | 51-3501-OM-4-obese |
| **25** | C1 | 51-3520-OM-4-obese |
| **26** | C2 | 51-3502-SQ-2-obese |
| **27** | C3 | 51-3527-OM-2-obese |
| **28** | C4 | 51-3503-OM-2-lean |
| **29** | C5 | 51-3501-SQ-2-obese |
| **30** | C6 | 51-3504-OM-2-lean |
| **31** | C7 | 51-3516-OM-1-obese |
| **32** | C8 | 51-3529-SQ-1-obese |
| **33** | C9 | 51-3527-SQ-1-obese |
| **34** | C10 | 51-3517-SQ-4-obese |
| **35** | C11 | 51-3522-SQ-1-obese |
| **36** | C12 | 51-3505-OM-4-obese |
| **37** | D1 | 51-3524-OM-2-obese |

| | | |
|---|---|---|
| **38** | D2 | 51-3528-OM-2-lean |
| **39** | D3 | 51-3521-SQ-3-lean |
| **40** | D4 | 51-3516-SQ-4-obese |
| **41** | D5 | 51-3532-SQ-2-obese |
| **42** | D6 | 51-3509-SQ-1-obese |
| **43** | D7 | 51-3515-OM-1-lean |
| **44** | D8 | 51-3517-OM-4-obese |
| **45** | D9 | 51-3514-SQ-2-obese |
| **46** | D10 | 51-3512-SQ-1-obese |
| **47** | D11 | 51-3510-OM-2-obese |
| **48** | D12 | 51-3515-SQ-4-lean |
| **49** | E1 | 51-3519-SQ-4-obese |
| **50** | E2 | 51-3504-SQ-4-lean |
| **51** | E3 | 51-3520-SQ-4-obese |
| **52** | E4 | 51-3531-SQ-4-obese |
| **53** | E5 | 51-3508-SQ-1-obese |
| **54** | E6 | 51-3518-OM-1-obese |
| **55** | E7 | 51-3507-SQ-4-obese |
| **56** | E8 | 51-3509-OM-1-obese |
| **57** | E9 | 51-3513-SQ-4-lean |
| **58** | E10 | 51-3524-SQ-1-obese |
| **59** | E11 | 51-3510-SQ-1-obese |

Supplementary Table S2: Peptides and protein detections identified by EncyclopeDIA per LC-MS/MS injection at 1% FDR.

| Sample Name | Peptide Detections | Protein Detections |
|---|---|---|
| **51-3514-SQ-2-obese** | 24255 | 2765 |
| **51-3520-OM-4-obese** | 32073 | 3425 |
| **51-3516-OM-1-obese** | 35161 | 3688 |
| **51-3504-SQ-4-lean** | 21271 | 2249 |
| **51-3515-OM-1-lean** | 28730 | 3106 |
| **51-3518-OM-1-obese** | 38395 | 3928 |
| **51-3508-OM-3-obese** | 36646 | 3712 |
| **51-3530-SQ-2-lean** | 36187 | 3666 |
| **51-3528-OM-2-lean** | 33440 | 3567 |
| **51-3524-OM-2-obese** | 33666 | 3577 |
| **51-3502-OM-1-obese** | 30655 | 3173 |
| **51-3522-SQ-1-obese** | 21053 | 2417 |
| **51-3507-SQ-4-obese** | 32941 | 3472 |

| | | |
|---|---|---|
| **51-3514-OM-4-obese** | 34900 | 3638 |
| **51-3512-SQ-1-obese** | 32488 | 3372 |
| **51-3525-SQ-1-obese** | 34311 | 3506 |
| **51-3523-OM-4-obese** | 39164 | 3950 |
| **51-3505-SQ-2-obese** | 28322 | 2918 |
| **51-3518-SQ-2-obese** | 27844 | 2990 |
| **51-3501-OM-4-obese** | 37334 | 3909 |
| **51-3502-SQ-2-obese** | 30563 | 3141 |
| **51-3506-SQ-4-obese** | 29911 | 3252 |
| **51-3533-SQ-2-lean** | 30202 | 3238 |
| **51-3521-SQ-3-lean** | 27543 | 2936 |
| **51-3529-SQ-1-obese** | 29387 | 3089 |
| **51-3504-OM-2-lean** | 36188 | 3754 |
| **51-3503-OM-2-lean** | 21786 | 2636 |
| **51-3509-OM-1-obese** | 34615 | 3506 |
| **51-3509-SQ-1-obese** | 24008 | 2829 |
| **51-3510-OM-2-obese** | 34742 | 3647 |
| **51-3519-SQ-4-obese** | 24532 | 2741 |
| **51-3528-SQ-1-lean** | 24610 | 2777 |
| **51-3507-OM-1-obese** | 31876 | 3336 |
| **51-3506-OM-2-obese** | 39807 | 4051 |
| **51-3531-SQ-4-obese** | 25815 | 2835 |
| **51-3515-SQ-4-lean** | 25658 | 2710 |
| **51-3532-SQ-2-obese** | 30567 | 3270 |
| **51-3531-OM-1-obese** | 36788 | 3782 |
| **51-3523-SQ-4-obese** | 36071 | 3707 |
| **51-3519-OM-2-obese** | 33084 | 3438 |
| **51-3521-OM-2-lean** | 32336 | 3445 |
| **51-3517-OM-4-obese** | 24005 | 2625 |
| **51-3525-OM-1-obese** | 36146 | 3806 |
| **51-3508-SQ-1-obese** | 33840 | 3424 |
| **51-3505-OM-4-obese** | 35603 | 3632 |
| **51-3516-SQ-4-obese** | 24329 | 2808 |
| **51-3524-SQ-1-obese** | 27226 | 2876 |
| **51-3520-SQ-4-obese** | 30417 | 3171 |
| **51-3517-SQ-4-obese** | 27201 | 2996 |
| **51-3522-OM-2-obese** | 25648 | 2754 |
| **51-3503-SQ-1-lean** | 30523 | 3239 |
| **51-3532-OM-1-obese** | 35429 | 3614 |
| **51-3513-SQ-4-lean** | 28238 | 3000 |
| **51-3501-SQ-2-obese** | 36336 | 3858 |

| | | |
|---|---|---|
| **51-3529-OM-4-obese** | 38457 | 3946 |
| **51-3512-OM-2-obese** | 36408 | 3807 |
| **51-3510-SQ-1-obese** | 20741 | 2403 |
| **51-3527-SQ-1-obese** | 31498 | 3231 |
| **51-3527-OM-2-obese** | 36038 | 3761 |
